# An integrated, transcript-resolution atlas of prostate cancer progression unmasks associations with oncogenic pathways

**DOI:** 10.64898/2026.09.18.752617

**Authors:** Jichang Zhang, Claudio Lorenzi, Daniela Bossi, Simone Nicastri, Matteo Corpetti, Nicolo Formaggio, Jacopo Sgrignani, Andrea Cavalli, Marco Bolis, Jean-Philippe Theurillat

## Abstract

Most genes produce multiple non-coding and coding RNA transcripts with potentially divergent biological functions, yet their contribution to cancer remains largely unexplored. Here, we built an integrated, transcript-resolution RNA-sequencing atlas of prostate cancer spanning 1,140 clinical samples from 10 independent studies, covering the full range of disease progression from normal tissue to primary, hormone-sensitive, and castration-resistant disease. Cross-study integration removes technical batch effects while retaining the biological heterogeneity that distinguishes these disease states, a property we validate against long-read sequencing and leverage to place newly incorporated external cohorts within this disease-progression context. We describe transcript patterns of oncogenic and tumor-suppressive long non-coding RNAs along disease progression and identify 41 protein-coding genes with significant isoform switches in cancer-relevant pathways, including WNT2B, RNF43, RELA, BRCA1, ROCK2, and TRIM11. Finally, we provide the Prostate Cancer Atlas (www.prostatecanceratlas.org), an interactive web resource that lets researchers contextualize their own transcriptomic data against this atlas.

**Highlights:** ● Data reprocessing removes batch effects and preserves prostate cancer heterogeneity

● Atlas maps lncRNA transcript changes along disease progression

● Isoform switches reveal domain-specific roles in tumor suppressors and oncogenes

● A public web tool contextualizes new external data within the atlas

**eTOC Blurb:** *Zhang et al. build an integrated, transcript-resolution atlas of prostate cancer from 14 source datasets, retaining ten paired-end datasets for the transcript-level atlas after excluding single-end libraries. Preserving heterogeneity while resolving individual transcripts uncovers isoform-specific oncogenic pathway associations and lets researchers map new external data onto the atlas*.

## Introduction

Gene transcripts mediate between the genome and biological effectors, and most genes produce multiple variants through alternative transcription start/stop sites and exon splicing. A paradigm for the functional consequences of isoform choice is the developmental and cancer-associated switch of pyruvate kinase from the adult (PKM1) to the embryonal (PKM2) isoform, which shifts enzyme activity from constitutive glycolytic flux to the allosterically regulated state that supports biosynthesis and the Warburg effect in proliferating cells^1,2^.

In prostate cancer, alternative isoforms of the androgen receptor (AR) — the disease’s key lineage-specific oncogene and therapeutic target — drive resistance to AR-directed therapies^3,4^; the ligand-binding-domain-truncated AR-V7, for instance, restores oncogenic signaling alongside full-length AR in castration-resistant disease^5,6^. Ancestry-specific splice variants have similarly been linked to differences in tumorigenic potential^7^. Yet the broader functional relevance of alternative transcripts to oncogenic pathways and disease progression remains largely uncharacterized.

Building such an association map requires integrating independent cohorts without erasing true biological heterogeneity. Recent efforts manage this only at gene or cell-type resolution — across prostate cancer cohorts^8^, in single-cell prostate cancer atlases^9^, and in an analogous breast cancer atlas^15^ — and none link isoform expression to pathway activity.

Conversely, efforts that reach transcript resolution have not solved cross-study harmonization. Pan-cancer splicing atlases built on single-consortium data catalog splice events but stop short of full transcript quantification^10–12^, while true isoform quantification elsewhere is either aggregated back to the gene level ^13^ or confined to a single cohort^14^. No resource unites transcript resolution, cross-study harmonization, and full disease-stage coverage while preserving heterogeneity — precisely what’s needed to mine isoform-pathway associations systematically.

Here, we take advantage of a deeply diverse, integrated RNA sequencing atlas of prostate cancer that we built at transcript resolution across the entire range of disease progression, from normal tissue through primary and metastatic disease to neuroendocrine and double-negative castration-resistant states, extending our previously published gene-level atlas^16^. We show that harmonizing independently generated cohorts removes technical batch effects while retaining the biological heterogeneity that distinguishes disease states, a property we validate both against long-read sequencing and by the atlas’s ability to correctly position newly integrated external cohorts of primary and hormone-sensitive disease. We then use this resource to systematically correlate transcript-specific expression with oncogenic pathway activity along disease progression, unmasking a thus far underappreciated role of alternative transcripts of key oncogenes and tumor suppressors in prostate cancer.

## Results

### A reprocessed, harmonized atlas resolves disease stages while removing cross-study batch effects

Gene-count tables released by individual bulk RNA-sequencing studies retained pronounced cross-study differences even after standard batch correction, exceeding the biological variance observed within any single study (Fig. S1A–D). We therefore reprocessed raw sequencing data from 14 bulk RNA-sequencing studies spanning normal prostate tissue, primary disease, and castration-resistant prostate cancer (CRPC) using a uniform computational pipeline^17^ (STAR alignment^18^ and Salmon-based gene and transcript quantification^19^; Fig. S1F–G).

This reprocessing produced an integrated atlas in which samples from the same disease stage overlapped closely while distinct disease stages separated along a continuous, inferred trajectory from normal tissue through primary disease to CRPC, without requiring further batch correction, in agreement with our previous gene-level atlas^16^ (Fig. 1A; Fig. S1F–L). Based on gene expression, CRPC samples were further stratified into androgen receptor-positive (ARPC), neuroendocrine (NEPC)^20^, and double-negative (DNPC)^21^ tumors, the latter lacking both AR and neuroendocrine markers (Fig. 1A; Fig. S1M, N). Restricting this analysis to protein-coding or long non-coding RNA (lncRNA) genes alone reproduced the overall entropy seen for the full gene expression atlas, but with a higher adjusted Rand index for lncRNAs, suggesting that lncRNA expression tracks disease stage even more closely than expression of the gene expression atlas as a whole (Fig. S2A–E).

**Figure 1.**
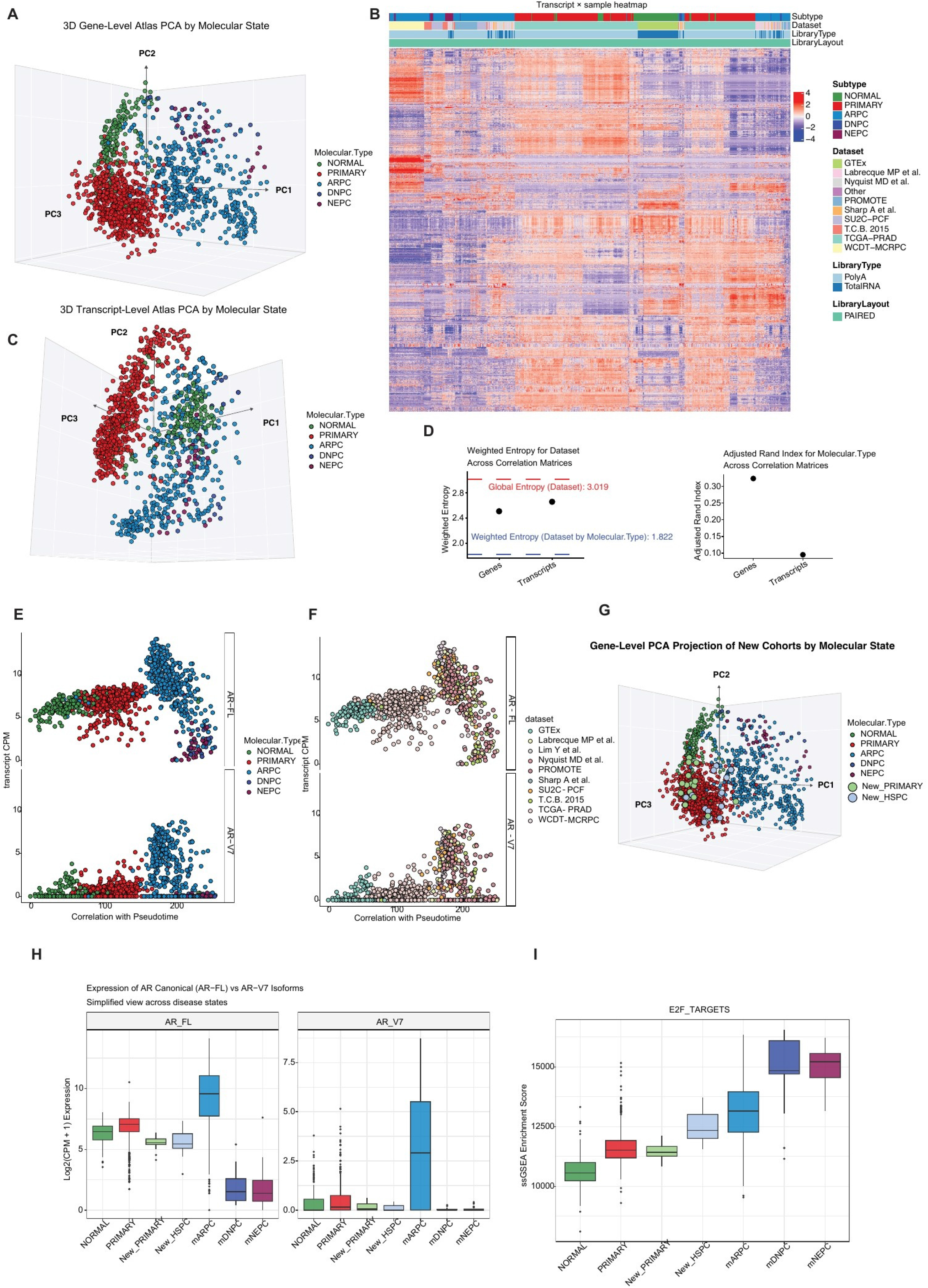
Construction and validation of an integrated transcript-level prostate cancer atlas. (A) Three-dimensional principal-component analysis (PCA) of gene-level expression in the integrated RNA-sequencing cohort, colored by molecular state (normal prostate, primary prostate cancer, androgen receptor-positive castration-resistant prostate cancer [ARPC], double-negative prostate cancer [DNPC], or neuroendocrine prostate cancer [NEPC]). (B) Transcript × sample heatmap based on the 2,000 most variable transcripts, with annotations for molecular state, source dataset, RNA-library type, and library layout. (C) Three-dimensional PCA based on transcript-level expression, colored as in (A). (D) Weighted entropy for source-dataset identity and adjusted Rand index for molecular state across gene-and transcript-level correlation matrices. Higher weighted entropy indicates greater mixing of datasets, whereas higher adjusted Rand index indicates stronger recovery of annotated molecular states. (E and F) Full-length androgen receptor (AR-FL) and AR-V7 expression plotted against prostate cancer pseudotime and colored by molecular state (E) or source dataset (F). (G) Projection of newly added primary and hormone-sensitive prostate cancer samples into the reference gene-level PCA space. (H) AR-FL and AR-V7 expression across the indicated disease states. (I) E2F_TARGETS ssGSEA scores across disease states. Expression is shown as log2(CPM + 1). Box plots show the median, interquartile range, and 1.5 × interquartile-range whiskers; points beyond the whiskers are shown individually.

### Extending the atlas to transcript resolution

To benchmark Salmon’s ability to resolve individual transcripts^19^, we compared its short-read-based expression estimates with matched MAS-Seq long-read data, which captures full-length poly-adenylated transcripts, across a panel of prostate cancer cell lines. Gene-and transcript-level expression correlated well between platforms (R = 0.8–0.83 for genes, R = 0.65 for transcripts), while the correlation dropped further for lncRNAs (R = 0.59 at gene level, R = 0.41 at transcript level), consistent with the frequent absence of poly(A) tails on lncRNAs and their consequent under-representation in long-read data (Fig. S3A–G). Salmon therefore provides a reasonable approximation of transcript-level expression.

We then screened the transcript-level atlas for residual technical batch effects. Poly(A)-selected and total-RNA libraries showed no measurable batch effect across studies (Fig. S4A, B), but single-versus paired-end library layout markedly separated primary tumors (Fig. S4C, D). We therefore excluded 225 single-end samples from four datasets (the NEPC lncRNA cohort [Trento/Cornell/Broad, 2015], Stelloo et al., EIMB RAS, and ANTE), retaining 1,140 samples from ten datasets for all subsequent transcript-level analyses (Fig. 1B, C; Fig. S4E–G). The resulting transcript-level atlas showed entropy comparable to the gene-level atlas but a lower adjusted Rand index, indicating that individual transcripts vary less sharply with disease progression than aggregate gene expression (Fig. 1D). CRPC samples nonetheless separated clearly from normal tissue and primary tumors along PC1–PC2, while PC3 separated normal tissue from primary disease; within CRPC, the most pronounced separation reflected treatment history, distinguishing androgen receptor pathway inhibitor (ARPI)-naive from ARPI-resistant tumors, with dataset and tissue of origin contributing comparatively little (Fig. S5A–D).

We validated the transcript-resolved atlas using full-length androgen receptor (AR-FL) and its constitutively active splice variant AR-V7^5,6^. As expected, AR-FL was robustly expressed in normal tissue and primary disease and further upregulated in ARPC, whereas AR-V7 was selectively upregulated in ARPC, a pattern consistent across contributing studies (Fig. 1E, F).

### The atlas accurately contextualizes newly generated external cohorts

Finally, we tested whether the atlas could correctly position independently generated data. We reprocessed a recently published bulk RNA-sequencing cohort of primary and hormone-sensitive de novo metastatic prostate cancer (New_PRIMARY and New_HSPC)^22^ and projected it onto the reference atlas. New_HSPC samples localized between primary tumors and early CRPC, and AR-FL and AR-V7 expression matched that of primary disease, consistent with these patients not having been exposed to hormonal therapy (Fig. 1G, H; Fig. S5E, F). By contrast, proliferation-associated gene sets (E2F targets, G2M checkpoint, and MYC targets) were more enriched in de novo metastatic disease than in primary disease alone (Fig. 1I; Fig. S5G, H). Together with the accompanying Prostate Cancer Atlas web tool enabling upload of external samples (www.prostatecanceratlas.org), these analyses establish a validated, transcript-resolution framework for placing new prostate cancer transcriptomic data within the context of disease progression.

### The lncRNA transcriptome atlas resolves prostate cancer progression

Restricting the transcript-resolution atlas to lncRNAs alone reproduced the dataset entropy and molecular-state recovery seen for lncRNA genes, indicating that the atlas integrates and resolves disease stages similarly at the lncRNA transcript level (Fig. 2A–C). Consistently, PCA of lncRNA transcript expression separated normal tissue, primary disease, and CRPC, with no discernible effect of poly(A)-selected versus total-RNA library preparation (Fig. 2D; Fig. S6A).

**Figure 2.**
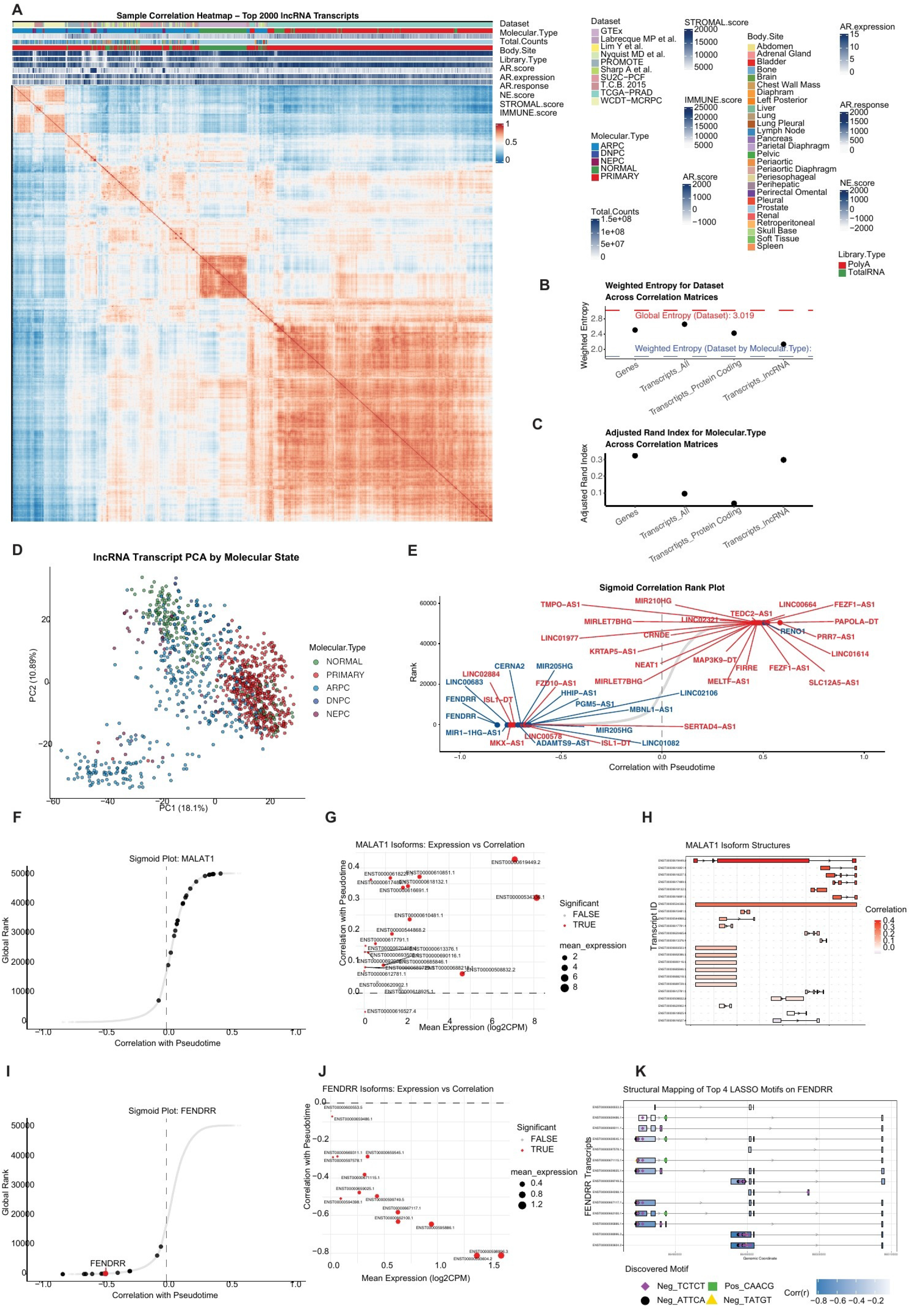
Transcript-resolved analysis identifies progression-associated long non-coding RNA isoforms. (A) Sample correlation heatmap based on the 2,000 most variable long non-coding RNA (lncRNA) transcripts. Annotation tracks indicate dataset, molecular state, library size, biopsy site, library type, AR score, AR expression, AR-response score, neuroendocrine score, stromal score, and immune score. (B and C) Weighted entropy for dataset identity (B) and adjusted Rand index for molecular state (C) across gene-level, all-transcript, protein-coding-transcript, and lncRNA-transcript matrices. (D) PCA of lncRNA transcript expression colored by molecular state. (E) Rank plot of Pearson correlations between individual lncRNA transcript abundance and pseudotime; selected positively and negatively associated lncRNAs are labeled. (F) Correlation-rank plot highlighting MALAT1; transcripts were ordered by Pearson coefficient and assigned consecutive ranks. (G) Mean log2(CPM + 1) expression and pseudotime correlation of MALAT1 transcripts. Point size denotes mean expression, and color denotes Benjamini–Hochberg-adjusted p <0.05. (H) MALAT1 transcript structures ordered by pseudotime correlation. (I) Correlation-rank plot highlighting FENDRR. (J) Mean expression and pseudotime correlation of FENDRR transcripts, displayed as in (G). (K) Genomic positions of the four highest-weight 5-mers selected by LASSO regression across FENDRR transcripts; transcript-specific pseudotime correlations are shown by the color scale.

To identify lncRNAs linked to disease progression, we first correlated individual lncRNA transcript abundance with pseudotime. LncRNAs with established oncogenic roles highlighted in red dominated the most positively correlated ones, while lncRNAs with reported tumor-suppressive functions highlighted in blue were enriched among the most negatively correlated ones (Fig. 2E). We selected three of the top oncogenic candidates — MALAT1, NEAT1, and FIRRE — and the top tumor-suppressive candidate, FENDRR, for transcript-level interrogation; all four showed the expected gene-level expression changes across normal, primary, ARPC, DNPC, and NEPC samples (Fig. S6B–E).

### Transcript-and exon-resolved analysis nominates functional regions within progression-associated lncRNAs

MALAT1 is an established oncogenic lncRNA whose expression in prostate cancer correlates with Gleason score, prostate-specific antigen levels, and progression to castration resistance^23^. Among MALAT1 transcripts, the most highly expressed isoform showed the strongest positive correlation with pseudotime (Fig. 2F, G), and mapping this correlation onto transcript structure showed that exons at the 3’ end contributed most strongly (Fig. 2H) — the region encoding the triple-helical stability element that protects MALAT1’s 3’ end from exonucleolytic decay and promotes its nuclear accumulation^24^, suggesting this stability determinant becomes increasingly important as disease progresses.

NEAT1, an estrogen-receptor-alpha-regulated lncRNA that sustains an oncogenic transcriptional program in prostate cancer and blunts sensitivity to AR-directed therapy^25^, showed a similar pattern: its most upregulated transcript was the canonical isoform, and although individual-transcript correlations with pseudotime were weaker than for MALAT1, the strongest associations again mapped to 3’-end exons (Fig. S6F–H) — the same triple-helix-forming region that stabilizes the long NEAT1_2 isoform required for paraspeckle formation^24^, pointing to a conserved, structure-linked mode of progression-associated regulation.

FIRRE, which organizes multi-chromosomal contacts through a nuclear-matrix-interacting repeat domain^26^, showed positive pseudotime correlations across essentially all transcripts, strengthening with expression as for MALAT1 (Fig. S6I, J). Unlike MALAT1 and NEAT1, no single exon dominated this association; unbiased k-mer regression instead identified discrete sequence motifs, mapping to discontinuous positions across FIRRE transcripts, whose enrichment tracked most closely with disease progression (Fig. S6K, L) — reminiscent of the modular, repeat-element architecture through which FIRRE exerts its functions^27^.

Conversely, FENDRR, a chromatin-modifying tumor suppressor lncRNA^28^ that is silenced in several epithelial cancers^29^, was strongly downregulated with pseudotime (Fig. 2E, I; Fig. S6D), and its most highly expressed transcript showed the most negative correlation (Fig. 2J). As for FIRRE, no single exon explained this pattern; k-mer regression instead identified discrete motifs distributed across the FENDRR transcript structure that were most negatively associated with progression (Fig. 2K; Fig. S6M).

Together, these analyses show that unbiased, transcript-and sequence-level interrogation of lncRNAs can pinpoint the specific exons or sequence elements that underlie their progression-associated, oncogenic or tumor-suppressive effects.

### Canonical transcripts dominate progression-associated changes in protein-coding isoforms

The protein-coding transcript atlas showed entropy and dataset integration comparable to the lncRNA and gene-level atlases (Fig. 2B; Fig. 3A; Fig. S7A), with good mixing across contributing datasets and no discernible effect of poly(A)-selected versus total-RNA library preparation (Fig. S7B, C).

**Figure 3.**
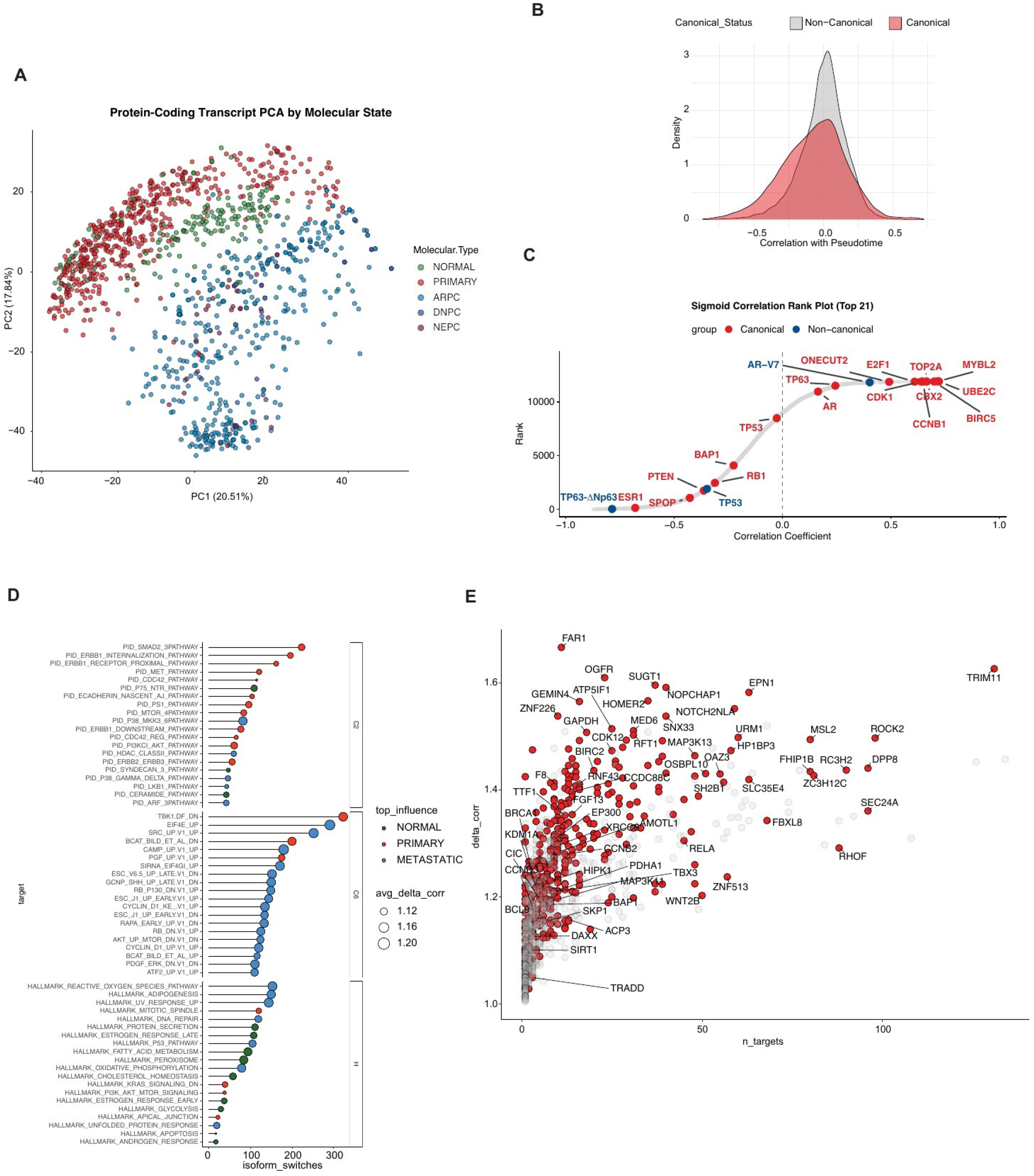
Systematic identification of pathway-associated isoform switches in prostate cancer. (A) PCA of protein-coding transcript expression, colored by molecular state. (B) Density distributions of transcript– pseudotime correlations for all analyzed protein-coding transcripts, stratified by canonical status. (C) Correlation-rank plot of transcript–pseudotime Pearson coefficients. Transcripts were ordered by correlation coefficient and assigned consecutive ranks. Red points denote canonical transcripts, which generally represent the most extreme positively or negatively correlated isoform within the corresponding gene; blue points denote non-canonical transcripts that instead show the most extreme correlation within their genes. Selected prostate-cancer-relevant examples, including AR-V7, TP63ΔNp63, and TP53, are labeled. For each of the three highlighted non-canonical transcripts, the corresponding canonical transcript is also indicated in red. (D) Numbers of detected isoform switches associated with the indicated C6 oncogenic, C2 PID, and Hallmark gene-set scores. Point color denotes the disease compartment with the greatest influence on the switch, point size denotes the average delta correlation, and the 20 gene sets with the largest switch counts in each category are shown. (E) Delta correlation for each representative isoform pair plotted against the number of associated pathway targets. Highlighted red points denote selected genes, and cancer-relevant genes are labeled. Candidate switches were restricted to protein-coding transcripts contributing >40% of parent-gene expression in >5% of samples and having >1 count in >5% of samples; at least one transcript required r >0.5 and another r <−0.5 for the same target, and retained genes encoded at least two distinct products.

Ranking protein-coding transcripts by their correlation with pseudotime showed that canonical isoforms accounted for most of the top up-and downregulated transcripts along disease progression, as expected (Fig. 3B, C; Fig. S7D, E). Among the canonical isoforms, previously noted genes related to proliferation (MYBL2, CCNB1, TOP2, UBE2C, CDK1), survival (BIRC5), and AR were among the top upregulated canonical isoforms. Among the downregulated canonical isoforms, we found many tumor suppressors such as BAP1, RB1, PTEN, and SPOP, as expected (Fig. 3C).

Gene set enrichment analysis performed separately on canonical-and non-canonical-driven genes showed that non-canonical transcripts were selectively enriched for epithelial-mesenchymal transition and myogenesis signatures (Fig. S7F), whereas canonical transcripts recovered cell-proliferation pathways consistent with our previous gene-level observations (Fig. S7G)^16^, suggesting a more pronounced role for non-canonical isoforms in pathways governing cell identity. Among genes for which a non-canonical transcript became progressively dominant over its canonical counterpart, the androgen receptor AR-V7 was the only one among the upregulated transcripts (Fig. 3C). Among the downregulated transcripts, the shorter TP63-DNp63 of TP63 specific to basal cells was lost during tumorigenesis and negatively correlated with pseudotime, as expected.

### Isoform switches nominate oncogenes and tumor suppressors across disease progression

We next leveraged the integrated protein-coding transcriptome to systematically associate isoform switches with gene signatures reflecting normal biological processes (Hallmark) and oncogenic states (curated C2 and oncogenic C6 gene sets). Switches associated with Hallmark signatures were predominantly detected in normal tissue, switches associated with C2 pathways mainly in primary disease, and switches associated with C6 oncogenic signatures mostly in metastatic (ARPC, DNPC, NEPC) disease, consistent with the increasingly cancer-specific, transformation-related processes these gene sets capture (Fig. 3D; Fig. S8A, B).

The genes most frequently affected by these signature-associated switches included established oncogenes — RELA^30^, SKP1^31^, BIRC2^32^, TRIM11^33^, ROCK2^34^, and WNT2B^35^ — and tumor suppressors — BAP1^36,37^, BRCA1^38^, RNF43^39^, CDK12^40^, and CIC^41^ (Fig. 3E). We dissect a subset of these gene pairs — WNT2B and RNF43, RELA and BRCA1, and ROCK2 and TRIM11 — in the sections that follow, illustrating how the atlas can nominate and help validate thus far underappreciated isoform-specific functions for bona fide cancer genes.

### WNT2B isoform switching tracks divergent WNT-pathway activity

WNT2B is a canonical WNT ligand that activates β-catenin-dependent signaling^35^. The atlas resolved two WNT2B transcripts differing in their N-terminus: the canonical isoform (WNT2B-202) retains an N-terminal extension folded as an alpha helix that is absent from the shorter isoform (WNT2B-203) (Fig. 4A, B). Long-read sequencing confirmed both isoforms in prostate cancer cell lines, with the canonical transcript predominant and short-read Salmon quantification yielding concordant relative abundances (Fig. S9A).

**Figure 4.**
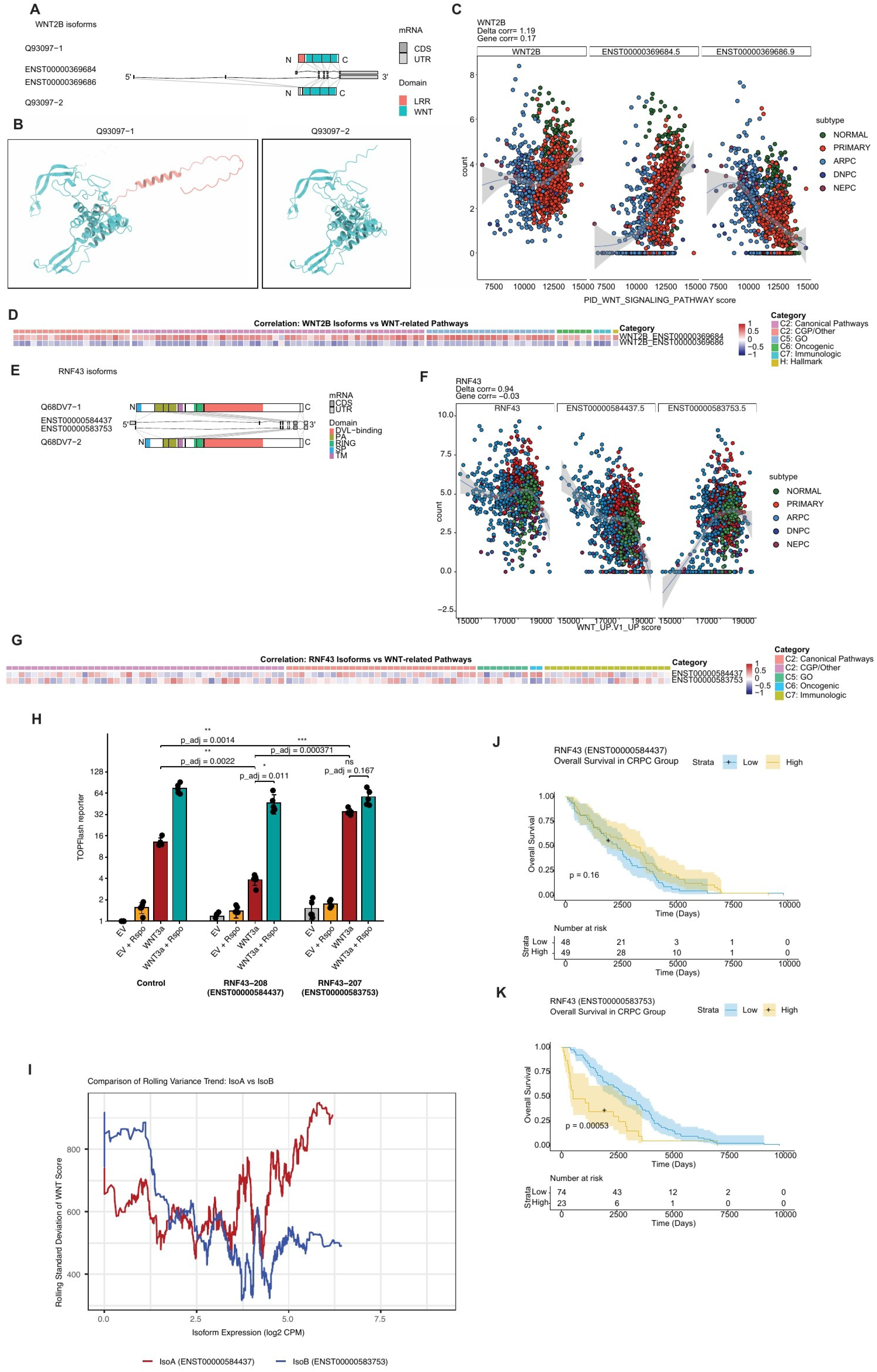
WNT2B and RNF43 isoform switching is associated with divergent WNT-pathway activity and clinical outcome. (A) Transcript and protein-domain organization of WNT2B-202 (ENST00000369684.5; Q93097-1) and WNT2B-203 (ENST00000369686.9; Q93097-2). (B) AlphaFold-predicted structures of the two WNT2B protein isoforms, visualized in UCSF ChimeraX v1.12. (C) Gene-level WNT2B and the two indicated transcripts plotted against PID_WNT_SIGNALING_PATHWAY ssGSEA scores. Points represent tumors and are colored by molecular state; gray curves and bands denote smoothed trends and confidence intervals. Gene-level correlation and delta correlation are indicated. (D) Correlations of WNT2B-202 and WNT2B-203 with WNT-related pathway scores from the indicated MSigDB collections. (E) Transcript and domain organization of RNF43-208 (ENST00000584437.5; Q68DV7-1) and RNF43-207 (ENST00000583753.5; Q68DV7-2). (F) Gene-level RNF43 and the two transcripts plotted against WNT_UP.V1_UP scores, displayed as in (C). (G) Correlations of RNF43-208 and RNF43-207 with WNT-related pathway scores. (H) TOPFlash reporter activity in HEK293T cells expressing control, canonical RNF43-208 (ENST00000584437), or truncated RNF43-207 (ENST00000583753) constructs under empty-vector (EV), EV plus R-spondin-1, WNT3A, or WNT3A plus R-spondin-1 conditions. R-spondin-1 (200 ng/mL) was added 2 h after transfection, and luminescence was measured 24 h after transfection. Firefly luciferase activity was normalized to Renilla activity and then expressed relative to the empty-vector (EV) condition in the control arm. Bars show mean ± SD, and points represent five biological replicates. Five prespecified comparisons were analyzed using two-sided paired t tests with Šidák correction: Control versus RNF43-208 under WNT3A, adjusted p = 0.0022 (**); Control versus RNF43-207 under WNT3A, adjusted p = 0.0014 (**); RNF43-208 versus RNF43-207 under WNT3A, adjusted p = 0.000371 (***); WNT3A versus WNT3A plus R-spondin-1 within RNF43-208, adjusted p = 0.0109 (*); and the same comparison within RNF43-207, adjusted p = 0.167 (ns). *adjusted p < 0.05, **adjusted p < 0.01, ***adjusted p < 0.001; ns, not significant. (I) Centered rolling standard deviation of the original WNT_UP.V1_UP score after separate ordering by RNF43-208 (IsoA, red) or RNF43-207 (IsoB, blue) expression; the window was max(50, floor[0.05 × n]) samples with a one-sample step. (J and K) Kaplan–Meier estimates of overall survival in CRPC stratified by low or high RNF43-208 (J) or RNF43-207 (K) expression. Shading denotes 95% confidence intervals, tick marks denote censoring, and numbers at risk are shown.

Consistent with a canonical-activating role, expression of the canonical WNT2B transcript correlated positively with WNT pathway activity, whereas the shorter transcript correlated inversely, a pattern reproduced across independent WNT gene sets and related pathways such as epithelial-to-mesenchymal transition (EMT), JAK-STAT3-, and TGF-beta-signaling (Fig. 4C, D; Fig. S9B, C).

### RNF43 isoform switching uncouples WNT restraint from R-spondin regulation and predicts survival

RNF43 is a transmembrane E3 ubiquitin ligase that restrains WNT signaling by ubiquitinating and degrading Frizzled receptors, a restraint relieved by R-spondin binding to LGR4/5^42,43^; loss-of-function alterations in this axis, including R-spondin gene fusions, recur in advanced prostate cancer^44^. We identified a shorter RNF43 transcript lacking part of the N-terminal domain required for membrane anchoring (Fig. 4E). Long-read sequencing and the Salmon quantification showed again concordant relative abundances in cell lines (Fig. S9D). Unlike the canonical transcript, whose expression correlated negatively with WNT pathway activity as expected for a pathway inhibitor, the short isoform correlated positively with WNT activity (Fig. 4F), although this relationship was less consistent across the broader panel of WNT-related gene sets (Fig. 4G).

A TOPFlash reporter assay confirmed this divergence: the canonical isoform repressed WNT3A-induced signaling, while the short isoform enhanced it (Fig. 4H). R-spondin-1 restored full WNT activity in cells expressing the canonical isoform but had little additional effect on the short isoform, which alone already drove signaling to near-maximal levels — consistent with a model in which the short isoform antagonizes endogenous RNF43.

In line with R-spondin-dependent regulation of the canonical isoform, the spread of WNT pathway activity across samples increased with canonical RNF43-208 expression, whereas the short RNF43-207 isoform showed increasing activity with less variability at higher expression (Fig. 4I; Fig. S9G, H). Clinically, high expression of the short isoform was associated with significantly poorer overall survival in CRPC, while the canonical isoform showed no such association (Fig. 4J, K); this relationship did not reach significance for progression-free survival in primary disease (Fig. S9I, J), consistent with a CRPC-specific, R-spondin-independent driver of WNT activity.

### RELA isoform switching reduces NF-κB pathway activity through loss of the TAD2 transactivation domain

RELA (p65) is a bona fide oncogene and the principal transactivating subunit of NF-κB^30^. The atlas resolved two RELA transcripts, also detected by long-read sequencing, whose expression inversely correlated with NF-κB activity signatures and related pathways (Fig. 5A–D; Fig. S10A–E).

**Figure 5.**
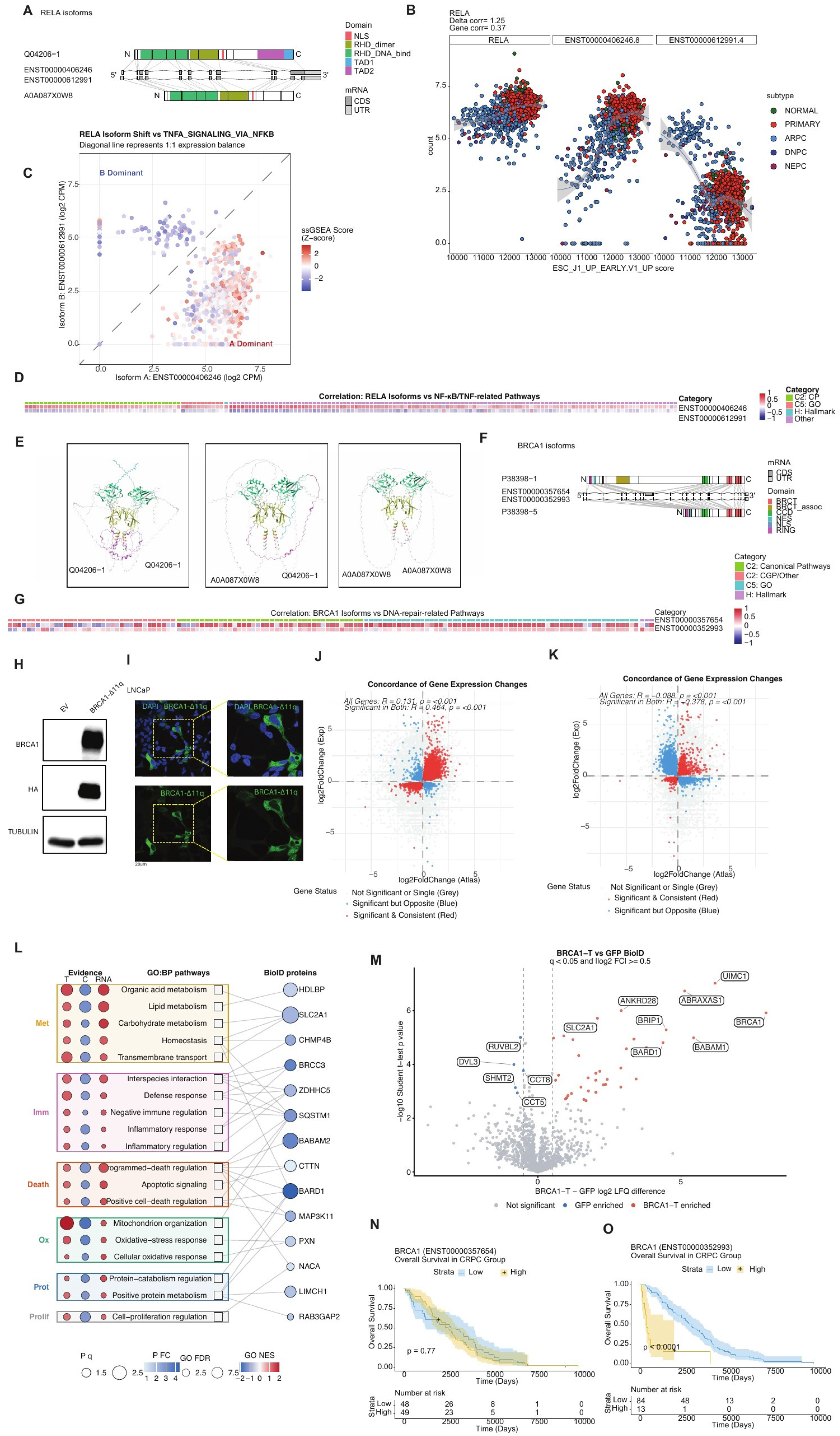
RELA and BRCA1 isoforms show opposing pathway associations and distinct functional and clinical phenotypes. (A) Transcript and protein-domain organization of RELA-202 (ENST00000406246.8; Q04206-1) and RELA-226 (ENST00000612991.4; A0A087X0W8). (B) Gene-level RELA and the two indicated transcripts plotted against ESC_J1_UP_EARLY.V1_UP scores. Points represent tumors and are colored by molecular state; gray curves and bands denote smoothed trends and confidence intervals. Gene-level and delta correlations are indicated. (C) RELA isoform-dominance plot colored by the z-scored TNFA_SIGNALING_VIA_NFKB ssGSEA score; the diagonal denotes equal transcript expression. (D) Correlations of RELA-202 and RELA-226 with NF-κB/TNF-related pathway scores from the indicated MSigDB collections. (E) AlphaFold-predicted homodimeric structures of RELA-202 and RELA-226 and the predicted RELA-202–RELA-226 heterodimer, visualized in UCSF ChimeraX v1.12. (F) Transcript and protein-domain organization of canonical/full-length BRCA1-203 (BRCA1-C; ENST00000357654.9; P38398-1) and truncated BRCA1-201/BRCA1-Δ11q (BRCA1-T; ENST00000352993.7; P38398-5). (G) Correlations of BRCA1-C and BRCA1-T with DNA-repair-related pathway scores. (H) Immunoblot of BRCA1 and HA-tagged BRCA1-Δ11q in LNCaP cells infected with empty-vector or BRCA1-Δ11q lentivirus; tubulin is the loading control. The experiment was performed once. (I) Representative immunofluorescence images of LNCaP cells expressing HA-tagged BRCA1-Δ11q. DAPI labels nuclei and green fluorescence denotes anti-HA staining; the boxed region is enlarged at right. Scale bar, 20 μm. The imaging experiment was performed once. For the perturbation RNA-seq experiment, three independent biological replicates were analyzed per condition. (J) Concordance between gene-expression changes associated with high versus low BRCA1-T expression in the atlas and changes induced by BRCA1-T/Δ11q overexpression. Red denotes genes significant and directionally concordant in both datasets, blue denotes significant but discordant genes, and gray denotes non-significant or single-dataset genes. Pearson correlations are shown for all genes and for genes significant in both datasets. (K) Concordance between gene-expression changes induced by BRCA1-T/Δ11q overexpression and those associated with high versus low BRCA1-C expression in the atlas, displayed and classified as in (J). (L) Bipartite network linking GO Biological Process terms supported by conditional BRCA1 isoform associations in the atlas and by BRCA1-T-overexpression RNA-seq to significant non-bait BRCA1-T-enriched proximity-labeled proteins. The three evidence columns represent Atlas-T, Atlas-C, and experimental BRCA1-T RNA-seq; fill denotes normalized enrichment score (NES) and size denotes −log10(FDR). Protein-node fill denotes the BRCA1-T-minus-GFP BioID log2 LFQ difference, protein-node size denotes −log10(q value), and edges denote MSigDB GO:BP membership. Shaded boxes denote descriptive biological-process modules. (M) Volcano plot of differential proximity labeling between BRCA1-T BioID and GFP BioID controls across four replicate samples per group. The x axis shows the BRCA1-T-minus-GFP log2 LFQ difference and the y axis shows −log10(Student’s t test p value). Proteins meeting Benjamini–Hochberg q < 0.05 and |log2 fold change| ≥ 0.5 are colored by enrichment direction; selected proteins are labeled. (N and O) Kaplan–Meier estimates of overall survival in CRPC stratified by low or high BRCA1-C (N) or BRCA1-T (O) expression, with 95% confidence intervals, censor marks, and risk tables.

RELA binds DNA as a homodimer through its Rel homology domain (RHD)^45^, a configuration reinforced by the C-terminal TAD2 transactivation domain^46^, which is absent from the shorter isoform (Fig. 5A). Consistent with this, AlphaFold-predicted structures showed a stable homodimer for the canonical isoform, while the short isoform’s homodimer lacked the corresponding C-terminal region (Fig. 5E). Pathway activity decreased gradually with increasing expression of the short isoform relative to the canonical one (Fig. 5C; Fig. S10C, D), a pattern compatible with the formation of less-active heterodimers between the two isoforms through their intact RHD domains.

### A BRCA1 isoform lacking exon 11 retains partial function, engages cytoplasmic interactors, and predicts poor survival in CRPC

BRCA1 is a tumor suppressor central to the resolution of DNA double-strand breaks by homologous recombination^38^. We detected significant pathway-activity differences between canonical, full-length BRCA1 and a previously described shorter isoform lacking exon 11 (BRCA1-D11)^47^ (Fig. 3E, 5F; Fig. S10F, G). In line with a dampened but partially proficient DNA-repair capacity — the same property that has been linked to PARP-inhibitor and cisplatin resistance^48^ — the BRCA1-D11 isoform showed reduced activity of BRCA1-and DNA-repair-related signatures, including the G2M checkpoint, relative to the canonical isoform in the atlas (Fig. 5G; Fig. S10H).

To functionally validate this isoform, we overexpressed the less-characterized BRCA1-D11 isoform in LNCaP prostate cancer cells (Fig. 5H). Consistent with the absence of a nuclear localization signal within exon 11 (Fig. 5F), the short isoform localized predominantly to the cytoplasm (Fig. 5I). RNA sequencing of the LNCaP overexpression model showed transcriptional changes that concorded with those associated with BRCA1-D11 but not the canonical full-length isoform in the atlas (Fig. 5J, K; Fig. S10I).

To search for biological pathways specific to BRCA1-D11 but not the canonical isoform, we fitted a linear model to disentangle each isoform’s contribution and subsequently interrogated Gene Ontology biological process signatures for differences that replicated specifically for the BRCA1-D11 in the LNCaP experiment (Fig. 5L). Among the processes enriched for the short isoform were cytoplasmic functions such as energy and cell metabolism, inflammatory response, and programmed cell death — consistent with the predominantly cytoplasmic localization of BRCA1-Δ11q.

To identify effector proteins associated with the short isoform, we performed proximity-dependent biotin identification (BioID) in LNCaP cells expressing either the short BRCA1 isoform or GFP (Fig. 5M). The resulting interactome recovered established BRCA1-associated DNA-repair factors, including BARD1, ABRAXAS1, UIMC1, BABAM1, and BRIP1, confirming specificity. Alongside these, BioID identified interactors not previously linked to BRCA1 — among them HDLBP (Vigilin), SLC2A1 (GLUT1), CTTN (cortactin), ZDHHC5, MAP3K11 (MLK3), PXN (paxillin), NACA, LIMCH1, and RAB3GAP2 — whose functions overlap with the biological processes nominated by the linear model (Fig. 5L). Clinically, high expression of the short isoform, but not the canonical isoform, was associated with significantly poorer overall survival in CRPC (Fig. 5N, O), an association not observed for progression-free survival in primary disease (Fig. S10J, K). Together, these data may point to a thus far unrecognized function of the short BRCA1 isoform in cancer outside of DNA repair.

### ROCK2 isoform switching identifies a truncated kinase domain that dominantly restrains Rho-kinase activity

ROCK2 is a Rho-associated coiled-coil kinase that drives actomyosin contractility and cell motility, promoting epithelial-to-mesenchymal transition (EMT), invasion, and metastatic spread downstream of and in cooperation with WNT, TGF-β, and NOTCH signaling^34,35,51^ (Fig. 3E). The atlas resolved two ROCK2 transcripts differing in their N-terminus: the canonical isoform (ROCK2-202) retains the intact kinase domain, whereas the shorter isoform (ROCK2-203) lacks part of the N-terminal kinase-domain extension, confirmed by long-read sequencing in prostate cancer cell lines (Fig. 6A; Fig. S11A). Canonical ROCK2 expression remained comparatively stable across disease stages, while the activity of EMT and other ROCK-associated Hallmark and pathway signatures instead tracked inversely with expression of the shorter isoform in an isoform-dominance analysis (Fig. 6B, C; Fig. S11B–F), suggesting that the truncated isoform exerts a dominant-negative effect on ROCK2 signaling.

**Figure 6.**
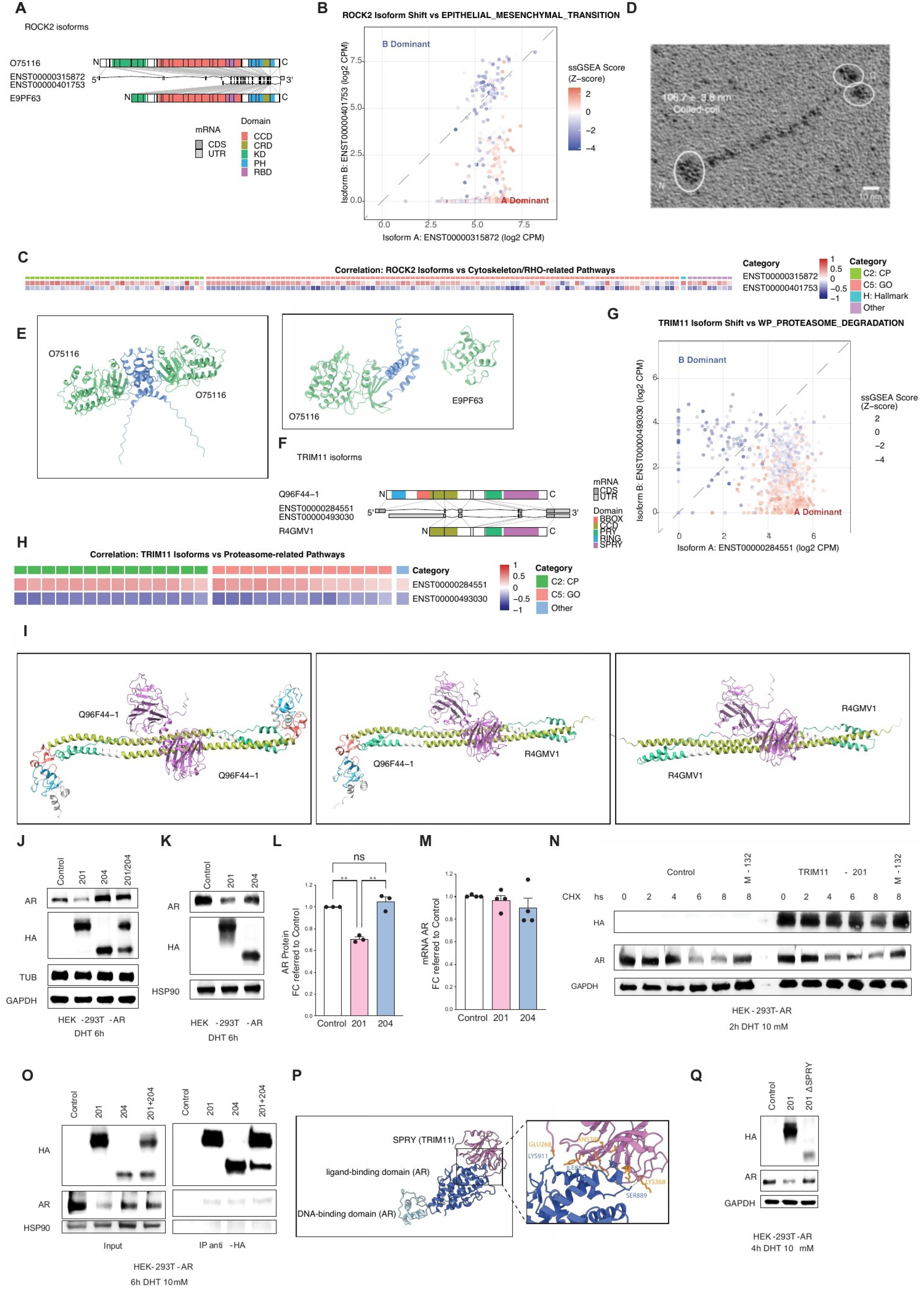
ROCK2 and TRIM11 isoform switching identifies structurally divergent proteins and a TRIM11– AR interaction. (A) Transcript and protein-domain organization of ROCK2-202 (ENST00000315872.11; O75116) and ROCK2-203 (ENST00000401753.5; E9PF63). (B) ROCK2 isoform-dominance plot colored by the z-scored HALLMARK_EPITHELIAL_MESENCHYMAL_TRANSITION score. (C) Correlations of the two ROCK2 transcripts with cytoskeleton/RHO-related pathway scores from the indicated MSigDB collections. (D) Rotary-shadowing electron micrograph of a representative full-length human ROCK2 particle, showing an extended dimer with a long central coiled-coil and asymmetric terminal globular densities; circled regions highlight terminal domains. Scale bar, 4 nm. Reproduced from Figure 1e of Truebestein et al. (2015), Nature Communications, under the Creative Commons Attribution 4.0 International License. (E) AlphaFold-predicted N-terminal structures of the full-length ROCK2-202 homodimer and the ROCK2-202–ROCK2-203 heterodimer, visualized in UCSF ChimeraX v1.12; only the N-terminal regions are displayed. (F) Transcript and domain organization of TRIM11-201 (ENST00000284551.11; Q96F44-1) and TRIM11-204 (ENST00000493030.6; R4GMV1). (G) TRIM11 isoform-dominance plot colored by the z-scored WP_PROTEASOME_DEGRADATION score. (H) Correlations of TRIM11-201 and TRIM11-204 with proteasome-related pathway scores. (I) AlphaFold-predicted homo-and heteromeric structures of TRIM11-201 and TRIM11-204, visualized in UCSF ChimeraX v1.12. (J) Immunoblot analysis of AR and HA-tagged TRIM11 in HEK293T-AR cells transfected with empty-vector control, TRIM11-201, TRIM11-204, or both isoforms. (K) Immunoblot analysis of AR and HA-tagged TRIM11 in HEK293T-AR cells transfected with empty-vector control, TRIM11-201, or TRIM11-204. For (J) and (K), cells were maintained in phenol-red-free DMEM supplemented with 10% charcoal-stripped serum, analyzed 24 h after transfection, and treated with 10 nM dihydrotestosterone (DHT) for 6 h; loading controls are shown. (L) AR protein abundance in (K), normalized to GAPDH and expressed relative to control. (M) AR mRNA abundance measured by one-step RT-qPCR, normalized to GAPDH and expressed relative to control. Data in (L) and (M) are mean ± SEM from at least three independent experiments; comparisons were performed by one-way ANOVA. Significance annotations in (L): **p < 0.01; ns, not significant. (N) Cycloheximide-chase analysis of AR stability in HEK293T-AR cells transfected with empty-vector control or TRIM11-201. Cells were treated with 10 nM DHT for 2 h before addition of cycloheximide (50 μg/mL), with MG132 (10 μM) included where indicated, and collected at the displayed time points. (O) Co-immunoprecipitation of HA-tagged TRIM11 isoforms with AR. HEK293T-AR cells were transfected with empty-vector control, TRIM11-201, TRIM11-204, or both isoforms, treated with 10 nM DHT for 6 h, and subjected to anti-HA immunoprecipitation; inputs and immunoprecipitates were immunoblotted for HA, AR, and HSP90. (P) AlphaFold 3 prediction of the interaction between full-length human TRIM11 (UniProt Q96F44; 468 amino acids) and full-length human AR (UniProt P10275), modeled at 1:1 stoichiometry. The top-ranked of five models returned by AlphaFold-beta-20231127 under server-default settings is shown. The model was visualized in UCSF ChimeraX v1.12 using the publication preset. Candidate interface residues were defined by inter-chain atom pairs within 4.0 Å and displayed as sticks; detected inter-chain hydrogen bonds are shown in yellow, and displayed interface residues are labeled by residue name and sequence position. No ligands, metal ions, nucleic acids, or post-translational modifications were specified. (Q) Immunoblot analysis comparing empty-vector control, full-length TRIM11-201, and a TRIM11-201 construct lacking the SPRY domain in HEK293T-AR cells under the conditions used in (J) and (K).

ROCK2 is a constitutive, elongated dimer held together by a long central coiled-coil, with the paired kinase domains positioned at one end (Fig. 6D)^49^. Kinase activity additionally depends on an intramolecular interaction between the hydrophobic motif of the kinase domain and its N-terminal extension^50^ — the region truncated in the short isoform. AlphaFold-predicted structures of ROCK2 homo-and heterodimers highlighted the altered architecture of this kinase-proximal region in the short isoform (Fig. 6E), consistent with a model in which heterodimerization between canonical and truncated ROCK2 disrupts this interaction and thereby also restrains the kinase activity of the intact subunit within the dimer.

### TRIM11 isoform switching uncovers a RING-and SPRY-domain-dependent mechanism of androgen receptor degradation

TRIM11 is a tripartite-motif E3 ubiquitin ligase that ubiquitinates its substrates through an N-terminal RING domain, targeting them for proteasomal degradation^33,53^, and was the gene most extensively associated with isoform-switch-linked pathway and signature changes in the atlas (Fig. 3E). The atlas resolved a canonical transcript (TRIM11-201) containing the B-box, coiled-coil (CCD), RING, PRY, and SPRY domains, and a shorter transcript (TRIM11-204) lacking the RING domain, both confirmed by long-read sequencing (Fig. 6F; Fig. S11G). In agreement with a RING-dependent function, expression of the canonical isoform correlated positively with pathways governing protein turnover, proteasomal degradation, and ubiquitination, an association not observed — and in part reversed — for the truncated isoform (Fig. 6G, H; Fig. S11H–N).

Like other TRIM-family ligases, TRIM11 is predicted to dimerize through its coiled-coil domain, a feature retained in both isoforms (Fig. 6I), and RING-domain dimerization is required for the catalytic activity of many TRIM E3 ligases^52^. Because TRIM11-204 retains the coiled-coil but lacks the RING domain, heterodimers between the two isoforms may sequester canonical TRIM11-201 in complexes with reduced ubiquitin-ligase activity, providing a plausible mechanism for the truncated isoform’s opposing pathway associations.

The truncated isoform was specifically upregulated in castration-resistant disease and associated with increased androgen receptor (AR) signaling (Fig. S11N; Fig. S12A–D), nominating TRIM11 as a candidate regulator of AR, a transcription factor whose stability is itself governed by the ubiquitin-proteasome system^54^.

Supporting this, forced expression of TRIM11-204 alone or together with TRIM11-201 increased AR protein levels in HEK293T cells stably expressing AR, whereas TRIM11-201 alone reduced them (Fig. 6J). This effect occurred at the protein rather than the transcript level (Fig. 6K–M) and reflected accelerated AR degradation, as cycloheximide-chase experiments showed faster AR turnover in the presence of TRIM11-201 that was rescued by proteasome inhibition with MG132 (Fig. 6N).

Co-immunoprecipitation confirmed that both TRIM11-201 and TRIM11-204 physically interact with AR (Fig. 6O), and structural modeling predicted an interface between the TRIM11 SPRY domain and the AR ligand-and DNA-binding domains (Fig. 6P). Deletion of the SPRY domain abolished the ability of TRIM11-201 to reduce AR levels (Fig. 6Q), indicating that SPRY-mediated substrate recognition together with an intact RING domain are both required for AR degradation. Together, these data support a model in which canonical TRIM11-201 degrades AR via SPRY-domain-mediated substrate binding and RING-dependent ubiquitination, while the RING-deficient TRIM11-204 isoform antagonizes this activity — both directly, by competing for AR binding, and indirectly, through heterodimerization with TRIM11-201.

## Discussion

Most cancer genes can generate multiple RNA transcripts with distinct, sometimes opposing, functions, yet whether and how this transcript diversity is remodeled during disease progression in specific cancer types has remained largely unexplored, because no existing resource combines transcript resolution, cross-study harmonization, and full disease-stage coverage while preserving genuine biological heterogeneity. By reprocessing raw RNA-sequencing data from 10 independent studies spanning normal prostate tissue through primary, hormone-sensitive, and castration-resistant disease with a uniform computational pipeline, we built an integrated, transcript-resolution atlas that removes largely technical batch effects while retaining the biological heterogeneity that distinguishes these disease states.

This harmonized structure is what makes it possible to correlate transcript-specific expression, rather than gene-level expression alone, with disease-progression pseudotime and oncogenic pathway activity across a diverse patient population — an association map that collapses if cross-study batch effects are not first removed without erasing genuine disease-stage differences. That the atlas correctly positioned newly incorporated external cohorts of primary and hormone-sensitive metastatic disease, and agreed with long-read sequencing and with our previous gene-level atlas^16^, indicates that this framework generalizes beyond the studies used to build it and can serve as a stable reference onto which new prostate cancer transcriptomic data can be mapped.

At the lncRNA level, this resolution let us move beyond the gene-level associations already established for individual lncRNAs — such as MALAT1’s correlation with Gleason grade and castration resistance^23^ or NEAT1’s role in sustaining an oncogenic, AR-therapy-resistant transcriptional program^25^ — to ask which specific transcripts, and even which exons or sequence motifs, underlie these associations. That the strongest pseudotime correlations for both MALAT1 and NEAT1 mapped onto the same 3’ triple-helix-forming element that stabilizes both transcripts^24^ suggests that this structural determinant, rather than transcript abundance alone, becomes increasingly important as disease progresses, whereas the discrete, repeat-associated motifs we nominated for FIRRE and FENDRR point to a distinct, more modular mode of regulation^26,27^. More broadly, these analyses illustrate how transcript-and exon-resolved interrogation of a harmonized atlas can generate specific, testable hypotheses about the sequence elements that underlie a lncRNA’s oncogenic or tumor-suppressive activity, complementing gene-level lncRNA atlases with a finer-grained, mechanistically actionable layer.

At the protein-coding level, the atlas let us systematically test whether isoform switches, rather than changes in total gene expression, track oncogenic pathway activity. A recurring theme among the genes we examined is that a shorter isoform lacking a specific functional domain modulates the activity of its canonical counterpart, most plausibly through heterodimerization. Dominant-negative isoform pairs of this kind have precedent in other systems — for example, a truncated TEAD4 variant that retains the YAP-interaction domain but loses DNA binding and thereby attenuates Hippo-YAP signaling^55^, and a dominant-negative PPARγ1 splice variant^56^ — and our findings extend this concept to ROCK2, RELA, and TRIM11 in prostate cancer.

The kinetics of the switch, however, differed between genes. For ROCK2, canonical expression remained comparatively stable across disease stages while pathway activity instead tracked the shorter isoform, consistent with a dominant-negative effect exerted onto a constitutively expressed canonical pool. For RELA and TRIM11, pathway activity declined more gradually as expression shifted from a canonical-dominant to a short-isoform-dominant state, suggesting a more continuous titration of dimer composition along disease progression. Both patterns are compatible with a shared structural logic: in each case, the truncated isoform retains the dimerization interface — the Rel homology domain of RELA, the coiled-coil of TRIM11, the elongated coiled-coil of ROCK2 — while losing a domain required for full activity (TAD2, RING, or the kinase-domain N-terminal extension, respectively), predicting that canonical-short heterodimers have intermediate or suppressed activity relative to canonical homodimers.

Functional follow-up remains limited for several of these candidates, and the atlas points to specific, testable hypotheses for each. For RNF43, the short isoform’s resistance to R-spondin-mediated repression and its association with poor CRPC survival suggest that isoform switching may represent an alternative route to disrupting RNF43-mediated WNT restraint, distinct from the recurrent RNF43 mutations described in colorectal and endometrial cancer^39^ and from the R-spondin gene fusions recurrent in advanced prostate cancer^44^; whether short-isoform upregulation and loss-of-function mutation are mutually exclusive or co-occurring events in individual tumors remains an open question.

For BRCA1, the concordance between the short isoform’s atlas-derived pathway associations, its cytoplasmic localization, transcriptional consequences upon overexpression, and the BioID-identified interactome in LNCaP cells supports a model in which BRCA1-Δ11q partitions BRCA1 function between nuclear DNA repair and a possibly distinct more cytoplasmic roles in metabolism and energy supply. The recovery of established DNA-repair partners (BARD1, ABRAXAS1, BRIP1) alongside novel cytoplasmic interactors such as SLC2A1 (GLUT1), CTTN (cortactin), and MAP3K11 (MLK3) — whose functions mirror the metabolic, inflammatory, and cell-death processes independently nominated by the linear model — suggests that BRCA1-Δ11q engages effectors beyond the canonical homologous-recombination machinery, consistent with a dampened but partially proficient DNA-repair capacity and its previously reported link to PARP-inhibitor and cisplatin resistance^48^.

For TRIM11, our data nominate the androgen receptor as a substrate degraded by canonical TRIM11 through coordinated SPRY-domain substrate recognition and RING-domain-dependent ubiquitination, a mechanism specifically disabled by the RING-deficient short isoform that predominates in castration-resistant disease. Further evidence for stabilization of AR in advanced disease comes from the downregulation of the canonical isoform of SPOP (Fig. 3C), another ubiquitin ligase adaptor protein, shown to degrade AR^57^ and associated co-activators such as TRIM24^58^ and SRC-3^59^.

Beyond these specific findings, we see the principal contribution of this work as methodological: demonstrating that a sufficiently large and diverse set of independently generated RNA-sequencing cohorts can be harmonized to transcript resolution without sacrificing the biological heterogeneity needed to resolve disease stages, and that this harmonized structure can then be mined systematically for isoform-pathway associations rather than relying on gene-by-gene literature curation. To make this approach usable beyond our own analyses, we built the Prostate Cancer Atlas web tool (www.prostatecanceratlas.org), which lets researchers explore transcript-and gene-level expression across disease stages and, by reprocessing their own external cohorts through the same pipeline, contextualize new prostate cancer transcriptomic data within this disease-progression framework — either directly through the tool or using the underlying pipeline and code.

## Limitations of the study

Transcript-level quantification in this atlas is inferred from short-read RNA sequencing, which cannot always unambiguously resolve individual isoforms; although we observed good concordance with long-read sequencing in a panel of prostate cancer cell lines, this agreement was weaker for lncRNAs, likely reflecting their frequent lack of poly(A) tails, and emerging approaches that jointly leverage short-and long-read data may further improve isoform-level accuracy in future iterations of the atlas^60^.

The associations we describe between transcript expression and pathway or signature activity are correlative and derived from bulk tissue, which cannot distinguish cell-intrinsic isoform switching from shifts in cellular composition; single-cell and long-read approaches will be needed to resolve this further^9^.

Finally, the insights derived from the atlas are principally descriptive, and although we provide direct functional validation for RNF43, BRCA1, and TRIM11, this validation is deliberately limited and punctual — intended to demonstrate how a diverse, harmonized transcriptome resource can nominate and help validate transcript function in cancer, rather than to exhaustively characterize each gene. Broader functional and clinical validation, including in additional cell line and patient-derived models, will be required to establish the causal contribution of these isoform switches to prostate cancer progression.

## METHODS

### Study design and clinical transcriptome atlas

#### Clinical RNA-sequencing cohorts and metadata

We assembled a prostate cancer RNA-sequencing atlas spanning normal prostate tissue, primary prostate cancer, hormone-sensitive prostate cancer, and castration-resistant prostate cancer. After removal of ineligible hybrid-capture libraries and duplicate representations identified during cohort harmonization, the gene-level atlas comprised 1,365 samples from 14 studies. Clinical and technical metadata were harmonized to record source dataset, molecular state, anatomical site, pretreatment status, sample-acquisition method, RNA-library type, sequencing layout, and available clinical endpoints. Dataset membership, sample counts, metadata fields, and analysis eligibility are reported in Supplementary Table S1.

Four datasets were represented exclusively by single-end sequencing: NEPC lncRNA (Trento/Cornell/Broad 2015; n = 96), Stelloo et al. (n = 92), EIMB RAS (n = 31), and ANTE (n = 6). These 225 samples were retained for gene-level quality-control analyses but excluded from transcript-level analyses. The paired-end transcript-level atlas therefore comprised 1,140 samples from ten studies. Analyses of de-identified public data were performed in accordance with the access conditions and ethics approvals of the source studies.

#### Uniform processing of clinical short-read RNA sequencing

Raw FASTQ files were uniformly reprocessed with nf-core/rnaseq v3.6 against the GRCh38 primary assembly and GENCODE v39 annotation. TCGA-PRAD and WCDT-MCRPC alignment files were downloaded with gdc-client v1.6.1 and converted to FASTQ with SAMtools v1.14; other public datasets were retrieved with fasterq-dump v2.11.3. Reads were aligned with STAR v2.6.1d in two-pass mode, and transcript abundance was quantified with Salmon v1.5.2. Salmon estimated counts were imported with tximport using its default count construction; transcript counts were summarized to genes with the transcript-to-gene mapping derived from GENCODE v39.

Counts were converted independently at gene and transcript levels to counts per million (CPM) and transformed as log2(CPM + 1). No global feature filter was applied before generation of the core CPM matrices. Analysis-specific filters are described below. Unless stated otherwise, transcript-level analyses used the 1,140 paired-end samples.

#### Processing comparison and batch-effect assessment

Three processing configurations were compared: count tables supplied by the source datasets; the same supplied tables after ComBat-seq correction; and expression matrices generated by uniform FASTQ reprocessing. ComBat-seq was applied with source dataset as the batch variable and molecular state as the preserved biological group (full_mod = TRUE, shrink = FALSE, shrink.disp = FALSE).

For PCA and sample-correlation heatmaps, log2(CPM + 1) values were used and the 2,000 features with the largest row variance were selected. PCA was performed with prcomp on the transposed expression matrix. Heatmap input features were standardized by row where indicated in the figure legends; Pearson sample correlations were calculated with pairwise-complete observations and clustered by complete linkage. Sample dendrograms were cut into five clusters. Agreement with molecular-state labels was quantified by adjusted Rand index, and mixing of source datasets by cluster-size-weighted Shannon entropy using base-2 logarithms.

#### Molecular-state assignment and pathway activity

Metastatic samples were classified using androgen-receptor (AR) and neuroendocrine (NE) signatures. The AR signature comprised KLK3, KLK2, TMPRSS2, FKBP5, NKX3-1, PLPP1, PMEPA1, PART1, ALDH1A3, and STEAP4; the NE signature comprised SYP, CHGA, CHGB, ENO2, CHRNB2, SCG3, SCN3A, PCSK1, ELAVL4, and NKX2-1. ssGSEA scores were calculated from genes with >3 counts in at least 5% of samples and rescaled within CRPC samples to −1 to 1. Samples with NE score >0.5 were classified as NEPC; those with NE score ≤0.5 and AR score <0 were classified as DNPC; and the remaining CRPC samples were classified as ARPC.

Pathway activity was estimated by single-sample GSEA using the GSVA implementation (method = "ssgsea", mx.diff = TRUE, ssgsea.norm = FALSE). Hallmark, C2 curated/PID, C6 oncogenic, Gene Ontology, and other displayed gene sets were used as specified for each analysis. For the broad pathway matrix, genes detected in >5% of samples were retained.

#### Dimensionality reduction and disease-progression trajectory

Disease progression was represented by a slingshot trajectory fitted to PC1 and PC2 from the gene-level atlas. Samples were partitioned by k-means with three centers, and a single lineage was fitted with start.clus = 3, end.clus = 2, and allow.breaks = FALSE. The first returned lineage pseudotime was used without additional rescaling. For 1,046 samples shared with the preceding atlas version, old and updated pseudotime values were compared by Pearson correlation. Newly incorporated primary and hormone-sensitive samples were projected into the saved reference PCA and trajectory space. Cluster-to-state labels and the k-means seed are not required to interpret the reported trajectory and are not used in downstream statistical definitions.

### Experimental models and biological assays

#### Matched long-read and short-read RNA sequencing

Total RNA was obtained from independent cultures of LNCaP, VCaP, 22Rv1, PC3, and H660 prostate cancer cell lines, with two biological replicates per cell line. RNA quantity and quality were assessed using a Qubit 4.0 fluorometer with the Qubit RNA BR Assay Kit (Q10211; Thermo Fisher Scientific) and a Fragment Analyzer with the RNA Kit (DNF-471; Agilent), respectively.

Kinnex full-length RNA libraries were prepared according to PacBio procedure 103-238-700, Rev08. For each sample, 300 ng total RNA was converted to first-strand cDNA, PCR-amplified, and barcoded with Iso-Seq primers. Equal masses of barcoded cDNA were pooled to 55 ng and amplified in eight parallel Kinnex PCR reactions. Amplified cDNA segments were enzymatically joined to barcoded terminal adapters, followed by nuclease treatment and bead cleanup.

Final libraries were quantified with the Qubit dsDNA HS Assay Kit (Q32854; Thermo Fisher Scientific), and fragment size was assessed with a FEMTO Pulse Genomic DNA 165 kb Kit (FP-1002-0275; Agilent). Sequencing complexes were prepared with a Revio polymerase kit (103-496-900; PacBio) and cleanup beads (102-158-300; PacBio), diluted in SMRT Link v26.1, and loaded at 160 pM by adaptive loading onto a Revio SPRQ sequencing plate-Nx (103-726-200; PacBio) with four SMRT Cell 25M cells (102-202-200; PacBio). HiFi sequencing was performed on a PacBio Revio instrument using a 30-h movie. Procedures from post-extraction library preparation through SMRT Link processing were performed at the Next Generation Sequencing Platform, University of Bern.

Kinnex barcode demultiplexing, cDNA Iso-Seq demultiplexing, read segmentation, and Iso-Seq analysis were performed in SMRT Link v26.1.0.284828. Barcoded samples were not clustered; poly(A) tails were required and trimmed. Reads were mapped to hg38 with GENCODE v39. Filters required minimum CCS predicted Phred accuracy 20, maximum fuzzy-junction difference 5 bp, minimum gap-compressed identity 95%, minimum mapped coverage 99%, and minimum mapped length 50 bp. The downstream concordance scripts used these SMRT Link transcript outputs directly and did not apply an additional isoform-collapse or classification step.

For matched short-read sequencing, poly(A)+ RNA was isolated with the NEBNext Poly(A) mRNA Magnetic Isolation Module (E7490; New England BioLabs). Libraries were prepared with the NEBNext UltraExpress RNA Library Prep Kit for Illumina (E3330S; New England BioLabs) and NEBNext Multiplex Oligos for Illumina, assessed with an Agilent 2100 Bioanalyzer, and sequenced on a SURFSeq 5000 platform (Genemind) as single-end 120-bp reads. Reads were processed with nf-core/rnaseq v3.15.1 using Singularity, STAR–Salmon, the GRCh38 primary assembly, and GENCODE v39.

#### RNF43 TOPFlash reporter assays

HEK293T cells were transiently transfected in 96-well plates using jetPRIME (Polyplus). Each well received 50 ng each of M50 Super 8× TOPFlash (Addgene #12456), pRL-CMV-Renilla (Promega, E2271), and LGR4 plasmid (VectorBuilder), together with 50 ng of each plasmid required for the indicated condition: pCW107 empty vector (Addgene #62511), pcDNA-Wnt3A (Addgene #35908), RNF43-207, or RNF43-208 (VectorBuilder). R-spondin-1 was added at 200 ng/mL 2 h after transfection where indicated. Firefly and Renilla luminescence were measured 24 h after transfection with the Dual-Glo Luciferase Assay System (Promega, E2920). Firefly activity was normalized to Renilla activity within each well and then expressed relative to the empty-vector (EV) condition in the control arm. Data are mean ± SD from five biological replicates. Five prespecified comparisons were assessed by two-sided paired t tests with Šidák family-wise correction.

#### BRCA1-Δ11q cell model and lentiviral delivery

LNCaP cells were used for BRCA1-Δ11q perturbation experiments, and HEK293T cells were used for lentiviral production. The empty-vector control was pCW107 (Addgene #62511). The HA-tagged BRCA1-Δ11q construct was provided by Neil Johnson and corresponds to the construct described in PMID 27197267.

HEK293T cells were seeded at 4 × 10^6 cells per 100-mm dish and incubated overnight at 37°C in 5% CO2. After 24 h, 3 μg expression vector, 2.7 μg pCMV-dR8.2 packaging plasmid, and 0.7 μg pVSV-G envelope plasmid were mixed in 300 μL Opti-MEM I (31985-070; Gibco) with polyethyleneimine (919012; Sigma-Aldrich; 1.25 mM) at 4 μL per μg DNA. Complexes were incubated for 15 min at room temperature and added to the cells. Viral supernatants were collected 48 h later, passed through a 0.45-μm filter, and applied to target cells with 8 μg/mL polybrene (H9268; Sigma-Aldrich) for 72 h. No antibiotic selection was applied. Immunofluorescence, immunoblotting, and RNA sequencing were performed 7 days after infection.

#### BRCA1-Δ11q RNA sequencing

RNA was extracted with the RNeasy Mini Kit (QIAGEN) according to the manufacturer’s instructions and assessed with an Agilent 2100 Bioanalyzer. Three independent biological replicates per condition were analyzed. Poly(A)+ RNA isolation, library preparation, and sequencing were performed as described above for the matched short-read samples, using NEBNext E7490 and E3330S reagents and single-end 120-bp sequencing on a SURFSeq 5000 instrument. Reads were processed with nf-core/rnaseq v3.15.1 using the Singularity profile, STAR–Salmon, GRCh38 primary assembly, and GENCODE v39.

#### BRCA1 immunofluorescence and immunoblotting

For immunofluorescence, 13-mm coverslips in 24-well plates were coated overnight with poly-D-lysine. LNCaP cells (1 × 10^5 per well) were seeded for 48 h, fixed in 10% formalin for 8–10 min, washed with 0.05% Tween 20, permeabilized with 0.1% Triton X-100 for 20 min, and blocked with DAKO Protein Block Serum-Free (X0909) for 15 min. HA-tagged BRCA1-Δ11q was detected with rabbit anti-HA H6908 (Sigma-Aldrich; RRID: AB_260070; 1:1,000) and Alexa Fluor 488 anti-rabbit secondary antibody (Invitrogen; 1:1,000). Coverslips were mounted with Fluoromount-G containing DAPI and imaged with a Leica THUNDER microscope. The imaging experiment was performed once.

For immunoblotting, snap-frozen pellets were lysed in RIPA buffer containing phosphatase inhibitor (4906845001; Roche) and protease inhibitor (5892953001; Roche). Protein concentration was measured by BCA assay (A52255; Thermo Fisher Scientific). Lysate (50 μg) was resolved on 8% SDS-polyacrylamide gels, transferred to PVDF membrane (88518; Thermo Fisher Scientific), blocked for 30 min at room temperature in 5% milk/TBST, incubated with primary antibodies overnight at 4°C and HRP-conjugated secondary antibodies for 1 h at room temperature, and visualized with WesternBright Quantum (K-12042-D20; Advansta) on a Fusion Solo IV system. Primary antibodies were anti-HA H6908 (Sigma-Aldrich; RRID: AB_260070), anti-BRCA1 MS110 (Millipore; RRID: AB_10682944), and anti-α-tubulin DM1A (Cell Signaling Technology; RRID: AB_1904178); secondary antibodies were Promega W401B and W402B. Antibodies were used at 1:1,000. The immunoblot experiment was performed once.

#### BRCA1-Δ11q BioID labeling and streptavidin pulldown

LNCaP cells (50 × 10^6) were plated and treated the following day with doxycycline (100 ng/mL to 1 μg/mL; D3447; Merck) to induce comparable expression of GFP-BioID control or BRCA1-Δ11q–BioID. Biotin (50 μM; B4639; Merck) was added at induction, and cells were collected 24 h later. Cells were washed twice with PBS and lysed in complete RIPA buffer. Lysates were sonicated with a Bioruptor Plus (BIOSENSE) at high power for 30 cycles of 30 s on and 30 s off, and protein concentration was determined by BCA assay.

For each pulldown, 4 mg protein in 4 mL RIPA buffer was incubated overnight with 50 μL Dynabeads M-280 Streptavidin (11205D; Thermo Fisher Scientific) under rotation. Beads were washed twice with RIPA buffer, twice with 2% SDS, four times with lithium high-salt buffer (50 mM Tris, pH 8.0, 500 mM LiCl, and 2 mM EDTA), four times with sodium high-salt buffer (50 mM Tris, pH 8.0, 500 mM NaCl, and 2 mM EDTA), and twice with low-salt buffer (50 mM Tris, pH 8.0, and 150 mM NaCl). Washed beads were dried and snap-frozen in liquid nitrogen.

Downstream on-bead digestion, liquid chromatography–tandem mass spectrometry (LC–MS/MS), and label-free quantification followed the FOXA1/FOXA2 BioID workflow reported by Formaggio et al. (Cell Reports, 2025; https://doi.org/10.1016/j.celrep.2025.116324). Beads were resuspended in 8 M urea and 50 mM ammonium bicarbonate, reduced with 10 mM dithiothreitol for 60 min at 37°C, and alkylated with 50 mM iodoacetamide for 30 min at room temperature. Proteins were digested on-bead sequentially with 1 μg LysC for 2 h at 37°C and 1 μg trypsin overnight at 37°C. Peptides were purified on C18 StageTips, eluted with 80% acetonitrile and 0.5% acetic acid, dried, and resuspended in 2% acetonitrile, 0.5% acetic acid, and 0.1% trifluoroacetic acid.

For each sample, 1 μg peptide was analyzed on a Q Exactive HF mass spectrometer coupled to an EASY-nLC 1200 system by nanoelectrospray ionization. Peptides were separated on an in-house packed 75-μm-inner-diameter, 50-cm column maintained at 50°C, using a 150-min linear gradient from 5% to 30% buffer B at 250 nL/min (buffer A, 0.1% formic acid; buffer B, 80% acetonitrile and 0.1% formic acid). Data-dependent acquisition comprised survey scans from m/z 300–1,650 at 60,000 resolution (at m/z 200), a maximum injection time of 20 ms, and an automatic gain-control target of 3 × 10^6. The ten most intense ions with charge states 2–5 were isolated in a 1.8-m/z window and fragmented by higher-energy collisional dissociation at normalized collision energy 27. MS/MS scans were acquired at 15,000 resolution with a maximum injection time of 55 ms and automatic gain-control target of 1 × 10^5; dynamic exclusion was 30 s.

Raw files were processed with MaxQuant v1.6.7.0 against the June 2019 human UniProt database supplemented with common contaminants. Peptide-and protein-level false-discovery rates were controlled at 1%. Trypsin/P specificity, up to two missed cleavages, and a minimum peptide length of seven amino acids were used. Carbamidomethylation of cysteine was fixed; protein N-terminal acetylation, methionine oxidation, and lysine biotinylation were variable. Match between runs was enabled with a 0.7-min match window and 20-min alignment window. MaxLFQ quantification used a minimum peptide-ratio count of one. Perseus v1.6.2.3 was used to remove proteins identified only by site, reverse-database matches, and contaminants; LFQ intensities were log2 transformed, and proteins with at least three valid values in one group were retained. Missing values were imputed from a normal distribution with width 0.3 and downshift 1.8. The data export used for this study contained four BRCA1-Δ11q–BioID samples and four GFP-BioID samples. Differential proximity labeling was assessed using the exported two-sided two-sample Student’s t-test p values and Benjamini–Hochberg-adjusted q values as described below.

#### TRIM11–AR cell models and constructs

HEK293T cells were obtained from ATCC, used at early passage (≤5), and tested monthly for mycoplasma with the MycoAlert Mycoplasma Detection Kit (LT07-318; Lonza); all tests were negative. HEK293T and HEK293T-AR cells were maintained at 37°C and 5% CO2 in DMEM (61965-026; Gibco) supplemented with 10% fetal bovine serum and 1% penicillin–streptomycin. For androgen-response experiments, HEK293T-AR cells were transferred to phenol-red-free DMEM (31053-028; Gibco) containing 10% charcoal-stripped serum and 1% penicillin–streptomycin. The AR expression vector pLENTI-AR was obtained from Addgene (#6889). Empty pLV-Puro-RFP-HA control, pLV-Puro-EF1A-TRIM11-201-HA, pLV-Puro-EF1A-TRIM11-204-HA, and pLV-Puro-EF1A-TRIM11-201-ΔSPRY-HA constructs were synthesized by VectorBuilder.

Stable HEK293T-AR cells were generated by PEI-mediated lentiviral production in HEK293T cells using 3 μg pLENTI-AR, 2.7 μg pCMV-dR8.2, and 0.7 μg pVSV-G. Viral supernatant was collected after 48 h, filtered through 0.45 μm, and applied to HEK293T cells with 8 μg/mL polybrene for 72 h. Cells were selected for at least 10 days with 10 μg/mL blasticidin (15205; Sigma-Aldrich).

Transient transfections were performed with jetPRIME (101000046; Polyplus/Sartorius). Cells received 5 μg plasmid DNA per 100-mm dish for immunoblotting and immunoprecipitation or 1 μg per well of a six-well plate for protein-stability assays. Cells were analyzed 24 h after transfection and treated with 10 nM dihydrotestosterone for 6 h before immunoblotting or immunoprecipitation, or for 2 h before cycloheximide treatment.

#### TRIM11 immunoblotting, co-immunoprecipitation, protein stability, and RT-qPCR

Cells were lysed in RIPA buffer supplemented with Roche phosphatase-and protease-inhibitor cocktails. Protein concentration was determined by BCA assay, and 30–50 μg protein was resolved on 10% SDS–PAGE and transferred to PVDF. Membranes were blocked in 5% milk/TBST, incubated with primary antibodies overnight at 4°C and HRP-conjugated secondary antibodies for 1 h at room temperature, and imaged with WesternBright Quantum on a Fusion Solo IV system. Primary antibodies were anti-GAPDH (sc-47724; Santa Cruz Biotechnology), anti-tubulin (3873; Cell Signaling Technology), anti-HSP90 (4877; Cell Signaling Technology), anti-AR (ab133273; Abcam), and anti-HA (H6908; Sigma-Aldrich); secondary antibodies were Promega W401B and W402B.

For co-immunoprecipitation, cells were harvested 24 h after transfection following 6 h of 10 nM dihydrotestosterone and lysed in 50 mM Tris-HCl (pH 7.5), 100 mM NaCl, 0.5% NP-40, 1 mM EDTA, and 10% glycerol with protease and phosphatase inhibitors. One milligram lysate was incubated overnight with 3 μg anti-HA antibody, captured with Dynabeads Protein A (10001D; Invitrogen), washed five times, and eluted in 2× SDS sample buffer before immunoblotting.

For cycloheximide chase, 0.4 × 10^6 HEK293T-AR cells were seeded per well of a six-well plate and transfected with empty vector or TRIM11-201. At 24 h, cells were treated with 10 nM dihydrotestosterone for 2 h, followed by 50 μg/mL cycloheximide with or without 10 μM MG132, and harvested at the indicated time points. AR abundance was normalized to GAPDH using Image Lab v6.1 (Bio-Rad).

For RT-qPCR, total RNA was isolated with the RNeasy Kit (74104; QIAGEN), and one-step reverse transcription and amplification were performed with KAPA SYBR FAST One-Step reagent (KK4652; Sigma-Aldrich). Primers were selected from PrimerBank and GAPDH was the reference gene. Data are mean ± SEM from at least three independent experiments and were analyzed by one-way ANOVA in GraphPad Prism 10.

### Computational and statistical analyses

#### Progression-associated transcripts and canonical-status analyses

Pearson correlations were calculated between log2(CPM + 1) transcript abundance and pseudotime, with gene-level correlations calculated in parallel. Protein-coding transcripts were retained when they contributed >10% of parent-gene expression in at least one sample and had >3 counts in >5% of samples. Progression-associated transcripts were defined by |transcript correlation| >0.5 and |gene correlation| <0.4. lncRNA transcript correlations were analyzed analogously, and transcript structures were plotted in genomic coordinates.

For complementary ordered-state analysis, protein-coding transcript abundance was tested against normal, primary, ARPC, DNPC, and NEPC states encoded 1–5 using Kendall correlation with exact = FALSE in paired-end samples. Tau, the asymptotic test statistic, and nominal two-sided p value were retained; no multiple-testing correction was applied to this descriptive analysis. Canonical status was assigned by version-stripped matching to canonical ids.tsv, treated as the latest source. For the canonical-position density analysis, transcripts were ranked by decreasing Kendall test statistic, canonical relative ranks were calculated, and kernel density was estimated with bandwidth 0.01 against a uniform-density reference.

#### K-mer/LASSO analysis of lncRNA isoforms

For each target lncRNA, transcripts with a non-missing pseudotime correlation and matching sequence were included; at least five transcripts were required. Normalized k-mer frequencies were calculated from transcript sequences, and k-mers present in at least 20% of included transcripts were retained. Gaussian LASSO regression used alpha = 1, seed = 42, and min(10, number of transcripts) cross-validation folds. The lambda.min solution was selected. The four non-zero k-mers with the largest absolute coefficients were mapped to transcript and genomic coordinates. Five-mers were analyzed for FENDRR and 10-mers for FIRRE. No correlation-significance or FDR filter was applied before fitting.

#### Identification and prioritization of pathway-associated isoform switches

Transcript usage was calculated as transcript abundance divided by summed parent-gene abundance. Protein-coding transcripts were eligible when they contributed >40% of parent-gene expression in >5% of samples and had >1 count in >5% of samples. Pearson correlations were calculated between transcript usage and pseudotime or pathway scores. A candidate switch required at least one transcript with r >0.5 and another transcript from the same gene with r <−0.5 for the same target. Retained genes had at least two qualifying transcripts and at least two distinct annotated protein products. The most positively and negatively associated transcripts defined the representative pair, and their correlation difference defined delta correlation. Pathway targets with rescaled score SD ≤10 were excluded.

The 41-gene manuscript subset was used for biological presentation rather than as an additional statistical significance test. It was the union of 34 prespecified cancer-relevant candidates in candidates.txt and the ten genes with the largest representative delta correlations; three genes overlapped, yielding 41 unique genes. The complete computational switch catalogue and sensitivity analyses are provided in Supplementary Tables S4–S5.

#### Isoform dominance, AWSI, and pathway networks

Isoform-dominance plots display expression of the two transcripts in a selected pair, with the diagonal denoting equal expression. For grouped analyses, the expression difference between the designated canonical/longer isoform and alternative/truncated isoform was calculated; the upper and lower quartiles defined dominance groups and the middle 50% was excluded. Displayed ssGSEA scores were compared by two-sided Wilcoxon rank-sum tests.

For isoform pair A and B, the abundance-weighted splicing index (AWSI) was calculated as [CPM_A/(CPM_A + CPM_B + 10−5) − 0.5] × log2(CPM_A + CPM_B + 1). AWSI was correlated with Hallmark pathway scores by Spearman correlation. P values were adjusted by Benjamini–Hochberg, and pathways with FDR ≤0.05 were retained. Approximate 95% confidence intervals were obtained by Fisher z transformation with standard error 1/sqrt(n − 3).

For WNT2B, RNF43, RELA, ROCK2, and TRIM11, significant AWSI–Hallmark associations were represented as networks. Node color denotes the Spearman coefficient and node size −log10(FDR). Pairwise Hallmark gene-set overlap was quantified by Jaccard index; edges with Jaccard similarity ≥0.05 were retained. Communities were identified by weighted Louvain clustering with a fixed layout seed. Community labels are descriptive and were not used as statistical filters.

#### RNF43 score-dependent variability

Samples were sorted separately by RNF43-208 or RNF43-207 abundance. A centered rolling standard deviation of the original WNT_UP.V1_UP score was calculated using a window of max(50, floor[0.05 × n]) samples and a one-sample step. The displayed curves compare variability under the two isoform-specific orderings.

#### Long-read/short-read concordance

Long-read and Salmon-derived short-read measurements were converted to log2(CPM + 1). Transcript and gene version suffixes were removed where required, duplicate identifiers were aggregated, and comparisons were restricted to intersecting features. Protein-coding and lncRNA classes were assigned from the GENCODE v39-derived annotation table. Pearson and Spearman correlations were calculated for two biological replicates each of LNCaP, VCaP, 22Rv1, PC3, and H660; Pearson statistics are displayed. Target-isoform panels used the same matched matrices.

#### BRCA1 atlas and perturbation RNA-seq analyses

BRCA1-203 (ENST00000357654.9) was treated as canonical/full-length BRCA1-C, and BRCA1-201/Δ11q (ENST00000352993.7) as truncated BRCA1-T. For expression-stratified atlas contrasts, samples in the upper and lower expression quartiles of the indicated transcript were compared and the middle 50% was excluded. Raw gene counts were analyzed with DESeq2 using design ∼ GROUP and independent filtering; genes with Benjamini–Hochberg-adjusted p <0.05 were considered differentially expressed. Genes were ranked by the Wald statistic for Hallmark GSEA with clusterProfiler.

BRCA1-Δ11q perturbation reads were processed with nf-core/rnaseq v3.15.1 using Singularity, a maximum of 16 CPUs and 100 GB memory, STAR–Salmon, GRCh38 primary assembly, and GENCODE v39. Differential expression used raw gene counts and DESeq2 with design ∼ BATCH + TREAT and the 11q-versus-pCW107 contrast. One re-sequenced library replaced an outlier. Variance-stabilizing transformation followed by limma removeBatchEffect while preserving treatment was used only for visualization. Hallmark GSEA used DESeq2 Wald statistics, clusterProfiler, pvalueCutoff = 0.05, eps = 1 × 10−50, and nPermSimple = 10,000.

Figure 5J compares BRCA1-T overexpression with the atlas BRCA1-T high-versus-low signature; Figure 5K compares the same perturbation with the atlas BRCA1-C high-versus-low signature. Version-stripped Ensembl identifiers were matched. Genes were classified as significant in both datasets when adjusted p <0.05 in each. Pearson correlations were calculated for all matched genes and separately for genes significant in both.

#### BRCA1 BioID differential analysis and integrated GO Biological Process network

Processed Perseus output from four BRCA1-T BioID replicate samples and four GFP BioID replicate samples was analyzed at protein level. The archived table contained 1,174 proteins with complete LFQ measurements across all eight samples. Two-group Student’s t test p values, Benjamini–Hochberg q values, BRCA1-T-minus-GFP log2 LFQ differences, and t statistics were taken from the Perseus export. Proteins were classified as BRCA1-T enriched or GFP enriched at q <0.05 and log2 fold change ≥0.5 or ≤−0.5, respectively. Figure 5M plots −log10(p value) against log2 LFQ difference; dashed lines denote ±0.5. Up to eight proteins per direction were labeled by decreasing [−log10(p value) + 0.15 × | log2 fold change|], with BRCA1 retained when present.

To estimate conditional continuous atlas associations, raw gene counts from all 1,365 atlas samples were modeled with DESeq2 using design ∼ BRCA1_C_z + BRCA1_T_z after exclusion of genes with zero total counts. BRCA1_C_z and BRCA1_T_z were standardized transcript-level log2(CPM + 1) values. No dataset, molecular-state, purity, or library covariate was included; coefficients are therefore associations adjusted only for the other BRCA1 isoform and are not interpreted causally. Genes were ranked by the corresponding Wald statistic.

Atlas-T, Atlas-C, experimental BRCA1-T-overexpression, and BRCA1-T-versus-GFP BioID ranked lists were analyzed with fgseaMultilevel against MSigDB 2026.1.Hs C5 GO:BP gene sets using minSize = 15, no upper size limit, eps = 0, nPermSimple = 10,000, and deterministic seeds based on 20260827. For Figure 5L, pathways were retained when Atlas-T and Atlas-C each had FDR <0.10 and opposite NES signs, experimental BRCA1-T-overexpression RNA-seq had FDR <0.10 with NES matching Atlas-T, and at least one significant non-bait BRCA1-T-enriched BioID protein occurred in the BioID GO:BP leading edge. BioID proteins required q <0.05 and log2 fold change ≥0.5. Network edges represent MSigDB GO:BP membership. The displayed network contains 19 pathways, 14 non-bait proteins, and 38 edges; shaded modules are descriptive groupings.

#### Survival analysis

Primary-tumor panels used progression-free survival fields PFS_Time and PFS_Event; CRPC panels used overall-survival fields OS_Time and OS_Event. Samples with missing expression, missing event status, missing survival time, or time ≤0 were excluded. Displayed primary analyses included 359 evaluable tumors with 35 progression events, and CRPC analyses included 97 evaluable tumors with 96 deaths. For transcripts with a non-zero median, groups were split at the cohort-specific transcript median; for zero-median transcripts, detected and undetected samples were compared. Kaplan–Meier estimates include 95% confidence intervals, censor marks, and risk tables. Groups were compared by nominal two-sided log-rank tests.

#### Transcript and protein structural analysis

Transcript structures and protein-domain diagrams were generated from GENCODE v39 transcript models, Ensembl/biomaRt annotations, UniProt isoform identifiers, and Pfam-derived domain records, with targeted manual domain labels recorded in Supplementary Table S6. Protein structures for the displayed isoforms were obtained from AlphaFold predictions corresponding to the indicated isoform sequences and visualized in UCSF ChimeraX v1.12. Structural predictions were used qualitatively to illustrate domain retention, truncation, and candidate interfaces and were not treated as experimental structural validation.

For the TRIM11–AR model, full-length human TRIM11 (UniProt Q96F44; 468 amino acids) and full-length human AR (UniProt P10275) were submitted to the AlphaFold Server as a two-chain 1:1 job. AlphaFold 3 model AlphaFold-beta-20231127 was run with server-default multiple-sequence alignments, 10 recycling iterations, five returned models, no user-specified seed, and no ligands, metal ions, nucleic acids, or post-translational modifications. The top-ranked model was displayed. In ChimeraX, candidate interface residues were defined by inter-chain atom pairs within 4.0 Å and displayed as sticks; hydrogen bonds detected by hbonds were shown in yellow with radius 0.05. Interface residues were labeled by residue name and sequence position. Confidence metrics were used only as model-quality guidance and the interface is presented as a testable prediction.

The ROCK2 rotary-shadowing electron micrograph in Figure 6D was reproduced from Figure 1e of Truebestein et al. (Nature Communications, 2015) under the Creative Commons Attribution 4.0 International License; the original scale bar is 4 nm.

#### Statistical analysis and computational environment

Statistical tests were two-sided unless explicitly stated. Multiple-testing correction used the Benjamini– Hochberg method where reported. Box plots show medians, interquartile ranges, and whiskers extending to 1.5 × the interquartile range unless otherwise indicated. Exact tests, thresholds, replicate numbers, and display conventions are specified in the corresponding method subsection or figure legend. Analysis scripts, machine-readable source-data files, and a panel-to-code manifest are supplied with the reproducibility package. Versioned workflow software is reported above; a current-workstation package inventory is included for audit purposes but is not represented as the historical environment of every analysis run.

## Data and code availability

The interactive atlas is available at www.prostatecanceratlas.org. Newly generated short-read and long-read RNA-sequencing data have been submitted to the NCBI Sequence Read Archive under BioProject PRJNA1519081; the records are currently private, and reviewer access is available through the NCBI reviewer-access link: https://dataview.ncbi.nlm.nih.gov/object/PRJNA1519081? reviewer=sj4k333m6ocaucrhjc1lusgc5b. The data will be released publicly upon publication. Public source-study identifiers and sample eligibility are provided in Supplementary Table S1. The versioned analysis-code package and processed source-data tables are supplied with the submission for editorial and peer review and will be released through a public repository no later than publication. Processed BRCA1 BioID protein-level results are provided in Supplementary Table S11; raw LC–MS/MS files and associated search and quantification outputs will be made available upon reasonable request and deposited in an appropriate ProteomeXchange repository before publication. Access to controlled source cohorts remains governed by the original data providers.

Figure S1. Quality control and trajectory validation of the integrated prostate cancer atlas. (A and B) PCA of count tables provided by the source datasets, colored by molecular state (A) or dataset (B). (C and D) PCA of the same count tables after ComBat-seq correction with dataset as batch and molecular state as the preserved biological group, colored by molecular state (C) or dataset (D). (E) Treemap showing sample contributions by dataset and molecular state; single-end and hybrid-capture datasets are indicated. (F and G) PCA of counts generated by uniform reprocessing of FASTQ files with nf-core/rnaseq v3.6, colored by molecular state (F) or dataset (G). (H) Sample correlation heatmap of the final atlas, annotated by molecular state, dataset, library type, and library layout. (I and J) Gene-level PCA colored by library layout (I) or library type (J). (K) Gene-level PCA colored by slingshot pseudotime. (L) Comparison of pseudotime for 1,046 samples shared by the preceding and updated atlases (Pearson r = 0.960). (M and N) Gene-level PCA colored by neuroendocrine score (M) or AR score (N).

Figure S2. Gene-level correlation structures in the integrated atlas. (A–C) Sample correlation heatmaps based on the 2,000 most variable genes of all biotypes (A), protein-coding genes (B), or lncRNA genes (C). Annotation tracks indicate dataset, molecular state, library size, body site, library type, AR score, AR expression, AR-response score, neuroendocrine score, stromal score, and immune score. (D and E) Adjusted Rand index for molecular state (D) and weighted entropy for dataset identity (E) across the three feature classes.

Figure S3. Concordance between MAS-Seq long-read and matched short-read transcript quantification in prostate cancer cell lines. (A) Pearson correlations between Kinnex/MAS-Seq long-read and Salmon-quantified short-read expression across LNCaP, VCaP, 22Rv1, PC3, and H660 cells, with two biological replicates per cell line, for all genes, all transcripts, protein-coding genes and transcripts, and lncRNA genes and transcripts. (B–G) Representative LNCaP replicate-1 comparisons for all genes (B), protein-coding genes (C), lncRNA genes (D), all transcripts (E), protein-coding transcripts (F), and lncRNA transcripts (G). Values were transformed as log2(CPM + 1); version suffixes were removed where required, duplicate identifiers were aggregated, and intersecting features were retained. Pearson r and two-sided p values are shown. Kinnex libraries were sequenced on PacBio Revio and processed in SMRT Link v26.1.0.284828 with hg38/GENCODE v39; exact filters are provided in STAR Methods.

Figure S4. Technical and biological structure of the transcript-level prostate cancer atlas. (A–D) PCA of the initial transcript-level dataset colored by library type (A), source dataset (B), library layout (C), or molecular state (D). (E and F) PCA of the filtered transcript-level atlas after removal of all 225 single-end libraries, showing PC1 versus PC2 (E) or PC1 versus PC3 (F), colored by molecular state. The filtered atlas comprised 1,140 paired-end samples from ten datasets. (G) Sample correlation heatmap based on the 2,000 most variable transcripts, with the indicated technical, clinical, lineage, and microenvironment annotations. Counts were transformed as log2(CPM + 1), features were row-wise z-scored, and Pearson sample correlations were clustered by complete linkage.

Figure S5. Clinical annotation and validation of newly incorporated prostate cancer samples. (A–D) PCA of integrated protein-coding and lncRNA transcript expression, colored by pretreatment status (A), sample-acquisition method (B), source dataset (C), or organ/site (D). (E and F) Gene-level PCA projection of newly incorporated primary and hormone-sensitive prostate cancer samples into PC1–PC2 (E) and PC1–PC3 (F) spaces learned from the reference cohort. (G and H) HALLMARK_G2M_CHECKPOINT (G) and HALLMARK_MYC_TARGETS_V1 (H) ssGSEA scores across the indicated disease states. Box plots show medians, interquartile ranges, and 1.5 × interquartile-range whiskers.

Figure S6. Expression, progression association, and sequence features of prostate cancer-associated lncRNA isoforms. (A) PCA of lncRNA transcript expression colored by poly(A)-selected or total-RNA library type. (B–E) Gene-level MALAT1 (B), NEAT1 (C), FIRRE (D), and FENDRR (E) expression across molecular states. (F) Correlation-rank plot of NEAT1 transcript–pseudotime Pearson coefficients. (G) Mean log2(CPM + 1) expression and pseudotime correlation of NEAT1 transcripts; point size denotes mean expression and color denotes adjusted p <0.05. (H) NEAT1 transcript structures ordered and colored by pseudotime correlation. (I) Correlation-rank plot for FIRRE transcripts. (J) Mean expression and pseudotime correlation of FIRRE transcripts. (K) Genomic mapping of the four FIRRE 10-mers with the largest absolute non-zero LASSO coefficients. (L and M) Non-zero coefficients from cross-validated Gaussian LASSO models for FIRRE 10-mers (L) and FENDRR 5-mers (M). K-mers present in at least 20% of transcripts were fitted with alpha = 1, seed = 42, up to 10-fold cross-validation, and lambda.min selection.

Figure S7. Robustness and positional properties of canonical and non-canonical isoform switches. (A) Sample correlation heatmap based on the 2,000 most variable protein-coding transcripts. (B and C) PCA of protein-coding transcripts colored by dataset (B) or library type (C). (D) Kendall tau estimates and asymptotic test statistics for the relationship between log2(CPM + 1) transcript abundance and ordered molecular state (normal, primary, ARPC, DNPC, and NEPC), stratified by canonical annotation. Analyses were restricted to paired-end samples; p values were two-sided and were not adjusted for multiple testing. (E) Density of canonical transcript positions after ranking protein-coding transcripts by decreasing Kendall test statistic. Relative ranks were estimated with bandwidth 0.01; the dashed line denotes a uniform-density reference. (F) Hallmark enrichment among genes classified as non-canonical driven, defined by a non-canonical isoform with |r| >0.4 when the canonical transcript was neither the most positive nor most negative isoform. (G) Hallmark enrichment among canonical-driven genes, defined by a canonical transcript that was the most positive or negative isoform with |r| >0.4. Over-representation results are shown at nominal p <0.05 and q <0.20.

Figure S8. Sensitivity of pathway-associated isoform-switch detection to pathway variability and disease compartment. (A) Numbers of isoform switches detected for the indicated C2, C6, and Hallmark gene sets. Point color denotes the disease compartment with greatest influence and point size denotes average delta correlation. (B) Number of detected switches plotted against the standard deviation of each gene-set score. Gene sets with score standard deviation <10 were excluded from downstream analysis.

Figure S9. Independent expression and pathway analyses support divergent WNT2B and RNF43 isoform associations. (A) WNT2B isoform expression measured by Kinnex/MAS-Seq long-read and Salmon-based short-read sequencing across prostate cancer cell lines (n = 2 biological replicates per cell line). (B) WNT2B-202 and WNT2B-203 expression ratios across PID_WNT_SIGNALING_PATHWAY scores; Pearson correlations are shown. (C) Hallmark pathway network of WNT2B AWSI associations. Nodes represent pathways with AWSI– pathway FDR ≤0.05; node color denotes Spearman correlation and node size denotes −log10(FDR). Edges connect pathways with Jaccard similarity ≥0.05, and shaded regions denote weighted Louvain communities annotated for lipid metabolism, epithelial-to-mesenchymal transition, cell cycle, and inflammation/STAT signaling. (D) RNF43 isoform expression measured by long-and short-read sequencing as in (A). (E) RNF43-208 and RNF43-207 expression ratios across WNT_UP.V1_UP scores. (F) Hallmark pathway network of RNF43 AWSI associations, displayed as in (C); communities include energy metabolism, DNA repair/epithelial-to-mesenchymal transition, and cell-cycle/MYC pathways. (G and H) Original WNT_UP.V1_UP score plotted against RNF43-208 (G) or RNF43-207 (H) expression, with the 10th, 50th, and 90th conditional percentiles. (I and J) Kaplan–Meier estimates of progression-free survival in primary prostate cancer stratified by RNF43-208 (I) or RNF43-207 (J) expression. Shading denotes 95% confidence intervals and p values are from two-sided log-rank tests.

Figure S10. Validation of RELA and BRCA1 isoform-specific pathway and survival associations. (A) RELA isoform expression measured by Kinnex/MAS-Seq and matched short-read sequencing across prostate cancer cell lines (n = 2 biological replicates per cell line). (B) RELA-202 and RELA-226 expression ratios across ESC_J1_UP_EARLY.V1_UP scores; Pearson correlations are shown. (C and D) RELA isoform-dominance plots colored by z-scored JAK_STAT3_SIGNALING (C) or INFLAMMATORY_RESPONSE (D) ssGSEA scores. (E) Hallmark pathway network of RELA AWSI associations. Node color denotes Spearman correlation, node size denotes −log10(FDR), edges denote Jaccard similarity ≥ 0.05, and shaded weighted Louvain communities are annotated for inflammation/STAT signaling, DNA repair, cell-cycle/MYC, lipid metabolism, and cholesterol homeostasis. (F and G) Gene-level BRCA1 and transcript-level BRCA1-C and BRCA1-T expression (F), and their isoform ratios (G), plotted against TBK1.DF_DN scores in the atlas. Gene-level and delta correlations are indicated. (H) Atlas-only Hallmark GSEA comparing samples with BRCA1-T-versus BRCA1-C-associated expression signatures. Circle position and color denote NES, and circle size denotes −log10(adjusted p value). (I) Hallmark GSEA comparing experimental BRCA1-Δ11q/BRCA1-T overexpression with pCW107 empty vector in LNCaP cells (n = 3 independent biological replicates per condition). Circle position and color denote NES, and circle size denotes −log10(adjusted p value). (J and K) Kaplan–Meier estimates of progression-free survival in primary prostate cancer stratified by canonical BRCA1-C (J) or truncated BRCA1-T (K) expression. Shading denotes 95% confidence intervals, tick marks denote censoring, and numbers at risk are shown.

Figure S11. Independent validation of ROCK2 and TRIM11 isoform switches. (A) ROCK2 isoform expression measured by Kinnex/MAS-Seq and matched short-read sequencing across prostate cancer cell lines (n = 2 biological replicates per cell line). (B and C) Gene-level ROCK2 and transcript-level ROCK2-202 and ROCK2-203 expression (B), and their ratios (C), plotted against CYCLIN_D1_KE_.V1_UP scores. (D) Hallmark pathway network of ROCK2 AWSI associations; communities include inflammation/STAT signaling, lipid metabolism, DNA repair/epithelial-to-mesenchymal transition, hypoxia, and E2F targets. (E and F) ROCK2 isoform-dominance plots colored by z-scored HALLMARK_MYOGENESIS (E) or HALLMARK_APICAL_JUNCTION (F) scores. (G) TRIM11 isoform expression measured by long-and short-read sequencing as in (A). (H and I) Gene-level TRIM11 and transcript-level TRIM11-201 and TRIM11-204 expression (H), and their ratios (I), plotted against ESC_V6.5_UP_LATE.V1_DN scores. (J and K) Correlations of TRIM11-201 and TRIM11-204 with ubiquitin-related (J) and autophagy-related (K) pathway scores from the indicated MSigDB collections. (L and M) TRIM11 isoform-dominance plots colored by z-scored REACTOME_SELECTIVE_AUTOPHAGY (L) or GOBP_POSITIVE_REGULATION_OF_PROTEIN_UBIQUITINATION (M) scores. (N) Hallmark pathway network of TRIM11 AWSI associations; communities include cholesterol-homeostasis/mTOR signaling, DNA repair/inflammation, cell-cycle/MYC, and energy metabolism, with androgen response additionally labeled. Network encodings and thresholds are as in Figure S9.

Figure S12. TRIM11 isoform dynamics and androgen-receptor pathway activity in the prostate cancer atlas. (A) Gene-level TRIM11 and transcript-level TRIM11-201 (ENST00000284551.11) and TRIM11-204 (ENST00000493030.8) expression plotted against prostate cancer pseudotime and colored by molecular state. Gray curves and bands denote generalized additive model trends and confidence intervals; the gene-level pseudotime correlation and delta correlation between transcripts are shown. (B–D) HALLMARK_ANDROGEN_RESPONSE (B), ANDROGEN_RECEPTOR_SIGNALING_PATHWAY (C), and REGULATION_OF_ANDROGEN_RECEPTOR_SIGNALING_PATHWAY (D) ssGSEA scores in TRIM11 isoform-dominance groups. Dominance was defined from the TRIM11-201 minus TRIM11-204 expression difference; upper and lower quartiles defined the two groups and the middle half was excluded (n = 121 per group). Boxes show medians and interquartile ranges, whiskers extend to 1.5 × the interquartile range, and points denote samples. Two-sided Wilcoxon rank-sum p values are shown.

### General statistical and nomenclature notes

Unless stated otherwise, each point in atlas analyses represents one biological sample; transcript expression is log2(CPM + 1), and pathway activity was estimated by ssGSEA. Transcript-level analyses excluded all 225 single-end samples and used 1,140 paired-end samples from ten datasets. Primary-tumor survival panels show progression-free survival, whereas CRPC panels show overall survival. AWSI, abundance-weighted splicing index; AR, androgen receptor; ARPC, AR-positive castration-resistant prostate cancer; CI, confidence interval; DNPC, double-negative prostate cancer; FDR, false discovery rate; GSEA, gene set enrichment analysis; NEPC, neuroendocrine prostate cancer; NES, normalized enrichment score; PCA, principal-component analysis; ssGSEA, single-sample gene set enrichment analysis.

## Supporting information

Machine-readable data

Reproducible code

