## Supplementary Figures for "An integrated, transcript-resolution atlas of prostate cancer progression unmasks associations with oncogenic pathways"

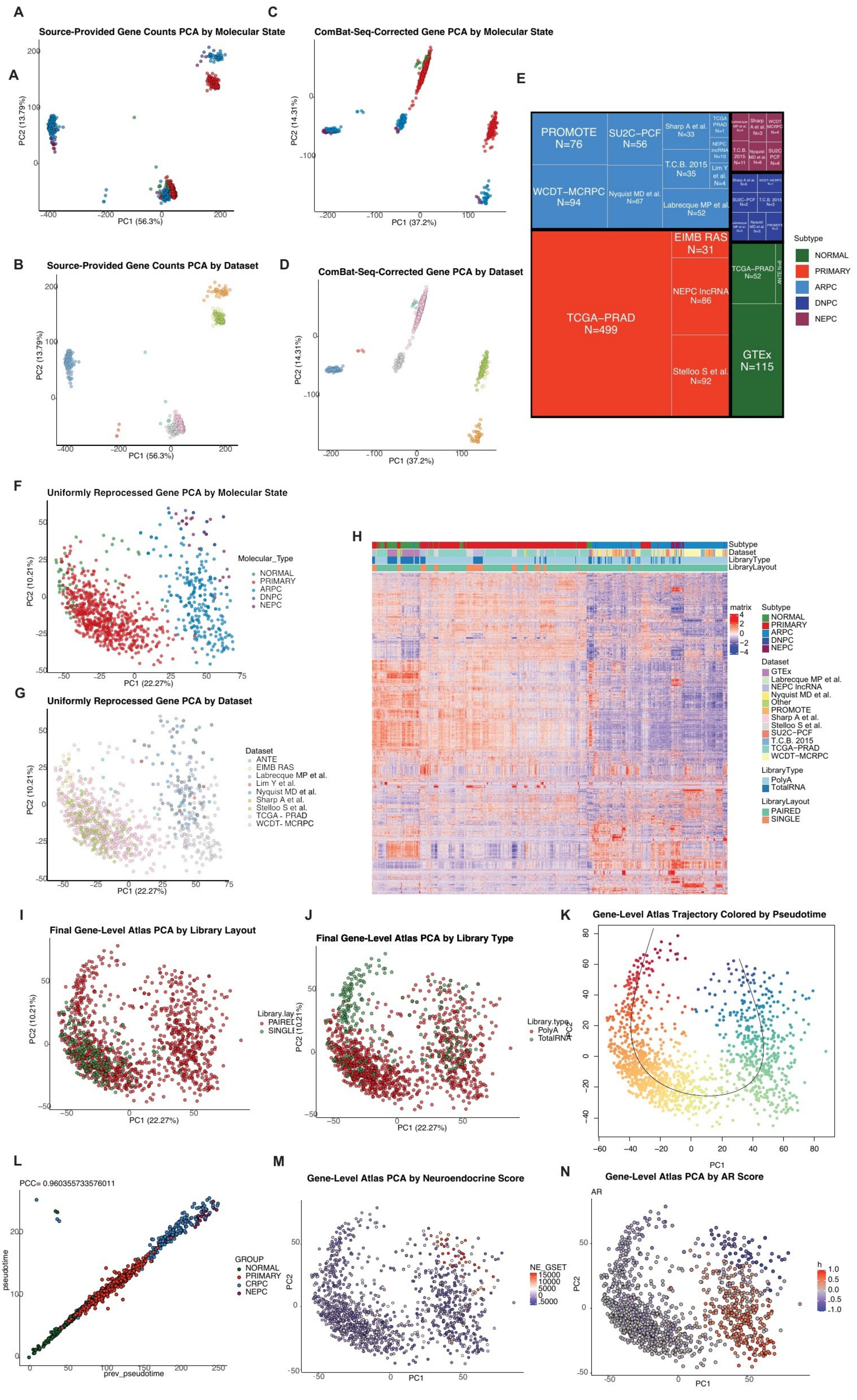

A
Source-Provided Gene Counts PCA by Molecular State
C
ComBat-Seq-Corrected Gene PCA by Molecular State
E
PROMOTE N=76
SU2C-PCF N=56
Sharp A et al. N=33
WCDT-MCRPC N=94
Nyquist MD et al. N=67
T.C.B. 2015 N=35
Labrecque MP et al. N=52
EIMB RAS N=31
TCGA-PRAD N=499
NEPC lncRNA N=86
Stelloo S et al. N=92
TCGA-PRAD N=52
GTEx N=115
Subtype
NORMAL
PRIMARY
ARPC
DNPC
NEPC
B
Source-Provided Gene Counts PCA by Dataset
D
ComBat-Seq-Corrected Gene PCA by Dataset
F
Uniformly Reprocessed Gene PCA by Molecular State
Molecular_Type
G
Uniformly Reprocessed Gene PCA by Dataset
Dataset
ANTE
EIMB RAS
Labrecque MP et al.
Lim Y et al.
Nyquist MD et al.
Sharp A et al.
Stelloo S et al.
TCGA - PRAD
WCDT- MCRPC
H
Subtype
Dataset
LibraryType
LibraryLayout
matrix
GTEx
NEPC lncRNA
Other
PROMOTE
SU2C-PCF
T.C.B. 2015
TCGA-PRAD
WCDT-MCRPC
PolyA
TotalRNA
PAIRED
SINGLE
I
Final Gene-Level Atlas PCA by Library Layout
J
Final Gene-Level Atlas PCA by Library Type
K
Gene-Level Atlas Trajectory Colored by Pseudotime
L
PCC= 0.960355733576011
pseudotime
prev_pseudotime
GROUP
NORMAL
PRIMARY
CRPC
NEPC
M
Gene-Level Atlas PCA by Neuroendocrine Score
NE_GSET
N
Gene-Level Atlas PCA by AR Score
AR
PC1 (56.3%)
PC2 (13.79%)
PC1 (37.2%)
PC2 (14.31%)
PC1 (22.27%)
PC2 (10.21%)
PC1
PC2

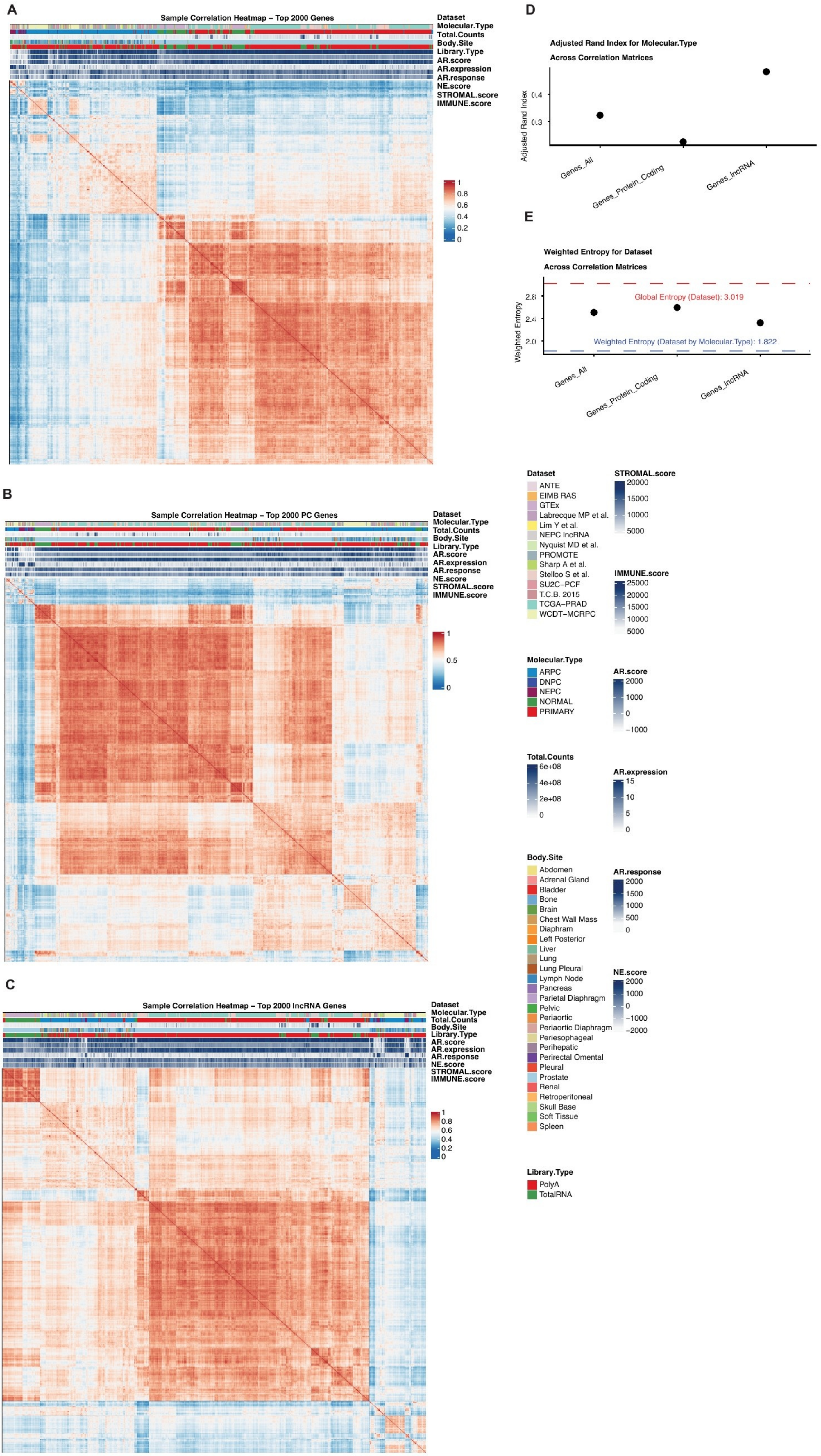
A
Sample Correlation Heatmap – Top 2000 Genes
Dataset
Molecular.Type
Total.Counts
Body.Site
Library.Type
AR.score
AR.expression
AR.response
NE.score
STROMAL.score
IMMUNE.score
1
0.8
0.6
0.4
0.2
0
B
Sample Correlation Heatmap – Top 2000 PC Genes
1
0.5
0
C
Sample Correlation Heatmap – Top 2000 lncRNA Genes
D
Adjusted Rand Index for Molecular.Type
Across Correlation Matrices
Adjusted Rand Index
0.4
0.3
Genes_All
Genes_Protein_Coding
Genes_lncRNA
E
Weighted Entropy for Dataset
Across Correlation Matrices
Weighted Entropy
2.8
2.4
2.0
Global Entropy (Dataset): 3.019
Weighted Entropy (Dataset by Molecular.Type): 1.822
Dataset
ANTE
EIMB RAS
GTEx
Labrecque MP et al.
Lim Y et al.
NEPC lncRNA
Nyquist MD et al.
PROMOTE
Sharp A et al.
Stelloo S et al.
SU2C–PCF
T.C.B. 2015
TCGA–PRAD
WCDT–MCRPC
STROMAL.score
20000
15000
10000
5000
IMMUNE.score
25000
20000
15000
10000
5000
Molecular.Type
ARPC
DNPC
NEPC
NORMAL
PRIMARY
AR.score
2000
1000
0
–1000
Total.Counts
6e+08
4e+08
2e+08
0
AR.expression
15
10
5
0
Body.Site
Abdomen
Adrenal Gland
Bladder
Bone
Brain
Chest Wall Mass
Diaphram
Left Posterior
Liver
Lung
Lung Pleural
Lymph Node
Pancreas
Parietal Diaphragm
Pelvic
Periaortic
Periaortic Diaphragm
Periesophageal
Perihepatic
Perirectal Omental
Pleural
Prostate
Renal
Retroperitoneal
Skull Base
Soft Tissue
Spleen
AR.response
2000
1500
1000
500
0
NE.score
2000
1000
0
–1000
–2000
Library.Type
PolyA
TotalRNA

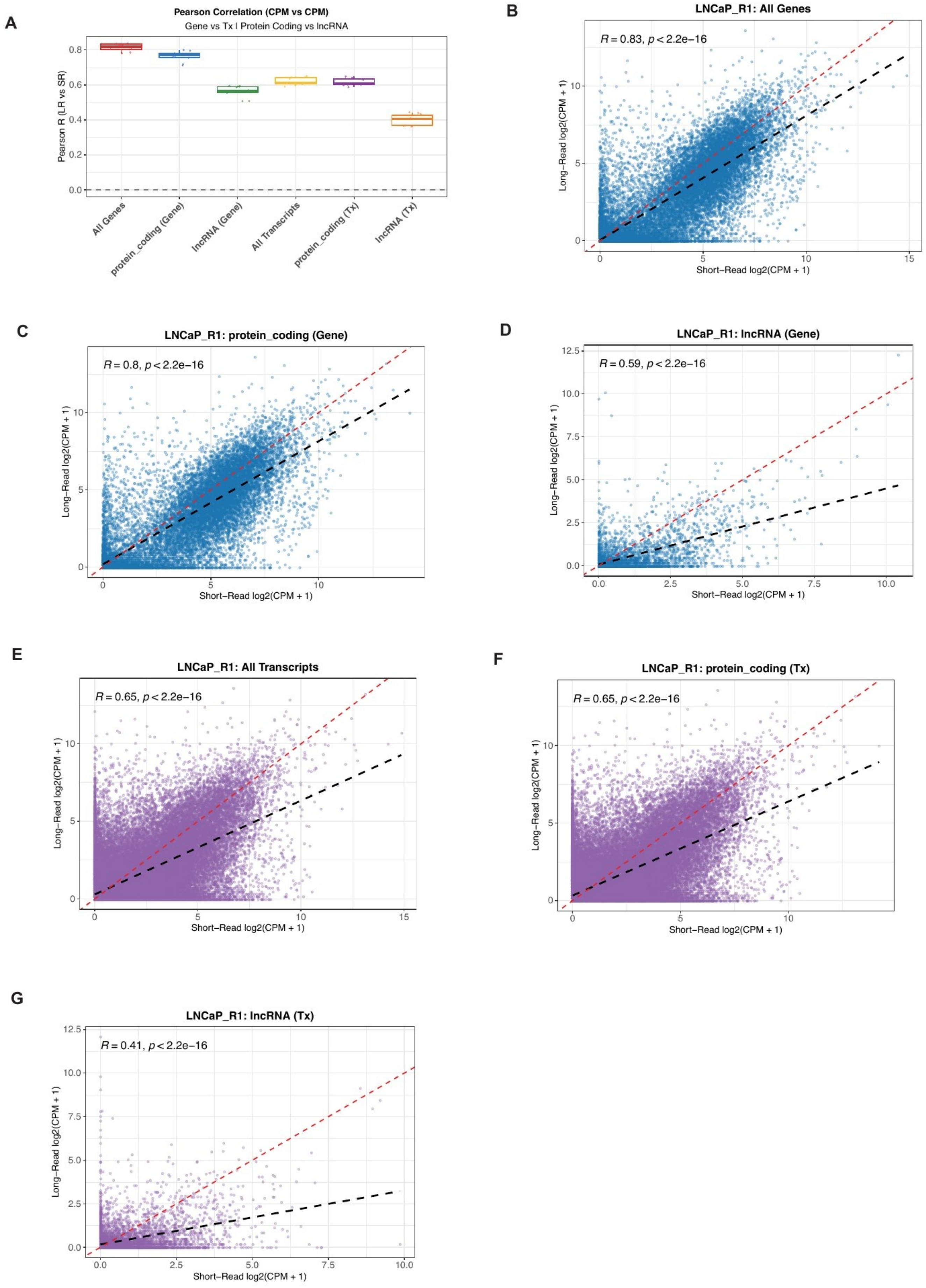

A
Pearson Correlation (CPM vs CPM)
Gene vs Tx | Protein Coding vs lncRNA
Pearson R (LR vs SR)
All Genes
protein_coding (Gene)
lncRNA (Gene)
All Transcripts
protein_coding (Tx)
lncRNA (Tx)
B
LNCaP_R1: All Genes
R = 0.83, p < 2.2e−16
Long−Read log2(CPM + 1)
Short−Read log2(CPM + 1)
C
LNCaP_R1: protein_coding (Gene)
R = 0.8, p < 2.2e−16
Long−Read log2(CPM + 1)
Short−Read log2(CPM + 1)
D
LNCaP_R1: lncRNA (Gene)
R = 0.59, p < 2.2e−16
Long−Read log2(CPM + 1)
Short−Read log2(CPM + 1)
E
LNCaP_R1: All Transcripts
R = 0.65, p < 2.2e−16
Long−Read log2(CPM + 1)
Short−Read log2(CPM + 1)
F
LNCaP_R1: protein_coding (Tx)
R = 0.65, p < 2.2e−16
Long−Read log2(CPM + 1)
Short−Read log2(CPM + 1)
G
LNCaP_R1: lncRNA (Tx)
R = 0.41, p < 2.2e−16
Long−Read log2(CPM + 1)
Short−Read log2(CPM + 1)

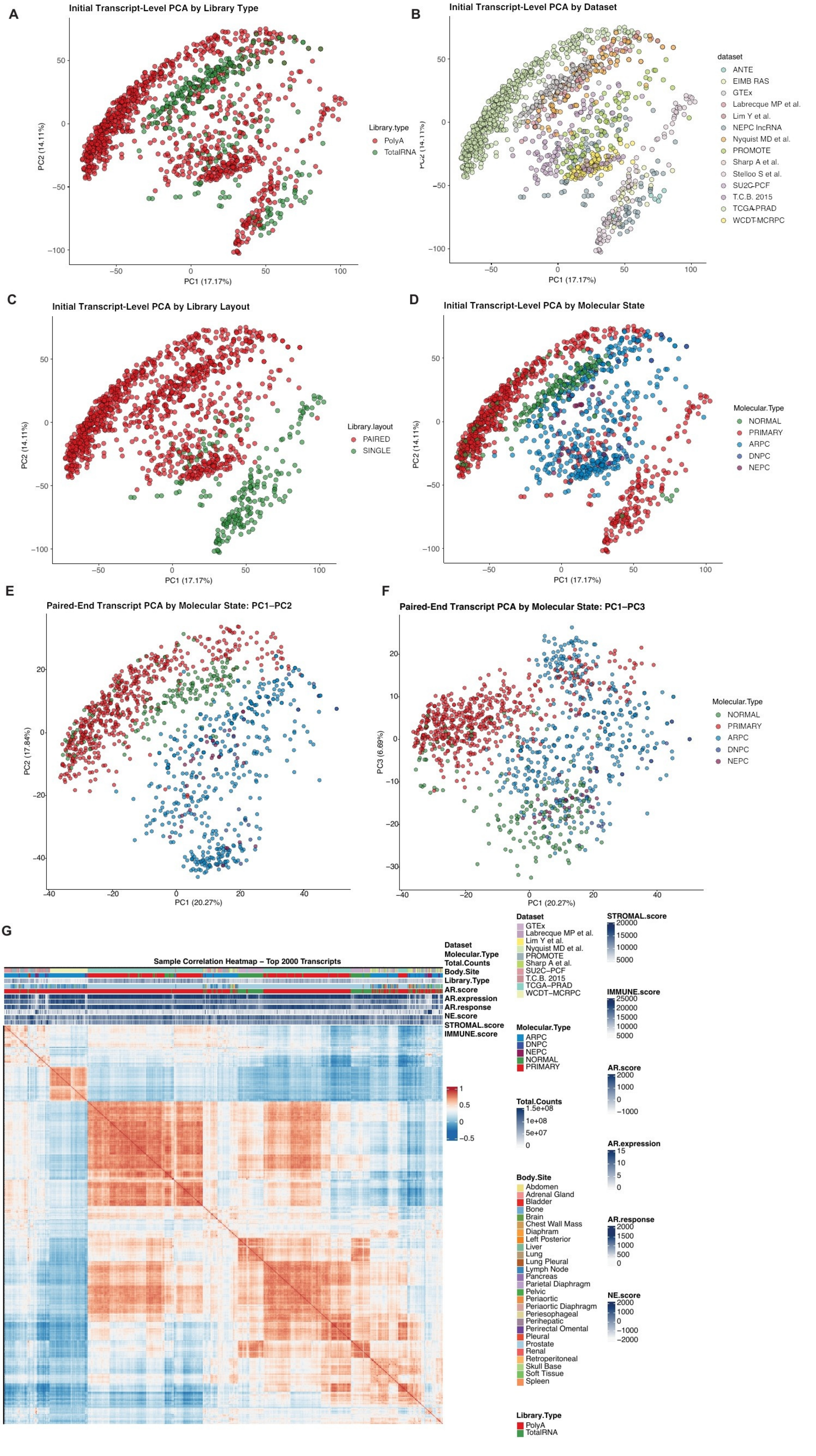

A
Initial Transcript-Level PCA by Library Type
PC1 (17.17%)
PC2 (14.11%)
Library.type
PolyA
TotalRNA
B
Initial Transcript-Level PCA by Dataset
dataset
ANTE
EIMB RAS
GTEx
Labrecque MP et al.
Lim Y et al.
NEPC lncRNA
Nyquist MD et al.
PROMOTE
Sharp A et al.
Stelloo S et al.
SU2C-PCF
T.C.B. 2015
TCGA-PRAD
WCDT-MCRPC
C
Initial Transcript-Level PCA by Library Layout
Library.layout
PAIRED
SINGLE
D
Initial Transcript-Level PCA by Molecular State
Molecular.Type
NORMAL
PRIMARY
ARPC
DNPC
NEPC
E
Paired-End Transcript PCA by Molecular State: PC1–PC2
PC1 (20.27%)
PC2 (17.84%)
F
Paired-End Transcript PCA by Molecular State: PC1–PC3
PC3 (6.69%)
G
Sample Correlation Heatmap – Top 2000 Transcripts
Dataset
Molecular.Type
Total.Counts
Body.Site
Library.Type
AR.score
AR.expression
AR.response
NE.score
STROMAL.score
IMMUNE.score
Dataset
GTEx
Labrecque MP et al.
Lim Y et al.
Nyquist MD et al.
PROMOTE
Sharp A et al.
SU2C–PCF
T.C.B. 2015
TCGA–PRAD
WCDT–MCRPC
Molecular.Type
ARPC
DNPC
NEPC
NORMAL
PRIMARY
Total.Counts
Body.Site
Abdomen
Adrenal Gland
Bladder
Bone
Brain
Chest Wall Mass
Diaphram
Left Posterior
Liver
Lung
Lung Pleural
Lymph Node
Pancreas
Parietal Diaphragm
Pelvic
Periaortic
Periaortic Diaphragm
Periesophageal
Perihepatic
Perirectal Omental
Pleural
Prostate
Renal
Retroperitoneal
Skull Base
Soft Tissue
Spleen
Library.Type
PolyA
TotalRNA
STROMAL.score
IMMUNE.score
AR.score
AR.expression
AR.response
NE.score

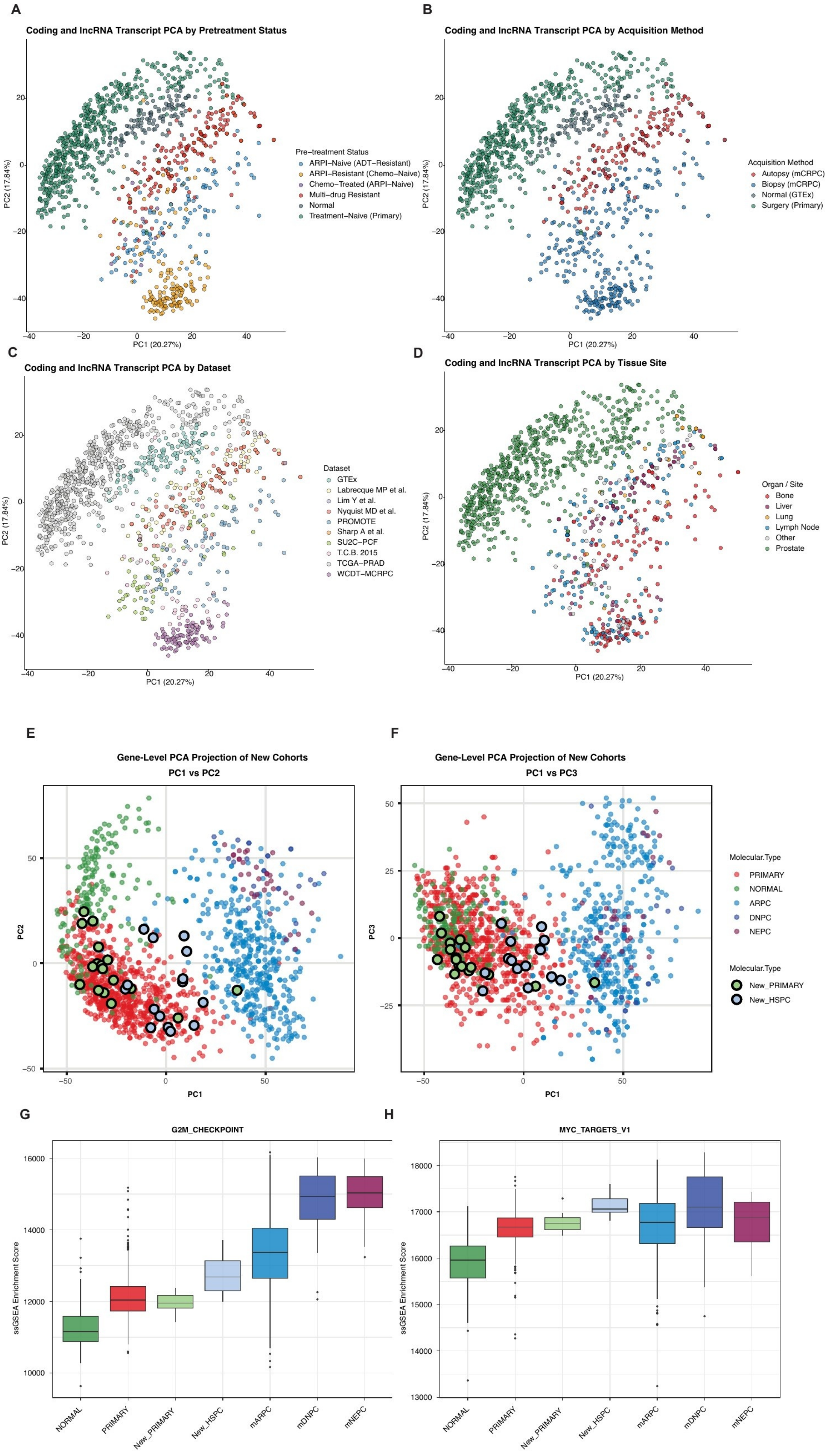
A
Coding and lncRNA Transcript PCA by Pretreatment Status
Pre−treatment Status
ARPI−Naive (ADT−Resistant)
ARPI−Resistant (Chemo−Naive)
Chemo−Treated (ARPI−Naive)
Multi−drug Resistant
Normal
Treatment−Naive (Primary)
PC2 (17.84%)
PC1 (20.27%)
B
Coding and lncRNA Transcript PCA by Acquisition Method
Acquisition Method
Autopsy (mCRPC)
Biopsy (mCRPC)
Normal (GTEx)
Surgery (Primary)
PC2 (17.84%)
PC1 (20.27%)
C
Coding and lncRNA Transcript PCA by Dataset
Dataset
GTEx
Labrecque MP et al.
Lim Y et al.
Nyquist MD et al.
PROMOTE
Sharp A et al.
SU2C−PCF
T.C.B. 2015
TCGA−PRAD
WCDT−MCRPC
PC2 (17.84%)
PC1 (20.27%)
D
Coding and lncRNA Transcript PCA by Tissue Site
Organ / Site
Bone
Liver
Lung
Lymph Node
Other
Prostate
PC2 (17.84%)
PC1 (20.27%)
E
Gene-Level PCA Projection of New Cohorts
PC1 vs PC2
PC2
PC1
F
Gene-Level PCA Projection of New Cohorts
PC1 vs PC3
PC3
PC1
Molecular.Type
PRIMARY
NORMAL
ARPC
DNPC
NEPC
Molecular.Type
New_PRIMARY
New_HSPC
G
G2M_CHECKPOINT
ssGSEA Enrichment Score
NORMAL
PRIMARY
New_PRIMARY
New_HSPC
mARPC
mDNPC
mNEPC
H
MYC_TARGETS_V1
ssGSEA Enrichment Score
NORMAL
PRIMARY
New_PRIMARY
New_HSPC
mARPC
mDNPC
mNEPC

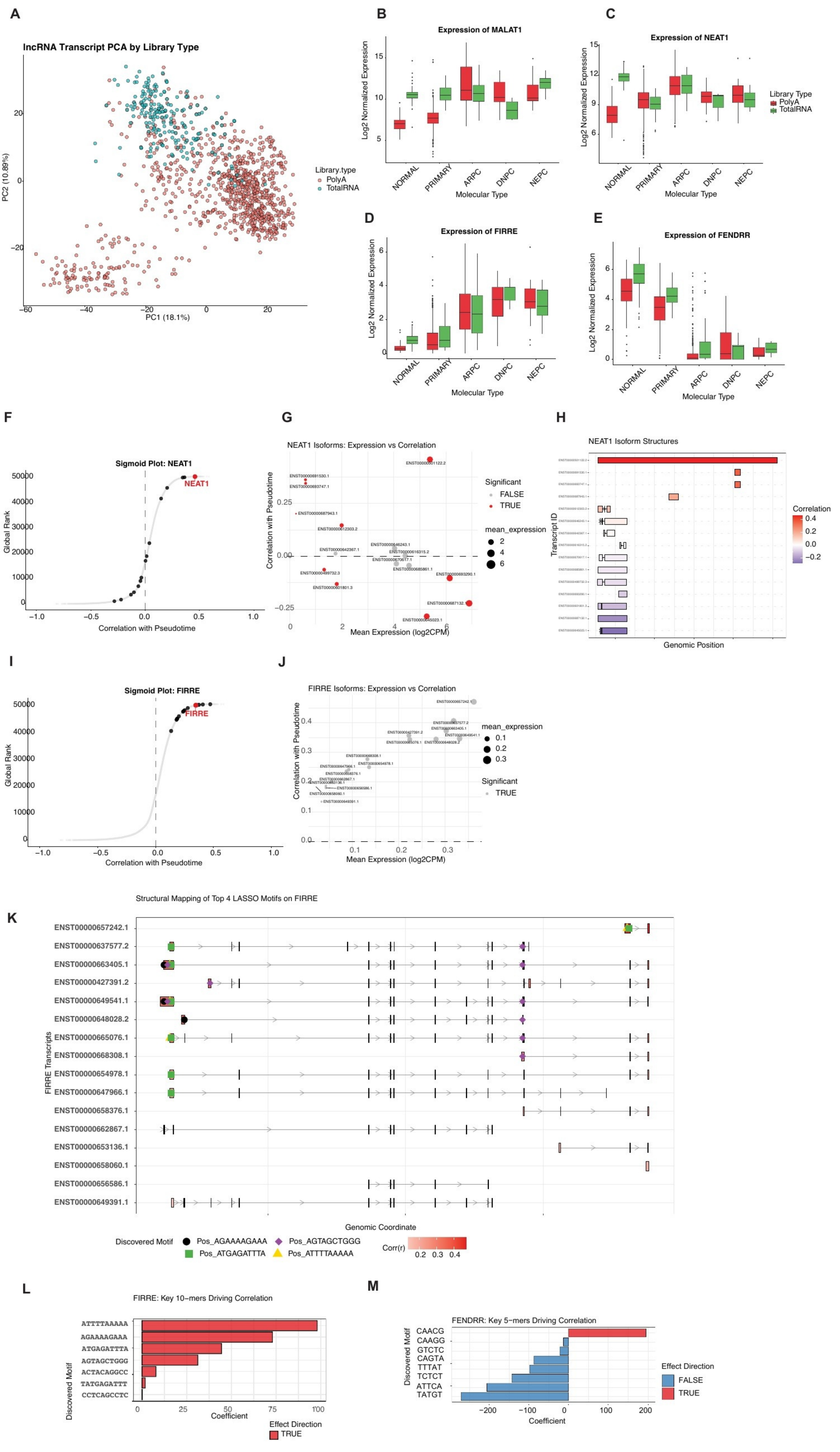
A
lncRNA Transcript PCA by Library Type
PC1 (18.1%)
PC2 (10.89%)
Library.type
PolyA
TotalRNA
B
Expression of MALAT1
Log2 Normalized Expression
Molecular Type
C
Expression of NEAT1
Library Type
PolyA
TotalRNA
D
Expression of FIRRE
E
Expression of FENDRR
NORMAL
PRIMARY
ARPC
DNPC
NEPC
F
Sigmoid Plot: NEAT1
NEAT1
Global Rank
Correlation with Pseudotime
G
NEAT1 Isoforms: Expression vs Correlation
Significant
FALSE
TRUE
mean_expression
Mean Expression (log2CPM)
H
NEAT1 Isoform Structures
Transcript ID
Genomic Position
Correlation
I
Sigmoid Plot: FIRRE
FIRRE
J
FIRRE Isoforms: Expression vs Correlation
K
Structural Mapping of Top 4 LASSO Motifs on FIRRE
ENST00000657242.1
ENST00000637577.2
ENST00000663405.1
ENST00000427391.2
ENST00000649541.1
ENST00000648028.2
ENST00000665076.1
ENST00000668308.1
ENST00000654978.1
ENST00000647966.1
ENST00000658376.1
ENST00000662867.1
ENST00000653136.1
ENST00000658060.1
ENST00000656586.1
ENST00000649391.1
FIRRE Transcripts
Genomic Coordinate
Discovered Motif
Pos_AGAAAAGAAA
Pos_AGTAGCTGGG
Pos_ATGAGATTTA
Pos_ATTTTAAAAA
Corr(r)
L
FIRRE: Key 10-mers Driving Correlation
ATTTTAAAAA
AGAAAAGAAA
ATGAGATTTA
AGTAGCTGGG
ACTACAGGCC
TATGAGATTT
CCTCAGCCTC
Discovered Motif
Coefficient
Effect Direction
TRUE
M
FENDRR: Key 5-mers Driving Correlation
CAACG
CAAGG
GTCTC
CAGTA
TTTAT
TCTCT
ATTCA
TATGT
Effect Direction
FALSE
TRUE

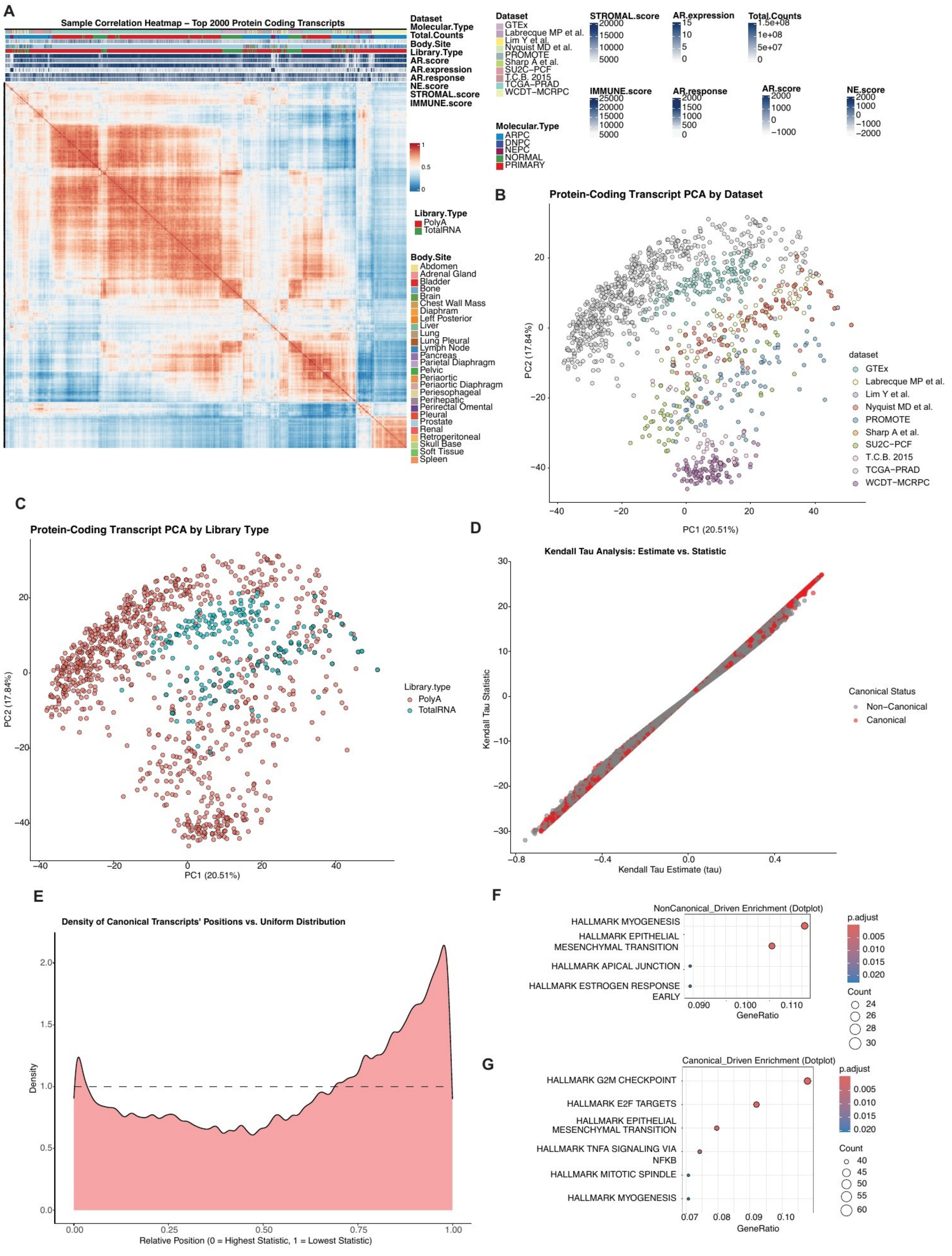
A
Sample Correlation Heatmap – Top 2000 Protein Coding Transcripts
Dataset
Molecular.Type
Total.Counts
Body.Site
Library.Type
AR.score
AR.expression
AR.response
NE.score
STROMAL.score
IMMUNE.score
Dataset
GTEx
Labrecque MP et al.
Lim Y et al.
Nyquist MD et al.
PROMOTE
Sharp A et al.
SU2C–PCF
T.C.B. 2015
TCGA–PRAD
WCDT–MCRPC
Molecular.Type
ARPC
DNPC
NEPC
NORMAL
PRIMARY
STROMAL.score
AR.expression
Total.Counts
IMMUNE.score
AR.response
AR.score
NE.score
Library.Type
PolyA
TotalRNA
Body.Site
Abdomen
Adrenal Gland
Bladder
Bone
Brain
Chest Wall Mass
Diaphram
Left Posterior
Liver
Lung
Lung Pleural
Lymph Node
Pancreas
Parietal Diaphragm
Pelvic
Periaortic
Periaortic Diaphragm
Periesophageal
Perihepatic
Perirectal Omental
Pleural
Prostate
Renal
Retroperitoneal
Skull Base
Soft Tissue
Spleen
B
Protein-Coding Transcript PCA by Dataset
PC1 (20.51%)
PC2 (17.84%)
dataset
C
Protein-Coding Transcript PCA by Library Type
Library.type
PolyA
TotalRNA
D
Kendall Tau Analysis: Estimate vs. Statistic
Kendall Tau Estimate (tau)
Kendall Tau Statistic
Canonical Status
Non–Canonical
Canonical
E
Density of Canonical Transcripts' Positions vs. Uniform Distribution
Density
Relative Position (0 = Highest Statistic, 1 = Lowest Statistic)
F
NonCanonical_Driven Enrichment (Dotplot)
HALLMARK MYOGENESIS
HALLMARK EPITHELIAL MESENCHYMAL TRANSITION
HALLMARK APICAL JUNCTION
HALLMARK ESTROGEN RESPONSE EARLY
GeneRatio
p.adjust
Count
G
Canonical_Driven Enrichment (Dotplot)
HALLMARK G2M CHECKPOINT
HALLMARK E2F TARGETS
HALLMARK EPITHELIAL MESENCHYMAL TRANSITION
HALLMARK TNFA SIGNALING VIA NFKB
HALLMARK MITOTIC SPINDLE
HALLMARK MYOGENESIS
GeneRatio
p.adjust
Count

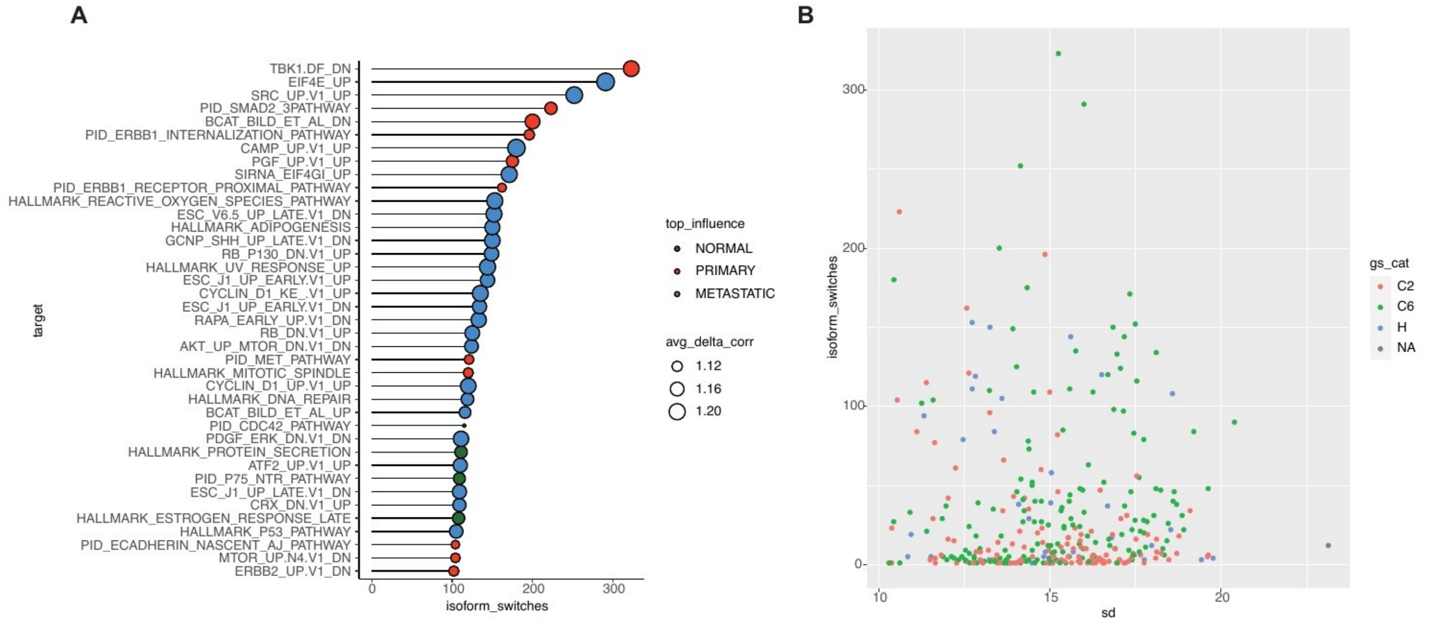
A
TBK1.DF_DN
EIF4E_UP
SRC_UP.V1_UP
PID_SMAD2_3PATHWAY
BCAT_BILD_ET_AL_DN
PID_ERBB1_INTERNALIZATION_PATHWAY
CAMP_UP.V1_UP
PGF_UP.V1_UP
SIRNA_EIF4GI_UP
PID_ERBB1_RECEPTOR_PROXIMAL_PATHWAY
HALLMARK_REACTIVE_OXYGEN_SPECIES_PATHWAY
ESC_V6.5_UP_LATE.V1_DN
HALLMARK_ADIPOGENESIS
GCNP_SHH_UP_LATE.V1_DN
RB_P130_DN.V1_UP
HALLMARK_UV_RESPONSE_UP
ESC_J1_UP_EARLY.V1_UP
CYCLIN_D1_KE_.V1_UP
ESC_J1_UP_EARLY.V1_DN
RAPA_EARLY_UP.V1_DN
RB_DN.V1_UP
AKT_UP_MTOR_DN.V1_DN
PID_MET_PATHWAY
HALLMARK_MITOTIC_SPINDLE
CYCLIN_D1_UP.V1_UP
HALLMARK_DNA_REPAIR
BCAT_BILD_ET_AL_UP
PID_CDC42_PATHWAY
PDGF_ERK_DN.V1_DN
HALLMARK_PROTEIN_SECRETION
ATF2_UP.V1_UP
PID_P75_NTR_PATHWAY
ESC_J1_UP_LATE.V1_DN
CRX_DN.V1_UP
HALLMARK_ESTROGEN_RESPONSE_LATE
HALLMARK_P53_PATHWAY
PID_ECADHERIN_NASCENT_AJ_PATHWAY
MTOR_UP.N4.V1_DN
ERBB2_UP.V1_DN
target
0
100
200
300
isoform_switches
top_influence
NORMAL
PRIMARY
METASTATIC
avg_delta_corr
1.12
1.16
1.20
B
isoform_switches
300
200
100
0
10
15
20
sd
gs_cat
C2
C6
H
NA

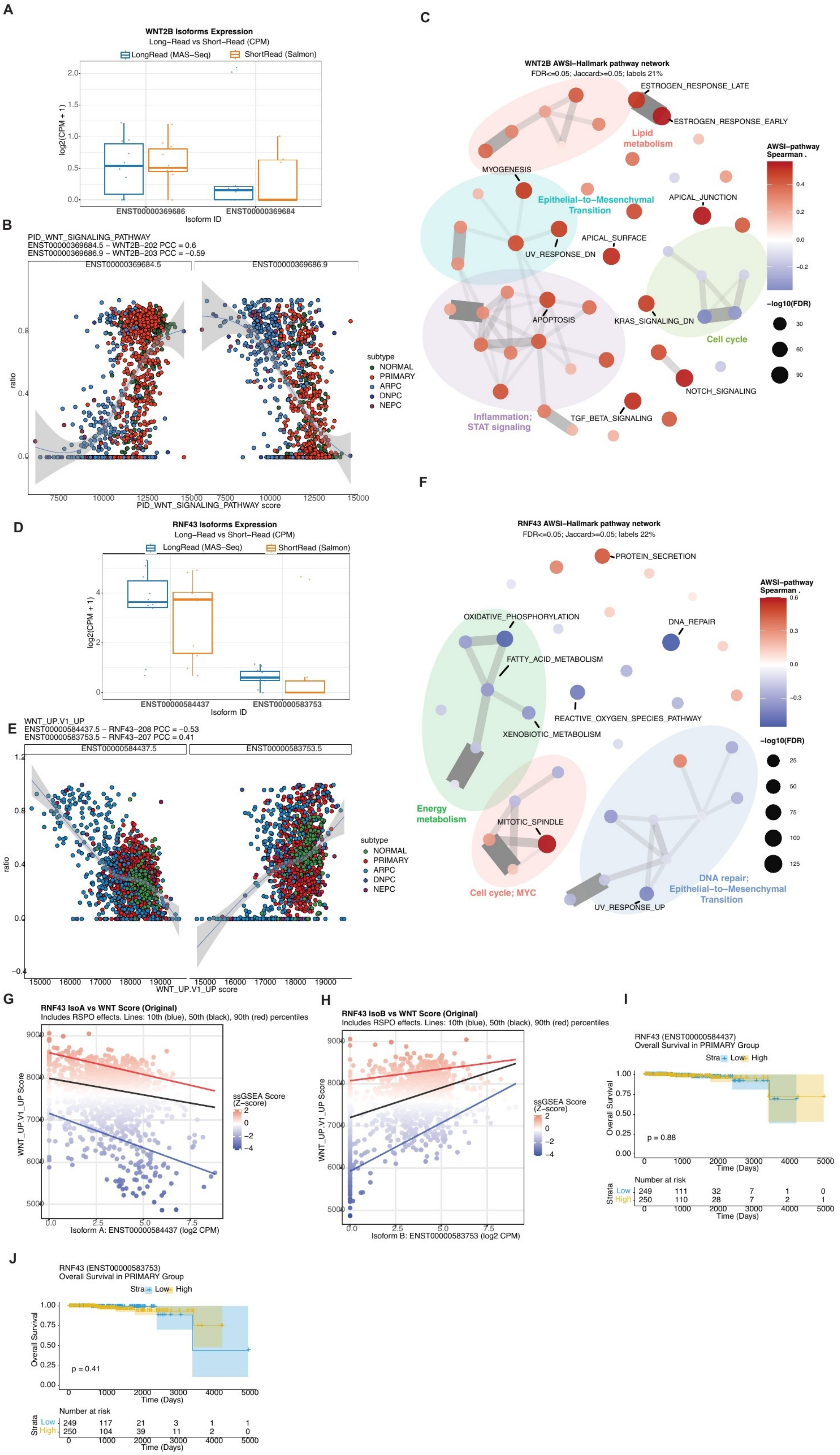
A
WNT2B Isoforms Expression
Long−Read vs Short−Read (CPM)
LongRead (MAS−Seq)
ShortRead (Salmon)
log2(CPM + 1)
ENST00000369686
ENST00000369684
Isoform ID
B
PID_WNT_SIGNALING_PATHWAY
ENST00000369684.5 – WNT2B−202 PCC = 0.6
ENST00000369686.9 – WNT2B−203 PCC = −0.59
ENST00000369684.5
ENST00000369686.9
ratio
PID_WNT_SIGNALING_PATHWAY score
subtype
NORMAL
PRIMARY
ARPC
DNPC
NEPC
C
WNT2B AWSI–Hallmark pathway network
FDR<=0.05; Jaccard>=0.05; labels 21%
ESTROGEN_RESPONSE_LATE
ESTROGEN_RESPONSE_EARLY
Lipid metabolism
MYOGENESIS
Epithelial−to−Mesenchymal Transition
APICAL_JUNCTION
APICAL_SURFACE
UV_RESPONSE_DN
APOPTOSIS
KRAS_SIGNALING_DN
Cell cycle
NOTCH_SIGNALING
TGF_BETA_SIGNALING
Inflammation; STAT signaling
AWSI−pathway Spearman .
−log10(FDR)
D
RNF43 Isoforms Expression
Long−Read vs Short−Read (CPM)
LongRead (MAS−Seq)
ShortRead (Salmon)
log2(CPM + 1)
ENST00000584437
ENST00000583753
Isoform ID
E
WNT_UP.V1_UP
ENST00000584437.5 – RNF43−208 PCC = −0.53
ENST00000583753.5 – RNF43−207 PCC = 0.41
ENST00000584437.5
ENST00000583753.5
ratio
WNT_UP.V1_UP score
subtype
NORMAL
PRIMARY
ARPC
DNPC
NEPC
F
RNF43 AWSI–Hallmark pathway network
FDR<=0.05; Jaccard>=0.05; labels 22%
PROTEIN_SECRETION
OXIDATIVE_PHOSPHORYLATION
DNA_REPAIR
FATTY_ACID_METABOLISM
REACTIVE_OXYGEN_SPECIES_PATHWAY
XENOBIOTIC_METABOLISM
Energy metabolism
MITOTIC_SPINDLE
Cell cycle; MYC
UV_RESPONSE_UP
DNA repair; Epithelial−to−Mesenchymal Transition
AWSI−pathway Spearman .
−log10(FDR)
G
RNF43 IsoA vs WNT Score (Original)
Includes RSPO effects. Lines: 10th (blue), 50th (black), 90th (red) percentiles
WNT_UP.V1_UP Score
Isoform A: ENST00000584437 (log2 CPM)
ssGSEA Score (Z−score)
H
RNF43 IsoB vs WNT Score (Original)
Includes RSPO effects. Lines: 10th (blue), 50th (black), 90th (red) percentiles
WNT_UP.V1_UP Score
Isoform B: ENST00000583753 (log2 CPM)
ssGSEA Score (Z−score)
I
RNF43 (ENST00000584437)
Overall Survival in PRIMARY Group
Strata Low High
Overall Survival
p = 0.88
Time (Days)
Number at risk
Low 249 111 32 7 1 0
High 250 110 28 7 2 1
J
RNF43 (ENST00000583753)
Overall Survival in PRIMARY Group
Strata Low High
Overall Survival
p = 0.41
Time (Days)
Number at risk
Low 249 117 21 3 1 1
High 250 104 39 11 2 0

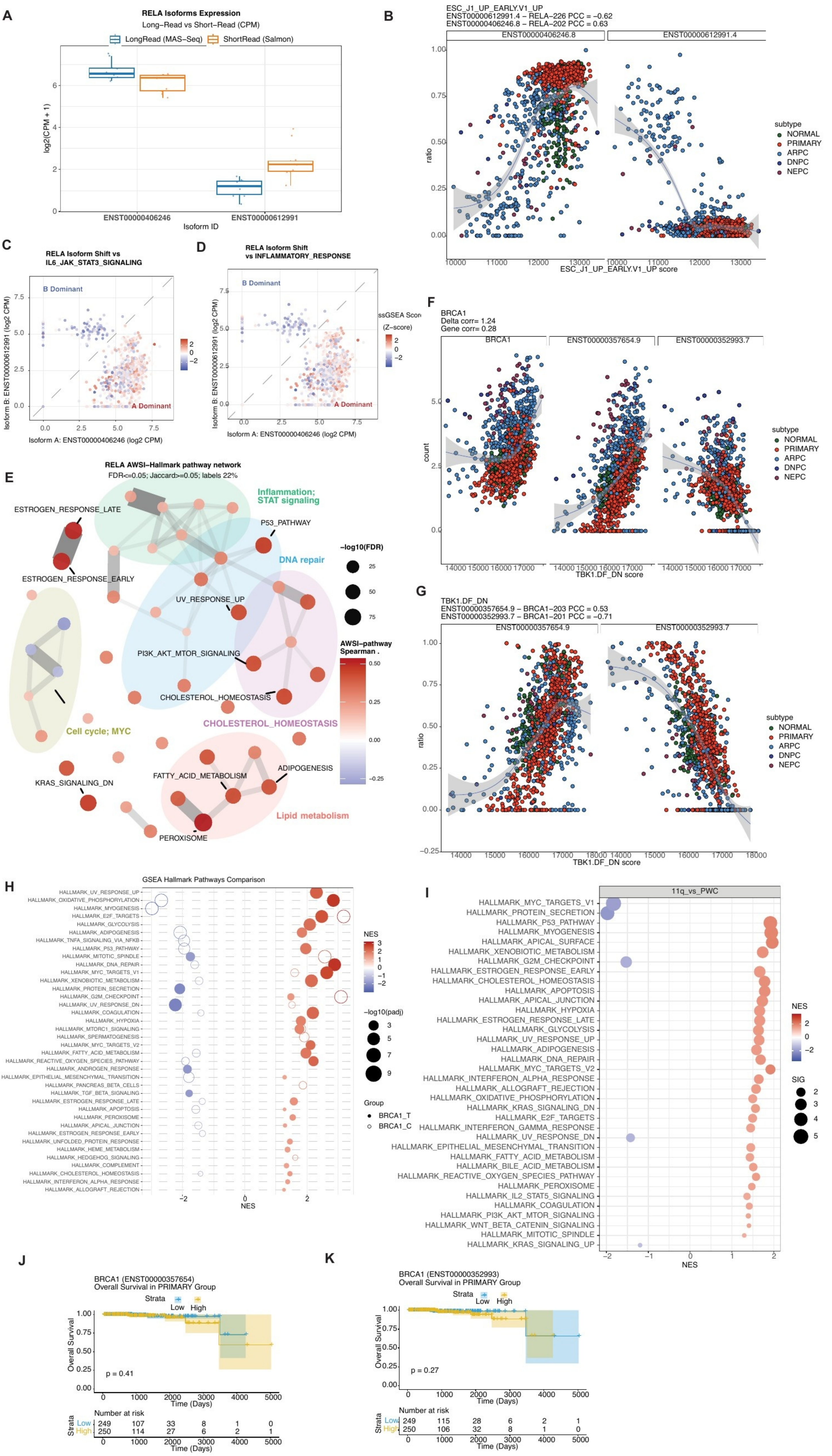

A
RELA Isoforms Expression
Long-Read vs Short-Read (CPM)
LongRead (MAS-Seq)
ShortRead (Salmon)
log2(CPM + 1)
ENST00000406246
ENST00000612991
Isoform ID
B
ESC_J1_UP_EARLY.V1_UP
ENST00000612991.4 – RELA-226 PCC = −0.62
ENST00000406246.8 – RELA-202 PCC = 0.63
ENST00000406246.8
ENST00000612991.4
ratio
ESC_J1_UP_EARLY.V1_UP score
subtype
NORMAL
PRIMARY
ARPC
DNPC
NEPC
C
RELA Isoform Shift vs
IL6_JAK_STAT3_SIGNALING
B Dominant
A Dominant
Isoform B: ENST00000612991 (log2 CPM)
Isoform A: ENST00000406246 (log2 CPM)
D
RELA Isoform Shift
vs INFLAMMATORY_RESPONSE
ssGSEA Score
(Z-score)
F
BRCA1
Delta corr= 1.24
Gene corr= 0.28
BRCA1
ENST00000357654.9
ENST00000352993.7
count
TBK1.DF_DN score
E
RELA AWSI–Hallmark pathway network
FDR<=0.05; Jaccard>=0.05; labels 22%
ESTROGEN_RESPONSE_LATE
ESTROGEN_RESPONSE_EARLY
Inflammation;
STAT signaling
P53_PATHWAY
DNA repair
UV_RESPONSE_UP
PI3K_AKT_MTOR_SIGNALING
CHOLESTEROL_HOMEOSTASIS
Cell cycle; MYC
KRAS_SIGNALING_DN
FATTY_ACID_METABOLISM
ADIPOGENESIS
Lipid metabolism
PEROXISOME
−log10(FDR)
AWSI–pathway
Spearman ρ
G
TBK1.DF_DN
ENST00000357654.9 – BRCA1-203 PCC = 0.53
ENST00000352993.7 – BRCA1-201 PCC = −0.71
H
GSEA Hallmark Pathways Comparison
HALLMARK_UV_RESPONSE_UP
HALLMARK_OXIDATIVE_PHOSPHORYLATION
HALLMARK_MYOGENESIS
HALLMARK_E2F_TARGETS
HALLMARK_GLYCOLYSIS
HALLMARK_ADIPOGENESIS
HALLMARK_TNFA_SIGNALING_VIA_NFKB
HALLMARK_P53_PATHWAY
HALLMARK_MITOTIC_SPINDLE
HALLMARK_DNA_REPAIR
HALLMARK_MYC_TARGETS_V1
HALLMARK_XENOBIOTIC_METABOLISM
HALLMARK_PROTEIN_SECRETION
HALLMARK_G2M_CHECKPOINT
HALLMARK_UV_RESPONSE_DN
HALLMARK_COAGULATION
HALLMARK_HYPOXIA
HALLMARK_MTORC1_SIGNALING
HALLMARK_SPERMATOGENESIS
HALLMARK_MYC_TARGETS_V2
HALLMARK_FATTY_ACID_METABOLISM
HALLMARK_REACTIVE_OXYGEN_SPECIES_PATHWAY
HALLMARK_ANDROGEN_RESPONSE
HALLMARK_EPITHELIAL_MESENCHYMAL_TRANSITION
HALLMARK_PANCREAS_BETA_CELLS
HALLMARK_TGF_BETA_SIGNALING
HALLMARK_ESTROGEN_RESPONSE_LATE
HALLMARK_APOPTOSIS
HALLMARK_PEROXISOME
HALLMARK_APICAL_JUNCTION
HALLMARK_ESTROGEN_RESPONSE_EARLY
HALLMARK_UNFOLDED_PROTEIN_RESPONSE
HALLMARK_HEME_METABOLISM
HALLMARK_HEDGEHOG_SIGNALING
HALLMARK_COMPLEMENT
HALLMARK_CHOLESTEROL_HOMEOSTASIS
HALLMARK_INTERFERON_ALPHA_RESPONSE
HALLMARK_ALLOGRAFT_REJECTION
NES
−log10(padj)
Group
BRCA1_T
BRCA1_C
I
11q_vs_PWC
HALLMARK_MYC_TARGETS_V1
HALLMARK_PROTEIN_SECRETION
HALLMARK_P53_PATHWAY
HALLMARK_MYOGENESIS
HALLMARK_APICAL_SURFACE
HALLMARK_XENOBIOTIC_METABOLISM
HALLMARK_G2M_CHECKPOINT
HALLMARK_ESTROGEN_RESPONSE_EARLY
HALLMARK_CHOLESTEROL_HOMEOSTASIS
HALLMARK_APOPTOSIS
HALLMARK_APICAL_JUNCTION
HALLMARK_HYPOXIA
HALLMARK_ESTROGEN_RESPONSE_LATE
HALLMARK_GLYCOLYSIS
HALLMARK_UV_RESPONSE_UP
HALLMARK_ADIPOGENESIS
HALLMARK_DNA_REPAIR
HALLMARK_MYC_TARGETS_V2
HALLMARK_INTERFERON_ALPHA_RESPONSE
HALLMARK_ALLOGRAFT_REJECTION
HALLMARK_OXIDATIVE_PHOSPHORYLATION
HALLMARK_KRAS_SIGNALING_DN
HALLMARK_E2F_TARGETS
HALLMARK_INTERFERON_GAMMA_RESPONSE
HALLMARK_UV_RESPONSE_DN
HALLMARK_EPITHELIAL_MESENCHYMAL_TRANSITION
HALLMARK_FATTY_ACID_METABOLISM
HALLMARK_BILE_ACID_METABOLISM
HALLMARK_REACTIVE_OXYGEN_SPECIES_PATHWAY
HALLMARK_PEROXISOME
HALLMARK_IL2_STAT5_SIGNALING
HALLMARK_COAGULATION
HALLMARK_PI3K_AKT_MTOR_SIGNALING
HALLMARK_WNT_BETA_CATENIN_SIGNALING
HALLMARK_MITOTIC_SPINDLE
HALLMARK_KRAS_SIGNALING_UP
NES
SIG
J
BRCA1 (ENST00000357654)
Overall Survival in PRIMARY Group
Strata
Low
High
Overall Survival
p = 0.41
Time (Days)
Number at risk
Strata
Low 249 107 33 8 1 0
High 250 114 27 6 2 1
K
BRCA1 (ENST00000352993)
Overall Survival in PRIMARY Group
p = 0.27
Low 249 115 28 6 2 1
High 250 106 32 8 1 0

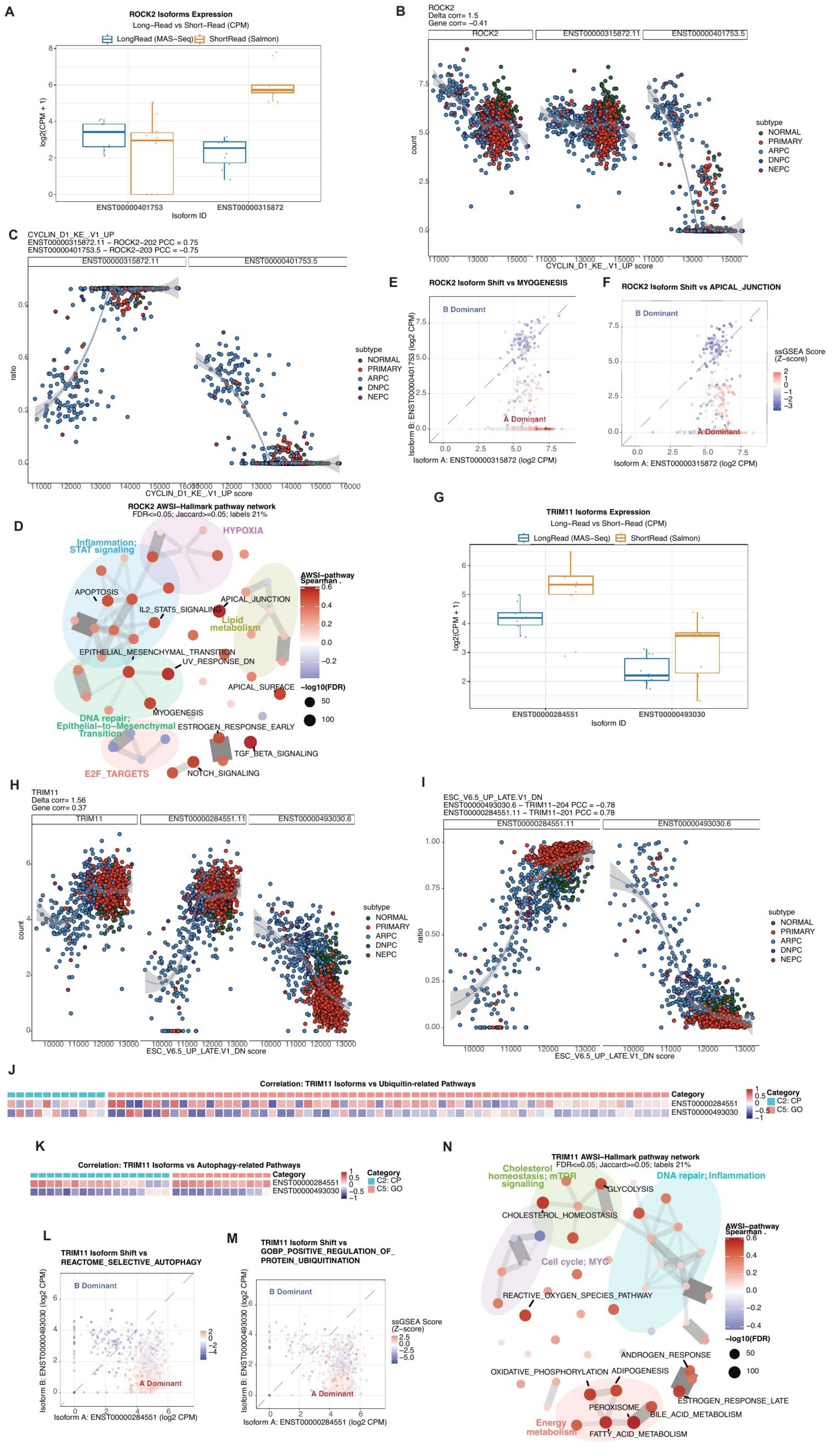

A
ROCK2 Isoforms Expression
Long−Read vs Short−Read (CPM)
LongRead (MAS−Seq)
ShortRead (Salmon)
log2(CPM + 1)
ENST00000401753
ENST00000315872
Isoform ID
B
ROCK2
Delta corr= 1.5
Gene corr= −0.41
ROCK2
ENST00000315872.11
ENST00000401753.5
count
CYCLIN_D1_KE_.V1_UP score
subtype
NORMAL
PRIMARY
ARPC
DNPC
NEPC
C
CYCLIN_D1_KE_.V1_UP
ENST00000315872.11 − ROCK2−202 PCC = 0.75
ENST00000401753.5 − ROCK2−203 PCC = −0.75
ratio
CYCLIN_D1_KE_.V1_UP score
E
ROCK2 Isoform Shift vs MYOGENESIS
B Dominant
A Dominant
Isoform B: ENST00000401753 (log2 CPM)
Isoform A: ENST00000315872 (log2 CPM)
F
ROCK2 Isoform Shift vs APICAL_JUNCTION
ssGSEA Score (Z−score)
D
ROCK2 AWSI−Hallmark pathway network
FDR<=0.05; Jaccard>=0.05; labels 21%
HYPOXIA
Inflammation; STAT signaling
APOPTOSIS
IL2_STAT5_SIGNALING
APICAL_JUNCTION
Lipid metabolism
EPITHELIAL_MESENCHYMAL_TRANSITION
UV_RESPONSE_DN
APICAL_SURFACE
MYOGENESIS
DNA repair; Epithelial−to−Mesenchymal Transition
ESTROGEN_RESPONSE_EARLY
TGF_BETA_SIGNALING
E2F_TARGETS
NOTCH_SIGNALING
AWSI−pathway Spearman
−log10(FDR)
G
TRIM11 Isoforms Expression
Long−Read vs Short−Read (CPM)
ENST00000284551
ENST00000493030
Isoform ID
H
TRIM11
Delta corr= 1.56
Gene corr= 0.37
TRIM11
ENST00000284551.11
ENST00000493030.6
ESC_V6.5_UP_LATE.V1_DN score
I
ESC_V6.5_UP_LATE.V1_DN
ENST00000493030.6 − TRIM11−204 PCC = −0.78
ENST00000284551.11 − TRIM11−201 PCC = 0.78
J
Correlation: TRIM11 Isoforms vs Ubiquitin-related Pathways
Category
C2: CP
C5: GO
K
Correlation: TRIM11 Isoforms vs Autophagy-related Pathways
L
TRIM11 Isoform Shift vs REACTOME_SELECTIVE_AUTOPHAGY
Isoform B: ENST00000493030 (log2 CPM)
Isoform A: ENST00000284551 (log2 CPM)
M
TRIM11 Isoform Shift vs GOBP_POSITIVE_REGULATION_OF_PROTEIN_UBIQUITINATION
N
TRIM11 AWSI−Hallmark pathway network
FDR<=0.05; Jaccard>=0.05; labels 21%
Cholesterol homeostasis; mTOR signalling
GLYCOLYSIS
DNA repair; Inflammation
CHOLESTEROL_HOMEOSTASIS
Cell cycle; MYC
REACTIVE_OXYGEN_SPECIES_PATHWAY
ANDROGEN_RESPONSE
OXIDATIVE_PHOSPHORYLATION
ADIPOGENESIS
ESTROGEN_RESPONSE_LATE
PEROXISOME
BILE_ACID_METABOLISM
Energy metabolism
FATTY_ACID_METABOLISM

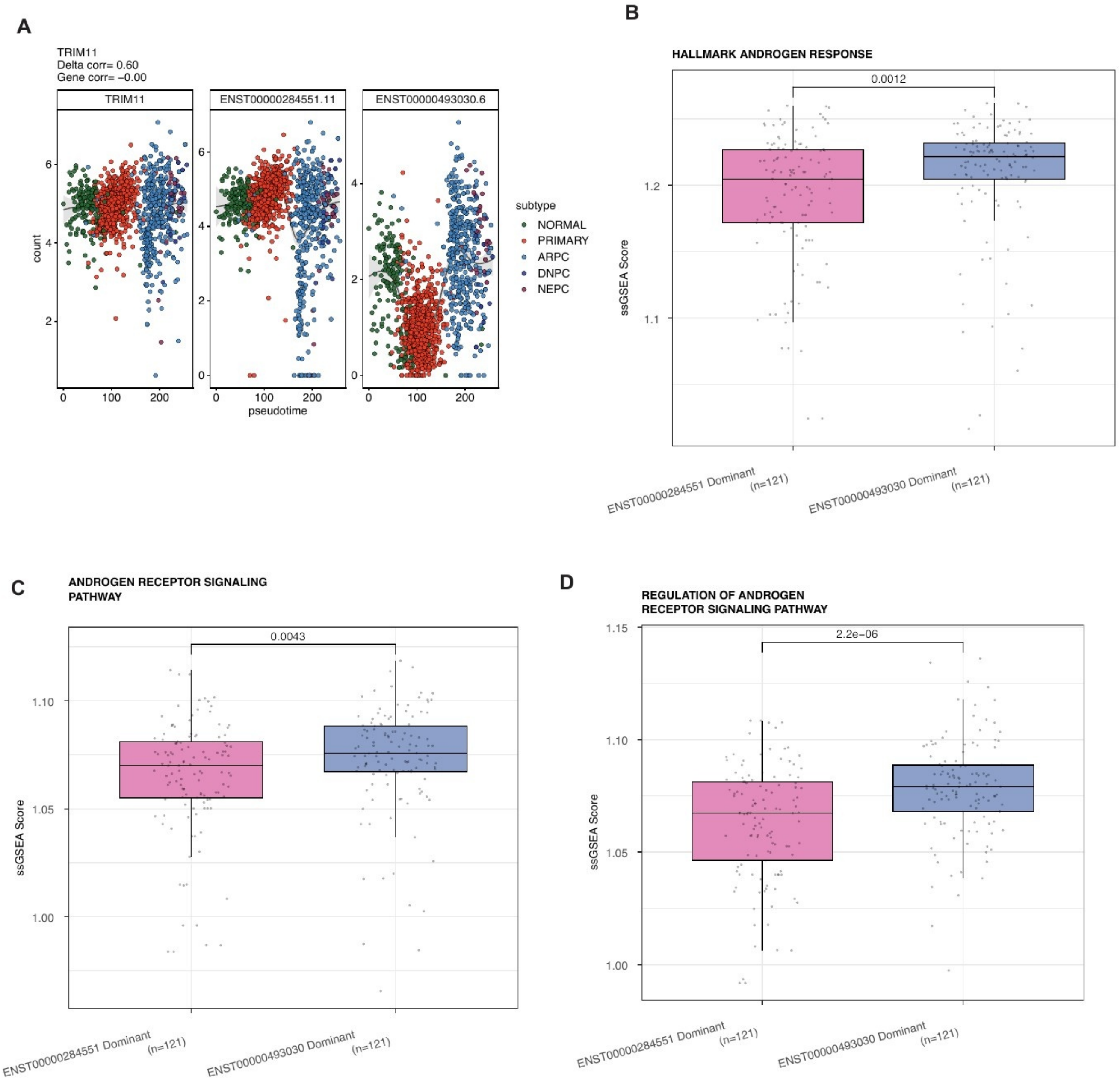
A
TRIM11
Delta corr= 0.60
Gene corr= −0.00
TRIM11
ENST00000284551.11
ENST00000493030.6
count
pseudotime
subtype
NORMAL
PRIMARY
ARPC
DNPC
NEPC
B
HALLMARK ANDROGEN RESPONSE
0.0012
ssGSEA Score
ENST00000284551 Dominant (n=121)
ENST00000493030 Dominant (n=121)
C
ANDROGEN RECEPTOR SIGNALING PATHWAY
0.0043
ssGSEA Score
ENST00000284551 Dominant (n=121)
ENST00000493030 Dominant (n=121)
D
REGULATION OF ANDROGEN RECEPTOR SIGNALING PATHWAY
2.2e−06
ssGSEA Score
ENST00000284551 Dominant (n=121)
ENST00000493030 Dominant (n=121)
