## Supplementary material for "An integrated, transcript-resolution atlas of prostate cancer progression unmasks associations with oncogenic pathways": Reproducible code: README.txt

Compact Hallmark-only AWSI pathway networks

This version retains the same Hallmark pathway nodes, FDR filter and Jaccard
edge definition as the primary Hallmark-only analysis. It changes display only:

  - Fruchterman–Reingold layout for a more compact arrangement;
  - pathway-label candidates require |Spearman rho| >= 0.2 and P <= 0.05;
    the top approximately 22% by |rho| × -log10(P) are labeled, bounded to
    20–40% of nodes in each figure;
  - all other nodes remain plotted without text labels;
  - Louvain community ellipses remain, with concise group labels containing at
    most two representative pathways;
  - ggraph edge widths are linearly rescaled to 0.8–2.0 (a 1.2-unit range);
  - output dimensions are 10 cm × 10 cm with 5-pt plot text.

The thresholds affect labels only; they do not remove nodes, edges or data.
The generating script is:

  ${PROJECT_ROOT}/WNT2B_pathway_network/WNT2B_pathway_network.R
