## Supplementary material for "An integrated, transcript-resolution atlas of prostate cancer progression unmasks associations with oncogenic pathways": Reproducible code: check_package.py

```
#!/usr/bin/env python3
import csv
from pathlib import Path
root = Path(__file__).resolve().parent
for manifest in (root/'manifest/code_manifest.csv', root/'manifest/input_manifest.csv'):
    with manifest.open(encoding='utf-8-sig') as fh:
        rows = list(csv.DictReader(fh))
    missing = [r for r in rows if r.get('located') != 'True']
    print(f'{manifest.name}: {len(rows)} entries; {len(missing)} missing')
    for row in missing:
        print('  ', row.get('original_path'))
```
