## Supplementary material for "An integrated, transcript-resolution atlas of prostate cancer progression unmasks associations with oncogenic pathways": Reproducible code: cor_calc_all.R

```
# Public submission copy.
# Analytical statements and parameter values are unchanged from the archived source.
# Legacy non-English comments and console messages were normalized for public release.
# Load required libraries
library(readr)
library(dplyr)
library(fst)
library(matrixStats)
library(ggplot2)
library(tidyverse)
library(gridExtra)

# Function to load and preprocess data
load_and_preprocess <- function(counts_file, metadata_file, graph_file, transcript_file, transcript_types = NULL,paired) {
  # Load metadata and filter
  mdata <- read.csv(metadata_file, sep = ",")

  id = !is.na(mdata$Library.layout )

  # Filter for PAIRED Library.Layout when using txi.counts.fst
  if (paired == TRUE) {
    id = mdata$Library.layout == "PAIRED"
    mdata = mdata[id,]
  }

  # Load counts and normalize
  txi <- list(counts = as.matrix(read.fst(counts_file)))
  ncount.tx <- log2(sweep(x = txi$counts, MARGIN = 2, STATS = colSums(txi$counts)/1000000, FUN = "/") + 1)
  ncount.tx = ncount.tx[,id]
  # Filter by transcript type if specified
  if (!is.null(transcript_types) && counts_file == "./txi.counts.fst") {
    ids <- read.csv(transcript_file, sep = ",", header = TRUE)
    idx <- ids$transcript_type %in% transcript_types
    ncount.tx <- ncount.tx[idx, ]
  }
  if (!is.null(transcript_types) && counts_file == "./gc_txi.counts.fst") {
    ids <- read.csv("genes_info.csv", sep = ",", header = TRUE)
    idx <- ids$gene_type %in% transcript_types
    ncount.tx <- ncount.tx[idx, ]
  }
  return(list(expr_matrix = ncount.tx, mdata = mdata))
}

calculate_cor <- function(expr_matrix, top_n = NULL) {
  if (!is.null(top_n)) {
    gene_vars <- rowVars(as.matrix(expr_matrix))
    top_genes <- order(gene_vars, decreasing = TRUE)[1:top_n]
    top_expr_matrix <- expr_matrix[top_genes, ]
    sample_cor <- cor(top_expr_matrix, method = "pearson")
  } else {
    sample_cor <- cor(expr_matrix, method = "pearson")
  }
  return(sample_cor)
}


# Main function to run analysis for a given combination
run_analysis <- function(counts_file, transcript_types, top_n, output_prefix,pair) {
  # Create pca_results directory if it doesn't exist
  results_dir <- "cor_results_all"
  if (!dir.exists(results_dir)) {
    dir.create(results_dir)
  }

  # Define file paths
  metadata_file <- "filtered_samples.csv"
  graph_file <- "graph.csv"
  transcript_file <- "transcripts_info.csv"

  # Load and preprocess data
  data <- load_and_preprocess(counts_file, metadata_file, graph_file, transcript_file, transcript_types,pair)


  ##### Calculate Correlation Coefficient####
  tmp <- calculate_cor(data$expr_matrix, top_n)


  # Generate output file names with pca_results path
  counts_type <- ifelse(grepl("gc", counts_file), "gc", "txi")
  trans_type <- if (is.null(transcript_types)) "all" else paste(transcript_types, collapse = "_")
  top_n_str <- if (is.null(top_n)) "all" else as.character(top_n)
  pair_str = if(pair==0) "all" else "paired"

  csv_file <- file.path(results_dir, sprintf("cor_%s_%s_%s_%s.csv", counts_type, trans_type, top_n_str,pair_str))


  # Save CSV
  write.csv(tmp, csv_file, row.names = FALSE)
}


# Define input combinations
counts_files <- c("./txi.counts.fst", "./gc_txi.counts.fst")
transcript_combinations <- list(
  c("lncRNA"),
  c("protein_coding"),
  c("lncRNA", "protein_coding"),
  c("lncRNA"),
  c("protein_coding"),
  NULL # For gc_txi.counts.fst, use all transcripts
)
top_n_values <- list(c(NULL),c(2000))
#top_n_values <- list(c(NULL))
# Run analysis for all combinations
for (counts_file in counts_files) {
  trans_options <- if (grepl("gc", counts_file)) transcript_combinations[4:6] else transcript_combinations[1:3]
  for (trans in trans_options) {
    for (top_n in top_n_values) {
      for(pair in c(1,0)){
        run_analysis(counts_file, trans, top_n,"pca_output",pair)
      }
    }
  }
}


#run_analysis("gc_txi.counts.fst", NULL, NULL,"pca_output",0)
```
