## Supplementary material for "An integrated, transcript-resolution atlas of prostate cancer progression unmasks associations with oncogenic pathways": Reproducible code: heatmap_cor_pseudobulk_norm.R

```
# Public submission copy.
# Analytical statements and parameter values are unchanged from the archived source.
# Legacy non-English comments and console messages were normalized for public release.
library(readr)
library(dplyr)
library(fst, quietly=T)
library(matrixStats)
library(ggplot2)
library(tidyverse)
library(pheatmap)
library(RColorBrewer)

mdata = read.csv("filtered_samples.csv")


txi = list(counts = as.matrix(read.fst("./gc_txi.counts.fst")))

genes = read.csv("genes_info.csv")
# Normalize and log-transform expression matrix
expr_matrix <- txi$counts
rownames(expr_matrix) =sub("\\..*$", "", genes$gene_id)


pseudo_txi = readRDS("./pseudobulk/sketchTumor_SAVER_PseudoMat.rds")


expr_matrix <- as.data.frame(expr_matrix)
pseudo_txi <- as.data.frame(pseudo_txi)

expr_matrix$rownames <- rownames(expr_matrix)
pseudo_txi$rownames <- rownames(pseudo_txi)

merged_data <- merge(expr_matrix, pseudo_txi, by = "rownames", all = TRUE)

rownames(merged_data) <- merged_data$rownames
merged_data$rownames <- NULL

#merged <- log2(sweep(x=merged_data, MARGIN=2, STATS=colSums(merged_data)/1000000, FUN="/") + 1)
merged <- log2(sweep(x=na.omit(merged_data), MARGIN=2, STATS=colSums(na.omit(merged_data))/1000000, FUN="/") + 1)
merged_data = merged
#head(merged_data)


sample_cor <- cor(merged_data, method = "pearson", use = "pairwise.complete.obs")
write.csv(sample_cor, "./pseudobulk/cor.csv",row.names = F)


pseudo_info = read.csv("./pseudobulk/sketchTumor_metaTYPE.csv")

# Select relevant columns from mdata
annotation_df <- mdata %>%
  select(
    dataset, Molecular.Type
  )
colnames(pseudo_info) = colnames(annotation_df)
rownames(pseudo_info) = pseudo_info$dataset
pseudo_info$dataset = "PSEUDO"
rownames(annotation_df) <- paste0("id.", seq_len(nrow(annotation_df)))

annotation_df = rbind(annotation_df,pseudo_info)


annotation_row <- data.frame(
  Dataset = annotation_df$dataset,
  Molecular.Type = annotation_df$Molecular.Type
)

assign_colors <- function(n, palette = "Set3") {
  if (n <= 12) {
    RColorBrewer::brewer.pal(min(n, 12), palette)
  } else {
    colorRampPalette(RColorBrewer::brewer.pal(12, palette))(n)
  }
}

# Colors for Dataset
n_dataset <- length(unique(annotation_row$Dataset))
dataset_colors <- assign_colors(n_dataset, "Set3")

# Colors for Molecular.Type (reusing specified colors)
molecular_type_colors <- c(
  "PRIMARY" = "#e20917",
  "ARPC" = "#088ac8",
  "NEPC" = "#8f1c5b",
  "NORMAL" = "#44984f",
  "DNPC" = "#0000ff",
  "CRPC" = "lightblue"
)


# Annotation colors list
ann_colors <- list(
  Dataset = setNames(dataset_colors, unique(annotation_row$Dataset)),
  Molecular.Type = molecular_type_colors
)

# Load the correlation matrix
# cor_matrix_df <- read.csv("sample_cor_lnc_2000.csv")
# sample_names_cor <- cor_matrix_df[[1]]
# cor_matrix_num <- as.matrix(cor_matrix_df[, -1])
# rownames(cor_matrix_num) <- sample_names_cor
# colnames(cor_matrix_num) <- sample_names_cor
# sample_cor <- cor_matrix_num
colnames(sample_cor) = rownames(annotation_row)
# Plot heatmap
pdf("./pseudobulk/sample_cor_heatmap_gene_all.pdf", width = 10, height = 25)
pheatmap(
  sample_cor,
  annotation_col = annotation_row,
  annotation_colors = ann_colors,
  color = colorRampPalette(rev(brewer.pal(9, "RdBu")))(100),
  clustering_method = "complete",
  show_rownames = FALSE,
  show_colnames = FALSE,
  main = "Sample Correlation Heatmap - All Genes"
)
dev.off()


gene_vars <- rowVars(as.matrix(merged_data))

top_genes <- order(gene_vars, decreasing = TRUE)[1:2000]
top_expr_matrix <- merged_data[top_genes, ]
# Calculate correlation matrix

sample_cor <- cor(top_expr_matrix, method = "pearson", use = "pairwise.complete.obs")

write.csv(sample_cor, "./pseudobulk/cor_2000.csv",row.names = F)


colnames(sample_cor) = rownames(annotation_row)
# Plot heatmap
pdf("./pseudobulk/sample_cor_heatmap_gene_2000.pdf", width = 10, height = 25)
pheatmap(
  sample_cor,
  annotation_col = annotation_row,
  annotation_colors = ann_colors,
  color = colorRampPalette(rev(brewer.pal(9, "RdBu")))(100),
  clustering_method = "complete",
  show_rownames = FALSE,
  show_colnames = FALSE,
  main = "Sample Correlation Heatmap - Top 2000 Genes"
)
dev.off()
```
