## Supplementary material for "An integrated, transcript-resolution atlas of prostate cancer progression unmasks associations with oncogenic pathways": Reproducible code: heatmap_gen_all.R

```
# Public submission copy.
# Analytical statements and parameter values are unchanged from the archived source.
# Legacy non-English comments and console messages were normalized for public release.
# Load required libraries
library(readr)
library(dplyr)
library(fst)
library(matrixStats)
library(ggplot2)
library(tidyverse)
library(gridExtra)
library(pheatmap)
library(RColorBrewer)

draw_heatmap <- function(cor,pair,pdf_file){

  # Load the metadata
  mdata <- read.csv("filtered_samples.csv")

  # Select relevant columns from mdata
  annotation_df <- mdata %>%
    select(
      dataset, Molecular.Type, totalcounts, Body.site, Library.type,
      Primary.gleason.grade, Library.layout, Patient.ethnicity,
      Secondary.gleason.grade, Pharmaceutical.therapy, Radiation.therapy,
      Age.at.diagnosis
    )

  AR_score = read.csv("./../WebToolStudy/AR_value.csv")
  AR_response = read.csv("./../WebToolStudy/AR_RES.csv")
  AR_expression = read.csv("./../WebToolStudy/AR_EXP.csv")
  NE_score = read.csv("./../WebToolStudy/NE_value.csv")
  STROMAL_score = read.csv("./../WebToolStudy/STROMA_value.csv")
  IMMUNE_score = read.csv("./../WebToolStudy/IMMUNE_value.csv")


  # Create annotation data frame
  annotation_row <- data.frame(
    Dataset = annotation_df$dataset,
    Molecular.Type = annotation_df$Molecular.Type,
    Total.Counts = annotation_df$totalcounts, # Keep as numeric
    Body.Site = annotation_df$Body.site,
    Library.Type = annotation_df$Library.type,
    Library.Layout = annotation_df$Library.layout,
    AR.score = as.numeric(AR_score$Score),
    AR.expression = as.numeric(AR_expression$Expression),
    AR.response = as.numeric(AR_response$Score),
    NE.score = as.numeric(NE_score$Score),
    STROMAL.score = as.numeric(STROMAL_score$Score),
    IMMUNE.score = as.numeric(IMMUNE_score$Score)
  )

  # Set row names
  rownames(annotation_row) <- paste0("id.", seq_len(nrow(annotation_row)))

  if (pair == "paired"){
    idx = mdata$Library.layout == "PAIRED"
    annotation_row = annotation_row[idx,]
  }


  # Define colors for annotations
  # Helper function to assign colors for discrete variables
  assign_colors <- function(n, palette = "Set3") {
    if (n <= 12) {
      RColorBrewer::brewer.pal(min(n, 12), palette)
    } else {
      colorRampPalette(RColorBrewer::brewer.pal(12, palette))(n)
    }
  }

  # Colors for Dataset
  n_dataset <- length(unique(annotation_row$Dataset))
  dataset_colors <- assign_colors(n_dataset, "Set3")

  # Colors for Molecular.Type (reusing specified colors)
  molecular_type_colors <- c(
    "PRIMARY" = "#e20917",
    "ARPC" = "#088ac8",
    "NEPC" = "#8f1c5b",
    "NORMAL" = "#44984f",
    "DNPC" = "#0000ff"
  )

  # Colors for Body.site
  n_body_site <- length(unique(annotation_row$Body.Site))
  body_site_colors <- assign_colors(n_body_site, "Paired")

  # Colors for Library.type
  n_library_type <- length(unique(annotation_row$Library.Type))
  library_type_colors <- c(
    "PolyA" = "#e20917",
    "TotalRNA" =  "#44984f")

  # Colors for Primary.gleason.grade
  n_primary_gleason <- length(unique(annotation_row$Primary.Gleason.Grade))
  primary_gleason_colors <- assign_colors(n_primary_gleason, "Set1")

  # Colors for Library.layout
  n_library_layout <- length(unique(annotation_row$Library.Layout))
  library_layout_colors <- c(
    "PAIRED" = "#e20917",
    "SINGLE" =  "#44984f")


  # Colors for Patient.ethnicity
  n_ethnicity <- length(unique(annotation_row$Patient.Ethnicity))
  ethnicity_colors <- assign_colors(n_ethnicity, "Set2")

  # Colors for Secondary.gleason.grade
  n_secondary_gleason <- length(unique(annotation_row$Secondary.Gleason.Grade))
  secondary_gleason_colors <- assign_colors(n_secondary_gleason, "Set1")

  # Colors for Pharmaceutical.therapy
  n_pharma_therapy <- length(unique(annotation_row$Pharmaceutical.Therapy))
  pharma_therapy_colors <- assign_colors(n_pharma_therapy, "Pastel1")

  # Colors for Radiation.therapy
  n_radiation_therapy <- length(unique(annotation_row$Radiation.Therapy))
  radiation_therapy_colors <- assign_colors(n_radiation_therapy, "Pastel2")

  # Colors for continuous variables (Total.Counts and Age.at.Diagnosis)
  # Use a gradient for continuous variables
  continuous_colors <- colorRampPalette(c("#f7f7f7", "#08306b"))(100)

  # Annotation colors list
  ann_colors <- list(
    Dataset = setNames(dataset_colors, unique(annotation_row$Dataset)),
    Molecular.Type = molecular_type_colors,
    Total.Counts = continuous_colors, # Assign the gradient here
    Body.Site = setNames(body_site_colors, unique(annotation_row$Body.Site)),
    Library.Type = setNames(library_type_colors, unique(annotation_row$Library.Type)),

    Library.Layout = setNames(library_layout_colors, unique(annotation_row$Library.Layout)),

    AR.score = continuous_colors,
    AR.response = continuous_colors,
    AR.expression = continuous_colors,
    NE.score = continuous_colors,
    STROMAL.score = continuous_colors,
    IMMUNE.score = continuous_colors
  )

  rownames(cor) = colnames(cor)

  # Plot heatmap


  pdf(pdf_file, width = 10, height = 25)
  pheatmap(
    cor,
    annotation_col = annotation_row,
    annotation_colors = ann_colors,
    color = colorRampPalette(rev(brewer.pal(9, "RdBu")))(100),
    clustering_method = "complete",
    show_rownames = FALSE,
    show_colnames = FALSE,
    main = "Sample Correlation Heatmap"
  )
  dev.off()
}


# Main function to run analysis for a given combination
run_analysis <- function(counts_file, transcript_types, top_n, output_prefix,pair) {
  # Create pca_results directory if it doesn't exist
  results_dir <- "heatmap_results_all"
  if (!dir.exists(results_dir)) {
    dir.create(results_dir)
  }

  # Define file paths
  metadata_file <- "filtered_samples.csv"
  graph_file <- "graph.csv"
  transcript_file <- "transcripts_info.csv"


  # Generate output file names with pca_results path
  counts_type <- ifelse(grepl("gc", counts_file), "gc", "txi")
  trans_type <- if (is.null(transcript_types)) "all" else paste(transcript_types, collapse = "_")
  top_n_str <- if (is.null(top_n)) "all" else as.character(top_n)
  pair_str = if(pair==0) "all" else "paired"
  print(pair_str)
  csv_file_input <- file.path("cor_results_all", sprintf("cor_%s_%s_%s_%s.csv", counts_type, trans_type, top_n_str,pair_str))
  result = read.csv(csv_file_input)
  pdf_file <- file.path("heatmap_results_all", sprintf("heatmap_%s_%s_%s_%s.pdf", counts_type, trans_type, top_n_str,pair_str))

  ##### Calculate Correlation Coefficient####


  draw_heatmap(result, pair_str,pdf_file)


  # Save CSV
  #write.csv(tmp, csv_file, row.names = FALSE)
}


# Define input combinations
counts_files <- c("./txi.counts.fst", "./gc_txi.counts.fst")
transcript_combinations <- list(
  c("lncRNA"),
  c("protein_coding"),
  c("lncRNA", "protein_coding"),
  c("lncRNA"),
  c("protein_coding"),
  NULL # For gc_txi.counts.fst, use all transcripts
)
top_n_values <- list(c(NULL),c(2000))
#top_n_values <- list(c(NULL))
# Run analysis for all combinations
for (counts_file in counts_files) {
  trans_options <- if (grepl("gc", counts_file)) transcript_combinations[4:6] else transcript_combinations[1:3]
  for (trans in trans_options) {
    for (top_n in top_n_values) {
      for(pair in c(1,0)){
        run_analysis(counts_file, trans, top_n,"pca_output",pair)
      }
    }
  }
}


#run_analysis("gc_txi.counts.fst", NULL, NULL,"pca_output",0)
```
