## Supplementary material for "An integrated, transcript-resolution atlas of prostate cancer progression unmasks associations with oncogenic pathways": Reproducible code: measure_cluster_all.R

```
# Public submission copy.
# Analytical statements and parameter values are unchanged from the archived source.
# Legacy non-English comments and console messages were normalized for public release.
library(readr)
library(dplyr)
library(fst, quietly=TRUE)
library(matrixStats)
library(ggplot2)
library(tidyverse)
library(pheatmap)
library(RColorBrewer)
library(dendextend)
library(cluster)
library(mclust)
library(entropy)

# Load the metadata
mdata <- read.csv("filtered_samples.csv")

# Select relevant columns from mdata
annotation_df <- mdata %>%
  select(
    dataset, Molecular.Type, totalcounts, Body.site, Library.type,
    Primary.gleason.grade, Library.layout, Patient.ethnicity,
    Secondary.gleason.grade, Pharmaceutical.therapy, Radiation.therapy,
    Age.at.diagnosis
  )

# Load additional data
AR_score <- read.csv("AR_value.csv")
AR_response <- read.csv("AR_RES.csv")
AR_expression <- read.csv("AR_EXP.csv")
NE_score <- read.csv("NE_value.csv")
STROMAL_score <- read.csv("STROMA_value.csv")
IMMUNE_score <- read.csv("IMMUNE_value.csv")

# Create annotation data frame
annotation_row <- data.frame(
  Dataset = annotation_df$dataset,
  Molecular.Type = annotation_df$Molecular.Type,
  Total.Counts = annotation_df$totalcounts,
  Body.Site = annotation_df$Body.site,
  Library.Type = annotation_df$Library.type,
  Library.Layout = annotation_df$Library.layout,
  AR.score = as.numeric(AR_score$Score),
  AR.expression = as.numeric(AR_expression$Expression),
  AR.response = as.numeric(AR_response$Score),
  NE.score = as.numeric(NE_score$Score),
  STROMAL.score = as.numeric(STROMAL_score$Score),
  IMMUNE.score = as.numeric(IMMUNE_score$Score)
)

# Set row names
rownames(annotation_row) <- paste0("id.", seq_len(nrow(annotation_row)))

# Define colors for annotations
assign_colors <- function(n, palette = "Set3") {
  if (n <= 12) {
    RColorBrewer::brewer.pal(min(n, 12), palette)
  } else {
    colorRampPalette(RColorBrewer::brewer.pal(12, palette))(n)
  }
}

# Colors for Dataset
n_dataset <- length(unique(annotation_row$Dataset))
dataset_colors <- assign_colors(n_dataset, "Set3")

# Colors for Molecular.Type
molecular_type_colors <- c(
  "PRIMARY" = "#e20917",
  "ARPC" = "#088ac8",
  "NEPC" = "#8f1c5b",
  "NORMAL" = "#44984f",
  "DNPC" = "#0000ff"
)

# Colors for Body.site
n_body_site <- length(unique(annotation_row$Body.Site))
body_site_colors <- assign_colors(n_body_site, "Paired")

# Colors for Library.type
n_library_type <- length(unique(annotation_row$Library.Type))
library_type_colors <- c(
  "PolyA" = "#e20917",
  "TotalRNA" = "#44984f"
)

# Colors for Library.layout
n_library_layout <- length(unique(annotation_row$Library.Layout))
library_layout_colors <- c(
  "PAIRED" = "#e20917",
  "SINGLE" = "#44984f"
)

# Colors for continuous variables
continuous_colors <- colorRampPalette(c("#f7f7f7", "#08306b"))(100)

# Annotation colors list
ann_colors <- list(
  Dataset = setNames(dataset_colors, unique(annotation_row$Dataset)),
  Molecular.Type = molecular_type_colors,
  Total.Counts = continuous_colors,
  Body.Site = setNames(body_site_colors, unique(annotation_row$Body.Site)),
  Library.Type = setNames(library_type_colors, unique(annotation_row$Library.Type)),
  Library.Layout = setNames(library_layout_colors, unique(annotation_row$Library.Layout)),
  AR.score = continuous_colors,
  AR.response = continuous_colors,
  AR.expression = continuous_colors,
  NE.score = continuous_colors,
  STROMAL.score = continuous_colors,
  IMMUNE.score = continuous_colors
)

# List of CSV files to process
csv_files <- c(
  "sample_cor_gene.csv", "sample_cor_gene_2000.csv",
  "sample_cor_all.csv", "sample_cor_all_2000.csv", "sample_cor_all_pair_2000.csv",
  "sample_cor_pc.csv", "sample_cor_pc_2000.csv", "sample_cor_pc_pair_2000.csv",
  "sample_cor_lnc.csv", "sample_cor_lnc_2000.csv", "sample_cor_lnc_pair_2000.csv"
)

# Initialize data frame to store metrics
metrics_df <- data.frame(
  CSV_File = character(),
  ARI_Molecular = numeric(),
  Weighted_Entropy_Dataset = numeric(),
  Weighted_Entropy_Library_Type = numeric(),
  Weighted_Entropy_Library_Layout = numeric(),
  stringsAsFactors = FALSE
)


# Process each CSV file
for (csv_file in csv_files) {
  # Load the correlation matrix
  cor_matrix_df <- read.csv(csv_file)
  sample_names_cor <- cor_matrix_df[[1]]
  cor_matrix_num <- as.matrix(cor_matrix_df[, -1])
  rownames(cor_matrix_num) <- sample_names_cor
  colnames(cor_matrix_num) <- sample_names_cor
  sample_cor <- cor_matrix_num

  # Generate heatmap to get dendrogram
  original_heatmap <- pheatmap(
    sample_cor,
    annotation_col = annotation_row,
    annotation_colors = ann_colors,
    color = colorRampPalette(rev(brewer.pal(9, "RdBu")))(100),
    clustering_method = "complete",
    show_rownames = FALSE,
    show_colnames = FALSE,
    main = paste("Sample Correlation Heatmap -", csv_file),
    silent = TRUE
  )

  # Extract dendrogram
  dend <- as.dendrogram(original_heatmap$tree_row)

  # Match annotation_row to dendrogram labels
  labels_dend <- labels(dend)
  annotation_row_matched <- annotation_row[match(labels_dend, rownames(annotation_row)), ]

  # Cut the dendrogram into k=5 clusters
  k <- 5
  clusters <- cutree(dend, k = k, order_clusters_as_data = FALSE)

  # Compute ARI for Molecular.Type
  molecular_type_labels <- as.factor(annotation_row_matched$Molecular.Type)
  ari_molecular <- adjustedRandIndex(clusters, molecular_type_labels)

  # Compute Weighted Entropy for Dataset
  dataset_labels <- as.factor(annotation_row_matched$Dataset)
  cluster_list <- split(dataset_labels, clusters)

  compute_cluster_entropy <- function(cluster_labels) {
    if (length(cluster_labels) == 0) return(0)
    freq_table <- table(cluster_labels) / length(cluster_labels)
    entropy::entropy(freq_table, unit = "log2")
  }

  entropies <- sapply(cluster_list, compute_cluster_entropy)
  weighted_entropy_dataset <- sum(entropies * (sapply(cluster_list, length) / length(dataset_labels)))


  # Compute Weighted Entropy for Library.Type
  library_type_labels <- as.factor(annotation_row_matched$Library.Type)
  cluster_list_library_type <- split(library_type_labels, clusters)
  entropies_library_type <- sapply(cluster_list_library_type, compute_cluster_entropy)
  weighted_entropy_library_type <- sum(entropies_library_type * (sapply(cluster_list_library_type, length) / length(library_type_labels)))

  # Compute Weighted Entropy for Library.Layout
  library_layout_labels <- as.factor(annotation_row_matched$Library.Layout)
  cluster_list_library_layout <- split(library_layout_labels, clusters)
  entropies_library_layout <- sapply(cluster_list_library_layout, compute_cluster_entropy)
  weighted_entropy_library_layout <- sum(entropies_library_layout * (sapply(cluster_list_library_layout, length) / length(library_layout_labels)))

  # Store metrics
  metrics_df <- rbind(metrics_df, data.frame(
    CSV_File = csv_file,
    ARI_Molecular = ari_molecular,
    Weighted_Entropy_Dataset = weighted_entropy_dataset,
    Weighted_Entropy_Library_Type = weighted_entropy_library_type,
    Weighted_Entropy_Library_Layout = weighted_entropy_library_layout
  ))
}


# After the loop for processing CSV files, compute global entropy for Dataset
dataset_labels_all <- as.factor(annotation_row$Dataset)
freq_table_all <- table(dataset_labels_all) / length(dataset_labels_all)
global_entropy <- entropy::entropy(freq_table_all, unit = "log2")


molecular_type_labels_all <- as.factor(annotation_row$Molecular.Type)
cluster_list_molecular <- split(dataset_labels_all, molecular_type_labels_all)
entropies_molecular <- sapply(cluster_list_molecular, compute_cluster_entropy)
weighted_entropy_dataset_by_molecular <- sum(entropies_molecular * (sapply(cluster_list_molecular, length) / length(dataset_labels_all)))


# Compute global entropy for Library.Type
library_type_labels_all <- as.factor(annotation_row$Library.Type)
freq_table_library_type <- table(library_type_labels_all) / length(library_type_labels_all)
global_entropy_library_type <- entropy::entropy(freq_table_library_type, unit = "log2")

# Compute weighted entropy for Library.Type by Molecular.Type
cluster_list_library_type_molecular <- split(library_type_labels_all, molecular_type_labels_all)
entropies_library_type_molecular <- sapply(cluster_list_library_type_molecular, compute_cluster_entropy)
weighted_entropy_library_type_by_molecular <- sum(entropies_library_type_molecular * (sapply(cluster_list_library_type_molecular, length) / length(library_type_labels_all)))

# Compute global entropy for Library.Layout
library_layout_labels_all <- as.factor(annotation_row$Library.Layout)
freq_table_library_layout <- table(library_layout_labels_all) / length(library_layout_labels_all)
global_entropy_library_layout <- entropy::entropy(freq_table_library_layout, unit = "log2")

# Compute weighted entropy for Library.Layout by Molecular.Type
cluster_list_library_layout_molecular <- split(library_layout_labels_all, molecular_type_labels_all)
entropies_library_layout_molecular <- sapply(cluster_list_library_layout_molecular, compute_cluster_entropy)
weighted_entropy_library_layout_by_molecular <- sum(entropies_library_layout_molecular * (sapply(cluster_list_library_layout_molecular, length) / length(library_layout_labels_all)))


# Save metrics to a CSV file (unchanged)
write.csv(metrics_df, "cluster_metrics_3.csv", row.names = FALSE)

# Plot 1: Adjusted Rand Index for Molecular.Type (unchanged)
pdf("ari_molecular_plot_2.pdf", width = 10, height = 6)
ggplot(metrics_df, aes(x = factor(CSV_File, levels = csv_files), y = ARI_Molecular)) +
  geom_point(size = 3) +
  theme_minimal() +
  theme(axis.text.x = element_text(angle = 45, hjust = 1)) +
  labs(
    title = "Adjusted Rand Index for Molecular.Type Across Correlation Matrices",
    x = "Correlation Matrix File",
    y = "Adjusted Rand Index"
  )
dev.off()

# Plot 2: Weighted Entropy for Dataset with Global Entropy Line
pdf("entropy_dataset_with_molecular_plot_3.pdf", width = 10, height = 6)
ggplot(metrics_df, aes(x = factor(CSV_File, levels = csv_files), y = Weighted_Entropy_Dataset)) +
  geom_point(size = 3, color = "black") +
  geom_line(aes(group = 1), color = "black") +
  geom_hline(yintercept = global_entropy, linetype = "dashed", color = "red", size = 1) +
  geom_hline(yintercept = weighted_entropy_dataset_by_molecular, linetype = "dashed", color = "blue", size = 1) +
  theme_minimal() +
  theme(axis.text.x = element_text(angle = 45, hjust = 1)) +
  labs(
    title = "Weighted Entropy for Dataset Across Correlation Matrices",
    x = "Correlation Matrix File",
    y = "Weighted Entropy"
  ) +
  annotate("text", x = 1, y = global_entropy + 0.05, label = sprintf("Global Entropy (Dataset): %.3f", global_entropy),
           hjust = 0, vjust = -0.5, color = "red") +
  annotate("text", x = 1, y = weighted_entropy_dataset_by_molecular + 0.05,
           label = sprintf("Weighted Entropy (Dataset by Molecular.Type): %.3f", weighted_entropy_dataset_by_molecular),
           hjust = 0, vjust = -0.5, color = "blue")
dev.off()


# Plot 3: Weighted Entropy for Library.Type
pdf("entropy_library_type_plot_3.pdf", width = 10, height = 6)
ggplot(metrics_df, aes(x = factor(CSV_File, levels = csv_files), y = Weighted_Entropy_Library_Type)) +
  geom_point(size = 3, color = "black") +
  geom_line(aes(group = 1), color = "black") +
  geom_hline(aes(yintercept = global_entropy_library_type, linetype = "Global (Library.Type)"), color = "red", size = 1) +
  geom_hline(aes(yintercept = weighted_entropy_library_type_by_molecular, linetype = "By Molecular.Type (Library.Type)"), color = "blue", size = 1) +
  scale_linetype_manual(values = c("Global (Library.Type)" = "dashed", "By Molecular.Type (Library.Type)" = "dotted")) +
  theme_minimal() +
  theme(axis.text.x = element_text(angle = 45, hjust = 1), legend.position = "top") +
  labs(
    title = "Weighted Entropy for Library.Type Across Correlation Matrices",
    x = "Correlation Matrix File",
    y = "Weighted Entropy",
    linetype = "Entropy Type"
  ) +
  annotate("text", x = 1, y = global_entropy_library_type + 0.05,
           label = sprintf("Global Entropy (Library.Type): %.3f", global_entropy_library_type),
           hjust = 0, vjust = -0.5, color = "red") +
  annotate("text", x = 1, y = weighted_entropy_library_type_by_molecular + 0.05,
           label = sprintf("Weighted Entropy (Library.Type by Molecular.Type): %.3f", weighted_entropy_library_type_by_molecular),
           hjust = 0, vjust = -0.5, color = "blue")
dev.off()

# Plot 4: Weighted Entropy for Library.Layout
pdf("entropy_library_layout_plot_3.pdf", width = 10, height = 6)
ggplot(metrics_df, aes(x = factor(CSV_File, levels = csv_files), y = Weighted_Entropy_Library_Layout)) +
  geom_point(size = 3, color = "black") +
  geom_line(aes(group = 1), color = "black") +
  geom_hline(aes(yintercept = global_entropy_library_layout, linetype = "Global (Library.Layout)"), color = "red", size = 1) +
  geom_hline(aes(yintercept = weighted_entropy_library_layout_by_molecular, linetype = "By Molecular.Type (Library.Layout)"), color = "blue", size = 1) +
  scale_linetype_manual(values = c("Global (Library.Layout)" = "dashed", "By Molecular.Type (Library.Layout)" = "dotted")) +
  theme_minimal() +
  theme(axis.text.x = element_text(angle = 45, hjust = 1), legend.position = "top") +
  labs(
    title = "Weighted Entropy for Library.Layout Across Correlation Matrices",
    x = "Correlation Matrix File",
    y = "Weighted Entropy",
    linetype = "Entropy Type"
  ) +
  annotate("text", x = 1, y = global_entropy_library_layout + 0.05,
           label = sprintf("Global Entropy (Library.Layout): %.3f", global_entropy_library_layout),
           hjust = 0, vjust = -0.5, color = "red") +
  annotate("text", x = 1, y = weighted_entropy_library_layout_by_molecular + 0.05,
           label = sprintf("Weighted Entropy (Library.Layout by Molecular.Type): %.3f", weighted_entropy_library_layout_by_molecular),
           hjust = 0, vjust = -0.5, color = "blue")
dev.off()
```
