## Supplementary material for "An integrated, transcript-resolution atlas of prostate cancer progression unmasks associations with oncogenic pathways": Reproducible code: pathway_and_isoform_visualization.R

```
# Public submission copy.
# Analytical statements and parameter values are unchanged from the archived source.
# Legacy non-English comments and console messages were normalized for public release.
library(fst)
library(pheatmap)
library(msigdbr)


txi_df <- read.fst("./txi.counts.fst")
txi_info <- read.csv("./transcripts_info.csv")
rownames(txi_df) <- txi_info$transcript_id
txi_counts <- as.matrix(txi_df)

ncount.tx <- log2(sweep(x = txi_counts, MARGIN = 2, STATS = colSums(txi_counts)/1000000, FUN = "/") + 1)

base_tx_ids <- gsub("\\..*$", "", rownames(ncount.tx))
rownames(ncount.tx) <- base_tx_ids

gc_df <- read.fst("./gc_txi.counts.fst")
gc_info <- read.csv("./genes_info.csv")
rownames(gc_df) <- gc_info$gene_id
gc_counts <- as.matrix(gc_df)

ncount.gc <- log2(sweep(x = gc_counts, MARGIN = 2, STATS = colSums(gc_counts)/1000000, FUN = "/") + 1)

base_gc_ids <- gsub("\\..*$", "", rownames(ncount.gc))
rownames(ncount.gc) <- base_gc_ids

gc_info$base_gene_id <- gsub("\\..*$", "", gc_info$gene_id)


#
# message("Running analysis step")
#
# all_msigdb_df <- msigdbr(species = "Homo sapiens")
#
# all_signatures_list <- split(x = all_msigdb_df$gene_symbol, f = all_msigdb_df$gs_name)
# all_signatures_list <- lapply(all_signatures_list, unique)
#
# message(sprintf("Running pathway analysis: %d", length(all_signatures_list)))
#
# message("Running analysis step")
#
#
# message("Running analysis step")
#
# keep_exprs <- rowSums(gc_counts > 3) >= max((ncol(gc_counts) * 0.05), 2)
#
# ensg_to_symbol <- setNames(gc_info$gene_name, gc_info$base_gene_id)
# symbols <- ensg_to_symbol[rownames(ncount.gc)]
#
# valid_rows <- !is.na(symbols) & (symbols != "") & keep_exprs
#
# ncount_sym <- ncount.gc[valid_rows, ]
# symbols_valid <- symbols[valid_rows]
#
# dup_idx <- duplicated(symbols_valid)
# ncount_sym <- ncount_sym[!dup_idx, ]
# rownames(ncount_sym) <- symbols_valid[!dup_idx]
#
# message(sprintf("Analysis completed and output saved: %d", nrow(ncount_sym)))
#
#
#
#
#
#
# ssgsea_all_par <- ssgseaParam(
#   exprData = ncount_sym,
#   geneSets = all_signatures_list,
#   normalize = FALSE
# )
#
#
#
# library(parallel)
#
# total_cores <- detectCores()
# message(sprintf("Analysis progress: %d", total_cores))
#
# use_cores <- max(1, total_cores - 2)
# message(sprintf("Analysis progress: %d", use_cores))
#
# library(BiocParallel)
# bp_param <- MulticoreParam(workers = use_cores, progressbar = TRUE)
#
#
# full_gsva_res <- gsva(ssgsea_all_par, verbose = TRUE,BPPARAM = bp_param)
#
# message("Analysis completed and output saved")
#
# output_file <- paste0("Full_GSVA_Analysis_Workspace_", Sys.Date(), ".Rdata")
#
# message(paste("Running analysis step", output_file))
#
# save(
#   file = output_file
# )
#
# message("Analysis completed and output saved")
#


load("Full_GSVA_Analysis_Workspace_2026-03-23.Rdata")

library(reshape2)

analysis_tasks <- list(
  c("WNT2B",  "ENST00000369684", "ENST00000369686"),
  c("TRIM11", "ENST00000284551", "ENST00000493030"),
  c("ROCK2",  "ENST00000315872", "ENST00000401753"),
  c("RNF43",  "ENST00000584437", "ENST00000583753"),
  c("CDK12",  "ENST00000447079", "ENST00000584632"),
  c("F8",     "ENST00000360256", "ENST00000330287"),
  c("RELA",   "ENST00000406246", "ENST00000612991"),
  c("BRCA1",  "ENST00000357654", "ENST00000352993")
)

all_antagonistic_results <- list()
all_full_results <- list()

n_samples <- ncol(full_gsva_res)

for (task in analysis_tasks) {
  gene_name <- task[1]
  iso1_id   <- task[2]
  iso2_id   <- task[3]

  message(sprintf("Running analysis step: %s %s %s", gene_name, iso1_id, iso2_id))

  if (!(iso1_id %in% rownames(ncount.tx)) || !(iso2_id %in% rownames(ncount.tx))) {
    warning(sprintf("Analysis warning; step skipped: %s", gene_name))
    next
  }

  target_features_exp <- ncount.tx[c(iso1_id, iso2_id), ]

  cor_mat_gsva <- cor(t(target_features_exp), t(full_gsva_res), method = "pearson")

  # t = r * sqrt((n-2)/(1-r^2))
  r_safe <- ifelse(abs(cor_mat_gsva) == 1, 0.999999, cor_mat_gsva)
  t_stat <- r_safe * sqrt((n_samples - 2) / (1 - r_safe^2))
  p_mat_gsva <- 2 * pt(-abs(t_stat), df = n_samples - 2)

  df_res <- data.frame(
    Target_Gene = gene_name,
    Pathway = colnames(cor_mat_gsva),
    Iso1_ID = iso1_id,
    Iso2_ID = iso2_id,
    Cor_Iso1 = cor_mat_gsva[1, ],
    P_Iso1 = p_mat_gsva[1, ],
    Cor_Iso2 = cor_mat_gsva[2, ],
    P_Iso2 = p_mat_gsva[2, ]
  )

  all_full_results[[gene_name]] <- df_res

  df_filtered <- subset(df_res,
                        P_Iso1 < 0.05 &
                          P_Iso2 < 0.05 &
                          (Cor_Iso1 * Cor_Iso2 < 0))

  if (nrow(df_filtered) > 0) {
    df_filtered$Antagonistic_Strength <- abs(df_filtered$Cor_Iso1 - df_filtered$Cor_Iso2)
    df_filtered <- df_filtered[order(-df_filtered$Antagonistic_Strength), ]

    all_antagonistic_results[[gene_name]] <- df_filtered
    message(sprintf("Significant pathways detected: %d", nrow(df_filtered)))
  } else {
    message("Required input was not found")
  }
}


if (length(all_antagonistic_results) > 0) {
  final_antagonistic_df <- do.call(rbind, all_antagonistic_results)
  rownames(final_antagonistic_df) <- NULL

  write.csv(final_antagonistic_df,
            file = "Antagonistic_Pathways_By_Isoforms.csv",
            row.names = FALSE, quote = FALSE)

  message("Running pathway analysis")
} else {
  message("Required input was not found")
}

 final_full_df <- do.call(rbind, all_full_results)
 write.csv(final_full_df, file = "All_Pathways_Correlation_Detailed.csv", row.names = FALSE)


 biology_keywords <- list(
   WNT2B  = "WNT|BETA_CATENIN|STEM_CELL|FZD|DEVELOPMENT|GASTRULATION|CELL_CYCLE",
   TRIM11 = "PROTEASOME|UBIQUITIN|PI3K|AKT|ONCOGENESIS|APOPTOSIS|CELL_CYCLE|NFKB",
   ROCK2  = "ACTIN|CYTOSKELETON|FOCAL_ADHESION|MIGRATION|RHO|ANGIOGENESIS|TGF_BETA",
   RNF43  = "WNT|BETA_CATENIN|UBIQUITIN|PROTEASOME|RECEPTOR_INTERNALIZATION|APC|AXIN|STEM",
   CDK12  = "TRANSCRIPTION|ELONGATION|RNA_POLYMERASE|DNA_REPAIR|SPLICING|HOMOLOGOUS_RECOMBINATION",
   F8     = "COAGULATION|HEMOSTASIS|PLATELET|BLOOD_COAGULATION|WOUND_HEALING",
   RELA   = "NFKB|TNF|INFLAMMATORY|IL6|JAK_STAT|APOPTOSIS|INTERFERON|IMMUNE",
   BRCA1  = "DNA_REPAIR|HOMOLOGOUS_RECOMBINATION|CHECKPOINT|P53|DNA_DAMAGE|CELL_CYCLE|BRCA"
 )

 message("Running analysis step")


 is_high_quality <- grepl("^HALLMARK_|^KEGG_|^REACTOME_|^GOBP_", final_antagonistic_df$Pathway)
 df_hq <- final_antagonistic_df[is_high_quality, ]

 p_threshold <- 0.05
 cor_threshold <- 0.1
 strength_threshold <- 0.5

 df_strict <- subset(df_hq,
                     P_Iso1 < p_threshold &
                       P_Iso2 < p_threshold &
                       abs(Cor_Iso1) > cor_threshold &
                       abs(Cor_Iso2) > cor_threshold &
                       Antagonistic_Strength > strength_threshold)
 table(df_strict$Target_Gene)
 message(sprintf("Analysis completed and output saved: %d %d",
                 nrow(final_antagonistic_df), nrow(df_strict)))

 filtered_gene_list <- list()

 for (gene in names(biology_keywords)) {

   gene_df <- subset(df_strict, Target_Gene == gene)

   if (nrow(gene_df) == 0) {
     message(sprintf("Running pathway analysis: %s", gene))
     filtered_gene_list[[gene]] <- NULL
     next
   }

   keywords_regex <- biology_keywords[[gene]]

   is_biologically_relevant <- grepl(keywords_regex, gene_df$Pathway, ignore.case = TRUE)
   gene_biology_df <- gene_df[is_biologically_relevant, ]

   if (nrow(gene_biology_df) > 0) {
     gene_biology_df <- gene_biology_df[order(-gene_biology_df$Antagonistic_Strength), ]

     filtered_gene_list[[gene]] <- gene_biology_df

     message(sprintf("Running pathway analysis: %s %d %s",
                     gene, nrow(gene_biology_df), gene_biology_df$Pathway[1]))
   } else {
     message(sprintf("Significant pathways detected: %s", gene))
     # filtered_gene_list[[gene]] <- gene_df[order(-gene_df$Antagonistic_Strength), ][1:min(5, nrow(gene_df)), ]
   }
 }

 saveRDS(filtered_gene_list, file = "Expert_Filtered_Antagonistic_Pathways_List.rds")
 message("Analysis completed and output saved")

  print(head(filtered_gene_list[["F8"]]))


  library(ggplot2)
  library(ggpubr)

  analysis_tasks <- list(
    c("WNT2B",  "ENST00000369684", "ENST00000369686"),
    c("TRIM11", "ENST00000284551", "ENST00000493030"),
    c("ROCK2",  "ENST00000315872", "ENST00000401753"),
    c("RNF43",  "ENST00000584437", "ENST00000583753"),
    c("CDK12",  "ENST00000447079", "ENST00000584632"),
    c("F8",     "ENST00000360256", "ENST00000330287"),
    c("RELA",   "ENST00000406246", "ENST00000612991"),
    c("BRCA1",  "ENST00000357654", "ENST00000352993")
  )


  plot_isoform_groups <- function(target_gene, target_pathway,
                                  tx_expr_mat = ncount.tx,
                                  gsva_mat = full_gsva_res,
                                  task_list = analysis_tasks) {

    task_info <- NULL
    for(t in task_list) {
      if(t[1] == target_gene) {
        task_info <- t
        break
      }
    }

    if(is.null(task_info)) {
      stop(sprintf("Analysis progress: %s", target_gene))
    }

    iso1_id <- task_info[2]
    iso2_id <- task_info[3]

    if(!(iso1_id %in% rownames(tx_expr_mat)) || !(iso2_id %in% rownames(tx_expr_mat))) {
      stop("Analysis progress")
    }
    if(!(target_pathway %in% rownames(gsva_mat))) {
      stop("Running pathway analysis")
    }

    iso1_expr <- tx_expr_mat[iso1_id, ]
    iso2_expr <- tx_expr_mat[iso2_id, ]

    iso1_group <- ifelse(iso1_expr > median(iso1_expr), "H", "L")
    iso2_group <- ifelse(iso2_expr > median(iso2_expr), "H", "L")

    combined_group <- paste0(iso1_id, "(", iso1_group, ")_", iso2_id, "(", iso2_group, ")")

    pathway_scores <- gsva_mat[target_pathway, ]

    df_plot <- data.frame(
      Sample = colnames(tx_expr_mat),
      Score = pathway_scores,
      Group = combined_group
    )

    factor_levels <- c(
      paste0(iso1_id, "(H)_", iso2_id, "(H)"),
      paste0(iso1_id, "(H)_", iso2_id, "(L)"),
      paste0(iso1_id, "(L)_", iso2_id, "(H)"),
      paste0(iso1_id, "(L)_", iso2_id, "(L)")
    )
    df_plot$Group <- factor(df_plot$Group, levels = factor_levels)

    my_comparisons <- combn(factor_levels, 2, simplify = FALSE)

    p <- ggplot(df_plot, aes(x = Group, y = Score, fill = Group)) +
      geom_boxplot(outlier.shape = NA, alpha = 0.7, width = 0.6) +
      geom_jitter(width = 0.15, size = 1.2, alpha = 0.5, color = "grey30") +

      stat_compare_means(comparisons = my_comparisons,
                         label = "p.signif",
                         method = "t.test",
                         step.increase = 0.1) +

      stat_compare_means(label.y = max(df_plot$Score) + 0.5,
                         method = "anova", size = 4, color = "darkred") +

      theme_bw(base_size = 13) +
      labs(title = target_pathway,
           subtitle = paste("Antagonistic effect of", target_gene, "isoforms"),
           x = paste(target_gene, "Isoform Combinations"),
           y = "GSVA Enrichment Score") +
      theme(
        axis.text.x = element_text(angle = 25, hjust = 1, color = "black", size = 10),
        axis.text.y = element_text(color = "black"),
        plot.title = element_text(face = "bold", size = 12),
        panel.grid.minor = element_blank()
      ) +
      scale_fill_brewer(palette = "Set2") +
      guides(fill = "none")

    output_filename <- paste0("Boxplot_", target_gene, "_", target_pathway, ".pdf")
    output_filename <- gsub("[ /:]", "_", output_filename)

    ggsave(output_filename, plot = p, width = 8, height = 7)
    message(sprintf("Analysis completed and output saved: %s", output_filename))

    return(p)
  }


  plot_rnf43 <- plot_isoform_groups(
    target_gene = "RNF43",
    target_pathway = "HALLMARK_WNT_BETA_CATENIN_SIGNALING"
  )

  plot_wnt2b <- plot_isoform_groups(
    target_gene = "WNT2B",
    target_pathway = "HALLMARK_WNT_BETA_CATENIN_SIGNALING"
  )

  plot_isoform_groups(
    target_gene = "BRCA1",
    target_pathway = "HALLMARK_DNA_REPAIR"
  )


  library(ggplot2)
  library(reshape2)


  plot_multi_gene_expr <- function(gene_list, expr_mat = ncount_sym, output_name = "MultiGene_Expression_Boxplot") {

    message(sprintf("Running analysis step: %d", length(gene_list)))

    valid_genes <- intersect(gene_list, rownames(expr_mat))
    missing_genes <- setdiff(gene_list, rownames(expr_mat))

    if(length(valid_genes) == 0) {
      stop("Analysis failed")
    }

    if(length(missing_genes) > 0) {
      warning(sprintf("Required input was not found: %s",
                      paste(missing_genes, collapse = ", ")))
    }

    message(sprintf("Analysis progress: %d", length(valid_genes)))

    target_expr <- expr_mat[valid_genes, , drop = FALSE]

    df_long <- melt(target_expr, varnames = c("Gene", "Sample"), value.name = "Expression")

    df_long$Gene <- factor(df_long$Gene, levels = valid_genes)

    p <- ggplot(df_long, aes(x = Gene, y = Expression, fill = Gene)) +
      geom_boxplot(outlier.shape = NA, alpha = 0.7, width = 0.6) +
      geom_jitter(width = 0.2, size = 1, alpha = 0.1, color = "grey30") +
      theme_bw(base_size = 14) +
      labs(title = "Expression Levels of Target Genes",
           x = "Gene Symbol",
           y = "Expression (log2 CPM + 1)") +
      theme(
        axis.text.x = element_text(angle = 45, hjust = 1, color = "black", face = "bold"),
        axis.text.y = element_text(color = "black"),
        plot.title = element_text(face = "bold", hjust = 0.5),
        panel.grid.minor = element_blank(),
        legend.position = "none"
      ) +
      scale_fill_viridis_d(option = "turbo", alpha = 0.7)

    file_name <- paste0(output_name, ".pdf")
    plot_width <- max(6, length(valid_genes) * 0.8)
    ggsave(file_name, plot = p, width = plot_width, height = 6)

    message(sprintf("Analysis completed and output saved: %s", file_name))

    return(p)
  }


  my_genes <- c("WNT2B", "TRIM11", "ROCK2", "RNF43", "CDK12", "F8", "RELA", "BRCA1", "LGR4")

  p_expr <- plot_multi_gene_expr(
    gene_list = my_genes,
    expr_mat = ncount_sym,
    output_name = "Target_8_Genes_Expression"
  )

  print(p_expr)


  my_genes <- c("RSPO1", "RSPO2", "RSPO3", "RSPO4", "LGR4", "LGR5", "LGR6")

  p_expr <- plot_multi_gene_expr(
    gene_list = my_genes,
    expr_mat = ncount_sym,
    output_name = "WNT2B_company_gene"
  )

  print(p_expr)
```
