## Supplementary material for "An integrated, transcript-resolution atlas of prostate cancer progression unmasks associations with oncogenic pathways": Reproducible code: pca_generate_new_style.R

```
# Public submission copy.
# Analytical statements and parameter values are unchanged from the archived source.
# Legacy non-English comments and console messages were normalized for public release.
# Load required libraries
library(readr)
library(dplyr)
library(fst)
library(matrixStats)
library(ggplot2)
library(tidyverse)
library(gridExtra)

# Function to load and preprocess data
load_and_preprocess <- function(counts_file, metadata_file, graph_file, transcript_file, transcript_types = NULL) {
  # Load metadata and filter
  mdata <- read.csv(metadata_file, sep = ",")

  # Note: Keeping the original filtering logic based on layout, but plotting will use type
  id = !is.na(mdata$Library.layout)
  # Filter for PAIRED Library.Layout when using txi.counts.fst
  if (counts_file == "./txi.counts.fst") {
    id = mdata$Library.layout == "PAIRED"
    mdata = mdata[id,]
  }

  # Load counts and normalize
  txi <- list(counts = as.matrix(read.fst(counts_file)))
  ncount.tx <- log2(sweep(x = txi$counts, MARGIN = 2, STATS = colSums(txi$counts)/1000000, FUN = "/") + 1)
  ncount.tx = ncount.tx[,id]
  # Filter by transcript type if specified
  if (!is.null(transcript_types) && counts_file == "./txi.counts.fst") {
    ids <- read.csv(transcript_file, sep = ",", header = TRUE)
    idx <- ids$transcript_type %in% transcript_types
    ncount.tx <- ncount.tx[idx, ]
  }

  return(list(expr_matrix = ncount.tx, mdata = mdata))
}

# Function to perform PCA analysis
perform_pca <- function(expr_matrix, top_n = Inf) {
  # Calculate gene variances
  gene_vars <- rowVars(as.matrix(expr_matrix))

  # Select top variable genes
  if (is.finite(top_n) && top_n < nrow(expr_matrix)) {
    top_genes <- order(gene_vars, decreasing = TRUE)[1:top_n]
    expr_matrix <- expr_matrix[top_genes, ]
  }

  # Standardize data
  expr_scaled <- t(scale(t(expr_matrix)))

  # Perform PCA
  pca_result <- prcomp(t(expr_scaled), scale. = FALSE)

  # Extract scores and variance
  pca_scores <- as.data.frame(pca_result$x[, 1:3])
  colnames(pca_scores) <- c("PC1", "PC2", "PC3")

  var_explained <- pca_result$sdev^2 / sum(pca_result$sdev^2)
  var_percent <- round(var_explained[1:3] * 100, 2)

  return(list(scores = pca_scores, var_percent = var_percent))
}

# Function to generate PCA plots (Only PC1 vs PC2 now)
generate_pca_plots <- function(tmp, var_percent, color_var, output_pdf, color_palette = NULL) {
  pdf(output_pdf, width = 10, height = 8) # 5:4 ratio, matching reference style

  # Define custom theme matching the reference image
  custom_theme <- theme_classic() +
    theme(
      plot.title = element_text(size = 16, face = "bold", hjust = 0), # Left-aligned title
      axis.title = element_text(size = 14),
      axis.text = element_text(size = 12, color = "black"),
      legend.title = element_text(size = 12),
      legend.text = element_text(size = 10),
      axis.line = element_line(colour = "black", linewidth = 0.5) # L-shape border
    )

  # ONLY PC1 vs PC2
  p1 <- ggplot(tmp, aes(x = .data[["PC1"]], y = .data[["PC2"]], fill = .data[[color_var]])) +
    geom_point(shape = 21, color = "black", size = 3, alpha = 0.7, stroke = 0.5) +
    labs(title = paste("PCA: PC1 vs PC2 colored by", color_var),
         x = paste0("PC1 (", var_percent[1], "%)"),
         y = paste0("PC2 (", var_percent[2], "%)")) +
    custom_theme

  # Add manual color scale ONLY if a palette is provided
  if (!is.null(color_palette)) {
    p1 <- p1 + scale_fill_manual(values = color_palette)
  }

  print(p1)
  dev.off()
}

# Main function to run analysis for a given combination
run_analysis <- function(counts_file, transcript_types, top_n, output_prefix) {
  # 1. Create directory for results
  results_dir <- "pca_results_new"
  if (!dir.exists(results_dir)) {
    dir.create(results_dir)
  }

  # Define file paths
  metadata_file <- "filtered_samples.csv"
  graph_file <- "graph.csv"
  transcript_file <- "./../WebToolStudy/transcripts_info.csv"

  # Load and preprocess data
  data <- load_and_preprocess(counts_file, metadata_file, graph_file, transcript_file, transcript_types)

  # Perform PCA
  pca <- perform_pca(data$expr_matrix, top_n)

  # ==========================================
  # NEW: Reverse PC1 direction if using gc_txi file
  # ==========================================
  if (counts_file == "./gc_txi.counts.fst") {
    pca$scores$PC1 <- pca$scores$PC1 * -1
  }

  # Combine metadata with PCA scores
  tmp <- cbind(data$mdata, pca$scores)

  # Define colors for Molecular.Type
  molecular_type_colors <- c(
    "PRIMARY" = "#e20917",
    "ARPC" = "#088ac8",
    "NEPC" = "#8f1c5b",
    "NORMAL" = "#44984f",
    "DNPC" = "#0000ff"
  )

  # Generate output file names
  counts_type <- ifelse(grepl("gc", counts_file), "gc", "txi")
  trans_type <- if (is.null(transcript_types)) "all" else paste(transcript_types, collapse = "_")
  top_n_str <- if (is.infinite(top_n)) "all" else as.character(top_n)

  # File paths
  csv_file <- file.path(results_dir, sprintf("pca_%s_%s_%s.csv", counts_type, trans_type, top_n_str))
  pdf_file_mol <- file.path(results_dir, sprintf("pca_scatter_MolType_%s_%s_%s.pdf", counts_type, trans_type, top_n_str))
  pdf_file_lib <- file.path(results_dir, sprintf("pca_scatter_LibType_%s_%s_%s.pdf", counts_type, trans_type, top_n_str))

  # Generate plot for Molecular.Type
  generate_pca_plots(tmp, pca$var_percent, "Molecular.Type", pdf_file_mol, molecular_type_colors)

  # Generate plot for Library.type
  if ("Library.type" %in% colnames(tmp)) {
    # Passing NULL for colors means ggplot will auto-assign colors for whatever types exist
    generate_pca_plots(tmp, pca$var_percent, "Library.type", pdf_file_lib, NULL)
  }

  # Save CSV
  write.csv(tmp, csv_file, row.names = FALSE)
}

# Define input combinations
counts_files <- c("./txi.counts.fst", "./gc_txi.counts.fst")
transcript_combinations <- list(
  c("lncRNA"),
  c("protein_coding"),
  c("lncRNA", "protein_coding"),
  NULL # For gc_txi.counts.fst, use all transcripts
)

# 2. Only keep the 2000 tier
top_n_values <- c(2000)

# Run analysis for all combinations
for (counts_file in counts_files) {
  trans_options <- if (grepl("gc", counts_file)) transcript_combinations[4] else transcript_combinations[1:3]
  for (trans in trans_options) {
    for (top_n in top_n_values) {
      run_analysis(counts_file, trans, top_n)
    }
  }
}
```
