## Supplementary material for "An integrated, transcript-resolution atlas of prostate cancer progression unmasks associations with oncogenic pathways": Reproducible code: slingshot.R

```
# Public submission copy.
# Analytical statements and parameter values are unchanged from the archived source.
# Legacy non-English comments and console messages were normalized for public release.
library(fst)
  library(slingshot)
  library(RColorBrewer)
  library(ggplot2)
  #
  # txi.gene = list(counts = as.matrix(read.fst("./gc_txi.counts.fst")))
  # pca= readRDS("pca.RDS")
  # mdata = read.csv("filtered_samples.csv")
  #
   keep <- rowSums(ncount > 3) >= (ncol(ncount)*0.05)
   #expr_matrix <- log2(sweep(x=ncount, MARGIN=2, STATS=colSums(ncount)/1000000, FUN="/") + 1)

  # ## start pseudoptime calculation
  #
  sim2d <- SingleCellExperiment(assays = List(counts = as.matrix(ncount[keep,]) ))
  #
  assays(sim2d)$norm <- as.matrix(expr_matrix[keep,])
  reducedDims(sim2d) <- SimpleList(PCA = pca$x[,1:2])
  #
   cl1= kmeans(pca$x[,1:2] , centers = 3)$cluster
   rd1 = pca$x[,1:2]
  #
   plot(rd1, col = brewer.pal(9,"Set1")[cl1], pch=16, asp = 1)
  #
   mdata$clus= cl1
  # pdf(paste0(outdir,"/clusters.pdf"))
  ggplot(mdata) + geom_bar(aes(x=clus, fill=PC.Type))
  ggsave("./version2/clusters.png")
  # dev.off()
   colData(sim2d)$kmeans <- cl1
   sim2d_2 <- slingshot(sim2d, reducedDim = 'PCA',clusterLabels = "kmeans",allow.breaks=F, start.clus=3 , end.clus = 2)
  #
   pdf("./version2/slingshot.pdf")
   colors <- colorRampPalette(brewer.pal(11,'Spectral')[-6])(100)
   plotcol <- colors[cut(sim2d_2$slingPseudotime_1, breaks=100)]
   plot(pca$x[,1:2], col = plotcol, pch=16, asp = 1)
   lines(SlingshotDataSet(sim2d_2), lwd=2,  col='black')
   ggplot(data.frame(x=sim2d_2$slingPseudotime_1, y=mdata$PC.Type)) + geom_point(aes(x=x, y=y))
   ggplot(data.frame(x=sim2d_2$slingPseudotime_1, y=mdata$PC.Type))+geom_histogram(aes(x=x, fill=y))
   dev.off()
   curva = slingCurves(sim2d_2)[[1]]$s
   ordine = slingCurves(sim2d_2)[[1]]$ord
   curve_i <- curva[ordine, ]
   colnames(curve_i) <- c("PCX", "PCY")
   curve_i = as.data.frame(curve_i)
  #
  # write(toJSON(list(curve_i$PCX, curve_i$PCY)), "./objects/slingshot_trajectory_curve.json" )
  #
   saveRDS(curve_i,"./version2/curve_i.RDS")
  #
  # ### To predict pseudotime for new data: predict(sds, pca$x)
   sds = SlingshotDataSet(sim2d_2)
   saveRDS(sim2d_2, "./version2/sim2d.RDS")
   saveRDS(sds,"./version2/slingshot.RDS")
   tmp = slingshot::predict(sds, pca$x[,1:2])
   pseudotimes = slingPseudotime(tmp)[,1]
   mdata$pseudotime = pseudotimes
# #
# # pt = data.frame(pseudotimes,old_pt = mdata$pseudotime)
# # pca = read.csv("./../Sampler_Zurich/pca_all_samples.csv",row.names =1 )
# # pca = read.csv("./../FFPE_integration/denovo_mPCa//pca.csv",row.names =1 )
#
# pseudotime.model = readRDS("./version1/slingshot.RDS")
# pseudotimes = predict(pseudotime.model, pca[,1:2, drop=F] )
# pca$pseudotime = slingPseudotime(pseudotimes)[,"Lineage1"]

# pca.model = readRDS("pca.RDS")
# pseudotimes = predict(pseudotime.model, pca.model$x[,1:2] )
# tmp =data.frame(PT= slingPseudotime(pseudotimes)[,"Lineage1"],
#                 PT_old = mdata$pseudotime)
# cor(tmp$PT,tmp$PT_old)
#
# write.csv(pca,"./../FFPE_integration/denovo_for_atlas/pca_all_samples_with_PT.csv")
#
# pca$totalcounts = colSums(ffpe_counts)
# write.csv(pca,"./../FFPE_integration/denovo_for_atlas/pca_all_samples_with_PT.csv")
```
