## Supplementary material for "An integrated, transcript-resolution atlas of prostate cancer progression unmasks associations with oncogenic pathways": Reproducible code: sva_seq_correction_atlas.R

```
# Public submission copy.
# Analytical statements and parameter values are unchanged from the archived source.
# Legacy non-English comments and console messages were normalized for public release.
library(sva)
library(ggplot2)
library(dplyr)
library(readr)
library(tibble)
library(fst, quietly=T)
library(plotly)

txi = list(counts = as.matrix(read.fst("./../WebToolStudy/gc_txi.counts.fst")))
mdata = read.csv("./../WebToolStudy/filtered_samples.csv")
genes_1 = read.csv("./../WebToolStudy/genes_info.csv")


counts_matrix <- round(txi$counts) %>%
  as.matrix()

metadata_df = mdata

batch_vector <- metadata_df$Dataset
condition_vector <- metadata_df$Molecular.Type


corrected_counts_matrix <- ComBat_seq(
  counts = counts_matrix,
  batch = batch_vector,
  group = condition_vector,
  full_mod = TRUE,
  shrink = FALSE,
  shrink.disp = FALSE
)

save(corrected_counts_matrix, file = "./../WebToolStudy/corrected_counts_matrix.RData")
load("./../WebToolStudy/corrected_counts_matrix.RData")
cpm <- function(counts) {
  sweep(counts, 2, colSums(counts) / 1e6, "/")
}

ff_cpm <- log2(cpm(corrected_counts_matrix) + 1)


gene_vars <- rowVars(as.matrix(ff_cpm))

top_genes <- order(gene_vars, decreasing = TRUE)[1:2000]
top_expr_matrix <- ff_cpm[top_genes, ]
pca = prcomp( t(top_expr_matrix))

to_draw = data.frame(pc1 = pca$x[,1],
                     pc2 = pca$x[,2],
                     pc3 = pca$x[,3],
                     m.type = mdata$Molecular.Type)
save(to_draw, file = "./../WebToolStudy/corrected_counts_matrix_to_draw.RData")


molecular_type_colors <- c(
  "PRIMARY" = "#e20917",
  "ARPC" = "#088ac8",
  "NEPC" = "#8f1c5b",
  "NORMAL" = "#44984f",
  "DNPC" = "#0000ff",
  "FF" = "yellow",
  "FFPE"= "gold")
plot_ly(data = to_draw,
        x = ~pc1,
        y = ~pc2,
        z = ~pc3,
        color = ~m.type,
        colors = molecular_type_colors,
        type = "scatter3d",
        mode = "markers",
        text = ~rownames(to_draw),
        hoverinfo = "text",
        marker = list(size = 5)) %>%
  layout(scene = list(
    xaxis = list(title = "PC1"),
    yaxis = list(title = "PC2"),
    zaxis = list(title = "PC3")
  ),
  title = "3D Scatter Plot of PC1, PC2, PC3 by Molecular Type")
```
