## Supplementary material for "An integrated, transcript-resolution atlas of prostate cancer progression unmasks associations with oncogenic pathways": Reproducible code: heatmap_vis_lnc.R

```
# Public submission copy.
# Analytical statements and parameter values are unchanged from the archived source.
# Legacy non-English comments and console messages were normalized for public release.
library(readr)
library(dplyr)
library(fst, quietly=T)
library(matrixStats)
library(ggplot2)
library(tidyverse)
library(pheatmap)
library(RColorBrewer)

# Load data
mdata = read.csv("filtered_samples.csv")
txi = list(counts = as.matrix(read.fst("./txi.counts.fst")))

# Normalize and log-transform expression matrix
expr_matrix <- log2(sweep(x=txi$counts, MARGIN=2, STATS=colSums(txi$counts)/1000000, FUN="/") + 1)

# Filter for protein-coding transcripts
ids = read.csv("transcripts_info.csv", sep=",", header=T)
idx = ids$transcript_type == "lncRNA"
expr_matrix <- expr_matrix[idx,]

# Calculate correlation matrix
sample_cor <- cor(expr_matrix, method="pearson")

# Prepare annotation data
annotation_df <- mdata %>% select(dataset, Molecular.Type)
annotation_row <- data.frame(
  Dataset = annotation_df$dataset,
  Molecular.Type = annotation_df$Molecular.Type
)
rownames(annotation_row) <- paste0("id.", seq_len(nrow(annotation_row)))

# Define colors for annotations
n_dataset <- length(unique(annotation_row$Dataset))
dataset_colors <- if (n_dataset <= 12) {
  RColorBrewer::brewer.pal(min(n_dataset, 12), "Set3")
} else {
  colorRampPalette(RColorBrewer::brewer.pal(12, "Set3"))(n_dataset)
}

# Use specified colors for Molecular.Type
molecular_type_colors <- c(
  "PRIMARY" = "#e20917",
  "ARPC" = "#088ac8",
  "NEPC" = "#8f1c5b",
  "NORMAL" = "#44984f",
  "DNPC" = "#0000ff"
)

ann_colors <- list(
  Dataset = setNames(dataset_colors, unique(annotation_row$Dataset)),
  Molecular.Type = molecular_type_colors
)

# Plot heatmap
pdf("sample_cor_heatmap_lncRNA.pdf", width=10, height=8)
pheatmap(
  sample_cor,
  annotation_col = annotation_row,
  annotation_colors = ann_colors,
  color = colorRampPalette(rev(brewer.pal(9, "RdBu")))(100),
  clustering_method = "complete",
  show_rownames = FALSE,
  show_colnames = FALSE,
  main = "Sample Correlation Heatmap"
)

# Generate individual heatmaps for each Molecular.Type
molecular_types <- unique(annotation_row$Molecular.Type)
for (mol_type in molecular_types) {
  # Subset data for the current Molecular.Type
  idx <- annotation_row$Molecular.Type == mol_type
  sub_cor <- sample_cor[idx, idx]
  sub_annotation <- annotation_row[idx, , drop=FALSE]

  # Plot heatmap for this Molecular.Type
  pheatmap(
    sub_cor,
    annotation_col = sub_annotation,
    annotation_colors = ann_colors,
    color = colorRampPalette(rev(brewer.pal(9, "RdBu")))(100),
    clustering_method = "complete",
    show_rownames = FALSE,
    show_colnames = FALSE,
    main = paste("Sample Correlation Heatmap -", mol_type)
  )
}
dev.off()

write.csv(sample_cor,"sample_cor_lnc.csv")


cor_matrix_df= read.csv("sample_cor_lnc.csv")
sample_names_cor <- cor_matrix_df[[1]]
cor_matrix_num <- as.matrix(cor_matrix_df[, -1])
rownames(cor_matrix_num) <- sample_names_cor
colnames(cor_matrix_num) <- sample_names_cor
sample_cor = cor_matrix_num
```
