## Supplementary material for "An integrated, transcript-resolution atlas of prostate cancer progression unmasks associations with oncogenic pathways": Reproducible code: kmer_lasso_motif_discovery_FENDRR.R

```
# Public submission copy.
# Analytical statements and parameter values are unchanged from the archived source.
# Legacy non-English comments and console messages were normalized for public release.
# =======================================================
# =======================================================

library(Biostrings)
library(glmnet)
library(dplyr)
library(ggplot2)
library(tibble)

# -------------------------------------------------------
# -------------------------------------------------------
kmer_length <- 5
min_prevalence <- 0.20

lnc <- read.csv("lncRNA_with_pseudotime_corr.csv", stringsAsFactors = FALSE)
lnc <- lnc %>% filter(!is.na(cor_pseudotime))

fasta_file <- "lncRNA_FASTA_output/FENDRR_transcripts.fasta"
seqs <- readDNAStringSet(fasta_file)
names(seqs) <- sapply(strsplit(names(seqs), "\\|"), `[`, 1)

# -------------------------------------------------------
# -------------------------------------------------------
print(paste0("Running analysis step", kmer_length, "Analysis progress"))

kmer_matrix <- oligonucleotideFrequency(seqs, width = kmer_length, as.prob = TRUE)

prevalence <- colMeans(kmer_matrix > 0)

kmer_matrix_filtered <- kmer_matrix[, prevalence >= min_prevalence]

print(paste0("Analysis progress", ncol(kmer_matrix), "Analysis progress",
             ncol(kmer_matrix_filtered), "Analysis progress"))

kmer_df <- as.data.frame(kmer_matrix_filtered)
kmer_df$transcript_id <- names(seqs)

# -------------------------------------------------------
# -------------------------------------------------------
model_data <- inner_join(lnc[, c("transcript_id", "cor_pseudotime")], kmer_df, by = "transcript_id")

if(nrow(model_data) < 10) {
  stop("Too few matched samples to continue")
}

X <- as.matrix(model_data %>% select(-transcript_id, -cor_pseudotime))
Y <- model_data$cor_pseudotime

# -------------------------------------------------------
# -------------------------------------------------------
print("Running analysis step")
set.seed(42)

cv_fit <- cv.glmnet(X, Y, alpha = 1, family = "gaussian")
best_lambda <- cv_fit$lambda.min
coef_matrix <- coef(cv_fit, s = best_lambda)

coef_df <- data.frame(
  K_mer = rownames(coef_matrix),
  Coefficient = as.vector(coef_matrix)
) %>%
  filter(Coefficient != 0) %>%
  filter(K_mer != "(Intercept)") %>%
  arrange(desc(Coefficient))

# -------------------------------------------------------
# -------------------------------------------------------
if(nrow(coef_df) == 0) {
  print("Required input was not found")
} else {
  print("Analysis progress")
  print(coef_df)

  coef_df$K_mer <- factor(coef_df$K_mer, levels = coef_df$K_mer[order(coef_df$Coefficient)])

  p_lasso <- ggplot(coef_df, aes(x = K_mer, y = Coefficient, fill = Coefficient > 0)) +
    geom_bar(stat = "identity", color = "black", alpha = 0.8) +
    coord_flip() +
    scale_fill_manual(values = c("TRUE" = "#E41A1C", "FALSE" = "#377EB8"),
                      name = "Effect Direction") +
    labs(title = paste0("Key ", kmer_length, "-mers Driving Correlation"),
         subtitle = paste0("Filtered for prevalence >= ", min_prevalence * 100, "%"),
         x = "Discovered Motif",
         y = "Coefficient (Impact on Pseudotime Correlation)") +
    theme_bw() +
    theme(axis.text.y = element_text(family = "mono", face = "bold", size = 12))

  print(p_lasso)
  ggsave(paste0("LASSO_Key_Motifs_k", kmer_length, ".png"), p_lasso, width = 8, height = 6)
}


#
#
# # =======================================================
# # =======================================================
#
# library(Biostrings)
# library(dplyr)
# library(ggplot2)
# library(tidyr)
#
# # -------------------------------------------------------
# # -------------------------------------------------------
# target_motifs <- c(
# )
#
# # -------------------------------------------------------
# # -------------------------------------------------------
# fasta_file <- "lncRNA_FASTA_output/FENDRR_transcripts.fasta"
# seqs <- readDNAStringSet(fasta_file)
# names(seqs) <- sapply(strsplit(names(seqs), "\\|"), `[`, 1)
#
# # -------------------------------------------------------
# # -------------------------------------------------------
# print("Running analysis step")
# motif_hits <- list()
#
# for (m_name in names(target_motifs)) {
#   m_seq <- target_motifs[[m_name]]
#
#   matches <- vmatchPattern(m_seq, seqs)
#
#   matches_df <- as.data.frame(matches)
#
#   if(nrow(matches_df) > 0) {
#     matches_df$transcript_id <- names(seqs)[matches_df$group]
#     matches_df$Motif_Type <- m_name
#     matches_df$Sequence <- m_seq
#     motif_hits[[m_name]] <- matches_df
#   }
# }
#
# motif_df <- bind_rows(motif_hits)
#
# # -------------------------------------------------------
# # -------------------------------------------------------
# lnc <- read.csv("lncRNA_with_pseudotime_corr.csv", stringsAsFactors = FALSE)
#
# tx_info <- data.frame(
#   transcript_id = names(seqs),
#   length = width(seqs)
# ) %>%
#   left_join(lnc[, c("transcript_id", "cor_pseudotime")], by = "transcript_id") %>%
#   filter(!is.na(cor_pseudotime)) %>%
#   arrange(desc(cor_pseudotime))
#
# motif_df <- motif_df %>% filter(transcript_id %in% tx_info$transcript_id)
#
# tx_info$transcript_id <- factor(tx_info$transcript_id, levels = tx_info$transcript_id)
# motif_df$transcript_id <- factor(motif_df$transcript_id, levels = levels(tx_info$transcript_id))
#
# # -------------------------------------------------------
# # -------------------------------------------------------
# print("Running analysis step")
#
# p_map <- ggplot() +
#   geom_segment(data = tx_info,
#                aes(x = 1, xend = length, y = transcript_id, yend = transcript_id),
#                color = "gray85", size = 4, lineend = "round") +
#
#   geom_point(data = motif_df,
#              aes(x = start, y = transcript_id, color = Motif_Type, shape = Motif_Type),
#              size = 4, alpha = 0.9) +
#
#   geom_text(data = tx_info,
#             aes(x = -150, y = transcript_id, label = sprintf("r = %.2f", cor_pseudotime)),
#             hjust = 1, size = 3.5, color = "black", fontface = "italic") +
#
#   scale_color_manual(values = c("Positive_Driver" = "#E41A1C", "Negative_Driver" = "#377EB8"),
#                      labels = c("Positive_Driver" = "CAACG (Drives Pos/Less Neg)",
#                                 "Negative_Driver" = "TATGT (Drives Strong Neg)")) +
#   scale_shape_manual(values = c("Positive_Driver" = 17, "Negative_Driver" = 16),
#                      labels = c("Positive_Driver" = "CAACG (Drives Pos/Less Neg)",
#                                 "Negative_Driver" = "TATGT (Drives Strong Neg)")) +
#   labs(title = "Location of Key Regulatory Motifs on FENDRR Transcripts",
#        subtitle = "Transcripts are ordered by Pseudotime Correlation (Top to Bottom: Highest to Lowest r)",
#        x = "Position along Mature RNA Transcript (bp)",
#        y = "Transcript ID",
#        color = "Discovered Motif", shape = "Discovered Motif") +
#   theme_minimal() +
#   theme(panel.grid.minor = element_blank(),
#         axis.text.y = element_text(face = "bold", size = 10),
#         legend.position = "top") +
#   coord_cartesian(clip = "off") +
#   theme(plot.margin = margin(10, 10, 10, 70))
#
# print(p_map)
# ggsave("FENDRR_Motif_Mapping.png", p_map, width = 11, height = 7)
#
# print("Analysis completed and output saved")


# =======================================================
# =======================================================

library(Biostrings)
library(dplyr)
library(ggplot2)
library(rtracklayer)
library(ggtranscript)

# -------------------------------------------------------
# -------------------------------------------------------
target_motifs <- c(
  "Pos_CAACG" = "CAACG",
  "Neg_TATGT" = "TATGT",
  "Neg_ATTCA" = "ATTCA",
  "Neg_TCTCT" = "TCTCT",
  "Neg_TTTAT" = "TTTAT"
)

fasta_file <- "lncRNA_FASTA_output/FENDRR_transcripts.fasta"
seqs <- readDNAStringSet(fasta_file)
names(seqs) <- sapply(strsplit(names(seqs), "\\|"), `[`, 1)

# -------------------------------------------------------
# -------------------------------------------------------
print("Running analysis step")
motif_hits <- list()
for (m_name in names(target_motifs)) {
  m_seq <- target_motifs[[m_name]]
  matches <- as.data.frame(vmatchPattern(m_seq, seqs))

  if(nrow(matches) > 0) {
    matches$transcript_id <- names(seqs)[matches$group]
    matches$Motif_Type <- m_name
    matches$Motif_Seq <- m_seq
    motif_hits[[m_name]] <- matches
  }
}
motif_tx_df <- bind_rows(motif_hits)

# -------------------------------------------------------
# -------------------------------------------------------
print("Loading input data")
gtf_path <- "__PCATLAS_DOCUMENTS_ROOT__/FFPE_integration/gencode.v39.annotation.gtf"
gtf <- rtracklayer::import(gtf_path)
gtf_df <- as.data.frame(gtf)

fendr_gtf <- gtf_df %>%
  filter(gene_name == "FENDRR", type == "exon")

# -------------------------------------------------------
# -------------------------------------------------------
lnc <- read.csv("lncRNA_with_pseudotime_corr.csv", stringsAsFactors = FALSE)
fendr_lnc <- lnc %>%
  filter(gene_name == "FENDRR", !is.na(cor_pseudotime)) %>%
  arrange(desc(cor_pseudotime))

fendr_gtf <- fendr_gtf %>% filter(transcript_id %in% fendr_lnc$transcript_id)

fendr_gtf$transcript_id <- factor(fendr_gtf$transcript_id, levels = fendr_lnc$transcript_id)
fendr_lnc$transcript_id <- factor(fendr_lnc$transcript_id, levels = fendr_lnc$transcript_id)

# -------------------------------------------------------
# -------------------------------------------------------
print("Running analysis step")

exon_mapping <- fendr_gtf %>%
  arrange(transcript_id, as.numeric(exon_number)) %>%
  group_by(transcript_id) %>%
  mutate(
    exon_len = end - start + 1,
    tx_end_pos = cumsum(exon_len),
    tx_start_pos = tx_end_pos - exon_len + 1
  ) %>%
  ungroup()

if(nrow(motif_tx_df) > 0) {
  motif_genomic_df <- motif_tx_df %>%
    inner_join(exon_mapping, by = "transcript_id", relationship = "many-to-many") %>%
    filter(start.x >= tx_start_pos & start.x <= tx_end_pos) %>%
    mutate(
      offset = start.x - tx_start_pos,
      genomic_pos = ifelse(strand == "+",
                           start.y + offset,
                           end.y - offset)
    )
}

# -------------------------------------------------------
# -------------------------------------------------------
print("Running analysis step")

plot_data <- fendr_gtf %>%
  left_join(fendr_lnc[, c("transcript_id", "cor_pseudotime")], by = "transcript_id")

p_map <- ggplot(plot_data, aes(xstart = start, xend = end, y = transcript_id)) +
  geom_range(fill = "white", height = 0.25, color = "gray60") +
  geom_intron(data = to_intron(plot_data, "transcript_id"), color = "gray60") +

  geom_range(aes(fill = cor_pseudotime), color = "black", size = 0.2) +
  scale_fill_gradient2(low = "#377EB8", mid = "white", high = "#E41A1C", midpoint = 0,
                       name = "Correlation\n(r)") +

  geom_point(data = motif_genomic_df,
             aes(x = genomic_pos, y = transcript_id, shape = Motif_Type, color = Motif_Type),
             size = 3.5, stroke = 1.2, inherit.aes = FALSE) +

  scale_color_manual(values = c(
    "Pos_CAACG" = "#FFD700",
    "Neg_TATGT" = "black",
    "Neg_ATTCA" = "#4DAF4A",
    "Neg_TCTCT" = "#984EA3"
  )) +
  scale_shape_manual(values = c(
    "Pos_CAACG" = 17,
    "Neg_TATGT" = 16,
    "Neg_ATTCA" = 15,
    "Neg_TCTCT" = 18
  )) +

  labs(title = "Structural Mapping of Multiple LASSO Motifs on FENDRR",
       subtitle = "Y-axis ordered by Pseudotime Correlation (Top: Less Negative | Bottom: Highly Negative)",
       x = "Genomic Coordinate (Chr16)",
       y = "FENDRR Transcripts",
       shape = "Discovered Motif", color = "Discovered Motif") +
  theme_bw() +
  theme(
    axis.text.y = element_text(face = "bold", size = 9),
    panel.grid.minor = element_blank(),
    legend.position = "bottom",
    legend.box = "horizontal",
    legend.text = element_text(size = 10)
  )

print(p_map)
ggsave("FENDRR_Structure_MultiMotif_Map.png", p_map, width = 13, height = 8)

print("Analysis completed and output saved")
```
