## Supplementary material for "An integrated, transcript-resolution atlas of prostate cancer progression unmasks associations with oncogenic pathways": Reproducible code: kmer_lasso_motif_discovery_FIRRE.R

```
# Public submission copy.
# Analytical statements and parameter values are unchanged from the archived source.
# Legacy non-English comments and console messages were normalized for public release.
# ====================================================================
# ====================================================================

library(Biostrings)
library(glmnet)
library(dplyr)
library(ggplot2)
library(tibble)
library(rtracklayer)
library(ggtranscript)

# =======================================================
# =======================================================
TARGET_GENE <- "FIRRE"
kmer_length <- 10
min_prevalence <- 0.2

print(paste0("Processing target gene", TARGET_GENE, "] <<<"))

# =======================================================
# =======================================================
print("Loading input data")
lnc <- read.csv("lncRNA_with_pseudotime_corr.csv", stringsAsFactors = FALSE)
gene_lnc <- lnc %>% filter(gene_name == TARGET_GENE, !is.na(cor_pseudotime))

fasta_file <- paste0("lncRNA_FASTA_output/", TARGET_GENE, "_transcripts.fasta")
if(!file.exists(fasta_file)) stop("Required input was not found")

seqs <- readDNAStringSet(fasta_file)
names(seqs) <- sapply(strsplit(names(seqs), "\\|"), `[`, 1)

print("Running analysis step")
kmer_matrix <- oligonucleotideFrequency(seqs, width = kmer_length, as.prob = TRUE)
prevalence <- colMeans(kmer_matrix > 0)
kmer_matrix_filtered <- kmer_matrix[, prevalence >= min_prevalence]

kmer_df <- as.data.frame(kmer_matrix_filtered)
kmer_df$transcript_id <- names(seqs)

model_data <- inner_join(gene_lnc[, c("transcript_id", "cor_pseudotime")], kmer_df, by = "transcript_id")

if(nrow(model_data) < 5) stop("Too few matched samples to continue")

X <- as.matrix(model_data %>% select(-transcript_id, -cor_pseudotime))
Y <- model_data$cor_pseudotime

set.seed(42)
cv_fit <- cv.glmnet(X, Y, alpha = 1, family = "gaussian")
coef_matrix <- coef(cv_fit, s = cv_fit$lambda.min)

coef_df <- data.frame(
  K_mer = rownames(coef_matrix),
  Coefficient = as.vector(coef_matrix)
) %>%
  filter(Coefficient != 0, K_mer != "(Intercept)") %>%
  arrange(desc(abs(Coefficient)))

if(nrow(coef_df) == 0) stop("Required input was not found")

coef_plot_df <- coef_df %>% arrange(Coefficient)
coef_plot_df$K_mer <- factor(coef_plot_df$K_mer, levels = coef_plot_df$K_mer)

p_lasso <- ggplot(coef_plot_df, aes(x = K_mer, y = Coefficient, fill = Coefficient > 0)) +
  geom_bar(stat = "identity", color = "black", alpha = 0.8) +
  coord_flip() +
  scale_fill_manual(values = c("TRUE" = "#E41A1C", "FALSE" = "#377EB8"), name = "Effect Direction") +
  labs(title = paste0(TARGET_GENE, ": Key ", kmer_length, "-mers Driving Correlation"),
       x = "Discovered Motif", y = "Coefficient") +
  theme_bw() + theme(axis.text.y = element_text(family = "mono", face = "bold", size = 12))

print(p_lasso)
ggsave(paste0(TARGET_GENE, "_LASSO_Motifs_10.png"), p_lasso, width = 8, height = 6)


 # =======================================================
# =======================================================
print("Running analysis step")

top_n <- min(2, nrow(coef_df))
top_motifs_df <- head(coef_df, top_n) %>%
  mutate(Motif_Name = paste0(ifelse(Coefficient > 0, "Pos_", "Neg_"), K_mer))

target_motifs <- setNames(top_motifs_df$K_mer, top_motifs_df$Motif_Name)

motif_hits <- list()
for (m_name in names(target_motifs)) {
  m_seq <- target_motifs[[m_name]]
  matches <- as.data.frame(vmatchPattern(m_seq, seqs))
  if(nrow(matches) > 0) {
    matches$transcript_id <- names(seqs)[matches$group]
    matches$Motif_Type <- m_name
    motif_hits[[m_name]] <- matches
  }
}
motif_tx_df <- bind_rows(motif_hits)

gtf_path <- "__PCATLAS_DOCUMENTS_ROOT__/FFPE_integration/gencode.v39.annotation.gtf"
gtf <- rtracklayer::import(gtf_path)
  gene_gtf <- as.data.frame(gtf) %>%
  filter(gene_name == TARGET_GENE, type == "exon") %>%
  filter(transcript_id %in% gene_lnc$transcript_id)

exon_mapping <- gene_gtf %>%
  arrange(transcript_id, as.numeric(exon_number)) %>%
  group_by(transcript_id) %>%
  mutate(
    exon_len = end - start + 1,
    tx_end_pos = cumsum(exon_len),
    tx_start_pos = tx_end_pos - exon_len + 1
  ) %>% ungroup()

motif_genomic_df <- data.frame()
if(nrow(motif_tx_df) > 0) {
  motif_genomic_df <- motif_tx_df %>%
    inner_join(exon_mapping, by = "transcript_id", relationship = "many-to-many") %>%
    filter(start.x >= tx_start_pos & start.x <= tx_end_pos) %>%
    mutate(
      offset = start.x - tx_start_pos,
      genomic_pos = ifelse(strand == "+", start.y + offset, end.y - offset)
    )
}

print("Running analysis step")

ordered_tx <- gene_lnc %>%
  arrange((cor_pseudotime)) %>%
  pull(transcript_id) %>%
  unique()

plot_data <- gene_gtf %>%
  left_join(gene_lnc[, c("transcript_id", "cor_pseudotime")], by = "transcript_id") %>%
  mutate(transcript_id = factor(transcript_id, levels = ordered_tx))

if(nrow(motif_genomic_df) > 0) {
  motif_genomic_df <- motif_genomic_df %>%
    mutate(transcript_id = factor(transcript_id, levels = ordered_tx))
}

custom_colors <- c("#FFD700", "black", "#4DAF4A", "#984EA3")
custom_shapes <- c(17, 16, 15, 18)

p_map <- ggplot(plot_data, aes(xstart = start, xend = end, y = transcript_id)) +
  geom_range(fill = "white", height = 0.25, color = "gray60") +
  geom_intron(data = to_intron(plot_data, "transcript_id"), color = "gray60") +
  geom_range(aes(fill = cor_pseudotime), color = "black", size = 0.2) +
  scale_fill_gradient2(low = "#377EB8", mid = "white", high = "#E41A1C", midpoint = 0, name = "Corr(r)")

if(nrow(motif_genomic_df) > 0) {
  p_map <- p_map +
    geom_point(data = motif_genomic_df,
               aes(x = genomic_pos, y = transcript_id, shape = Motif_Type, color = Motif_Type),
               size = 4, stroke = 1.2, inherit.aes = FALSE) +
    scale_color_manual(values = setNames(custom_colors[1:length(target_motifs)], names(target_motifs))) +
    scale_shape_manual(values = setNames(custom_shapes[1:length(target_motifs)], names(target_motifs)))
}

p_map <- p_map +
  labs(title = paste("Structural Mapping of Top LASSO Motifs on", TARGET_GENE),
       subtitle = "Y-axis ordered by Pseudotime Correlation (Top: Less Negative | Bottom: Highly Negative)",
       x = paste("Genomic Coordinate"), y = paste(TARGET_GENE, "Transcripts"),
       shape = "Discovered Motif", color = "Discovered Motif") +
  theme_bw() +
  theme(axis.text.y = element_text(face = "bold", size = 9),
        panel.grid.minor = element_blank(),
        legend.position = "bottom", legend.box = "horizontal")

print(p_map)
ggsave(paste0(TARGET_GENE, "_Structure_MultiMotif_Map_k10_top2.png"), p_map, width = 13, height = 8)

print("Analysis completed and output saved")


library(Biostrings)
library(stringr)

seqs <- readDNAStringSet("lncRNA_FASTA_output/FIRRE_transcripts.fasta")

count_A <- str_count(as.character(seqs), "AAACCAAT")
count_B <- str_count(as.character(seqs), "AAGAGATT")

print(table(count_A))
print(table(count_B))
```
