## Supplementary material for "An integrated, transcript-resolution atlas of prostate cancer progression unmasks associations with oncogenic pathways": Reproducible code: kmer_lasso_pipeline.R

```
# Public submission copy.
# Analytical statements and parameter values are unchanged from the archived source.
# Legacy non-English comments and console messages were normalized for public release.
# ====================================================================
# ====================================================================

library(Biostrings)
library(glmnet)
library(dplyr)
library(ggplot2)
library(tibble)
library(rtracklayer)
library(ggtranscript)
library(stringr)

# -------------------------------------------------------
# -------------------------------------------------------
output_dir <- "Kmer_LASSO_Analysis_Results"
if (!dir.exists(output_dir)) {
  dir.create(output_dir, recursive = TRUE)
  print(paste0("Analysis progress", output_dir))
}

print("Loading input data")
pheno_all <- read.csv("lncRNA_with_pseudotime_corr.csv", stringsAsFactors = FALSE) %>%
  filter(!is.na(cor_pseudotime))

gtf_path <- "__PCATLAS_DOCUMENTS_ROOT__/FFPE_integration/gencode.v39.annotation.gtf"
gtf_all <- as.data.frame(rtracklayer::import(gtf_path))


# =======================================================
# =======================================================
run_motif_pipeline <- function(TARGET_GENE, kmer_length, min_prevalence = 0.2, top_n_to_show = 4) {

  print(paste0("\n========================================================"))
  print(paste0("Processing target gene", TARGET_GENE, "] | kmer: ", kmer_length, " | Top Motifs: ", top_n_to_show))
  print(paste0("========================================================"))

  gene_lnc <- pheno_all %>% filter(gene_name == TARGET_GENE)

  fasta_file <- paste0("lncRNA_FASTA_output/", TARGET_GENE, "_transcripts.fasta")
  if(!file.exists(fasta_file)) stop(paste("Required input was not found", fasta_file))

  seqs <- readDNAStringSet(fasta_file)
  names(seqs) <- sapply(strsplit(names(seqs), "\\|"), `[`, 1)

  print("Running analysis step")
  kmer_matrix <- oligonucleotideFrequency(seqs, width = kmer_length, as.prob = TRUE)
  prevalence <- colMeans(kmer_matrix > 0)
  kmer_matrix_filtered <- kmer_matrix[, prevalence >= min_prevalence, drop = FALSE]

  kmer_df <- as.data.frame(kmer_matrix_filtered)
  kmer_df$transcript_id <- names(seqs)

  model_data <- inner_join(gene_lnc[, c("transcript_id", "cor_pseudotime")], kmer_df, by = "transcript_id")
  if(nrow(model_data) < 5) stop("Too few matched samples to continue")

  X <- as.matrix(model_data %>% select(-transcript_id, -cor_pseudotime))
  Y <- model_data$cor_pseudotime

  set.seed(42)
  n_folds <- min(10, nrow(X))
  cv_fit <- cv.glmnet(X, Y, alpha = 1, family = "gaussian", nfolds = n_folds)
  coef_matrix <- coef(cv_fit, s = cv_fit$lambda.min)

  coef_df <- data.frame(
    K_mer = rownames(coef_matrix),
    Coefficient = as.vector(coef_matrix)
  ) %>%
    filter(Coefficient != 0, K_mer != "(Intercept)") %>%
    arrange(desc(abs(Coefficient)))

  if(nrow(coef_df) == 0) stop("Required input was not found")

  coef_plot_df <- coef_df %>% arrange(Coefficient)
  coef_plot_df$K_mer <- factor(coef_plot_df$K_mer, levels = coef_plot_df$K_mer)

  p_lasso <- ggplot(coef_plot_df, aes(x = K_mer, y = Coefficient, fill = Coefficient > 0)) +
    geom_bar(stat = "identity", color = "black", alpha = 0.8) +
    coord_flip() +
    scale_fill_manual(values = c("TRUE" = "#E41A1C", "FALSE" = "#377EB8"), name = "Effect Direction") +
    labs(title = paste0(TARGET_GENE, ": Key ", kmer_length, "-mers Driving Correlation"), x = "Discovered Motif", y = "Coefficient") +
    theme_bw() + theme(axis.text.y = element_text(family = "mono", face = "bold", size = 12))

  ggsave(file.path(output_dir, paste0(TARGET_GENE, "_LASSO_Motifs_k", kmer_length, ".pdf")), p_lasso, width = 8, height = 6)

  print(paste0("Running analysis step", top_n_to_show, "Analysis progress"))
  actual_top_n <- min(top_n_to_show, nrow(coef_df))
  top_motifs_df <- head(coef_df, actual_top_n) %>%
    mutate(Motif_Name = paste0(ifelse(Coefficient > 0, "Pos_", "Neg_"), K_mer))

  target_motifs <- setNames(top_motifs_df$K_mer, top_motifs_df$Motif_Name)

  motif_hits <- list()
  for (m_name in names(target_motifs)) {
    m_seq <- target_motifs[[m_name]]
    matches <- as.data.frame(vmatchPattern(m_seq, seqs))
    if(nrow(matches) > 0) {
      matches$transcript_id <- names(seqs)[matches$group]
      matches$Motif_Type <- m_name
      motif_hits[[m_name]] <- matches
    }
  }

  motif_tx_df <- bind_rows(motif_hits)
  if(nrow(motif_tx_df) > 0) {
    motif_tx_df <- motif_tx_df %>% dplyr::rename(motif_start = start, motif_end = end)
  }

  gene_gtf <- gtf_all %>%
    filter(gene_name == TARGET_GENE, type == "exon") %>%
    filter(transcript_id %in% gene_lnc$transcript_id)

  exon_mapping <- gene_gtf %>%
    arrange(transcript_id, as.numeric(exon_number)) %>%
    group_by(transcript_id) %>%
    mutate(
      exon_len = end - start + 1,
      tx_end_pos = cumsum(exon_len),
      tx_start_pos = tx_end_pos - exon_len + 1
    ) %>% ungroup()

  motif_genomic_df <- data.frame()
  if(nrow(motif_tx_df) > 0) {
    motif_genomic_df <- motif_tx_df %>%
      inner_join(exon_mapping, by = "transcript_id", relationship = "many-to-many") %>%
      filter(motif_start >= tx_start_pos & motif_start <= tx_end_pos) %>%
      mutate(
        offset = motif_start - tx_start_pos,
        genomic_pos = ifelse(strand == "+", start + offset, end - offset)
      )
  }

  ordered_tx <- gene_lnc %>% arrange(cor_pseudotime) %>% pull(transcript_id) %>% unique()

  plot_data <- gene_gtf %>%
    left_join(gene_lnc[, c("transcript_id", "cor_pseudotime")], by = "transcript_id") %>%
    mutate(transcript_id = factor(transcript_id, levels = ordered_tx))

  if(nrow(motif_genomic_df) > 0) {
    motif_genomic_df <- motif_genomic_df %>% mutate(transcript_id = factor(transcript_id, levels = ordered_tx))
  }

  custom_colors <- c("#FFD700", "black", "#4DAF4A", "#984EA3", "#FF7F00", "#377EB8", "#E41A1C", "#A65628")
  custom_shapes <- c(17, 16, 15, 18, 19, 17, 16, 15)

  p_map <- ggplot(plot_data, aes(xstart = start, xend = end, y = transcript_id)) +
    geom_range(fill = "white", height = 0.25, color = "gray60") +
    geom_intron(data = to_intron(plot_data, "transcript_id"), color = "gray60") +
    geom_range(aes(fill = cor_pseudotime), color = "black", size = 0.2) +
    scale_fill_gradient2(low = "#377EB8", mid = "white", high = "#E41A1C", midpoint = 0, name = "Corr(r)")

  if(nrow(motif_genomic_df) > 0) {
    p_map <- p_map +
      geom_point(data = motif_genomic_df, aes(x = genomic_pos, y = transcript_id, shape = Motif_Type, color = Motif_Type),
                 size = 4, stroke = 1.2, inherit.aes = FALSE) +
      scale_color_manual(values = setNames(custom_colors[1:length(target_motifs)], names(target_motifs))) +
      scale_shape_manual(values = setNames(custom_shapes[1:length(target_motifs)], names(target_motifs)))
  }

  p_map <- p_map +
    labs(title = paste("Structural Mapping of Top", actual_top_n, "LASSO Motifs on", TARGET_GENE),
         subtitle = "Y-axis ordered by Pseudotime Correlation (Top: Less Negative | Bottom: Highly Negative)",
         x = paste("Genomic Coordinate"), y = paste(TARGET_GENE, "Transcripts"), shape = "Discovered Motif", color = "Discovered Motif") +
    theme_bw() +
    theme(axis.text.y = element_text(face = "bold", size = 9), panel.grid.minor = element_blank(), legend.position = "bottom", legend.box = "horizontal")

  ggsave(file.path(output_dir, paste0(TARGET_GENE, "_Structure_MultiMotif_Map_k", kmer_length, "_top", actual_top_n, ".pdf")), p_map, width = 13, height = 8)

  print("Running analysis step")
  for (m_name in names(target_motifs)) {
    m_seq <- target_motifs[[m_name]]
    counts <- str_count(as.character(seqs), m_seq)
    print(paste0(">> ", m_name, " (", m_seq, "Analysis progress"))
    print(table(counts))
  }
  print(paste0("✅ ", TARGET_GENE, "Analysis completed and output saved"))
}

# =======================================================
# =======================================================

run_motif_pipeline(TARGET_GENE = "FIRRE", kmer_length = 10, min_prevalence = 0.2, top_n_to_show = 4)

run_motif_pipeline(TARGET_GENE = "FENDRR", kmer_length = 5, min_prevalence = 0.2, top_n_to_show = 4)

print(paste0("Analysis completed and output saved", output_dir))
```
