## Supplementary material for "An integrated, transcript-resolution atlas of prostate cancer progression unmasks associations with oncogenic pathways": Reproducible code: lncRNA_isoform_plots.R

```
# Public submission copy.
# Analytical statements and parameter values are unchanged from the archived source.
# Legacy non-English comments and console messages were normalized for public release.
library(readr)
library(dplyr)
library(fst)
library(tibble)
library(ggplot2)
library(ggrepel)
library(ggtranscript)
library(rtracklayer)

# =======================================================
# =======================================================
output_dir <- "Isoform_Analysis_Results"
if (!dir.exists(output_dir)) {
  dir.create(output_dir, recursive = TRUE)
  print(paste0("Analysis progress", output_dir))
}

# =======================================================
# =======================================================
print("Loading input data")

counts_file <- "./txi.counts.fst"
meta <- read.csv("filtered_samples.csv", stringsAsFactors = FALSE)
transcripts_info <- read.csv("./../MFE/tx_info_full_2.0.csv", stringsAsFactors = FALSE)

txi <- list(counts = as.matrix(read.fst(counts_file)))
ncount.tx <- log2(sweep(x = txi$counts, MARGIN = 2, STATS = colSums(txi$counts)/1000000, FUN = "/") + 1)
rownames(ncount.tx) = transcripts_info$transcript_id

lnc = read.csv("lncRNA_with_pseudotime_corr.csv", stringsAsFactors = FALSE)

gtf_path <- "__PCATLAS_DOCUMENTS_ROOT__/FFPE_integration/gencode.v39.annotation.gtf"
gtf <- rtracklayer::import(gtf_path)
gtf_df <- as.data.frame(gtf)

# =======================================================
# =======================================================
target_genes <- c("NEAT1", "MALAT1", "FIRRE", "FENDRR")
genes_for_structure <- c("NEAT1", "MALAT1")

print(paste0("Running analysis step"))
print(paste0("Processing target gene", paste(target_genes, collapse = ", ")))

for (target_gene in target_genes) {

  message(paste0("Running analysis step\n", target_gene))

  gene_lnc <- lnc %>% filter(gene_name == target_gene)

  if (nrow(gene_lnc) == 0) {
    message(paste("Analysis progress", target_gene, "Analysis warning; step skipped"))
    next
  }

  gene_ids <- gene_lnc$transcript_id
  valid_ids <- gene_ids[gene_ids %in% rownames(ncount.tx)]

  if (length(valid_ids) == 0) {
    message(paste("Required input was not found", target_gene, "Analysis warning; step skipped"))
    next
  }

  gene_expr <- ncount.tx[valid_ids, , drop=FALSE]

  gene_stats <- data.frame(
    transcript_id = rownames(gene_expr),
    mean_expression = rowMeans(gene_expr)
  )

  gene_analysis <- left_join(gene_lnc, gene_stats, by = "transcript_id")

  p_expr <- ggplot(gene_analysis, aes(x = mean_expression, y = cor_pseudotime)) +
    geom_point(aes(color = padj_pseudotime < 0.05, size = mean_expression)) +
    geom_text_repel(aes(label = transcript_id), size = 3, max.overlaps = 30) +
    geom_hline(yintercept = 0, linetype = "dashed") +
    scale_color_manual(values = c("gray", "red"), name = "Significant") +
    labs(title = paste(target_gene, "Isoforms: Expression vs Correlation"),
         x = "Mean Expression (log2CPM)",
         y = "Correlation with Pseudotime") +
    theme_minimal()

  print(p_expr)

  expr_pdf_path <- file.path(output_dir, paste0(target_gene, "_expr_cor.pdf"))
  ggsave(expr_pdf_path, p_expr, width = 8, height = 6)
  message(paste0("Analysis completed and output saved", expr_pdf_path))

  if (target_gene %in% genes_for_structure) {

    if (exists("gtf_df")) {

      gene_gtf_data <- gtf_df %>%
        filter(gene_name == target_gene, type == "exon") %>%
        filter(transcript_id %in% gene_ids)

      if (nrow(gene_gtf_data) > 0) {

        gene_gtf_plot <- gene_gtf_data %>%
          left_join(gene_lnc[, c("transcript_id", "cor_pseudotime")], by = "transcript_id") %>%
          mutate(transcript_id = reorder(transcript_id, cor_pseudotime))

        p_str <- ggplot(gene_gtf_plot, aes(xstart = start, xend = end, y = transcript_id)) +
          geom_range(aes(y = transcript_id), fill = "white", height = 0.25) +
          geom_intron(data = to_intron(gene_gtf_plot, "transcript_id")) +
          geom_range(aes(fill = cor_pseudotime)) +
          scale_fill_gradient2(low = "blue", mid = "white", high = "red", midpoint = 0) +
          labs(title = paste(target_gene, "Isoform Structures"),
               subtitle = "Sorted by Correlation (Low/Blue -> Bottom, High/Red -> Top)",
               x = "Genomic Position",
               y = "Transcript ID",
               fill = "Correlation") +
          theme_bw() +
          theme(
            axis.text.y = element_text(size = 8),
            panel.grid.major.y = element_line(color = "grey90", linetype = "dotted")
          )

        print(p_str)

        str_pdf_path <- file.path(output_dir, paste0(target_gene, "_str_cor_sorted.pdf"))
        ggsave(str_pdf_path, p_str, width = 10, height = 8)
        message(paste0("Analysis completed and output saved", str_pdf_path))

      } else {
        message(paste("Required input was not found", target_gene, "Analysis progress"))
      }
    } else {
      stop("Analysis failed")
    }
  } else {
    message(paste0("Analysis warning; step skipped"))
  }
}

message(paste0("\n========================================================"))
message(paste0("Analysis completed and output saved"))
message(paste0("Analysis progress", output_dir))
message(paste0("========================================================"))
```
