## Supplementary material for "An integrated, transcript-resolution atlas of prostate cancer progression unmasks associations with oncogenic pathways": Reproducible code: 05_Isoform_Analysis.R

```
#!/usr/bin/Rscript
# Public submission copy.
# Analytical statements and parameter values are unchanged from the archived source.
# Legacy non-English comments and console messages were normalized for public release.
library("ggplot2")
library(ggrepel)
library(tidyverse)
source("./00_functions.R")


mdata=readRDS("./objects/mdata_signatures.RDS")
txi.tx = readRDS("./objects/txi.tx.RDS")
txi.gene = readRDS("./objects/txi.gene.RDS")
ncount = readRDS("./objects/ncount.RDS")
ncount.txi = readRDS("./objects/ncount.txi.RDS")

tx2gene = read.table("./metadata/salmon_tx2gene.tsv")
colnames(tx2gene) = c("transcript_id","gene_id", "gene_name")
tx_info = read.csv("./metadata/transcripts_info.csv")
tx_names = read.table("./metadata/transcript_protein.tsv", col.names = c("transcript_id", "protein_id", "transcript_name", "CDS"), sep="\t")
g_info= unique(tx2gene[,c("gene_id", "gene_name")])
rownames(mdata)=mdata$Name
mdata = mdata[colnames(txi.tx$counts), ]


### For protein coding transcripts, compute the correlation coefficients between transcripts normalized counts
### and pseudotime and the percentage of contribution of each transcript to the gene counts.
### Consider only Paired End samples

onlyPE = mdata$LibraryLayout == "PAIRED"
ptime = mdata$pseudotime[onlyPE]
keep.tx = rowSums(txi.tx$counts > 3) >= (ncol(txi.tx$counts)*0.05) &
  rownames(ncount.txi) %in% tx_info$transcript_id[tx_info$transcript_type == "protein_coding"]

perc_tx = t(sapply(rownames(txi.tx$counts)[keep.tx], function(x){txi.tx$counts[x,mdata$Name[onlyPE]] / txi.gene$counts[tx_info$gene_id[tx_info$transcript_id == x], mdata$Name[onlyPE]] } ))
perc_tx[is.na(perc_tx)] = 0


### consider only the Transcript that at least in one sample contribute to more than 10% to the gene count
keep.tx = keep.tx & rownames(ncount.txi) %in% rownames(perc_tx)[apply(perc_tx, 1, max) > 0.10]
dat = t(sapply(rownames(ncount.txi)[keep.tx],
            function(x){
              ct = cor.test(ptime, ncount.txi[x, mdata$Name[onlyPE]]) ;
              c(tx= x, corr=as.numeric(ct$estimate), pval = as.numeric(ct$p.value), stat = as.numeric(ct$statistic))
              }))
dat = as.data.frame(dat)
dat$corr = as.numeric(dat$corr)
dat$pval = as.numeric(dat$pval)
dat$stat = as.numeric(dat$stat)
if ( sum(is.na(dat$pval)) > 0 ){
  dat[is.na(dat$pval),]$pval = 1
  dat[is.na(dat$stat),]$stat = 0
  dat[is.na(dat$corr),]$corr = 0
}
dat$padj = p.adjust(dat$pval)
dat = merge(dat, data.frame(tx=tx_info$transcript_id, gid= tx_info$gene_id, tx_type = tx_info$transcript_type ), by="tx", all.x=T, all.y=F)
dat = dat %>% group_by(gid) %>% mutate(gCorr = cor(ptime, ncount[unique(gid), mdata$Name[onlyPE]]))

dat = dat[dat$tx_type == "protein_coding", ]
dat = merge(dat, data.frame(gid = g_info$gene_id, name = g_info$gene_name), by="gid", all.x=T, all.y=F)
dat = merge(dat, data.frame(tx = tx_names$transcript_id, protein = tx_names$protein_id, tx_name = tx_names$transcript_name), by="tx")

outdir="./results_05_Isoform_analysis/"

dir.create(outdir, showWarnings = F)
columns_to_write= c("tx", "tx_name", "protein", "gid", "name", "corr", "stat", "padj", "pval", "gCorr", "tx_type")
dat.filtered = dat[abs(dat$corr) > 0.5 & abs(dat$gCorr) < 0.4 ,columns_to_write]


write.csv(dat[order(dat$corr),columns_to_write], paste0(outdir,"Correlations.csv"), row.names = F)
write.csv(dat[abs(dat$corr) > 0.5 & abs(dat$gCorr) < 0.4 ,columns_to_write], paste0(outdir,"Correlations_Filtered.csv"), row.names = F)

nrow(dat.filtered)
sum(dat.filtered$corr < 0)
sum(dat.filtered$corr > 0)
to_plot = dat[abs(dat$gCorr) < 0.4,]
to_plot$color = "grey"
to_plot$color[abs(to_plot$corr) > 0.50 & abs(to_plot$gCorr) < 0.4]="red"
to_plot$color[to_plot$name == "AR"]="blue"
to_plot$label = paste(to_plot$tx, to_plot$tx_name, to_plot$name,sep="\n")
top_n = 5
to_show= c( to_plot$tx[to_plot$name == "AR"], to_plot$tx[to_plot$color == "red" &
                                                           to_plot$name %in% c("ERF","MXI1")],
            to_plot[order(to_plot$corr), "tx"][1:top_n],to_plot[order(to_plot$corr, decreasing = T), "tx"][1:top_n])
pdf(paste0(outdir,"Isoforms_correlations_Filtered.pdf"), width = 10, height = 7)
to_plot$to_show=to_plot$tx_name
to_plot$to_show[! to_plot$tx %in% to_show]=""
ggplot(to_plot, aes(x=corr, y=stat, label=to_show))+
  geom_point(data=to_plot[to_plot$color == "grey", ], color="grey", size=3 ) +
  geom_point(data=to_plot[to_plot$color == "red", ], aes(x=corr, y=stat), fill="red", size=4 , shape=21)+
  geom_point(data=to_plot[to_plot$color == "blue", ], aes(x=corr, y=stat), fill="blue", size=4, shape=21 )+
  geom_text_repel(max.overlaps = Inf, seed=123, size=4,  nudge_y = 1, nudge_x = .5, force = 5) + theme_classic()+
  geom_vline(xintercept = c(-0.5, 0.5), linetype=2)+
  xlab("Pearson Correlation Coefficient") + ylab("Pearson's product moment statistic")
dev.off()


to_show = to_plot$tx[to_plot$name %in% c("ERF")]
to_plot$color = "grey"
to_plot$color[abs(to_plot$corr) > 0.50 & abs(to_plot$gCorr) < 0.4]="red"
to_plot$color[to_plot$tx %in% to_show] = "blue"
to_plot$to_show=to_plot$tx_name
to_plot$to_show[! to_plot$tx %in% to_show]=""

pdf(paste0(outdir,"Isoforms_correlations_ERF.pdf"), width = 10, height = 7)
ggplot(to_plot, aes(x=corr, y=stat, label=to_show))+
  geom_point(data=to_plot[to_plot$color == "grey", ], color="grey", size=3 ) +
  geom_point(data=to_plot[to_plot$color == "red", ], aes(x=corr, y=stat), fill="red", size=4 , shape=21)+
  geom_point(data=to_plot[to_plot$color == "blue", ], aes(x=corr, y=stat), fill="blue", size=4, shape=21 )+
  geom_text_repel(max.overlaps = Inf, seed=123, size=4,  nudge_y = 1, nudge_x = 0.5, force=5) + theme_classic()+
  geom_vline(xintercept = c(-0.5, 0.5), linetype=2)+
  xlab("Pearson Correlation Coefficient") + ylab("Pearson's product moment statistic")
dev.off()


to_show = dat$tx[dat$name %in% c("MXI1")]
to_show = rowSums(ncount.txi[to_show,])
to_show = to_show[order(to_show, decreasing = T)]
to_show = names(to_show)[1:4]
to_plot$color = "grey"
to_plot$color[abs(to_plot$corr) > 0.50 & abs(to_plot$gCorr) < 0.4]="red"
to_plot$color[to_plot$tx %in% to_show] = "blue"
to_plot$to_show=to_plot$tx_name
to_plot$to_show[! to_plot$tx %in% to_show]=""

pdf(paste0(outdir,"Isoforms_correlations_MXI1.pdf"), width = 10, height = 7)
ggplot(to_plot, aes(x=corr, y=stat, label=to_show))+
  geom_point(data=to_plot[to_plot$color == "grey", ], color="grey", size=3 ) +
  geom_point(data=to_plot[to_plot$color == "red", ], aes(x=corr, y=stat), fill="red", size=4 , shape=21)+
  geom_point(data=to_plot[to_plot$color == "blue", ], aes(x=corr, y=stat), fill="blue", size=4 , shape=21)+
  geom_text_repel(max.overlaps = Inf, seed=123, size=4,  nudge_y = 1, nudge_x = 0.5, force=5) + theme_classic()+
  geom_vline(xintercept = c(-0.5, 0.5), linetype=2)+
  xlab("Pearson Correlation Coefficient") + ylab("Pearson's product moment statistic")
dev.off()

### ERF plots and DE analysis
gene_name = "ERF"
gene_id = g_info$gene_id[g_info$gene_name == gene_name]
isoforms = tx_names[tx_names$transcript_id %in% dat$tx[dat$name %in% c(gene_name)],]
target_tx = isoforms$transcript_id[isoforms$transcript_name == "ERF-202"]
dat_comp = data.frame(
  ncount = ncount.txi[target_tx, mdata$Name[onlyPE]],
  sample = mdata$Name[onlyPE],
  subtype = mdata$Subtype[onlyPE]
)

n_global = sum(dat_comp$ncount == 0 )
n_PRIMARY =  sum(dat_comp$ncount[dat_comp$subtype == "PRIMARY"] == 0 )
n_ARPC =  sum(dat_comp$ncount[dat_comp$subtype == "ARPC"] == 0 )

dat_comp$global_comparison = ""
dat_comp$PRIMARY_comparison = ""
dat_comp$ARPC_comparison = ""
dat_comp = dat_comp[order(dat_comp$ncount),]
dat_comp$global_comparison[1:n_global] = "Low"
dat_comp$PRIMARY_comparison[dat_comp$subtype == "PRIMARY"][1:n_PRIMARY] = "Low"
dat_comp$ARPC_comparison[dat_comp$subtype == "ARPC"][1:n_ARPC] = "Low"
dat_comp = dat_comp[order(dat_comp$ncount, decreasing = T),]
dat_comp$global_comparison[1:n_global] = "High"
dat_comp$PRIMARY_comparison[dat_comp$subtype == "PRIMARY"][1:n_PRIMARY] = "High"
dat_comp$ARPC_comparison[dat_comp$subtype == "ARPC"][1:n_ARPC] = "High"
rownames(dat_comp) = dat_comp$sample

to_plot= data.frame(
  id = gene_id,
  name = gene_name,
  ncount = ncount[gene_id, mdata$Name[onlyPE]],
  sample = mdata$Name[onlyPE],
  subtype = mdata$Subtype[onlyPE],
  pseudotime = mdata$pseudotime[onlyPE]
)
for ( i in 1:nrow(isoforms)){
  to_plot = rbind(to_plot, data.frame(
    id = isoforms$transcript_id[i],
    name = isoforms$transcript_name[i],
    ncount = ncount.txi[isoforms$transcript_id[i], mdata$Name[onlyPE]],
    sample = mdata$Name[onlyPE],
    subtype = mdata$Subtype[onlyPE],
    pseudotime = mdata$pseudotime[onlyPE]
  ))
}
to_plot = merge(to_plot, dat_comp[,c("sample", "global_comparison", "PRIMARY_comparison", "ARPC_comparison")], by="sample")
dir.create(paste0(outdir, "/ERF/"), showWarnings = F)
pdf(paste0(outdir, "/ERF/ncount_vs_pseudotime.pdf"), width = 8, height = 9)
ggplot(to_plot,aes(x=pseudotime, y=ncount)) + geom_point(aes(fill=subtype), size=3, shape=21)+ geom_smooth() +
  facet_grid(rows=vars(name))+ theme_bw() +scale_fill_manual(values = palette_subtypes)
ggplot() + geom_point(data=to_plot[to_plot$global_comparison == "",], aes(x=pseudotime, y=ncount), size=3, shape=21, alpha=0.5, fill="grey") +
   geom_point(data=to_plot[to_plot$global_comparison == "Low",], aes(x=pseudotime, y=ncount), size=3, shape=21, fill="blue") +
   geom_point(data=to_plot[to_plot$global_comparison == "High",], aes(x=pseudotime, y=ncount), size=3, shape=21,  fill="red") +
  facet_grid(rows=vars(name))+ theme_bw() + ggtitle("Samples division for the global comparison")
ggplot() + geom_point(data=to_plot[to_plot$PRIMARY_comparison == "",], aes(x=pseudotime, y=ncount), size=3, shape=21, alpha=0.5, fill="grey") +
  geom_point(data=to_plot[to_plot$PRIMARY_comparison == "Low",], aes(x=pseudotime, y=ncount), size=3, shape=21, fill="blue") +
  geom_point(data=to_plot[to_plot$PRIMARY_comparison == "High",], aes(x=pseudotime, y=ncount), size=3, shape=21,  fill="red") +
  facet_grid(rows=vars(name))+ theme_bw()+ ggtitle("Samples division for the PRIMARY samples comparison")
ggplot() + geom_point(data=to_plot[to_plot$ARPC_comparison == "",], aes(x=pseudotime, y=ncount), size=3, shape=21, alpha=0.5, fill="grey") +
  geom_point(data=to_plot[to_plot$ARPC_comparison == "Low",], aes(x=pseudotime, y=ncount), size=3, shape=21, fill="blue") +
  geom_point(data=to_plot[to_plot$ARPC_comparison == "High",], aes(x=pseudotime, y=ncount), size=3, shape=21,  fill="red") +
  facet_grid(rows=vars(name))+ theme_bw()+ ggtitle("Samples division for the ARPC samples comparison")
dev.off()

to_plot = merge(to_plot, mdata[,c("Name", "ESTIMATE.Stromal.Score", "ESTIMATE.Immune.Score")], by.x="sample", by.y="Name")
pdf(paste0(outdir,"/ERF/StromalContribution.pdf"))
ggplot(unique(to_plot[,c("name", "ESTIMATE.Stromal.Score", "global_comparison", "PRIMARY_comparison", "ARPC_comparison")]))+
  geom_boxplot(aes(x=global_comparison, y=ESTIMATE.Stromal.Score))+ theme_classic()
ggplot(unique(to_plot[,c("name", "ESTIMATE.Stromal.Score", "global_comparison", "PRIMARY_comparison", "ARPC_comparison")]))+
  geom_boxplot(aes(x=PRIMARY_comparison, y=ESTIMATE.Stromal.Score))+ theme_classic()
ggplot(unique(to_plot[,c("name", "ESTIMATE.Stromal.Score", "global_comparison", "PRIMARY_comparison", "ARPC_comparison")]))+
  geom_boxplot(aes(x=ARPC_comparison, y=ESTIMATE.Stromal.Score))+ theme_classic()
dev.off()
pdf(paste0(outdir,"/ERF/ImmuneContribution.pdf"))
ggplot(unique(to_plot[,c("name", "ESTIMATE.Immune.Score", "global_comparison", "PRIMARY_comparison", "ARPC_comparison")]))+
  geom_boxplot(aes(x=global_comparison, y=ESTIMATE.Immune.Score))+ theme_classic()
ggplot(unique(to_plot[,c("name", "ESTIMATE.Immune.Score", "global_comparison", "PRIMARY_comparison", "ARPC_comparison")]))+
  geom_boxplot(aes(x=PRIMARY_comparison, y=ESTIMATE.Immune.Score))+ theme_classic()
ggplot(unique(to_plot[,c("name", "ESTIMATE.Immune.Score", "global_comparison", "PRIMARY_comparison", "ARPC_comparison")]))+
  geom_boxplot(aes(x=ARPC_comparison, y=ESTIMATE.Immune.Score))+ theme_classic()
dev.off()


### DE and GSEA
library(clusterProfiler)
gsea.results = list()
de.results = list()
comparisons = data.frame(name = c("GLOBAL", "PRIMARY", "ARPC"), col = c("global_comparison", "PRIMARY_comparison", "ARPC_comparison"))
dir.create(paste0(outdir, "/ERF/Comparisons/"), recursive = T, showWarnings = F)
for ( i in 1:nrow(comparisons) ){
  comp_name = comparisons$name[i]
  comp_col =  comparisons$col[i]
  cmp_mdata = dat_comp[dat_comp[,comp_col] != "",c("sample", comp_col)]
  colnames(cmp_mdata) = c("sample", "condition")
  cmp_mdata$condition = factor(cmp_mdata$condition, levels = c("Low", "High"))
  tmp_txi = txi.gene
  for ( name in names(tmp_txi)[names(tmp_txi) != "countsFromAbundance"]){
    tmp_txi[[name]] = tmp_txi[[name]][,cmp_mdata$sample]
  }

  dds = DESeqDataSetFromTximport(tmp_txi, cmp_mdata, design = ~condition)
  keep <- rowSums(tmp_txi$counts > 10) >= max((ncol(tmp_txi$counts)*0.05), 2) & grepl("\\.[0-9]+$", rownames(tmp_txi$counts))
  sum(keep) / length(keep)
  dds <- DESeq(dds[keep,])
  res = as.data.frame(results(dds, independentFiltering = T))
  res$name = rownames(res)
  res = merge(res, g_info, by.x="name", by.y="gene_id")
  de.results[[comp_name]]=res
  res= res[order(res$log2FoldChange),]
  pdf(paste0(outdir, "/ERF/Comparisons/VolcanoPlot_", comp_name, ".pdf"))
  p<-ggplot(mapping = aes(x=log2FoldChange, y=-log10(padj)))+ geom_point(data= res[res$log2FoldChange < -1 & res$padj < 0.05 ,], color="blue")+
    geom_point(data= res[res$log2FoldChange > 1 & res$padj < 0.05 ,], color="red")+
    geom_point(data= res[abs(res$log2FoldChange) <= 1 | res$padj >= 0.05 ,], color="grey")+ theme_classic() +
    geom_text_repel(data= res[c(1:5, (nrow(res)-5):nrow(res) ), ], aes(label=gene_name))
  print(p)
  dev.off()
  write.csv(res, paste0(outdir, "/ERF/Comparisons/DESeq2_", comp_name, ".csv"), row.names = F)
  stat = res$stat
  names(stat) = gsub( "\\..*$", "", res$name)
  h_gs = msigdbr(species= "Homo sapiens", category="H")
  res.gsea = GSEA(sort(stat, decreasing=T), TERM2GENE = h_gs[,c("gs_name", "ensembl_gene")], minGSSize = 1, maxGSSize = 1000,seed=123, pvalueCutoff=1, verbose = F)
  gsea.results[[comp_name]] = res.gsea@result[,c("ID", "setSize", "enrichmentScore", "NES", "pvalue", "p.adjust", "qvalue", "rank")]
}
saveRDS(list(GSEA= gsea.results, DE = de.results, DAT_COMP = dat_comp ), "./objects/ERF_DE.RDS")
to_plot = NULL
for ( name in names(gsea.results)){
  gsea.results[[name]]$comparison = name
  to_plot = rbind(to_plot, gsea.results[[name]])
}
to_plot = merge(to_plot, h_groups, by.x="ID", by.y="hallmark")
to_plot$HALLMARK = gsub("_", " ", gsub("HALLMARK_", "", to_plot$ID))
to_plot$comparison = factor(to_plot$comparison , levels=c("GLOBAL", "PRIMARY", "ARPC"))
pdf(paste0(outdir, "/ERF/GSEA.pdf"), width = 10, height = 10)
ggplot(to_plot, aes(x = comparison, y= HALLMARK, size= -log10(p.adjust), color = enrichmentScore)) +
  geom_point(data=to_plot[to_plot$p.adjust <=0.05,], alpha=1) +
  geom_point(data=to_plot[to_plot$p.adjust >0.05,], alpha=0.5) +
  scale_color_gradient2(low="blue", high="red")+ facet_wrap(vars(group), scales = "free") + theme_classic() +
  theme( text =element_text(size=8), axis.text.x =element_text(size=7), axis.text.y = element_text(size=5)  )
ggplot(to_plot, aes(x = comparison, y= HALLMARK, size= -log10(p.adjust), color = enrichmentScore)) +
  geom_point(data=to_plot[to_plot$p.adjust <=0.05,], alpha=1) +
  geom_point(data=to_plot[to_plot$p.adjust >0.05,], alpha=0.1) +
  scale_color_gradient2(low="blue", high="red")+ facet_wrap(vars(group), scales = "free") + theme_classic() +
  theme( text =element_text(size=8), axis.text.x =element_text(size=7), axis.text.y = element_text(size=5)  )
ggplot(to_plot, aes(x = comparison, y= HALLMARK, size= -log10(p.adjust), color = enrichmentScore)) +
  geom_point(data=to_plot[to_plot$p.adjust <=0.05,], alpha=1) +
  scale_color_gradient2(low="blue", high="red")+ facet_wrap(vars(group), scales = "free") + theme_classic() +
  theme( text =element_text(size=8), axis.text.x =element_text(size=7), axis.text.y = element_text(size=5)  )
dev.off()

high_samples = mdata[mdata$pseudotime > 150,]
corr.dat = g_info[g_info$gene_id %in% rownames(txi.gene$counts)[rowSums(txi.gene$counts[,high_samples$Name] > 0) > (nrow(high_samples)*0.10) ],]
corr.dat = corr.dat[grepl("\\.[0-9]+$", corr.dat$gene_id),]
corr.dat = corr.dat %>% rowwise() %>% mutate(corr = cor(high_samples$pseudotime, ncount[gene_id,high_samples$Name]), stat = cor.test(high_samples$pseudotime, ncount[gene_id,high_samples$Name])$statistic )
stat = corr.dat$stat
names(stat) = gsub( "\\..*$", "", corr.dat$gene_id)
h_gs = msigdbr(species= "Homo sapiens", category="H")
res.gsea = GSEA(sort(stat, decreasing=T), TERM2GENE = h_gs[,c("gs_name", "ensembl_gene")], minGSSize = 1, maxGSSize = 1000,seed=123, pvalueCutoff=1, verbose = F)
res.gsea = res.gsea@result
res.gsea$comparison = "HighPseudotime"

to_plot = NULL
for ( name in names(gsea.results)){
  gsea.results[[name]]$comparison = name
  to_plot = rbind(to_plot, gsea.results[[name]])
}
colnames(to_plot)
to_plot = rbind(to_plot, res.gsea[,colnames(to_plot)])
to_plot = merge(to_plot, h_groups, by.x="ID", by.y="hallmark")
to_plot$HALLMARK = gsub("_", " ", gsub("HALLMARK_", "", to_plot$ID))
to_plot$comparison = factor(to_plot$comparison , levels=c("GLOBAL", "PRIMARY", "ARPC", "HighPseudotime"))


pdf(paste0(outdir, "/ERF/GSEA_plus.pdf"), width = 10, height = 10)
ggplot(to_plot, aes(x = comparison, y= HALLMARK, size= -log10(p.adjust), color = enrichmentScore)) +
  geom_point(data=to_plot[to_plot$p.adjust <=0.05,], alpha=1) +
  geom_point(data=to_plot[to_plot$p.adjust >0.05,], alpha=0.5) +
  scale_color_gradient2(low="blue", high="red")+ facet_wrap(vars(group), scales = "free") + theme_classic() +
  theme( text =element_text(size=8), axis.text.x =element_text(size=7), axis.text.y = element_text(size=5)  )
ggplot(to_plot, aes(x = comparison, y= HALLMARK, size= -log10(p.adjust), color = enrichmentScore)) +
  geom_point(data=to_plot[to_plot$p.adjust <=0.05,], alpha=1) +
  geom_point(data=to_plot[to_plot$p.adjust >0.05,], alpha=0.1) +
  scale_color_gradient2(low="blue", high="red")+ facet_wrap(vars(group), scales = "free") + theme_classic() +
  theme( text =element_text(size=8), axis.text.x =element_text(size=7), axis.text.y = element_text(size=5)  )
ggplot(to_plot, aes(x = comparison, y= HALLMARK, size= -log10(p.adjust), color = enrichmentScore)) +
  geom_point(data=to_plot[to_plot$p.adjust <=0.05,], alpha=1) +
  scale_color_gradient2(low="blue", high="red")+ facet_wrap(vars(group), scales = "free") + theme_classic() +
  theme( text =element_text(size=8), axis.text.x =element_text(size=7), axis.text.y = element_text(size=5)  )
ggplot(to_plot[to_plot$comparison == "HighPseudotime",], aes(x = comparison, y= HALLMARK, size= -log10(p.adjust), color = enrichmentScore)) +
  geom_point(data=to_plot[to_plot$comparison == "HighPseudotime" & to_plot$p.adjust <=0.05,], alpha=1) +
  geom_point(data=to_plot[to_plot$comparison == "HighPseudotime" & to_plot$p.adjust >0.05,], alpha=0.5) +
  scale_color_gradient2(low="blue", high="red")+ facet_wrap(vars(group), scales = "free") + theme_classic() +
  theme( text =element_text(size=8), axis.text.x =element_text(size=7), axis.text.y = element_text(size=5)  )
dev.off()

####use the bot 100 genes as signature

tmp = de.results$ARPC
tmp = tmp[tmp$padj < 0.05 ,]
tmp = tmp[order(tmp$log2FoldChange),]
tmp = tmp[!grepl("^ENSG", tmp$gene_name),]
ERF_LOW_SIGNATURE = tmp$gene_name[1:100]
ERF_LOW_SIGNATURE = tmp$name[1:100]
to_keep = rownames(txi.gene$counts)[rowSums(txi.gene$counts[,mdata$Name] > 10) > 10]
write.table(ERF_LOW_SIGNATURE, paste0(outdir, "/ERF/ERF_LOW_SIGNATURE.txt"), row.names = F, quote=F, col.names = F)
res = gsva(as.matrix(ncount[to_keep,mdata$Name,drop=F]), list(ERF_LOW_SIGNATURE=ERF_LOW_SIGNATURE), mx.diff=TRUE, method = "ssgsea", parallel.sz=4, ssgsea.norm=F)
res=t(res)
res = as.data.frame(res)
res$Name = rownames(res)
mdata.ssgsea = merge(mdata, res, by="Name")
pdf(paste0(outdir, "/ERF/ARPC_ERF_minor_signature.pdf"), width = 8)
ggplot(mdata.ssgsea) + geom_point(aes(x=PC1, y=PC2, fill=ERF_LOW_SIGNATURE), shape=21, size=3)+scale_fill_gradient2(low = "blue", high = "red")+ theme_classic()
ggplot(mdata.ssgsea) + geom_point(aes(x=pseudotime, y=ERF_LOW_SIGNATURE, fill=Subtype), shape=21, size=3)+scale_fill_manual(values=palette_subtypes) + theme_classic()
ggplot(mdata.ssgsea) + geom_point(aes(x=pseudotime, y=ERF_LOW_SIGNATURE, fill=Subtype), shape=21, size=3)+scale_fill_manual(values=palette_subtypes) + theme_classic() + xlim(c(150,250))
ggplot(mdata.ssgsea[mdata.ssgsea$Subtype %in% c("ARPC", "DNPC", "NEPC"),]) + geom_point(aes(x=pseudotime, y=ERF_LOW_SIGNATURE, fill=Subtype), shape=21, size=3)+
  scale_fill_manual(values=palette_subtypes[3:5]) + theme_classic() + xlim(c(150,250)) +
  geom_smooth(aes(x=pseudotime, y=ERF_LOW_SIGNATURE))+
  facet_grid(rows=vars(Subtype))+
  ggtitle(paste0("ARPC corr=",
                 round(cor(mdata.ssgsea$pseudotime[mdata.ssgsea$Subtype == "ARPC" & mdata.ssgsea$pseudotime > 150], mdata.ssgsea$ERF_LOW_SIGNATURE[mdata.ssgsea$Subtype == "ARPC" & mdata.ssgsea$pseudotime > 150]), 3),
                 " ( p-value ", cor.test(mdata.ssgsea$pseudotime[mdata.ssgsea$Subtype == "ARPC" & mdata.ssgsea$pseudotime > 150], mdata.ssgsea$ERF_LOW_SIGNATURE[mdata.ssgsea$Subtype == "ARPC" & mdata.ssgsea$pseudotime > 150])$p.value, ")",
                 "\nDNPC corr=",
                 round(cor(mdata.ssgsea$pseudotime[mdata.ssgsea$Subtype == "DNPC" & mdata.ssgsea$pseudotime > 150], mdata.ssgsea$ERF_LOW_SIGNATURE[mdata.ssgsea$Subtype == "DNPC" & mdata.ssgsea$pseudotime > 150]), 3),
                 " ( p-value ", cor.test(mdata.ssgsea$pseudotime[mdata.ssgsea$Subtype == "DNPC" & mdata.ssgsea$pseudotime > 150], mdata.ssgsea$ERF_LOW_SIGNATURE[mdata.ssgsea$Subtype == "DNPC" & mdata.ssgsea$pseudotime > 150])$p.value, ")",
                 "\nNEPC corr=",
                 round(cor(mdata.ssgsea$pseudotime[mdata.ssgsea$Subtype == "NEPC" & mdata.ssgsea$pseudotime > 150], mdata.ssgsea$ERF_LOW_SIGNATURE[mdata.ssgsea$Subtype == "NEPC" & mdata.ssgsea$pseudotime > 150]), 3),
                 " ( p-value ", cor.test(mdata.ssgsea$pseudotime[mdata.ssgsea$Subtype == "NEPC" & mdata.ssgsea$pseudotime > 150], mdata.ssgsea$ERF_LOW_SIGNATURE[mdata.ssgsea$Subtype == "NEPC" & mdata.ssgsea$pseudotime > 150])$p.value, ")"
                 ))

dev.off()


### MXI1 plots and DE analysis
gene_name = "MXI1"
gene_id = g_info$gene_id[g_info$gene_name == gene_name]
isoforms = tx_names[tx_names$transcript_id %in% dat$tx[dat$name %in% c(gene_name)],]
isoforms = isoforms[isoforms$transcript_name %in% c("MXI1-202", "MXI1-201", "MXI1-214"),]
target_tx = isoforms$transcript_id[isoforms$transcript_name == "MXI1-214"]
dat_comp = data.frame(
  ncount = ncount.txi[target_tx, mdata$Name[onlyPE]],
  sample = mdata$Name[onlyPE],
  subtype = mdata$Subtype[onlyPE]
)

n_ARPC =  min(sum(dat_comp$ncount[dat_comp$subtype == "ARPC"] == 0 ),sum(dat_comp$ncount[dat_comp$subtype == "ARPC"] > 0 ))
n_DNPC =  min(sum(dat_comp$ncount[dat_comp$subtype == "DNPC"] == 0 ),sum(dat_comp$ncount[dat_comp$subtype == "DNPC"] > 0 ))
n_NEPC =  min(sum(dat_comp$ncount[dat_comp$subtype == "NEPC"] == 0 ),sum(dat_comp$ncount[dat_comp$subtype == "NEPC"] > 0 ))

dat_comp$NEPC_comparison = ""
dat_comp$DNPC_comparison = ""
dat_comp$ARPC_comparison = ""
dat_comp$NEPC_comparison[dat_comp$subtype == "NEPC" & dat_comp$ncount == 0 ] = "Low"
dat_comp$DNPC_comparison[dat_comp$subtype == "DNPC" & dat_comp$ncount == 0 ] = "Low"
dat_comp$ARPC_comparison[dat_comp$subtype == "ARPC" & dat_comp$ncount == 0 ] = "Low"
dat_comp$NEPC_comparison[dat_comp$subtype == "NEPC" & dat_comp$ncount > 0 ] = "High"
dat_comp$DNPC_comparison[dat_comp$subtype == "DNPC" & dat_comp$ncount > 0 ] = "High"
dat_comp$ARPC_comparison[dat_comp$subtype == "ARPC" & dat_comp$ncount > 0 ] = "High"
rownames(dat_comp) = dat_comp$sample

to_plot= data.frame(
  id = gene_id,
  name = gene_name,
  ncount = ncount[gene_id, mdata$Name[onlyPE]],
  sample = mdata$Name[onlyPE],
  subtype = mdata$Subtype[onlyPE],
  pseudotime = mdata$pseudotime[onlyPE]
)
for ( i in 1:nrow(isoforms)){
  to_plot = rbind(to_plot, data.frame(
    id = isoforms$transcript_id[i],
    name = isoforms$transcript_name[i],
    ncount = ncount.txi[isoforms$transcript_id[i], mdata$Name[onlyPE]],
    sample = mdata$Name[onlyPE],
    subtype = mdata$Subtype[onlyPE],
    pseudotime = mdata$pseudotime[onlyPE]
  ))
}
to_plot = merge(to_plot, dat_comp[,c("sample", "NEPC_comparison", "DNPC_comparison", "ARPC_comparison")], by="sample")
dir.create(paste0(outdir, "/MXI1/"), showWarnings = F)
pdf(paste0(outdir, "/MXI1/ncount_vs_pseudotime.pdf"), width = 8, height = 9)
ggplot(to_plot,aes(x=pseudotime, y=ncount)) + geom_point(aes(fill=subtype),size=3, shape=21) + geom_smooth()+
  facet_grid(rows=vars(name))+ theme_bw() +scale_fill_manual(values = palette_subtypes)
ggplot() + geom_point(data=to_plot[to_plot$ARPC_comparison == "",], aes(x=pseudotime, y=ncount), size=3, shape=21, alpha=0.5, fill="grey") +
  geom_point(data=to_plot[to_plot$ARPC_comparison == "Low",], aes(x=pseudotime, y=ncount), size=3, shape=21, fill="blue") +
  geom_point(data=to_plot[to_plot$ARPC_comparison == "High",], aes(x=pseudotime, y=ncount), size=3, shape=21,  fill="red") +
  facet_grid(rows=vars(name))+ theme_bw() + ggtitle("Samples division for the ARPC samples comparison")
ggplot() + geom_point(data=to_plot[to_plot$NEPC_comparison == "",], aes(x=pseudotime, y=ncount), size=3, shape=21, alpha=0.5, fill="grey") +
  geom_point(data=to_plot[to_plot$NEPC_comparison == "Low",], aes(x=pseudotime, y=ncount), size=3, shape=21, fill="blue") +
  geom_point(data=to_plot[to_plot$NEPC_comparison == "High",], aes(x=pseudotime, y=ncount), size=3, shape=21,  fill="red") +
  facet_grid(rows=vars(name))+ theme_bw()+ ggtitle("Samples division for the NEPC samples comparison")
ggplot() + geom_point(data=to_plot[to_plot$DNPC_comparison == "",], aes(x=pseudotime, y=ncount), size=3, shape=21, alpha=0.5, fill="grey") +
  geom_point(data=to_plot[to_plot$DNPC_comparison == "Low",], aes(x=pseudotime, y=ncount), size=3, shape=21, fill="blue") +
  geom_point(data=to_plot[to_plot$DNPC_comparison == "High",], aes(x=pseudotime, y=ncount), size=3, shape=21,  fill="red") +
  facet_grid(rows=vars(name))+ theme_bw()+ ggtitle("Samples division for the DNPC samples comparison")
dev.off()

to_plot = merge(to_plot, mdata[,c("Name", "ESTIMATE.Stromal.Score", "ESTIMATE.Immune.Score")], by.x="sample", by.y="Name")
pdf(paste0(outdir,"/MXI1/StromalContribution.pdf"))
ggplot(unique(to_plot[,c("name", "ESTIMATE.Stromal.Score", "NEPC_comparison", "DNPC_comparison", "ARPC_comparison")]))+
  geom_boxplot(aes(x=ARPC_comparison, y=ESTIMATE.Stromal.Score))+ theme_classic()
ggplot(unique(to_plot[,c("name", "ESTIMATE.Stromal.Score", "NEPC_comparison", "DNPC_comparison", "ARPC_comparison")]))+
  geom_boxplot(aes(x=NEPC_comparison, y=ESTIMATE.Stromal.Score))+ theme_classic()
ggplot(unique(to_plot[,c("name", "ESTIMATE.Stromal.Score", "NEPC_comparison", "DNPC_comparison", "ARPC_comparison")]))+
  geom_boxplot(aes(x=DNPC_comparison, y=ESTIMATE.Stromal.Score))+ theme_classic()
dev.off()
pdf(paste0(outdir,"/MXI1/ImmuneContribution.pdf"))
ggplot(unique(to_plot[,c("name", "ESTIMATE.Immune.Score", "NEPC_comparison", "DNPC_comparison", "ARPC_comparison")]))+
  geom_boxplot(aes(x=ARPC_comparison, y=ESTIMATE.Immune.Score))+ theme_classic()
ggplot(unique(to_plot[,c("name", "ESTIMATE.Immune.Score", "NEPC_comparison", "DNPC_comparison", "ARPC_comparison")]))+
  geom_boxplot(aes(x=NEPC_comparison, y=ESTIMATE.Immune.Score))+ theme_classic()
ggplot(unique(to_plot[,c("name", "ESTIMATE.Immune.Score", "NEPC_comparison", "DNPC_comparison", "ARPC_comparison")]))+
  geom_boxplot(aes(x=DNPC_comparison, y=ESTIMATE.Immune.Score))+ theme_classic()
dev.off()


### DE and GSEA
library(clusterProfiler)
gsea.results = list()
de.results = list()
comparisons = data.frame(name = c("ARPC", "NEPC", "DNPC"), col = c("ARPC_comparison", "NEPC_comparison", "DNPC_comparison"))
dir.create(paste0(outdir, "/MXI1/Comparisons/"), recursive = T, showWarnings = F)
for ( i in 1:nrow(comparisons) ){
  comp_name = comparisons$name[i]
  comp_col =  comparisons$col[i]
  cmp_mdata = dat_comp[dat_comp[,comp_col] != "",c("sample", comp_col)]
  colnames(cmp_mdata) = c("sample", "condition")
  cmp_mdata$condition = factor(cmp_mdata$condition, levels = c("Low", "High"))
  tmp_txi = txi.gene
  for ( name in names(tmp_txi)[names(tmp_txi) != "countsFromAbundance"]){
    tmp_txi[[name]] = tmp_txi[[name]][,cmp_mdata$sample]
  }

  dds = DESeqDataSetFromTximport(tmp_txi, cmp_mdata, design = ~condition)
  keep <- rowSums(tmp_txi$counts > 10) >= max((ncol(tmp_txi$counts)*0.05), 2) & grepl("\\.[0-9]+$", rownames(tmp_txi$counts))
  sum(keep) / length(keep)
  dds <- DESeq(dds[keep,])
  res = as.data.frame(results(dds, independentFiltering = T))
  res$name = rownames(res)
  res = merge(res, g_info, by.x="name", by.y="gene_id")
  de.results[[comp_name]]=res
  res= res[order(res$log2FoldChange),]
  pdf(paste0(outdir, "/MXI1/Comparisons/VolcanoPlot_", comp_name, ".pdf"))
  p<-ggplot(mapping = aes(x=log2FoldChange, y=-log10(padj)))+ geom_point(data= res[res$log2FoldChange < -1 & res$padj < 0.05 ,], color="blue")+
    geom_point(data= res[res$log2FoldChange > 1 & res$padj < 0.05 ,], color="red")+
    geom_point(data= res[abs(res$log2FoldChange) <= 1 | res$padj >= 0.05 ,], color="grey")+ theme_classic() +
    geom_text_repel(data= res[c(1:5, (nrow(res)-5):nrow(res) ), ], aes(label=gene_name))
  print(p)
  dev.off()
  write.csv(res, paste0(outdir, "/MXI1/Comparisons/DESeq2_", comp_name, ".csv"), row.names = F)
  stat = res$stat
  names(stat) = gsub( "\\..*$", "", res$name)
  h_gs = msigdbr(species= "Homo sapiens", category="H")
  res.gsea = GSEA(sort(stat, decreasing=T), TERM2GENE = h_gs[,c("gs_name", "ensembl_gene")], minGSSize = 1, maxGSSize = 1000,seed=123, pvalueCutoff=1, verbose = F)
  gsea.results[[comp_name]] = res.gsea@result[,c("ID", "setSize", "enrichmentScore", "NES", "pvalue", "p.adjust", "qvalue", "rank")]
}
saveRDS(list(GSEA= gsea.results, DE = de.results, DAT_COMP = dat_comp ), "./objects/MXI1_DE.RDS")

to_plot = NULL
for ( name in names(gsea.results)){
  gsea.results[[name]]$comparison = name
  to_plot = rbind(to_plot, gsea.results[[name]])
}
to_plot = merge(to_plot, h_groups, by.x="ID", by.y="hallmark")
to_plot$HALLMARK = gsub("_", " ", gsub("HALLMARK_", "", to_plot$ID))
to_plot$comparison = factor(to_plot$comparison , levels=c("ARPC", "DNPC", "NEPC"))
pdf(paste0(outdir, "/MXI1/GSEA.pdf"), width = 10, height = 10)
ggplot(to_plot, aes(x = comparison, y= HALLMARK, size= -log10(p.adjust), color = enrichmentScore)) +
  geom_point(data=to_plot[to_plot$p.adjust <=0.05,], alpha=1) +
  geom_point(data=to_plot[to_plot$p.adjust >0.05,], alpha=0.5) +
  scale_color_gradient2(low="blue", high="red")+ facet_wrap(vars(group), scales = "free") + theme_classic() +
  theme( text =element_text(size=8), axis.text.x =element_text(size=7), axis.text.y = element_text(size=5)  )
ggplot(to_plot, aes(x = comparison, y= HALLMARK, size= -log10(p.adjust), color = enrichmentScore)) +
  geom_point(data=to_plot[to_plot$p.adjust <=0.05,], alpha=1) +
  geom_point(data=to_plot[to_plot$p.adjust >0.05,], alpha=0.1) +
  scale_color_gradient2(low="blue", high="red")+ facet_wrap(vars(group), scales = "free") + theme_classic() +
  theme( text =element_text(size=8), axis.text.x =element_text(size=7), axis.text.y = element_text(size=5)  )
ggplot(to_plot, aes(x = comparison, y= HALLMARK, size= -log10(p.adjust), color = enrichmentScore)) +
  geom_point(data=to_plot[to_plot$p.adjust <=0.05,], alpha=1) +
  scale_color_gradient2(low="blue", high="red")+ facet_wrap(vars(group), scales = "free") + theme_classic() +
  theme( text =element_text(size=8), axis.text.x =element_text(size=7), axis.text.y = element_text(size=5)  )
dev.off()


###

### AR V-7 analysis
palette_subtypes = c("#25681E", "#F63A21","#4889C7", "#1010C0", "#93355A")

to_plot = NULL
variants = list("AR-V7"="ENST00000504326.5", "AR-FL"="ENST00000374690.9")
for ( variant in names(variants)){
  tmp = mdata[onlyPE,c("Name", "PC.Type", "Subtype", "pseudotime")]
  tmp$isoform = variant
  tmp$ncount = ncount.txi[variants[[variant]], tmp$Name]
  tmp$perc = perc_tx[variants[[variant]], tmp$Name]
  to_plot = rbind(to_plot, tmp)
}
library(ggsignif)
library(gridExtra)

pdf(paste0(outdir, "AR_v7.pdf"))
to_plot$Subtype = factor(to_plot$Subtype, levels= c("NORMAL", "PRIMARY", "ARPC", "DNPC", "NEPC"))
ggplot(to_plot) + geom_boxplot(aes(x=Subtype, fill=isoform, y=ncount)) + theme_classic() + scale_fill_manual(values = c("blue", "red"))+ylab("transcript CPM")+
  facet_grid(rows=vars(isoform))
p1 = ggplot(to_plot[to_plot$isoform == "AR-FL",], aes(x=Subtype, fill=isoform, y=ncount)) + geom_boxplot()  + geom_signif(comparisons = list(c("PRIMARY", "ARPC"), c("NORMAL", "PRIMARY"), c("DNPC", "NEPC")), test = "t.test")+
  theme_classic() + scale_fill_manual(values = c("red"))+ylab("transcript CPM") +
  facet_grid(rows=vars(isoform))
p2 = ggplot(to_plot[to_plot$isoform == "AR-V7",], aes(x=Subtype, fill=isoform, y=ncount)) + geom_boxplot()  + geom_signif(comparisons = list(c("PRIMARY", "ARPC"), c("NORMAL", "PRIMARY"), c("DNPC", "NEPC")), test = "t.test")+
  theme_classic() + scale_fill_manual(values = c("blue"))+ylab("transcript CPM") +
  facet_grid(rows=vars(isoform))
grid.arrange(p1, p2, nrow=2)

ggplot(to_plot) + geom_point(aes(x=pseudotime, y=ncount, fill=Subtype), shape=21, size=3) + theme_classic() +
  scale_fill_manual(values = palette_subtypes) + facet_grid(rows = vars(isoform)) + xlab("Disease Progression") + ylab("transcript CPM")
ggplot(to_plot) + geom_point(aes(x=pseudotime, y=perc, fill=Subtype), shape=21, size=3) + theme_classic() +
  scale_fill_manual(values = palette_subtypes) + facet_grid(rows = vars(isoform)) + xlab("Disease Progression") + ylab("transcript/gene count")
dev.off()


### PKM1 -> adult tissues, PKM2 -> embryonic/tumor
pkm = tx_info[tx_info$gene_id == g_info$gene_id[g_info$gene_name == "PKM"] & tx_info$transcript_type == "protein_coding",]
pkm$maxExp = rowMax(ncount.txi[pkm$transcript_id,mdata$Name[onlyPE]])
pkm = pkm[order(pkm$maxExp, decreasing = T),]
pkm = pkm[pkm$length > 2000,]

to_plot = NULL
variants = list("PKM1"="ENST00000319622.10", "PKM2"="ENST00000335181.10")
for ( variant in names(variants)){
  tmp = mdata[onlyPE,c("Name", "PC.Type", "Subtype", "pseudotime")]
  tmp$isoform = variant
  tmp$ncount = ncount.txi[variants[[variant]], tmp$Name]
  tmp$perc = perc_tx[variants[[variant]], tmp$Name]
  to_plot = rbind(to_plot, tmp)
}

pdf(paste0(outdir, "PKM.pdf"))
to_plot$Subtype = factor(to_plot$Subtype, levels= c("NORMAL", "PRIMARY", "ARPC", "DNPC", "NEPC"))
ggplot(to_plot) + geom_boxplot(aes(x=Subtype, fill=isoform, y=ncount)) + theme_classic() + scale_fill_manual(values = c("blue", "red"))+ylab("transcript CPM")
p1 = ggplot(to_plot[to_plot$isoform == "PKM1",], aes(x=Subtype, fill=isoform, y=ncount)) + geom_boxplot()  + geom_signif(comparisons = list(c("PRIMARY", "ARPC"), c("NORMAL", "PRIMARY"), c("DNPC", "NEPC")), test = "t.test")+
  theme_classic() + scale_fill_manual(values = c("red"))+ylab("transcript CPM") +
  facet_grid(rows=vars(isoform))
p2 = ggplot(to_plot[to_plot$isoform == "PKM2",], aes(x=Subtype, fill=isoform, y=ncount)) + geom_boxplot()  + geom_signif(comparisons = list(c("PRIMARY", "ARPC"), c("NORMAL", "PRIMARY"), c("DNPC", "NEPC")), test = "t.test")+
  theme_classic() + scale_fill_manual(values = c("blue"))+ylab("transcript CPM") +
  facet_grid(rows=vars(isoform))
grid.arrange(p1, p2, nrow=2)

ggplot(to_plot) + geom_point(aes(x=pseudotime, y=ncount, fill=Subtype), shape=21, size=3) + theme_classic() +
  scale_fill_manual(values = palette_subtypes) + facet_grid(rows = vars(isoform)) + xlab("Disease Progression") + ylab("transcript CPM")
ggplot(to_plot) + geom_point(aes(x=pseudotime, y=perc, fill=Subtype), shape=21, size=3) + theme_classic() +
  scale_fill_manual(values = palette_subtypes) + facet_grid(rows = vars(isoform)) + xlab("Disease Progression") + ylab("transcript/gene count")
dev.off()


### AR V-7 analysis
palette_subtypes = c("#25681E", "#F63A21","#4889C7", "#1010C0", "#93355A")
gene_id = g_info$gene_id[g_info$gene_name == "AR"]
pkm = tx_info[tx_info$gene_id == gene_id ,]
pkm$maxExp = rowMax(ncount.txi[pkm$transcript_id,mdata$Name[onlyPE]])
pkm$mean = rowMeans(ncount.txi[pkm$transcript_id,mdata$Name[onlyPE]])
pkm = pkm[order(pkm$maxExp, decreasing = T),]
pkm

to_plot = NULL
variants = list("AR-dLBD"=c("ENST00000504326.5", "ENST00000613054.2","ENST00000514029.5"), "AR-LBD"=c("ENST00000374690.9", "ENST00000396043.3", "ENST00000612452.5"))
for ( variant in names(variants)){
  tmp = mdata[onlyPE,c("Name", "PC.Type", "Subtype", "pseudotime")]
  tmp$isoform = variant
  if ( length(variants[[variant]]) == 1 ){
    tmp$ncount = ncount.txi[variants[[variant]], tmp$Name]
    tmp$perc = perc_tx[variants[[variant]], tmp$Name]
  } else {
    tmp$ncount = colSums(txi.tx$counts[variants[[variant]], tmp$Name])
    tmp$perc = as.numeric(tmp$ncount/txi.gene$counts[gene_id, tmp$Name])
    tmp$ncount = log2( (tmp$ncount / (colSums(txi.tx$counts[,tmp$Name]) / 1000000))+1 )
  }
  to_plot = rbind(to_plot, tmp)
}

pdf(paste0(outdir, "AR_v7_grouped.pdf"))
to_plot$Subtype = factor(to_plot$Subtype, levels= c("NORMAL", "PRIMARY", "ARPC", "DNPC", "NEPC"))
ggplot(to_plot) + geom_boxplot(aes(x=Subtype, fill=isoform, y=ncount)) + theme_classic() + scale_fill_manual(values = c("blue", "red"))+ylab("transcript CPM")
ggplot(to_plot) + geom_point(aes(x=pseudotime, y=ncount, fill=Subtype), shape=21, size=3) + theme_classic() +
  scale_fill_manual(values = palette_subtypes) + facet_grid(rows = vars(isoform)) + xlab("Disease Progression") + ylab("transcript CPM")
ggplot(to_plot) + geom_point(aes(x=pseudotime, y=perc, fill=Subtype), shape=21, size=3) + theme_classic() +
  scale_fill_manual(values = palette_subtypes) + facet_grid(rows = vars(isoform)) + xlab("Disease Progression") + ylab("transcript/gene count")
dev.off()


### group similar isoforms PKM1 -> adult tissues, PKM2 -> embryonic/tumor
gene_id = g_info$gene_id[g_info$gene_name == "PKM"]
pkm = tx_info[tx_info$gene_id == gene_id & tx_info$transcript_type == "protein_coding",]
pkm$maxExp = rowMax(ncount.txi[pkm$transcript_id,mdata$Name[onlyPE]])
pkm = pkm[order(pkm$maxExp, decreasing = T),]
pkm = pkm[pkm$length > 2000,]

to_plot = NULL
variants = list("PKM-EX9"=c("ENST00000319622.10", "ENST00000565184.6", "ENST00000389093.7"), "PKM-EX10"=c("ENST00000335181.10"))
variant="PKM1"
for ( variant in names(variants)){
  tmp = mdata[onlyPE,c("Name", "PC.Type", "Subtype", "pseudotime")]
  tmp$isoform = variant
  if ( length(variants[[variant]]) == 1 ){
    tmp$ncount = ncount.txi[variants[[variant]], tmp$Name]
    tmp$perc = perc_tx[variants[[variant]], tmp$Name]
  } else {
    tmp$ncount = colSums(txi.tx$counts[variants[[variant]], tmp$Name])
    tmp$perc = as.numeric(tmp$ncount/txi.gene$counts[gene_id, tmp$Name])
    tmp$ncount = log2( (tmp$ncount / (colSums(txi.tx$counts[,tmp$Name]) / 1000000))+1 )
  }
  to_plot = rbind(to_plot, tmp)
}

pdf(paste0(outdir, "PKM_grouped.pdf"))
to_plot$Subtype = factor(to_plot$Subtype, levels= c("NORMAL", "PRIMARY", "ARPC", "DNPC", "NEPC"))
ggplot(to_plot) + geom_boxplot(aes(x=Subtype, fill=isoform, y=ncount)) + theme_classic() + scale_fill_manual(values = c("blue", "red"))+ylab("transcript CPM")+
  facet_grid(rows=vars(isoform))
ggplot(to_plot) + geom_boxplot(aes(x=Subtype, fill=isoform, y=perc)) + theme_classic() + scale_fill_manual(values = c("blue", "red"))+ ylab("transcript/gene count")+
  facet_grid(rows=vars(isoform))
ggplot(to_plot) + geom_point(aes(x=pseudotime, y=ncount, fill=Subtype), shape=21, size=3) + theme_classic() +
  scale_fill_manual(values = palette_subtypes) + facet_grid(rows = vars(isoform)) + xlab("Disease Progression") + ylab("transcript CPM")
ggplot(to_plot) + geom_point(aes(x=pseudotime, y=perc, fill=Subtype), shape=21, size=3) + theme_classic() +
  scale_fill_manual(values = palette_subtypes) + facet_grid(rows = vars(isoform)) + xlab("Disease Progression") + ylab("transcript/gene count")
dev.off()

###

tcga_psa = read.table("./metadata/nationwidechildrens.org_clinical_patient_prad.tsv", sep="\t", header=T)
pdf("./psa_levels_pseudotime.pdf")
ggplot(mdata.tcga) + geom_point(aes(x=pseudotime, y=log10(psa_value+1)))+ theme_bw()
ggplot(mdata.tcga) + geom_point(aes(x=pseudotime, y=psa_value))+ theme_bw()
dev.off()
summary(factor(mdata.tcga$biochemical_recurrence))
### ERF-202 expression
isoform_name = "ERF-202"
isoform_id = tx_names$transcript_id[tx_names$transcript_name == isoform_name]
mdata.tcga = mdata.tcga[mdata.tcga$biochemical_recurrence %in% c("NO", "YES"),]
to_plot = mdata.tcga
to_plot$ncount = ncount.txi[isoform_id, to_plot$Name]

pdf(paste0(outdir, "/BiochemicalRecurrence_ERF_202.pdf"))
ggplot(to_plot) + geom_boxplot(aes(x=biochemical_recurrence, y=ncount, fill=biochemical_recurrence)) + theme_classic() +
  ylab(paste0(isoform_name, " CPM normalized counts"))
dev.off()
```
