## Supplementary material for "An integrated, transcript-resolution atlas of prostate cancer progression unmasks associations with oncogenic pathways": Reproducible code: 06_IsoformSwitch.R

```
#!/usr/bin/Rscript
# Public submission copy.
# Analytical statements and parameter values are unchanged from the archived source.
# Legacy non-English comments and console messages were normalized for public release.
library(msigdbr, quietly = T)
library(tidyverse)
library("clusterProfiler")
library(ggrepel)
library(DESeq2)
library("GSVA", quietly = T)
library(sva, quietly = T)
source("./00_functions.R")

ncount = readRDS("./objects/ncount.RDS")
ncount.tx = readRDS("./objects/ncount.tx.RDS")
mdata=readRDS("./objects/mdata_signatures.RDS")
rownames(mdata)=mdata$Name
tx_info = read.csv("./metadata/transcripts_info.csv")
tx_to_protein = read.table("./metadata/transcript_protein.tsv",  col.names = c("transcript_id", "protein_id", "transcript_name", "CDS"))
tx_info = merge(tx_info, tx_to_protein, by="transcript_id", all.x=T)
g_info= read.csv("./metadata/genes_info.csv")
txi.tx.counts = readRDS("./objects/txi.tx.counts.RDS")
txi.gene.counts = readRDS("./objects/txi.gene.counts.RDS")
outdir="./results_06_IsoformSwitch/"

filters = list(
  samples = mdata$Name[mdata$LibraryLayout == "PAIRED"]
)

### Identify possible interesting transcripts by selecting samples having low/high ssGSEA
if ( file.exists("./objects/mdata.ssgsea.RDS")){
  mdata.ssgsea = readRDS("./objects/mdata.ssgsea.RDS")
  CS.names = readRDS("./objects/CS.names.RDS")
} else {
  hallmarks = as.data.frame(msigdbr(species = "Homo sapiens", category = "H"))
  hallmarks = rbind(hallmarks, as.data.frame(msigdbr(species = "Homo sapiens", category = "C2", subcategory = "PID")))
  hallmarks = rbind(hallmarks, as.data.frame(msigdbr(species = "Homo sapiens", category = "C6")))
  CS = list()
  # AR-SCORE NELSON (Bluemn et al., 2017) ---->
  CS$AR_GSET <- g_info$gene_id[g_info$gene_name %in% c("KLK3","KLK2","TMPRSS2","FKBP5","NKX3-1","PLPP1","PMEPA1","PART1","ALDH1A3","STEAP4")]
  # NE-SCORE NELSON (Bluemn et al., 2017) ---->
  CS$NE_GSET <- g_info$gene_id[g_info$gene_name %in% c("SYP","CHGA","CHGB","ENO2","CHRNB2","SCG3","SCN3A","PCSK1","ELAVL4","NKX2-1")]
  for ( h in unique(hallmarks$gs_name)){
    CS[[h]]=g_info$gene_id[g_info$gene_name %in% hallmarks$gene_symbol[hallmarks$gs_name == h]]
  }
  length(CS)
  mdata = mdata[filters$samples,]
  keep = rowSums(ncount[,mdata$Name] > 0) > nrow(mdata)*0.05
  sum(keep) / length(keep)
  THREADS=8
  res = gsva(as.matrix(ncount[keep,mdata$Name,drop=F]), CS, mx.diff=TRUE, method = "ssgsea", parallel.sz=THREADS, ssgsea.norm=F)
  res=t(res)
  res = as.data.frame(res)
  res$Name = rownames(res)
  colnames(mdata)
  mdata.ssgsea = merge(mdata[,c("Name", "pseudotime", "Subtype","ESTIMATE.Immune.Score", "ESTIMATE.Stromal.Score" )], res, by="Name")
  out_dir = paste0(outdir,"ssGSEA/")

  dir.create(out_dir, showWarnings = F, recursive = T)
  pdf(paste0(out_dir, "all_ssGSEA_hallmark_and_custom.pdf"))
  for ( name in names(CS)){
    p<-ggplot(data.frame(score=  mdata.ssgsea[,name], pseudotime = mdata.ssgsea$pseudotime, subtype = mdata.ssgsea$Subtype))+geom_point(aes(x=pseudotime, y=score, color=subtype)) +ggtitle(name)
    print(p)
  }
  dev.off()
  CS.names = names(CS)
  saveRDS(mdata.ssgsea, "./objects/mdata.ssgsea.RDS")
  saveRDS(CS.names, "./objects/CS.names.RDS")
}


### Only protein coding genes and transcripts

tx_info = tx_info[tx_info$transcript_type == "protein_coding",]

if ( file.exists("./objects/tx.contributions.RDS")){
  tx.contributions = readRDS("./objects/tx.contributions.RDS")
} else {
  tx.contributions = t(sapply(tx_info$transcript_id, function(x){txi.tx.counts[x,mdata.ssgsea$Name] / txi.gene.counts[tx_info$gene_id[tx_info$transcript_id == x], mdata.ssgsea$Name] } ))
  tx.contributions[is.na(tx.contributions)] = 0
  saveRDS(tx.contributions, "./objects/tx.contributions.RDS")
}

###
performIsoformSelection = function(metadata ,  out_dir , comparisons, tx_thr = 0.5, g_thr = 1, cor_method = "pearson"){
  out_dat = NULL
  dir.create(out_dir, showWarnings = F)
  tx.perc = tx.contributions[ rowSums(tx.contributions[,metadata$Name] > 0.4) > nrow(metadata) *0.05, metadata$Name ]
  tx.perc = tx.perc[rownames(tx.perc) %in% rownames(txi.tx.counts)[rowSums(txi.tx.counts[,metadata$Name] > 1) >  nrow(metadata) *0.05 ], metadata$Name]
  name= comparisons[1]
  tx.corr.base = merge(tx_info[,c("transcript_id", "gene_id", "transcript_type", "protein_id", "transcript_name", "CDS")], g_info[,c("gene_id", "gene_name")])
  tx.corr.base = tx.corr.base[tx.corr.base$transcript_id %in% rownames(tx.perc),]
  for ( name in comparisons ){
    target_val = metadata[,name]
    tx.corr = tx.corr.base %>% rowwise() %>% filter(sd(tx.perc[transcript_id, metadata$Name]) != 0) %>%
      mutate(corr = cor(target_val, tx.perc[transcript_id,metadata$Name], method = cor_method)) %>% filter(abs(corr) > tx_thr)
    if ( nrow( tx.corr ) > 0 ) {
      tx.corr = tx.corr %>% group_by(gene_id) %>% mutate(gcorr = cor(target_val, ncount[unique(gene_id),metadata$Name], method = cor_method) ,
                                                         n_tx = n(), n_tx_pos = sum(corr > tx_thr), n_tx_neg = sum(corr < -tx_thr), max_corr = max(corr), min_corr = min(corr), delta_corr = max_corr - min_corr  )
      tx.corr.filtered = tx.corr %>% filter(n_tx_pos > 0 & n_tx_neg > 0 & abs(gcorr) <= g_thr ) %>% group_by(gene_id) %>% mutate(n_tx_switch = n(), n_prot = ifelse( any(is.na(CDS)), length(unique(protein_id)), length(unique(CDS)) ) ) %>%
        filter(n_tx_switch > 1 & n_prot > 1)
      print(paste(name, nrow(tx.corr.filtered)))
      if ( nrow(tx.corr.filtered) > 0 ) {
        tx.corr.filtered = tx.corr.filtered[order(tx.corr.filtered$delta_corr, decreasing = T),]
        tx.corr.filtered$target = name
        subtype="NORMAL"
        for ( subtype in c("NORMAL", "PRIMARY", "METASTATIC")){
          if ( subtype == "METASTATIC") {
            sampleFilter = metadata$Subtype %in% c("ARPC", "DNPC", "NEPC")
          } else {
            sampleFilter = metadata$Subtype == subtype
          }
          tx.corr.filtered = tx.corr.filtered %>% rowwise() %>% mutate(st_corr = cor(target_val[sampleFilter], tx.perc[transcript_id, metadata$Name[sampleFilter]])) %>%
            group_by(gene_id) %>% mutate(st_valid = min(st_corr) < - tx_thr & max(st_corr) > tx_thr, st_delta = max(st_corr) - min(st_corr))
          if ( any(is.na(tx.corr.filtered$st_valid))){
            tx.corr.filtered$st_valid[is.na(tx.corr.filtered$st_valid)]=F
          }
          if ( any(is.na(tx.corr.filtered$st_delta))){
            tx.corr.filtered$st_delta[is.na(tx.corr.filtered$st_delta)]=0
          }
          if ( any(is.na(tx.corr.filtered$st_corr))){
            tx.corr.filtered$st_corr[is.na(tx.corr.filtered$st_corr)]=0
          }
          colnames(tx.corr.filtered)[grepl("st_",colnames(tx.corr.filtered))] = paste0(subtype, c("_corr", "_valid", "_delta"))
        }
        out_dat = rbind(out_dat, tx.corr.filtered)
      }
    }
  }
  return(out_dat)
}


plotIsoformSwitchByGene = function(isoSwitchRes,metadata, out_dir, out_gene_list, top_n = 10){
  dir.create(out_dir, showWarnings = F)
  for ( gene_name in out_gene_list){
    print(gene_name)
    pdf(paste0(out_dir, gene_name, ".pdf"), width = 10)
    gene_id = unique(isoSwitchRes$gene_id[isoSwitchRes$gene_name == gene_name])
    print(gene_id)
    full_gene_list = isoSwitchRes[isoSwitchRes$gene_id == gene_id,] %>% group_by(gene_id, delta_corr, gcorr, target) %>% reframe() %>% arrange(desc(delta_corr))
    names_to_plot = full_gene_list$target[1:min(top_n, nrow(full_gene_list))]
    print(names_to_plot)
    if ( "pseudotime" %in% full_gene_list$target ){
      names_to_plot = names_to_plot[names_to_plot != "pseudotime"]
      names_to_plot = c("pseudotime", names_to_plot)
    }
    for ( name in names_to_plot){
      tx.corr.filtered = isoSwitchRes[isoSwitchRes$target == name & isoSwitchRes$gene_name == gene_name, ]
      gene_list = tx.corr.filtered %>% group_by(gene_id, delta_corr, gcorr, target) %>% arrange(desc(delta_corr)) %>% reframe()
      to_plot = data.frame(
        count = ncount[gene_id, metadata$Name],
        score = metadata[, name],
        subtype = metadata$Subtype,
        ratio = 1,
        id =  gene_name
      )

      txs_ids=tx.corr.filtered$transcript_id[tx.corr.filtered$gene_id == gene_id]
      tx_title = name
      for ( tx in txs_ids){
        tx_title = paste0(tx_title, "\n", tx, " - ", tx_info$transcript_name[tx_info$transcript_id == tx], " PCC = ", round(tx.corr.filtered$corr[tx.corr.filtered$transcript_id == tx], 2))
        to_plot = rbind(to_plot, data.frame(
          count = ncount.tx[tx, metadata$Name],
          score = metadata[, name],
          ratio = as.numeric(tx.contributions[tx, metadata$Name]),
          subtype = metadata$Subtype,
          id = tx
        ))
      }
      to_plot$id = factor(to_plot$id, levels = c(gene_name, txs_ids))
      to_plot$subtype = factor(to_plot$subtype, levels = c("NORMAL", "PRIMARY", "ARPC", "DNPC", "NEPC"))
      p<- ggplot(to_plot,aes(x=score, y=count)) + geom_point(aes(fill = subtype), shape=21, size=3) + geom_smooth(method = 'gam', formula = y ~s(x, bs="cs"))+
        xlab(paste0(name, " score")) + facet_wrap(vars(id)) + scale_fill_manual(values=palette_subtypes)+
        ggtitle(paste0(gene_name, "\nDelta corr= ", round(gene_list$delta_corr[gene_list$gene_id == gene_id],2), "\nGene corr= ", round(gene_list$gcorr[gene_list$gene_id == gene_id],2) ))+
        theme_classic()
      print(p)
      to_plot = to_plot[to_plot$id != gene_name,]
      to_plot$id = factor(to_plot$id, levels = txs_ids)
      to_plot$subtype = factor(to_plot$subtype, levels = c("NORMAL", "PRIMARY", "ARPC", "DNPC", "NEPC"))
      p<- ggplot(to_plot,aes(x=score, y=ratio)) + geom_point(aes(fill=subtype), shape=21, size=3) + geom_smooth(method = 'gam', formula = y ~s(x, bs="cs"))+
        xlab(paste0(name, " score")) + facet_wrap(vars(id)) + ggtitle(tx_title)+
        theme_classic() + scale_fill_manual(values=palette_subtypes)
      print(p)
    }
    dev.off()
  }
}


### Filter genes sets

all_classes = c("pseudotime",  "ESTIMATE.Immune.Score", "ESTIMATE.Stromal.Score" , CS.names)

dat = mdata.ssgsea %>% pivot_longer(all_of(all_classes), names_to = "gset", values_to = "score") %>% group_by(gset) %>% mutate(rescaled.score = ((score-min(score))*100)/(max(score)-min(score)) )
dat.summ = dat %>% group_by(gset) %>% summarise(sd = sd(rescaled.score)) %>% arrange(desc(sd))
dat.summ$gset = factor(dat.summ$gset, levels = dat.summ$gset)
dat$gset = factor(dat$gset, levels = dat.summ$gset)
if ( ! file.exists(paste0(outdir,"/ssGSEA/standard_deviations.pdf"))){
  dir.create(paste0(outdir,"/ssGSEA/"), showWarnings = F, recursive = T)
  pdf(paste0(outdir,"/ssGSEA/standard_deviations.pdf"))
  ggplot(dat.summ)+ geom_point(aes(x=gset, y=sd))
  dev.off()
  pdf(paste0(outdir, "/ssGSEA/ssGSEA_vs_pseudotime.pdf"))
  i=2
  for ( i in 1:nrow(dat.summ) ){
    gsname = as.character(dat.summ$gset[i])
    p<-ggplot(data.frame(pseudotime = mdata.ssgsea$pseudotime, score =mdata.ssgsea[,gsname] , Subtype=mdata.ssgsea$Subtype)) +
      geom_point(aes(x=pseudotime, y=score, color=Subtype))+theme_bw()+ ylab(gsname) + ggtitle(paste0(i, " - " , gsname, "\n SD = ", dat.summ$sd[i]))
    print(p)
  }
  dev.off()
}


### Filter the classes with sd < 10
filtered_classes = as.character(dat.summ$gset[dat.summ$sd > 10])

if ( file.exists(paste0(outdir,"/IsoformSwitchAll/all_results.csv"))){
  all_res = read.csv(paste0(outdir,"/IsoformSwitchAll/all_results.csv"))
} else {
  all_res = performIsoformSelection(metadata = mdata.ssgsea, out_dir = paste0(outdir,"/IsoformSwitchAll/"),comparisons = filtered_classes,tx_thr =  0.5, g_thr = 1, cor_method = "pearson")
  write.csv(all_res , paste0(outdir,"/IsoformSwitchAll/all_results.csv"), row.names = F)
}
### In GENCODE v39 MTOR-202 is annotatated as protein coding but it's not ( It's retained_intron )
all_res= all_res[all_res$gene_name != "MTOR",]
write.csv(all_res , paste0(outdir,"/IsoformSwitchAll/all_results.csv"), row.names = F)

all_res.bygene = all_res %>% mutate(transcript = paste0(transcript_id, " (", transcript_name, ")")) %>% group_by(gene_id, gene_name) %>%
  mutate(n_targets = length(unique(target)), n_NORMAL = length(unique(target[NORMAL_valid])), n_PRIMARY = length(unique(target[PRIMARY_valid])),
         n_MET =length(unique(target[METASTATIC_valid])),  best_target = target[which(delta_corr == max(delta_corr))[1]], pseudotime_target = "pseudotime" %in% target  ) %>%
  ungroup() %>% filter(target == best_target) %>%
  group_by(gene_id, gene_name, n_targets, best_target, delta_corr, pseudotime_target, n_NORMAL, n_PRIMARY, n_MET) %>%
  summarise(iso_up_id = transcript_id[which(corr  == max(corr))] , iso_up_name = transcript_name[which(corr  == max(corr))],  iso_down_id = transcript_id[which(corr == min(corr))],  iso_down_name = transcript_name[which(corr == min(corr))], iso_up_corr = max(corr), iso_down_corr = min(corr) ) %>%
  arrange(desc(n_targets))


write.csv(all_res.bygene, paste0(outdir,"/IsoformSwitchAll/all_results_by_gene.csv"), row.names = F)

## plot interesting genes
candidates = read.table("./metadata/candidates.txt")$V1
all_res.bygene = all_res.bygene[order(all_res.bygene$delta_corr, decreasing = T),]
gene_list = unique(c(candidates,  all_res.bygene$gene_name[1:10], all_res$gene_name[all_res$target == "pseudotime"]))
sum(!gene_list %in% all_res$gene_name)
gene_list = gene_list[gene_list %in% all_res$gene_name]
length(gene_list)
plotIsoformSwitchByGene(isoSwitchRes = all_res, metadata = mdata.ssgsea, out_dir = paste0(outdir, "/IsoformSwitchAll/Genes/"), out_gene_list = gene_list)

## lollipop of the targets with the highest number of isoform switch
colnames(all_res)
all_res.bytarget = all_res %>% select(gene_id, delta_corr, target, NORMAL_valid, PRIMARY_valid, METASTATIC_valid) %>%
  distinct() %>% group_by(target) %>% summarise(isoform_switches = n(), avg_delta_corr = mean(delta_corr),
                                                n_NORMAL = sum(NORMAL_valid), n_PRIMARY = sum(PRIMARY_valid), n_METASTATIC = sum(METASTATIC_valid) ) %>% arrange(isoform_switches)

all_res.bytarget = merge(all_res.bytarget, dat.summ, by.x="target", by.y="gset")

write.csv(all_res.bytarget,  paste0(outdir,"/IsoformSwitchAll/all_results_by_target.csv"), row.names = F)


sum(all_res.bytarget$isoform_switches > 100)
hallmarks = as.data.frame(msigdbr(species = "Homo sapiens", category = "H"))
hallmarks = rbind(hallmarks, as.data.frame(msigdbr(species = "Homo sapiens", category = "C2", subcategory = "PID")))
hallmarks = rbind(hallmarks, as.data.frame(msigdbr(species = "Homo sapiens", category = "C6")))
str(hallmarks)
hallmarks = unique(hallmarks[,c("gs_name", "gs_cat")])
all_res.bytarget = merge(all_res.bytarget, hallmarks, by.x="target", by.y="gs_name", all.x=T, all.y=F)
pdf(paste0(outdir, "/IsoformSwitchAll/sd_vs_switches.pdf"))
ggplot(all_res.bytarget) + geom_point(aes(x=sd, y=isoform_switches, color=gs_cat))
dev.off()
all_res.bytarget= all_res.bytarget %>% rowwise() %>% mutate(top_influence = c("NORMAL", "PRIMARY", "METASTATIC")[which.max(c(n_NORMAL, n_PRIMARY, n_METASTATIC))])
all_res.bytarget$top_influence = factor(all_res.bytarget$top_influence,levels = c("NORMAL", "PRIMARY", "METASTATIC") )
all_res.bytarget = all_res.bytarget %>% arrange(isoform_switches)
all_res.bytarget$target = factor(all_res.bytarget$target, levels=all_res.bytarget$target)

pdf(paste0(outdir, "top_isoform_switches.pdf"), width=8, height = 6)
ggplot(all_res.bytarget[all_res.bytarget$isoform_switches > 100,], aes(x=isoform_switches, y=target)) +
  geom_segment(aes(x=0, xend=isoform_switches, y=target, yend=target))+
  geom_point(aes(size=avg_delta_corr, fill=top_influence), shape=21) +
  scale_fill_manual(values=palette_subtypes)+
  theme_classic()
dev.off()
pdf(paste0(outdir, "top_isoform_switches_2.pdf"), width=8, height = 10)
all_res.bytarget.small = all_res.bytarget %>% filter(!is.na(gs_cat))  %>% group_by(gs_cat) %>% slice_max(isoform_switches, n=20)
ggplot(all_res.bytarget.small, aes(x=isoform_switches, y=target)) +
  geom_segment(aes(x=0, xend=isoform_switches, y=target, yend=target))+
  geom_point(aes(size=avg_delta_corr, fill=top_influence), shape=21) +
  scale_fill_manual(values=palette_subtypes)+
  facet_grid(vars(gs_cat),scales = "free")+
  theme_classic()
dev.off()
```
