## Supplementary material for "An integrated, transcript-resolution atlas of prostate cancer progression unmasks associations with oncogenic pathways": Reproducible code: canonical_noncanonical_pathway_enrichment.R

```
# Public submission copy.
# Analytical statements and parameter values are unchanged from the archived source.
# Legacy non-English comments and console messages were normalized for public release.
# ==============================================================================
# ==============================================================================
library(tidyverse)
library(clusterProfiler)
library(msigdbr)
library(enrichplot)

patients <- read.csv("tx_info_with_pseudotime_corr.csv")

patients_pc <- patients %>%
  filter(transcript_type == "protein_coding")

m_t2g <- msigdbr(species = "Homo sapiens", category = "H") %>%
  dplyr::select(gs_name, gene_symbol)

# ==============================================================================
# ==============================================================================

genes_noncanon_driven <- patients_pc %>%
  group_by(gene_id) %>%
  filter({
    vals <- cor_pseudotime
    canon_idx <- which(is_canonical == 1)

    if (length(canon_idx) == 0) {
      FALSE
    } else {
      val_canon <- vals[canon_idx[1]]
      val_max <- max(vals, na.rm = TRUE)
      val_min <- min(vals, na.rm = TRUE)

      is_not_extreme <- (val_canon != val_max) & (val_canon != val_min)
      has_significant_correlation <- (val_max > 0.4) | (val_min < -0.4)

      is_not_extreme & has_significant_correlation
    }
  }) %>%
  pull(gene_name) %>%
  unique()


genes_canon_driven <- patients_pc %>%
  group_by(gene_id) %>%
  filter({
    vals <- cor_pseudotime
    canon_idx <- which(is_canonical == 1)

    if (length(canon_idx) == 0) {
      FALSE
    } else {
      val_canon <- vals[canon_idx[1]]
      val_max <- max(vals, na.rm = TRUE)
      val_min <- min(vals, na.rm = TRUE)

      is_extreme <- (val_canon == val_max) | (val_canon == val_min)
      is_significant <- abs(val_canon) > 0.4

      is_extreme & is_significant
    }
  }) %>%
  pull(gene_name) %>%
  unique()

cat("Analysis progress", length(genes_noncanon_driven), "\n")
cat("Analysis progress", length(genes_canon_driven), "\n")


# ==============================================================================
# ==============================================================================

plot_hallmark_pdfs <- function(gene_list, prefix) {

  if(length(gene_list) == 0) {
    message(paste("➡️ [", prefix, "Analysis warning; step skipped"))
    return()
  }

  res <- enricher(gene = gene_list,
                  TERM2GENE = m_t2g,
                  pvalueCutoff = 0.05,
                  qvalueCutoff = 0.2)

  if (is.null(res) || nrow(as.data.frame(res)) == 0) {
    message(paste("➡️ [", prefix, "No significant enrichment was detected"))
  } else {
    message(paste("➡️ [", prefix, "Significant pathways detected"))

    p1 <- dotplot(res, showCategory = 15) +
      ggtitle(paste(prefix, "Enrichment (Dotplot)")) +
      theme(axis.text.y = element_text(size = 10))
    ggsave(paste0(prefix, "_Hallmark_dotplot.pdf"), p1, width = 8, height = 6)

    p2 <- barplot(res, showCategory = 15) +
      ggtitle(paste(prefix, "Enrichment (Barplot)"))
    ggsave(paste0(prefix, "_Hallmark_barplot.pdf"), p2, width = 8, height = 6)

    try({
      p3 <- cnetplot(res, showCategory = 5, circular = FALSE, colorEdge = TRUE) +
        ggtitle(paste(prefix, "Enrichment (Cnetplot)"))
      ggsave(paste0(prefix, "_Hallmark_cnetplot.pdf"), p3, width = 10, height = 8)
    }, silent = TRUE)
  }
}

# ==============================================================================
# ==============================================================================

plot_hallmark_pdfs(genes_noncanon_driven, "NonCanonical_Driven")

plot_hallmark_pdfs(genes_canon_driven, "Canonical_Driven")

message("Analysis completed and output saved")
```
