## Supplementary material for "An integrated, transcript-resolution atlas of prostate cancer progression unmasks associations with oncogenic pathways": Reproducible code: canonical_rank_plot_rasterized.R

```
# Public submission copy.
# Analytical statements and parameter values are unchanged from the archived source.
# Legacy non-English comments and console messages were normalized for public release.
# ==========================================
# ==========================================
library(openxlsx)
library(ggplot2)
library(dplyr)
library(ggrepel)
library(ggrastr)

# ==========================================
# ==========================================
excel_file <- "lncRNA_with_pseudotime_corr 2.xlsx"
sheet_index <- 1

# ==========================================
# ==========================================
cat("Loading input data\n")
df <- read.xlsx(excel_file, sheet = sheet_index)

# ==========================================
# ==========================================
up_indices <- c(2,7,13,14,15,21,22,25,27,28,29,31,32,38,41,42,44,46,49,51,50384,50387,50394,50400,50404,50407,50409) - 1
down_indices <- c(12,50381,50382,50385,50388,50389,50392,50397,50398,50403,50410,50411,50413,50414) - 1

df$group <- "Background"

valid_up <- up_indices[up_indices > 0 & up_indices <= nrow(df)]
valid_down <- down_indices[down_indices > 0 & down_indices <= nrow(df)]

df$group[valid_up] <- "Up_Red"
df$group[valid_down] <- "Down_Blue"

# ==========================================
# ==========================================
plot_data <- df %>%
  arrange((cor_pseudotime)) %>%
  mutate(rank = 1:n())

highlight_data <- plot_data %>% filter(group != "Background")

# ==========================================
# ==========================================
p <- ggplot(plot_data, aes(x = cor_pseudotime, y = rank)) +

  rasterise(geom_point(data = subset(plot_data, group == "Background"),
                       color = "grey85", size = 1, alpha = 0.3), dpi = 300) +

  rasterise(geom_point(data = highlight_data,
                       aes(color = group),
                       size = 3, alpha = 1), dpi = 300) +

  scale_color_manual(values = c("Up_Red" = "#E41A1C",
                                "Down_Blue" = "#004c8c")) +

  geom_text_repel(
    data = highlight_data,
    aes(label = gene_name, color = group),
    size = 4,
    fontface = "bold",
    box.padding = 0.6,
    max.overlaps = Inf
  ) +

  geom_vline(xintercept = 0, linetype = "dashed", color = "grey40") +

  labs(
    title = "Sigmoid Correlation Rank Plot",
    x = "Correlation Coefficient",
    y = "Rank"
  ) +
  scale_x_continuous(limits = c(-1, 1), breaks = seq(-1, 1, 0.5)) +
  scale_y_continuous(limits = c(-10000, 60000)) +
  theme_classic() +
  theme(
    legend.position = "none",
    plot.title = element_text(hjust = 0.5, face = "bold")
  )

print(p)

ggsave("sigmoid_plot_rasterized.pdf", width = 10, height = 4, dpi = 300)


# ==========================================
# ==========================================
plot_data_all <- df %>%
  arrange(cor_pseudotime) %>%
  mutate(rank = 1:dplyr::n())

target_genes <- c("MALAT1", "NEAT1", "FIRRE", "FENDRR")

for (target in target_genes) {

  cat(paste0("Running analysis step", target, " ...\n"))

  current_plot_data <- plot_data_all %>%
    mutate(group = case_when(
      gene_name == target & is_canonical == 1 ~ "Target_Canonical",
      gene_name == target ~ "Target_Isoform",
      TRUE ~ "Background"
    ))

  canonical_data <- subset(current_plot_data, group == "Target_Canonical")
  isoform_data <- subset(current_plot_data, group == "Target_Isoform")

  p_loop <- ggplot(current_plot_data, aes(x = cor_pseudotime, y = rank)) +

    rasterise(geom_point(data = subset(current_plot_data, group == "Background"),
                         color = "grey90", size = 1, alpha = 0.3), dpi = 300) +

    rasterise(geom_point(data = isoform_data,
                         color = "black", size = 3, alpha = 0.8), dpi = 300) +

    rasterise(geom_point(data = canonical_data,
                         color = "#E41A1C", size = 4, alpha = 1), dpi = 300) +

    geom_text_repel(
      data = canonical_data,
      aes(label = gene_name),
      color = "#E41A1C",
      size = 5,
      fontface = "bold",
      box.padding = 0.8,
      min.segment.length = 0
    ) +

    geom_vline(xintercept = 0, linetype = "dashed", color = "grey40") +
    labs(
      title = paste0("Sigmoid Plot: ", target),
      x = "Correlation Coefficient",
      y = "Global Rank"
    ) +
    scale_x_continuous(limits = c(-1, 1), breaks = seq(-1, 1, 0.5)) +
    theme_classic() +
    theme(
      plot.title = element_text(hjust = 0.5, face = "bold")
    )

  print(p_loop)

  ggsave(paste0("Sigmoid_", target, "_Canonical_rasterized.pdf"), p_loop, width = 8, height = 5, dpi = 300)
}

cat("Analysis completed and output saved\n")


# ==========================================
# ==========================================

# library(openxlsx)
# library(ggplot2)
# library(dplyr)
# library(ggrepel)
# library(ggrastr)

excel_file_new <- "pc_tx_info_filter_5pct_JP.xlsx"
cat(paste0("Loading input data", excel_file_new, " ...\n"))
df_new <- read.xlsx(excel_file_new, sheet = 1)

df_new$group <- "Background"
target_rows <- 1:21

if(!"is_canonical" %in% colnames(df_new)) {
  stop("Analysis failed")
}

idx_canonical <- target_rows[df_new$is_canonical[target_rows] == 1]
idx_non_canonical <- target_rows[df_new$is_canonical[target_rows] == 0]

df_new$group[idx_canonical] <- "Canonical"
df_new$group[idx_non_canonical] <- "Non-canonical"

plot_data_new <- df_new %>%
  arrange(cor_pseudotime) %>%
  mutate(rank = 1:n())

highlight_data_new <- plot_data_new %>% filter(group != "Background")

# ==========================================
# ==========================================
p_new <- ggplot(plot_data_new, aes(x = cor_pseudotime, y = rank)) +

  rasterise(geom_point(data = subset(plot_data_new, group == "Background"),
                       color = "grey85", size = 1, alpha = 0.3), dpi = 300) +

  rasterise(geom_point(data = highlight_data_new,
                       aes(color = group),
                       size = 3, alpha = 1), dpi = 300) +

  scale_color_manual(values = c("Canonical" = "#E41A1C",
                                "Non-canonical" = "#004c8c")) +

  geom_text_repel(
    data = highlight_data_new,
    aes(label = gene_name, color = group),
    size = 4,
    fontface = "bold",
    box.padding = 0.6,
    max.overlaps = Inf
  ) +

  geom_vline(xintercept = 0, linetype = "dashed", color = "grey40") +

  labs(
    title = "Sigmoid Correlation Rank Plot (Top 21)",
    x = "Correlation Coefficient",
    y = "Rank"
  ) +

  scale_x_continuous(limits = c(-1, 1), breaks = seq(-1, 1, 0.5)) +
  scale_y_continuous(limits = c(0, 12500)) +

  theme_classic() +
  theme(
    legend.position = "top",
    plot.title = element_text(hjust = 0.5, face = "bold")
  )

print(p_new)

ggsave("sigmoid_plot_canonical_new_rasterized.pdf", plot = p_new, width = 10, height = 5, dpi = 300)

# ==========================================
# ==========================================
gene_range_df <- df_new %>%
  filter(!is.na(cor_pseudotime)) %>%
  group_by(gene_name) %>%
  summarise(
    range_value = max(cor_pseudotime) - min(cor_pseudotime),
    max_cor = max(cor_pseudotime),
    min_cor = min(cor_pseudotime),
    max_transcript_id = transcript_id[which.max(cor_pseudotime)],
    min_transcript_id = transcript_id[which.min(cor_pseudotime)]
  ) %>%
  arrange(desc(range_value))

write.csv(gene_range_df, "gene_cor_range_analysis.csv", row.names = FALSE)
write.xlsx(gene_range_df, "gene_cor_range_analysis.xlsx")

cat("Analysis completed and output saved\n")
```
