## Supplementary material for "An integrated, transcript-resolution atlas of prostate cancer progression unmasks associations with oncogenic pathways": Reproducible code: cononical_pop_up_gene.R

```
# Public submission copy.
# Analytical statements and parameter values are unchanged from the archived source.
# Legacy non-English comments and console messages were normalized for public release.
library(readr)
# ==============================================================================
library(tidyverse)
library(ggplot2)
library(ggrepel)
patients = read.csv("tx_info_with_pseudotime_corr.csv")


patients_pc = patients %>%
  filter(transcript_type == "protein_coding")

library(dplyr)

result_df <- patients_pc %>%
  group_by(gene_id) %>%

  filter({
    vals <- cor_pseudotime

    canon_idx <- which(is_canonical == 1)

    if (length(canon_idx) == 0) {
      FALSE
    } else {
      val_canon <- vals[canon_idx[1]]

      val_max <- max(vals, na.rm = TRUE)
      val_min <- min(vals, na.rm = TRUE)

      is_not_extreme <- (val_canon == val_max) | (val_canon == val_min)

      has_significant_correlation <- (val_max > 0.4) | (val_min < -0.4)

      is_not_extreme & has_significant_correlation
    }
  }) %>%

  ungroup()
length(table(result_df$gene_id))
head(result_df)

write.csv(result_df,"canonical_pop_up_pc_genes_0.4.csv",row.names = F)
ar = result_df %>% filter(gene_name == "AR")


library(clusterProfiler)
library(msigdbr)
library(dplyr)
library(ggplot2)
library(enrichplot)

# =======================================================
# =======================================================

target_genes <- unique(result_df$gene_name)

cat("Processing target gene", length(target_genes), "\n")
head(target_genes)

# =======================================================
# =======================================================

m_t2g <- msigdbr(species = "Homo sapiens", category = "H") %>%
  dplyr::select(gs_name, gene_symbol)

# =======================================================
# =======================================================

hallmark_result <- enricher(gene = target_genes,
                            TERM2GENE = m_t2g,
                            pvalueCutoff = 0.05,
                            qvalueCutoff = 0.2)

# =======================================================
# =======================================================

if (is.null(hallmark_result) || nrow(as.data.frame(hallmark_result)) == 0) {
  message("No significant enrichment was detected")
} else {
  message(paste("Analysis progress", nrow(as.data.frame(hallmark_result)), "Significant pathways detected"))

  print(head(as.data.frame(hallmark_result)))

  write.csv(as.data.frame(hallmark_result), "./canonical_pop_up/filtered_genes_hallmark_enrichment.csv")

  p1 <- dotplot(hallmark_result, showCategory = 15) +
    ggtitle("Hallmark Enrichment (Dotplot)") +
    theme(axis.text.y = element_text(size = 10))

  print(p1)
  ggsave("./canonical_pop_up/Hallmark_enrichment_dotplot.png", p1, width = 8, height = 6)


}
```
