## Supplementary material for "An integrated, transcript-resolution atlas of prostate cancer progression unmasks associations with oncogenic pathways": Reproducible code: correlation_with_pseudotime.R

```
# Public submission copy.
# Analytical statements and parameter values are unchanged from the archived source.
# Legacy non-English comments and console messages were normalized for public release.
library(fst)
library(dplyr)
library(tibble)

# =======================================================
# =======================================================

counts_file <- "./txi.counts.fst"

meta <- read.csv("filtered_samples.csv", stringsAsFactors = FALSE)

transcripts_info <- read.csv("./../MFE/tx_info_full_2.0.csv", stringsAsFactors = FALSE)

pseudotime_vec <- meta$pseudotime

txi <- list(counts = as.matrix(read.fst(counts_file)))
ncount.tx <- log2(sweep(x = txi$counts, MARGIN = 2, STATS = colSums(txi$counts)/1000000, FUN = "/") + 1)


print("Running analysis step")

r_values <- cor(t(ncount.tx), pseudotime_vec, method = "pearson")

n_samples <- ncol(ncount.tx)
df <- n_samples - 2
t_stats <- r_values * sqrt(df / (1 - r_values^2))
p_values <- 2 * pt(-abs(t_stats), df)

# =======================================================
# =======================================================

stat_res <- data.frame(
  cor_pseudotime = as.numeric(r_values),
  pval_pseudotime = as.numeric(p_values)
)

stat_res$padj_pseudotime <- p.adjust(stat_res$pval_pseudotime, method = "BH")

transcripts_info_new =cbind(transcripts_info,stat_res)

write.csv(transcripts_info_new, "tx_info_with_pseudotime_corr.csv", row.names = FALSE)


# lnc = transcripts_info_new %>%
#   filter(transcript_type == "lncRNA") %>%
#   dplyr::select(c("transcript_id","gene_id","transcript_type","gene_name","cor_pseudotime"))
lnc = transcripts_info_new %>%
  filter(transcript_type == "lncRNA") %>%
  filter(!is.na(cor_pseudotime))

write.csv(lnc, "lncRNA_with_pseudotime_corr.csv", row.names = FALSE)


pc = transcripts_info_new %>%
  filter(transcript_type == "protein_coding")


library(dplyr)
library(ggplot2)
library(ggpubr)
library(patchwork)

pc <- transcripts_info_new %>%
  filter(transcript_type == "protein_coding") %>%
  filter(!is.na(cor_pseudotime))
plot_data <- pc %>%
  mutate(
    Canonical_Status = ifelse(is_canonical == 1, "Canonical", "Non-Canonical"),
    Canonical_Status = factor(Canonical_Status, levels = c("Non-Canonical", "Canonical")),
    abs_cor = abs(cor_pseudotime)
  )


p1 <- ggplot(plot_data, aes(x = cor_pseudotime, fill = Canonical_Status)) +
  geom_density(alpha = 0.5) +
  scale_fill_manual(values = c("gray70", "#E41A1C")) +
  labs(title = "A. Distribution of Correlation",
       x = "Correlation with Pseudotime",
       y = "Density") +
  theme_minimal() +
  theme(legend.position = "top")

p2 <- ggplot(plot_data, aes(x = Canonical_Status, y = abs_cor, fill = Canonical_Status)) +
  geom_violin(alpha = 0.4, color = NA) +
  geom_boxplot(width = 0.2, outlier.shape = NA) +
  stat_compare_means(method = "wilcox.test", label.y = 1.05, label.x = 1.3) +
  scale_fill_manual(values = c("gray70", "#E41A1C")) +
  labs(title = "B. Magnitude of Correlation (|r|)",
       subtitle = "Are canonical transcripts more strongly correlated?",
       y = "Absolute Correlation (|r|)",
       x = NULL) +
  theme_minimal() +
  theme(legend.position = "none")

final_plot <- p1 + p2
print(final_plot)

ggsave("canonical_vs_pseudotime_correlation.pdf", final_plot, width = 10, height = 5)


test_res <- wilcox.test(abs_cor ~ Canonical_Status, data = plot_data, alternative = "less")

print(paste("P-value (Canonical is stronger):", test_res$p.value))

summary_stats <- plot_data %>%
  group_by(Canonical_Status) %>%
  summarise(
    count = n(),
    mean_abs_cor = mean(abs_cor),
    median_abs_cor = median(abs_cor),
    sd_cor = sd(cor_pseudotime)
  )

print(summary_stats)
```
