## Supplementary material for "An integrated, transcript-resolution atlas of prostate cancer progression unmasks associations with oncogenic pathways": Reproducible code: isoform_pair_DEG_GSEA_full.R

```
# Public submission copy.
# Analytical statements and parameter values are unchanged from the archived source.
# Legacy non-English comments and console messages were normalized for public release.
# =====================================================================
# =====================================================================
neededlibraries <- c("edgeR", "org.Hs.eg.db", "ape", "ggplot2", "reshape", "RColorBrewer",
                     "plotly", "pheatmap", "DESeq2", "AnnotationDbi", "ggrepel", "qusage",
                     "EGSEA", "GeneOverlap", "reshape2", "clipr", "svglite", "readtext",
                     "ggpubr", "clusterProfiler", "fst", "dplyr", "msigdbr", "DOSE", "stringr", "forcats")
lapply(neededlibraries, require, character.only = TRUE)

# =====================================================================
# =====================================================================
cat("Loading input data\n")

txi_df <- read.fst("./txi.counts.fst")
txi_info <- read.csv("./transcripts_info.csv")
rownames(txi_df) <- txi_info$transcript_id
txi_counts <- as.matrix(txi_df)

ncount.tx <- log2(sweep(x = txi_counts, MARGIN = 2, STATS = colSums(txi_counts)/1000000, FUN = "/") + 1)
base_tx_ids <- gsub("\\..*$", "", rownames(ncount.tx))

gc_df <- read.fst("./gc_txi.counts.fst")
gc_info <- read.csv("./genes_info.csv")
rownames(gc_df) <- gc_info$gene_id
gc_counts <- round(as.matrix(gc_df))
gc_counts <- gc_counts[rowSums(gc_counts) > 0, ]
gc_info$base_gene_id <- gsub("\\..*$", "", gc_info$gene_id)

m_t2g <- msigdbr(species = "Homo sapiens", category = "H") %>%
  dplyr::select(gs_name, entrez_gene)

cat("Loading input data\n")

# =====================================================================
# =====================================================================
run_deg_gsea_pipeline <- function(gene_name, enst_A, enst_B, pct = 0.1) {

  prefix <- paste0(enst_A, "_vs_", enst_B, "_Top", pct*100)
  cat(paste0("\n------------------------------------------------------\n"))
  cat(paste0("Running analysis step", gene_name, "]: ", prefix, "\n"))

  if (!dir.exists(gene_name)) {
    dir.create(gene_name)
  }

  match_idx_A <- which(base_tx_ids == enst_A)
  match_idx_B <- which(base_tx_ids == enst_B)

  if (length(match_idx_A) == 0 | length(match_idx_B) == 0) {
    cat(paste0("Analysis failed", enst_A, "Analysis progress", enst_B, "Analysis warning; step skipped\n"))
    return(NULL)
  }

  expr_A <- ncount.tx[match_idx_A[1], ]
  expr_B <- ncount.tx[match_idx_B[1], ]

  cutoff_high_A <- quantile(expr_A, 1 - pct); cutoff_low_A  <- quantile(expr_A, pct)
  cutoff_high_B <- quantile(expr_B, 1 - pct); cutoff_low_B  <- quantile(expr_B, pct)

  sample_annot <- data.frame(ID = colnames(ncount.tx), Expr_A = expr_A, Expr_B = expr_B, GROUP = NA, stringsAsFactors = FALSE)

  sample_annot$GROUP[sample_annot$Expr_A >= cutoff_high_A & sample_annot$Expr_B <= cutoff_low_B] <- "A_High_B_Low"
  sample_annot$GROUP[sample_annot$Expr_A <= cutoff_low_A & sample_annot$Expr_B >= cutoff_high_B] <- "A_Low_B_High"

  sample_annot <- sample_annot[!is.na(sample_annot$GROUP), ]

  count_A_high <- sum(sample_annot$GROUP == "A_High_B_Low")
  count_B_high <- sum(sample_annot$GROUP == "A_Low_B_High")
  cat(paste0("Assigning analysis groups", count_A_high, "Analysis progress", count_B_high, "Analysis progress\n"))

  if(count_A_high < 2 | count_B_high < 2) {
    cat(paste0("Analysis failed\n"))
    return(NULL)
  }

  sample_annot$GROUP <- factor(sample_annot$GROUP, levels = c("A_Low_B_High", "A_High_B_Low"))

  gc_counts_filtered <- gc_counts[, sample_annot$ID]
  gc_counts_filtered <- gc_counts_filtered[rowSums(gc_counts_filtered) > 0, ]

  dds.design <- model.matrix(~ 0 + GROUP, data = sample_annot)
  dds <- DESeqDataSetFromMatrix(countData = gc_counts_filtered, colData = sample_annot, design = dds.design)

  res_de <- tryCatch({
    dds <- DESeq(dds, betaPrior = FALSE, quiet = TRUE)
    results(dds, contrast = makeContrasts(GROUPA_High_B_Low - GROUPA_Low_B_High, levels = dds.design), independentFiltering = TRUE, alpha = 0.05)
  }, error = function(e) { return(NULL) })

  if(is.null(res_de)) {
    cat("Analysis failed\n")
    return(NULL)
  }

  res_df <- data.frame(res_de, stringsAsFactors = FALSE)
  res_df$ENSEMBL <- rownames(res_df)
  base_ensembl_res <- gsub("\\..*$", "", res_df$ENSEMBL)
  res_df$SYMBOL <- gc_info$gene_name[match(base_ensembl_res, gc_info$base_gene_id)]
  res_df$ENTREZ <- mapIds(org.Hs.eg.db, keys = res_df$SYMBOL, column = "ENTREZID", keytype = "SYMBOL", multiVals = "first")

  res_df <- res_df[!is.na(res_df$SYMBOL) & !duplicated(res_df$SYMBOL), ]
  res_df <- res_df[!is.na(res_df$ENTREZ) & !duplicated(res_df$ENTREZ), ]
  res_df <- res_df[!is.na(res_df$padj), ]

  write.csv(res_df, file.path(gene_name, paste0("DEGs_", prefix, ".csv")), row.names = FALSE)

  geneList <- res_df$stat
  names(geneList) <- as.character(res_df$ENTREZ)
  geneList <- sort(geneList, decreasing = TRUE)

  GSEA_res <- tryCatch({
    GSEA(geneList, TERM2GENE = m_t2g, pvalueCutoff = 0.05, eps = 1e-50, verbose = FALSE, seed = TRUE)
  }, error = function(e) { return(NULL) })

  if(!is.null(GSEA_res) && nrow(GSEA_res@result) > 0) {
    cat("Analysis completed and output saved\n")

    # Dotplot
    p_dotplot <- dotplot(GSEA_res, showCategory = 20, orderBy = "p.adjust", x = "enrichmentScore") +
      ggtitle(paste0(gene_name, " GSEA: ", prefix))
    ggsave(file.path(gene_name, paste0('GSEA_Dotplot_', prefix, '.pdf')), p_dotplot, width = 10, height = 8)

    max_nes <- max(abs(GSEA_res@result$NES), na.rm = TRUE)
    color_limit <- c(-max_nes, max_nes)

    dot_df <- GSEA_res@result %>% mutate(SIG = -log10(p.adjust))
    p_custom <- ggplot(dot_df, aes(x = NES, y = fct_reorder(Description, SIG))) +
      geom_point(aes(size = SIG, color = NES)) +
      theme_bw(base_size = 14) +
      scale_colour_gradient2(limits = color_limit, low = "#3953A4", mid = "white", high = "#b50404") +
      labs(title = paste0(gene_name, " GSEA NES"), y = NULL)
    ggsave(file.path(gene_name, paste0('GSEA_NES_', prefix, '.pdf')), p_custom, width = 14, height = 10)

  } else {
    cat("Significant pathways detected\n")
  }
}

# =====================================================================
# =====================================================================
analysis_tasks <- list(
  c("WNT2B",  "ENST00000369684", "ENST00000369686"),
  c("TRIM11", "ENST00000284551", "ENST00000493030"),
  c("ROCK2",  "ENST00000315872", "ENST00000401753"),
  c("RNF43",  "ENST00000584437", "ENST00000583753"),
  c("CDK12",  "ENST00000447079", "ENST00000584632"),
  c("F8",     "ENST00000360256", "ENST00000330287"),
  c("RELA",   "ENST00000406246", "ENST00000612991"),
  c("BRCA1",  "ENST00000357654", "ENST00000352993")
)
analysis_tasks <- list(

  c("CDK12",  "ENST00000447079", "ENST00000584632")
)
target_pct <- 0.1
target_pct <- 0.2
for (task in analysis_tasks) {
  run_deg_gsea_pipeline(gene_name = task[1],
                        enst_A = task[2],
                        enst_B = task[3],
                        pct = target_pct)
}

cat("\n======================================================\n")
cat("Analysis progress\n")
```
