## Supplementary material for "An integrated, transcript-resolution atlas of prostate cancer progression unmasks associations with oncogenic pathways": Reproducible code: kendall_redraw_rasterized.R

```
# Public submission copy.
# Analytical statements and parameter values are unchanged from the archived source.
# Legacy non-English comments and console messages were normalized for public release.
library(readr)
library(ggplot2)
library(ggrastr)

custom_theme <- theme_classic() +
  theme(
    plot.title = element_text(size = 16, face = "bold"),
    axis.title = element_text(size = 14),
    axis.text = element_text(size = 12),
    legend.title = element_text(size = 12),
    legend.text = element_text(size = 10),
    panel.grid = element_blank() # Remove grid lines
    #panel.border = element_rect(colour = "black", fill = NA, linewidth = 1) # Keep border
  )

results = read.csv("kendall_tau_results_pc_isoform.csv")
ggplot(data = results, aes(x = kendall_tau_estimate, y = kendall_tau_statistic, color = factor(is_canonical))) +
  rasterise(geom_point(alpha = 0.6, size = 2), dpi = 300) +
  scale_color_manual(values = c("0" = "#808080", "1" = "#FF0000"),
                     labels = c("0" = "Non-Canonical", "1" = "Canonical")) +
  labs(x = "Kendall Tau Estimate (tau)",
       y = "Kendall Tau Statistic",
       title = "Kendall Tau Analysis: Estimate vs. Statistic",
       color = "Canonical Status") +
  custom_theme +
  theme(
    plot.title = element_text(hjust = 0.5),
    axis.title = element_text(size = 12),
    axis.text = element_text(size = 10)
  )
ggsave("kendall_tau_results_pc_isoform_scatter_plot_redraw.pdf", width = 8, height = 6, dpi = 300)


library(dplyr)
v = results %>%
  arrange(desc(kendall_tau_statistic)) %>%
  select(is_canonical) %>%
  pull(is_canonical)
red_positions <- which(v == 1) / length(v)


total_counts <- length(red_positions)
num_bins <- 100
expected_count <- total_counts / num_bins

ggplot(data.frame(position = red_positions), aes(x = position)) +
  geom_density(fill = "#FF0000", alpha = 0.4, bw =0.01) +
  stat_function(fun = dunif, args = list(min = 0, max = 1), color = "black", linetype = "dashed") +
  labs(x = "Relative Position (0 = Highest Statistic, 1 = Lowest Statistic)",
       y = "Density",
       title = "Density of Canonical Transcripts' Positions vs. Uniform Distribution") +
  custom_theme +
  theme( plot.title = element_text(hjust = 0.5),
         axis.title = element_text(size = 12),
         axis.text = element_text(size = 10) )
ggsave("red_positions_density_redraw.pdf", width = 8, height = 6, dpi = 300)


results = read.csv("kendall_tau_results_lnc_gene.csv")
library(ggrepel)
library(readxl)

lnc_list <-read_excel("lncRNA_list_new.xlsx",
                      sheet = "Sheet2")

ggplot(data = results, aes(x = kendall_tau_estimate, y = kendall_tau_statistic)) +
  rasterise(geom_point(alpha = 0.6, size = 2), dpi = 300) +
  rasterise(geom_point(data = subset(results, gene_name %in% lnc_list$`Search Name`),
                       aes(x = kendall_tau_estimate, y = kendall_tau_statistic,
                           color = lnc_list$`Major impacts`[match(gene_name, lnc_list$`Search Name`)]),
                       size = 2), dpi = 300) +
  geom_text_repel(data = subset(results, gene_name %in% lnc_list$`Search Name`),
                  aes(label = lnc_list$LncRNA[match(gene_name, lnc_list$`Search Name`)],
                      color = lnc_list$`Major impacts`[match(gene_name, lnc_list$`Search Name`)]),
                  size = 3.5,
                  max.overlaps = Inf,
                  min.segment.length = 0,
                  box.padding = 1.0,
                  point.padding = 0.5,
                  force = 50,
                  segment.color = "grey50",
                  segment.size = 0.5,
                  segment.alpha = 0.8) +
  scale_color_manual(values = c("UP" = "red", "DOWN" = "blue"),
                     labels = c("UP" = "Positive", "DOWN" = "Negative"),
                     name = "Association with \nCancer Progression") +
  labs(x = "Kendall Tau Estimate (tau)",
       y = "Kendall Tau Statistic",
       title = "Kendall Tau Analysis: Estimate vs. Statistic") +
  custom_theme +
  theme(
    plot.title = element_text(hjust = 0.5),
    axis.title = element_text(size = 12),
    axis.text = element_text(size = 10)
  )
ggsave("kendall_tau_results_lnc_gene_scatter_plot_sel_redrawrasterized.pdf", width = 8, height = 6, dpi = 300)
```
