## Supplementary material for "An integrated, transcript-resolution atlas of prostate cancer progression unmasks associations with oncogenic pathways": Reproducible code: pc_isoform_kt_score.R

```
# Public submission copy.
# Analytical statements and parameter values are unchanged from the archived source.
# Legacy non-English comments and console messages were normalized for public release.
library(readr)
library(fst)

# Load data
mdata = read.csv("filtered_samples.csv")
txi = list(counts = as.matrix(read.fst("./txi.counts.fst")))
ids = read.csv("PCa.tx.corr_annot.csv", sep=",", header=T)

# Normalize and log-transform expression matrix
expr_matrix <- log2(sweep(x=txi$counts, MARGIN=2, STATS=colSums(txi$counts)/1000000, FUN="/") + 1)

# Filter for protein-coding transcripts

pair = mdata$Library.layout == "PAIRED"
expr_matrix <- expr_matrix[,pair]
mdata = mdata[pair,]


# Filter for protein-coding transcripts
pc = ids$transcript_type == "protein_coding"
expr_matrix <- expr_matrix[pc,]
ids = ids[pc,]

dim(expr_matrix)
mdata$Molecular.Type <- factor(mdata$Molecular.Type,
                               levels = c("NORMAL", "PRIMARY", "ARPC", "DNPC", "NEPC"),
                               ordered = TRUE)

n_transcripts <- nrow(expr_matrix)
results <- data.frame(
  transcript_id = ids$transcript_id,
  kendall_tau_statistic = numeric(n_transcripts),
  kendall_tau_estimate = numeric(n_transcripts),
  p_value = numeric(n_transcripts)
)

for (i in 1:n_transcripts) {
  kendall_result <- cor.test(expr_matrix[i, ],
                             as.numeric(mdata$Molecular.Type),
                             method = "kendall",
                             exact = FALSE)
  results$kendall_tau_statistic[i] <- kendall_result$statistic
  results$kendall_tau_estimate[i] <- kendall_result$estimate
  results$p_value[i] <- kendall_result$p.value
}

results$ensembl = ids$ensembl
results$is_canonical = ids$is_canonical
results$gene_name = ids$gene_name
write.csv(results, "kendall_tau_results_pc_isoform.csv", row.names = FALSE)


library(ggplot2)


ggplot(data = results, aes(x = kendall_tau_estimate, y = kendall_tau_statistic, color = factor(is_canonical))) +
  geom_point(alpha = 0.6, size = 2) +
  scale_color_manual(values = c("0" = "#808080", "1" = "#FF0000"),
                     labels = c("0" = "Non-Canonical", "1" = "Canonical")) +
  labs(x = "Kendall Tau Estimate (tau)",
       y = "Kendall Tau Statistic",
       title = "Kendall Tau Analysis: Estimate vs. Statistic",
       color = "Canonical Status") +
  theme_minimal() +
  theme(
    plot.title = element_text(hjust = 0.5),
    axis.title = element_text(size = 12),
    axis.text = element_text(size = 10)
  )

ggsave("kendall_tau_results_pc_isoform_scatter_plot.pdf", width = 8, height = 6, dpi = 300)


library(dplyr)
v = results %>%
  arrange(desc(kendall_tau_statistic)) %>%
  select(is_canonical) %>%
  pull(is_canonical)
red_positions <- which(v == 1) / length(v)


total_counts <- length(red_positions)
num_bins <- 100
expected_count <- total_counts / num_bins
# length(v)/length(red_positions)* 100  =
ggplot(data.frame(position = red_positions), aes(x = position)) +
  geom_histogram(bins = num_bins, fill = "#FF0000", color = "black", alpha = 0.6) +
  geom_hline(yintercept = expected_count, color = "red", linetype = "solid", size = 1) +
  labs(x = "Relative Position (0 = Highest Statistic, 1 = Lowest Statistic)",
       y = "Count",
       title = "Distribution of Canonical Transcripts' Positions") +
  theme_minimal() +
  theme(
    plot.title = element_text(hjust = 0.5),
    axis.title = element_text(size = 12),
    axis.text = element_text(size = 10)
  )

ggsave("red_positions_histogram.png", width = 8, height = 6, dpi = 300)

ggplot(data.frame(position = red_positions), aes(x = position)) +
  geom_density(fill = "#FF0000", alpha = 0.4, bw =0.01) +
  stat_function(fun = dunif, args = list(min = 0, max = 1), color = "black", linetype = "dashed") +
  #geom_rug(sides = "b", color = "black", size = 0.5, alpha = 0.6) +
  labs(x = "Relative Position (0 = Highest Statistic, 1 = Lowest Statistic)",
       y = "Density",
       title = "Density of Canonical Transcripts' Positions vs. Uniform Distribution") +
  theme_minimal() +
  theme( plot.title = element_text(hjust = 0.5),
         axis.title = element_text(size = 12),
         axis.text = element_text(size = 10) )
ggsave("red_positions_density.png", width = 8, height = 6, dpi = 300)

ks_test <- ks.test(red_positions, "punif", min = 0, max = 1)
print("Kolmogorov-Smirnov Test for Uniform Distribution:")
print(ks_test)

tail_proportion <- 0.1
n_total <- length(v)
n_red <- sum(v == 1)
head_tail_count <- sum(red_positions <= tail_proportion | red_positions >= (1 - tail_proportion))
head_tail_proportion <- head_tail_count / n_red
expected_proportion <- 2 * tail_proportion

cat("Proportion of canonical transcripts in head/tail 10%:", head_tail_proportion, "\n")
cat("Expected proportion under uniform distribution:", expected_proportion, "\n")

binom_test <- binom.test(head_tail_count, n_red, p = expected_proportion, alternative = "greater")
print("Binomial Test for Head/Tail Enrichment:")
print(binom_test)
```
