## Supplementary material for "An integrated, transcript-resolution atlas of prostate cancer progression unmasks associations with oncogenic pathways": Reproducible code: AWSI_forest_plot.R

```
# Public submission copy.
# Analytical statements and parameter values are unchanged from the archived source.
# Legacy non-English comments and console messages were normalized for public release.
# =====================================================================
# =====================================================================
library(fst)
library(dplyr)
library(ggplot2)
library(stringr)
library(tibble)
library(forcats)

cat("Loading input data\n")

txi_df <- read.fst("./txi.counts.fst")
txi_info <- read.csv("./transcripts_info.csv")
rownames(txi_df) <- txi_info$transcript_id
txi_counts <- as.matrix(txi_df)

ncount.tx <- log2(sweep(x = txi_counts, MARGIN = 2, STATS = colSums(txi_counts)/1000000, FUN = "/") + 1)

base_tx_ids <- gsub("\\..*$", "", rownames(ncount.tx))
rownames(ncount.tx) <- base_tx_ids

load("Full_GSVA_Analysis_Workspace_2026-03-23.Rdata")

cat("Analysis completed and output saved\n")

# =====================================================================
# =====================================================================

run_isoform_pathway_cor_with_ci <- function(gene_name, enst_A, enst_B, expr_matrix, ssgsea_mat, fdr_cutoff = 0.05, cor_cutoff = 0.3) {

  cat(paste0("\n------------------------------------------------------\n"))
  cat(paste0("Running analysis step", gene_name, "Analysis progress\n"))

  if(!enst_A %in% rownames(expr_matrix) | !enst_B %in% rownames(expr_matrix)) {
    cat(paste0("Analysis failed\n"))
    return(NULL)
  }

  expr_A <- expr_matrix[enst_A, ]
  expr_B <- expr_matrix[enst_B, ]

  cpm_A <- (2^expr_A) - 1
  cpm_B <- (2^expr_B) - 1
  total_cpm <- cpm_A + cpm_B

  AWSI <- ((cpm_A / (total_cpm + 1e-5)) - 0.5) * log2(total_cpm + 1)
  metric_df <- data.frame(Sample = names(AWSI), AWSI = AWSI, stringsAsFactors = FALSE)

  common_samples <- intersect(colnames(ssgsea_mat), metric_df$Sample)
  if(length(common_samples) < 10) return(NULL)

  mat_sub <- ssgsea_mat[, common_samples]
  vec_awsi <- metric_df$AWSI[match(common_samples, metric_df$Sample)]
  n_samples <- length(vec_awsi)

  cor_results <- lapply(rownames(mat_sub), function(pathway) {
    pathway_scores <- mat_sub[pathway, ]
    test_res <- cor.test(vec_awsi, pathway_scores, method = "spearman", exact = FALSE)
    r <- test_res$estimate

    r_safe <- pmin(pmax(r, -0.999), 0.999)
    z <- atanh(r_safe)
    se <- 1 / sqrt(n_samples - 3)
    ci_lower <- tanh(z - 1.96 * se)
    ci_upper <- tanh(z + 1.96 * se)

    data.frame(
      Pathway = pathway,
      Correlation = r,
      P_value = test_res$p.value,
      CI_lower = ci_lower,
      CI_upper = ci_upper
    )
  })

  res_df <- bind_rows(cor_results)
  res_df$FDR <- p.adjust(res_df$P_value, method = "fdr")
  res_df <- res_df %>% arrange(FDR, desc(abs(Correlation)))

  if (!dir.exists(gene_name)) dir.create(gene_name)
  write.csv(res_df, file.path(gene_name, paste0(gene_name, "_Continuous_AWSI_with_CI.csv")), row.names = FALSE)

  return(res_df)
}

plot_significant_hallmark_forest <- function(res_df, gene_name, enst_A, enst_B) {

  hallmark_sig_df <- res_df %>%
    filter(grepl("^HALLMARK_", Pathway)) %>%
    filter(FDR <= 0.05) %>%
    mutate(Short_Path = gsub("HALLMARK_", "", Pathway)) %>%
    mutate(Short_Path = str_trunc(Short_Path, 40))

  if(nrow(hallmark_sig_df) == 0) {
    cat(paste0("    ⚠️ ", gene_name, "Significant pathways detected\n"))
    return(NULL)
  }

  hallmark_sig_df <- hallmark_sig_df %>%
    mutate(Short_Path = fct_reorder(Short_Path, Correlation))

  hallmark_sig_df$SIG <- -log10(hallmark_sig_df$FDR + 1e-300)

  max_val <- max(abs(c(hallmark_sig_df$CI_lower, hallmark_sig_df$CI_upper)), na.rm = TRUE)

  p_forest <- ggplot(hallmark_sig_df, aes(x = Correlation, y = Short_Path)) +
    geom_vline(xintercept = 0, linetype = "dashed", color = "grey60", linewidth = 0.8) +
    geom_errorbarh(aes(xmin = CI_lower, xmax = CI_upper), height = 0.25, color = "grey50", linewidth = 0.8) +
    geom_point(aes(size = SIG, color = Correlation)) +
    scale_color_gradient2(limits = c(-max_val, max_val),
                          low = "#3953A4", mid = "white", high = "#b50404", midpoint = 0) +
    theme_bw(base_size = 13) +
    labs(
      title = paste0(gene_name, " Significant Hallmark Pathways (FDR < 0.05)"),
      subtitle = paste0("Spearman Rho & 95% CI | AWSI (", enst_A, " vs ", enst_B, ")"),
      x = "Spearman Correlation (Rho) with 95% CI",
      y = "Pathways",
      size = "-log10(FDR)",
      color = "Rho"
    ) +
    theme(
      axis.text.y = element_text(size = 11, face = "bold"),
      panel.grid.major.y = element_blank(),
      panel.grid.minor = element_blank(),
      legend.position = "right"
    )

  save_name <- file.path(gene_name, paste0(gene_name, "_Sig_Hallmark_Forest_Plot.pdf"))
  plot_height <- max(5, nrow(hallmark_sig_df) * 0.25)
  ggsave(save_name, p_forest, width = 11, height = plot_height)

  cat(paste0("Analysis completed and output saved", save_name, "\n"))
  return(p_forest)
}

# =====================================================================
# =====================================================================

analysis_tasks <- list(
  # c("RELA",   "ENST00000406246", "ENST00000612991"),
  # c("RNF43",  "ENST00000584437", "ENST00000583753"),
  # c("ROCK2",  "ENST00000315872", "ENST00000401753"),
  # c("TRIM11", "ENST00000284551", "ENST00000493030")
  c("BRCA1","ENST00000357654","ENST00000352993"),
  c("WNT2B","ENST00000369684","ENST00000369686")
  # c("GENE_NAME", "ENST_XXXX", "ENST_YYYY")
)

for (task in analysis_tasks) {
  gene  <- task[1]
  iso_A <- task[2]
  iso_B <- task[3]

  res_df_ci <- run_isoform_pathway_cor_with_ci(
    gene_name = gene,
    enst_A = iso_A,
    enst_B = iso_B,
    expr_matrix = ncount.tx,
    ssgsea_mat = full_gsva_res
  )

  if (!is.null(res_df_ci)) {
    plot_significant_hallmark_forest(res_df_ci, gene, iso_A, iso_B)
  }
}

cat("Analysis progress\n")
```
