## Supplementary material for "An integrated, transcript-resolution atlas of prostate cancer progression unmasks associations with oncogenic pathways": Reproducible code: isoform_single_comparison_GOCC_GSEA.R

```
# Public submission copy.
# Analytical statements and parameter values are unchanged from the archived source.
# Legacy non-English comments and console messages were normalized for public release.
# =====================================================================
# =====================================================================

library(dplyr)
library(ggplot2)
library(clusterProfiler)
library(msigdbr)
library(stringr)
library(forcats)

# =====================================================================
# =====================================================================
cat("Running analysis step\n")
m_t2g_cc <- msigdbr(species = "Homo sapiens", category = "C5", subcategory = "GO:CC") %>%
  dplyr::select(gs_name, entrez_gene) %>%
  mutate(gs_name = str_replace(gs_name, "^GOCC_", "")) %>%
  mutate(gs_name = str_replace_all(gs_name, "_", " "))

# =====================================================================
# =====================================================================
files_list <- list(
  "BRCA1_T" = "./DEGs_ENST00000352993_Top25_vs_Low25.csv",
  "BRCA1_C" = "./DEGs_ENST00000357654_Top25_vs_Low25.csv"
)

# =====================================================================
# =====================================================================
gsea_results_list <- list()

for (comp_name in names(files_list)) {
  csv_file <- files_list[[comp_name]]

  cat(paste0("Running analysis step", comp_name, "...\n"))

  if (!file.exists(csv_file)) {
    warning(paste("Required input was not found", csv_file))
    next
  }

  res_df <- read.csv(csv_file)

  valid_df <- res_df %>% filter(!is.na(stat) & !is.na(ENTREZ))

  geneList <- sort(setNames(valid_df$stat, as.character(valid_df$ENTREZ)), decreasing = TRUE)

  gsea_result <- tryCatch({
    GSEA(geneList, TERM2GENE = m_t2g_cc, pvalueCutoff = 0.05, seed = 123, verbose = FALSE)
  }, error = function(e) {
    cat(paste0("Analysis progress", comp_name, "Analysis progress", e$message, "\n"))
    return(NULL)
  })

  if(!is.null(gsea_result) && nrow(gsea_result@result) > 0) {
    gene_count <- gsea_result@result %>%
      group_by(ID) %>%
      summarise(count = sum(str_count(core_enrichment, "/")) + 1)

    dot_df <- left_join(gsea_result@result, gene_count, by = "ID") %>%
      mutate(GeneRatio = count/setSize)

    gsea_results_list[[comp_name]] <- dot_df
    cat(paste0("  -> ", comp_name, "Analysis progress", nrow(dot_df), "Significant pathways detected\n"))
  } else {
    cat(paste0("  -> ", comp_name, "No significant enrichment was detected\n"))
  }
}

# =====================================================================
# =====================================================================
if(length(gsea_results_list) > 0) {
  cat("Running analysis step\n")

  merged.res <- as.data.frame(merge_result(gsea_results_list))

  merged.res$SIG = -log10(merged.res$p.adjust)

  top_terms <- merged.res %>%
    group_by(Cluster) %>%
    slice_max(order_by = abs(NES), n = 15) %>%
    pull(Description) %>%
    unique()

  plot_data <- merged.res %>% filter(Description %in% top_terms)

  p <- ggplot(plot_data, aes(x = NES, y = fct_reorder(Description, SIG))) +
    geom_point(aes(size = SIG, color = NES, shape = Cluster), stroke = 1.2) +
    theme_bw(base_size = 14) +
    scale_colour_gradient2(limits=c(min(plot_data$NES), max(plot_data$NES)),
                           low = "#3953A4", mid = "white", high = "#b50404") +
    scale_shape_manual(values = c("BRCA1_T" = 16, "BRCA1_C" = 1)) +
    scale_size(range = c(3, 10)) +
    ylab(NULL) +
    labs(title = "Subcellular Spatial Signatures (GO-CC)",
         subtitle = "BRCA1_T vs BRCA1_C Comparison",
         size = "-log10(padj)",
         color = "NES",
         shape = "Group") +
    theme(
      legend.position = "right",
      panel.grid.major.y = element_line(linetype = "dashed", color = "gray80"),
      axis.text.y = element_text(size = 11, color = "black")
    )

  print(p)
  ggsave("./GSEA_Spatial_GO_CC_Combined_Style.pdf", p, width = 12, height = 9)
  ggsave("./GSEA_Spatial_GO_CC_Combined_Style.png", p, width = 12, height = 9, dpi = 300)

  write.csv(merged.res, "./GSEA_Spatial_GO_CC_merged_results.csv", row.names = FALSE)

  cat("Analysis completed and output saved\n")
} else {
  cat("No significant enrichment was detected\n")
}
```
