## Supplementary material for "An integrated, transcript-resolution atlas of prostate cancer progression unmasks associations with oncogenic pathways": Reproducible code: isoform_single_comparison_GSEA.R

```
# Public submission copy.
# Analytical statements and parameter values are unchanged from the archived source.
# Legacy non-English comments and console messages were normalized for public release.
# =====================================================================
# =====================================================================
neededlibraries <- c("edgeR", "org.Hs.eg.db", "ggplot2", "DESeq2", "AnnotationDbi",
                     "clusterProfiler", "fst", "dplyr", "msigdbr", "stringr", "forcats")
lapply(neededlibraries, require, character.only = TRUE)

# =====================================================================
# =====================================================================
cat("Loading input data\n")
txi_df <- read.fst("./txi.counts.fst")
txi_info <- read.csv("./transcripts_info.csv")
rownames(txi_df) <- txi_info$transcript_id
txi_counts <- as.matrix(txi_df)

ncount.tx <- log2(sweep(x = txi_counts, MARGIN = 2, STATS = colSums(txi_counts)/1000000, FUN = "/") + 1)

gc_df <- read.fst("./gc_txi.counts.fst")
gc_info <- read.csv("./genes_info.csv")
rownames(gc_df) <- gc_info$gene_id
gc_counts <- round(as.matrix(gc_df))
gc_counts <- gc_counts[rowSums(gc_counts) > 0, ]
gc_info$base_gene_id <- gsub("\\..*$", "", gc_info$gene_id)

m_t2g <- msigdbr(species = "Homo sapiens", category = "H") %>%
  dplyr::select(gs_name, entrez_gene)

# =====================================================================
# =====================================================================
run_single_iso_gsea <- function(target_enst, top_pct = 0.25, bottom_pct = 0.25) {

  prefix <- paste0(gsub("\\..*$", "", target_enst), "_Top", top_pct*100, "_vs_Low", bottom_pct*100)
  cat(paste0("Running analysis step", target_enst, " <<<\n"))

  if (target_enst %in% rownames(ncount.tx)) {
    expr_vec <- ncount.tx[target_enst, ]
  } else {
    base_ids <- gsub("\\..*$", "", rownames(ncount.tx))
    idx <- which(base_ids == gsub("\\..*$", "", target_enst))
    if (length(idx) == 0) stop("Required input was not found")
    expr_vec <- ncount.tx[idx[1], ]
  }

  cutoff_high <- quantile(expr_vec, 1 - top_pct)
  cutoff_low  <- quantile(expr_vec, bottom_pct)

  sample_annot <- data.frame(
    ID = colnames(ncount.tx),
    Expr = expr_vec,
    GROUP = NA,
    stringsAsFactors = FALSE
  )

  sample_annot$GROUP[sample_annot$Expr >= cutoff_high] <- "High"
  sample_annot$GROUP[sample_annot$Expr <= cutoff_low] <- "Low"

  sample_annot <- sample_annot[!is.na(sample_annot$GROUP), ]
  sample_annot$GROUP <- factor(sample_annot$GROUP, levels = c("Low", "High"))

  cat(paste0("Assigning analysis groups", sum(sample_annot$GROUP == "High"),
             "Analysis progress", sum(sample_annot$GROUP == "Low"), "Analysis progress\n"))

  selected_samples <- sample_annot$ID
  dds <- DESeqDataSetFromMatrix(countData = gc_counts[, selected_samples],
                                colData = sample_annot,
                                design = ~ GROUP)

  dds <- DESeq(dds, quiet = TRUE)
  res_de <- results(dds, contrast = c("GROUP", "High", "Low"))

  res_df <- as.data.frame(res_de)
  res_df$SYMBOL <- gc_info$gene_name[match(gsub("\\..*$", "", rownames(res_df)), gc_info$base_gene_id)]
  res_df$ENTREZ <- mapIds(org.Hs.eg.db, keys = res_df$SYMBOL, column = "ENTREZID", keytype = "SYMBOL", multiVals = "first")

  res_df <- res_df %>% filter(!is.na(SYMBOL), !is.na(ENTREZ), !is.na(padj)) %>%
    distinct(SYMBOL, .keep_all = TRUE) %>%
    distinct(ENTREZ, .keep_all = TRUE)

  write.csv(res_df, paste0("DEGs_", prefix, ".csv"))

  geneList <- sort(setNames(res_df$stat, res_df$ENTREZ), decreasing = TRUE)

  GSEA_res <- tryCatch({
    GSEA(geneList, TERM2GENE = m_t2g, pvalueCutoff = 0.05, seed = TRUE, verbose = FALSE)
  }, error = function(e) return(NULL))

  if(!is.null(GSEA_res) && nrow(GSEA_res@result) > 0) {
    dot_df <- GSEA_res@result %>%
      mutate(SIG = -log10(p.adjust))

    p <- ggplot(dot_df, aes(x = NES, y = fct_reorder(Description, NES))) +
      geom_point(aes(size = SIG, color = NES)) +
      scale_colour_gradient2(low = "blue", mid = "white", high = "red") +
      theme_minimal() +
      labs(title = paste0("GSEA Hallmark: ", prefix), y = NULL)

    ggsave(paste0("GSEA_Plot_", prefix, ".pdf"), p, width = 10, height = 8)
    cat("Analysis completed and output saved\n")
  } else {
    cat("No significant enrichment was detected\n")
  }
}

# =====================================================================
# =====================================================================
#run_single_iso_gsea("ENST00000352993.7", top_pct = 0.25, bottom_pct = 0.25)
run_single_iso_gsea("ENST00000357654.9", top_pct = 0.25, bottom_pct = 0.25)


# =====================================================================
# =====================================================================
library(dplyr)
library(ggplot2)
library(clusterProfiler)
library(msigdbr)
library(stringr)
library(forcats)

# =====================================================================
# =====================================================================
csv_file <- "./DEGs_ENST00000352993_Top25_vs_Low25.csv"
comp_name <- "ENST00000352993_High_vs_Low"

m_t2g <- msigdbr(species = "Homo sapiens", category = "H") %>%
  dplyr::select(gs_name, entrez_gene)

# =====================================================================
# =====================================================================
# =====================================================================
# =====================================================================
gsea_results_list <- list()

res_df <- read.csv(csv_file)

print("Analysis progress")
print(colnames(res_df))

geneList <- sort(setNames(res_df$stat, res_df$ENTREZ), decreasing = TRUE)

gsea_result <- tryCatch({
  GSEA(geneList, TERM2GENE = m_t2g, pvalueCutoff = 0.05, seed = TRUE, verbose = FALSE)
}, error = function(e) return(NULL))


if(!is.null(gsea_result) && nrow(gsea_result@result) > 0) {
  gene_count <- gsea_result@result %>%
    group_by(ID) %>%
    summarise(count = sum(str_count(core_enrichment, "/")) + 1)

  dot_df <- left_join(gsea_result@result, gene_count, by = "ID") %>%
    mutate(GeneRatio = count/setSize)

  gsea_results_list[[comp_name]] <- dot_df
}

# =====================================================================
# =====================================================================
if(length(gsea_results_list) > 0) {

  merged.res <- as.data.frame(merge_result(gsea_results_list))

  merged.res$type = "upregulated"
  merged.res$type[merged.res$NES < 0] = "downregulated"
  merged.res$SIG = -log10(merged.res$p.adjust)

  p <- ggplot(merged.res, aes(x = NES, y = fct_reorder(Description, SIG))) +
    geom_point(aes(size = SIG, color = NES)) +
    theme_bw(base_size = 14) +
    scale_colour_gradient2(limits=c(min(merged.res$NES), max(merged.res$NES)),
                           low = "#3953A4", mid = "white", high = "#b50404") +
    ylab(NULL) +
    facet_grid(.~Cluster) +
    scale_size(range = c(3, 12)) +
    labs(title = "GSEA Hallmark Pathways",
         size = "-log10(padj)",
         color = "NES")

  print(p)
  ggsave("./GSEA_NES_Style.png", p, width = 12, height = 10)

  write.csv(merged.res, "./GSEA_merged_results.csv", row.names = FALSE)

  cat("Analysis completed and output saved\n")
} else {
  cat("No significant enrichment was detected\n")
}


library(dplyr)
library(ggplot2)
library(clusterProfiler)
library(msigdbr)
library(stringr)
library(forcats)

# =====================================================================
# =====================================================================
m_t2g <- msigdbr(species = "Homo sapiens", category = "H") %>%
  dplyr::select(gs_name, entrez_gene)

files_list <- list(
  "BRCA1_T" = "./DEGs_ENST00000352993_Top25_vs_Low25.csv",
  "BRCA1_C" = "./DEGs_ENST00000357654_Top25_vs_Low25.csv"
)

# =====================================================================
# =====================================================================
gsea_results_list <- list()

for (comp_name in names(files_list)) {
  csv_file <- files_list[[comp_name]]

  if (!file.exists(csv_file)) {
    warning(paste("Required input was not found", csv_file))
    next
  }

  res_df <- read.csv(csv_file)

  geneList <- sort(setNames(res_df$stat, res_df$ENTREZ), decreasing = TRUE)

  gsea_result <- tryCatch({
    GSEA(geneList, TERM2GENE = m_t2g, pvalueCutoff = 0.05, seed = TRUE, verbose = FALSE)
  }, error = function(e) return(NULL))

  if(!is.null(gsea_result) && nrow(gsea_result@result) > 0) {
    gene_count <- gsea_result@result %>%
      group_by(ID) %>%
      summarise(count = sum(str_count(core_enrichment, "/")) + 1)

    dot_df <- left_join(gsea_result@result, gene_count, by = "ID") %>%
      mutate(GeneRatio = count/setSize)

    gsea_results_list[[comp_name]] <- dot_df
  }
}

# =====================================================================
# =====================================================================
if(length(gsea_results_list) > 0) {

  merged.res <- as.data.frame(merge_result(gsea_results_list))

  merged.res$type = "upregulated"
  merged.res$type[merged.res$NES < 0] = "downregulated"
  merged.res$SIG = -log10(merged.res$p.adjust)

  p <- ggplot(merged.res, aes(x = NES, y = fct_reorder(Description, SIG))) +
    geom_point(aes(size = SIG, color = NES, shape = Cluster), stroke = 1.2) +
    theme_bw(base_size = 14) +
    scale_colour_gradient2(limits=c(min(merged.res$NES), max(merged.res$NES)),
                           low = "#3953A4", mid = "white", high = "#b50404") +
    scale_shape_manual(values = c("BRCA1_T" = 16, "BRCA1_C" = 1)) +
    scale_size(range = c(3, 12)) +
    ylab(NULL) +
    # facet_grid(.~Cluster) +
    labs(title = "GSEA Hallmark Pathways Comparison",
         size = "-log10(padj)",
         color = "NES",
         shape = "Group") +
    theme(
      legend.position = "right",
      panel.grid.major.y = element_line(linetype = "dashed", color = "gray80")
    )

  print(p)
  ggsave("./GSEA_NES_Combined_Style.pdf", p, width = 12, height = 10)

 # write.csv(merged.res, "./GSEA_merged_combined_results.csv", row.names = FALSE)

  cat("Analysis completed and output saved\n")
} else {
  cat("No significant enrichment was detected\n")
}
```
