## Supplementary material for "An integrated, transcript-resolution atlas of prostate cancer progression unmasks associations with oncogenic pathways": Reproducible code: RNF43_Heteroscedasticity_4_Separate_Plots.R

```
# Public submission copy.
# Analytical statements and parameter values are unchanged from the archived source.
# Legacy non-English comments and console messages were normalized for public release.
library(fst)
library(dplyr)
library(ggplot2)
library(ggrepel)
library(quantreg)

cat("Loading input data\n")
txi_df <- read.fst("./txi.counts.fst")
txi_info <- read.csv("./transcripts_info.csv")
rownames(txi_df) <- txi_info$transcript_id
txi_counts <- as.matrix(txi_df)
ncount.tx <- log2(sweep(x = txi_counts, MARGIN = 2, STATS = colSums(txi_counts)/1000000, FUN = "/") + 1)
rownames(ncount.tx) <- gsub("\\..*$", "", rownames(ncount.tx))

gc_df <- read.fst("./gc_txi.counts.fst")
gc_info <- read.csv("./genes_info.csv")
rownames(gc_df) <- gc_info$gene_id
gc_counts <- as.matrix(gc_df)
ncount.gc <- log2(sweep(x = gc_counts, MARGIN = 2, STATS = colSums(gc_counts)/1000000, FUN = "/") + 1)

gc_info$base_gene_id <- gsub("\\..*$", "", gc_info$gene_id)
gene_means <- rowMeans(ncount.gc)
gc_mapped <- data.frame(
  base_gene_id = gsub("\\..*$", "", rownames(ncount.gc)),
  mean_expr = gene_means,
  stringsAsFactors = FALSE
) %>%
  left_join(gc_info %>% select(base_gene_id, gene_name) %>% distinct(), by = "base_gene_id") %>%
  filter(!is.na(gene_name) & gene_name != "") %>%
  group_by(gene_name) %>%
  slice_max(order_by = mean_expr, n = 1, with_ties = FALSE) %>%
  ungroup()

ncount.gc_symbol <- ncount.gc[rownames(ncount.gc) %in% gc_info$gene_id[match(gc_mapped$base_gene_id, gc_info$base_gene_id)], ]
rownames(ncount.gc_symbol) <- gc_mapped$gene_name[match(gsub("\\..*$", "", rownames(ncount.gc_symbol)), gc_mapped$base_gene_id)]

rspo_genes <- c("RSPO1", "RSPO2", "RSPO3", "RSPO4")
valid_rspo <- intersect(rspo_genes, rownames(ncount.gc_symbol))
rspo_expr_matrix <- ncount.gc_symbol[valid_rspo, , drop = FALSE]
rspo_mean_expr <- colMeans(rspo_expr_matrix)

load("Full_GSVA_Analysis_Workspace_2026-03-23.Rdata")

iso_A <- "ENST00000584437"
iso_B <- "ENST00000583753"
target_pathway <- "WNT_UP.V1_UP"

expr_A <- ncount.tx[iso_A, ]
expr_B <- ncount.tx[iso_B, ]
pathway_scores <- full_gsva_res[target_pathway, ]

common_samples <- intersect(intersect(names(expr_A), names(pathway_scores)), names(rspo_mean_expr))

df <- data.frame(
  Sample = common_samples,
  IsoA = expr_A[common_samples],
  IsoB = expr_B[common_samples],
  RSPO = rspo_mean_expr[common_samples],
  Score_Original = pathway_scores[common_samples]
)

cat("Analysis progress\n")
lm_rspo <- lm(Score_Original ~ RSPO, data = df)
df$Score_Adjusted <- resid(lm_rspo)

df$Score_Original_Z <- scale(df$Score_Original)[, 1]
df$Score_Adjusted_Z <- scale(df$Score_Adjusted)[, 1]

out_dir <- "RNF43_WNT_NonHallmark_2D"
dir.create(out_dir, showWarnings = FALSE)

cat("Analysis progress\n")
plot_single_quantile <- function(df, x_var, y_var, color_var, x_label, y_label, title, subtitle) {
  ggplot(df, aes(x = .data[[x_var]], y = .data[[y_var]])) +
    geom_point(aes(color = .data[[color_var]]), alpha = 0.8, size = 3) +
    scale_color_gradient2(
      low = "#3953A4", mid = "white", high = "#b50404", midpoint = 0,
      name = "ssGSEA Score\n(Z-score)"
    ) +
    geom_quantile(quantiles = 0.1, linewidth = 1.2, color = "blue", alpha=0.8) +
    geom_quantile(quantiles = 0.5, linewidth = 1.2, color = "black", alpha=0.8) +
    geom_quantile(quantiles = 0.9, linewidth = 1.2, color = "red", alpha=0.8) +
    theme_bw(base_size = 14) +
    labs(
      title = title,
      subtitle = subtitle,
      x = x_label,
      y = y_label
    ) +
    theme(
      panel.grid.minor = element_blank(),
      legend.position = "right",
      plot.title = element_text(face = "bold", size = 11)
    )
}

# 1. IsoA Original
p1 <- plot_single_quantile(df, "IsoA", "Score_Original", "Score_Original_Z",
                           paste0("Isoform A: ", iso_A, " (log2 CPM)"),
                           paste0(target_pathway, " Score"),
                           "RNF43 IsoA vs WNT Score (Original)",
                           "Includes RSPO effects. Lines: 10th (blue), 50th (black), 90th (red) percentiles")
ggsave(file.path(out_dir, "Quantile_Scatter_IsoA_Original.pdf"), p1, width = 8, height = 7)

# 2. IsoB Original
p2 <- plot_single_quantile(df, "IsoB", "Score_Original", "Score_Original_Z",
                           paste0("Isoform B: ", iso_B, " (log2 CPM)"),
                           paste0(target_pathway, " Score"),
                           "RNF43 IsoB vs WNT Score (Original)",
                           "Includes RSPO effects. Lines: 10th (blue), 50th (black), 90th (red) percentiles")
ggsave(file.path(out_dir, "Quantile_Scatter_IsoB_Original.pdf"), p2, width = 8, height = 7)

# 3. IsoA Adjusted
p3 <- plot_single_quantile(df, "IsoA", "Score_Adjusted", "Score_Adjusted_Z",
                           paste0("Isoform A: ", iso_A, " (log2 CPM)"),
                           paste0(target_pathway, " Score (RSPO Adjusted)"),
                           "RNF43 IsoA vs WNT Score (RSPO Adjusted)",
                           "Regressed out RSPO. Lines: 10th (blue), 50th (black), 90th (red) percentiles")
ggsave(file.path(out_dir, "Quantile_Scatter_IsoA_RSPO_Adjusted.pdf"), p3, width = 8, height = 7)

# 4. IsoB Adjusted
p4 <- plot_single_quantile(df, "IsoB", "Score_Adjusted", "Score_Adjusted_Z",
                           paste0("Isoform B: ", iso_B, " (log2 CPM)"),
                           paste0(target_pathway, " Score (RSPO Adjusted)"),
                           "RNF43 IsoB vs WNT Score (RSPO Adjusted)",
                           "Regressed out RSPO. Lines: 10th (blue), 50th (black), 90th (red) percentiles")
ggsave(file.path(out_dir, "Quantile_Scatter_IsoB_RSPO_Adjusted.pdf"), p4, width = 8, height = 7)

cat("Analysis completed and output saved\n")
```
