## Supplementary material for "An integrated, transcript-resolution atlas of prostate cancer progression unmasks associations with oncogenic pathways": Reproducible code: TRIM11_isoform_AR_analysis_ssGSEA.R

```
# Public submission copy.
# Analytical statements and parameter values are unchanged from the archived source.
# Legacy non-English comments and console messages were normalized for public release.
# =====================================================================
# TRIM11 Isoforms vs AR Pathways (Using ssGSEA)
# =====================================================================

neededlibraries <- c("fst", "dplyr", "ggplot2", "ggpubr", "msigdbr", "tidyr", "patchwork", "GSVA")
for (pkg in neededlibraries) {
  if (!requireNamespace(pkg, quietly = TRUE)) {
    if (pkg == "GSVA") BiocManager::install("GSVA", update = FALSE, ask = FALSE)
    else install.packages(pkg, repos = "https://cloud.r-project.org", ask = FALSE, update = FALSE)
  }
  library(pkg, character.only = TRUE)
}

cat("Loading input data\n")
txi_df <- read.fst("./txi.counts.fst")
txi_info <- read.csv("./transcripts_info.csv")
rownames(txi_df) <- txi_info$transcript_id
txi_counts <- as.matrix(txi_df)
ncount.tx <- log2(sweep(x = txi_counts, MARGIN = 2, STATS = colSums(txi_counts)/1000000, FUN = "/") + 1)

gc_df <- read.fst("./gc_txi.counts.fst")
gc_info <- read.csv("./genes_info.csv")
rownames(gc_df) <- gc_info$gene_id
gc_counts <- as.matrix(gc_df)
gc_counts <- gc_counts[rowSums(gc_counts) > 0, ]
ncount.gc <- log2(sweep(x = gc_counts, MARGIN = 2, STATS = colSums(gc_counts)/1000000, FUN = "/") + 1)

gc_info$base_gene_id <- gsub("\\..*$", "", gc_info$gene_id)
gene_means <- rowMeans(ncount.gc)
gc_mapped <- data.frame(
  base_gene_id = gsub("\\..*$", "", rownames(ncount.gc)),
  mean_expr = gene_means,
  stringsAsFactors = FALSE
) %>%
  left_join(gc_info %>% select(base_gene_id, gene_name) %>% distinct(), by = "base_gene_id") %>%
  filter(!is.na(gene_name) & gene_name != "") %>%
  group_by(gene_name) %>%
  slice_max(order_by = mean_expr, n = 1, with_ties = FALSE) %>%
  ungroup()

ncount.gc_symbol <- ncount.gc[rownames(ncount.gc) %in% gc_info$gene_id[match(gc_mapped$base_gene_id, gc_info$base_gene_id)], ]
rownames(ncount.gc_symbol) <- gc_mapped$gene_name[match(gsub("\\..*$", "", rownames(ncount.gc_symbol)), gc_mapped$base_gene_id)]

# =====================================================================
# =====================================================================
iso1_id <- "ENST00000284551"
iso2_id <- "ENST00000493030"

get_iso_expr <- function(id, mat) {
  idx <- which(gsub("\\..*$", "", rownames(mat)) == id)
  if (length(idx) == 0) return(rep(NA, ncol(mat)))
  return(mat[idx[1], ])
}
expr_iso1 <- get_iso_expr(iso1_id, ncount.tx)
expr_iso2 <- get_iso_expr(iso2_id, ncount.tx)

delta_expr <- expr_iso1 - expr_iso2

# =====================================================================
# =====================================================================
cat("Running pathway analysis\n")

gene_sets <- list()

# 4.1 Custom AR Gene Set
custom_ar_genes <- c("KLK3", "KLK2", "TMPRSS2", "FKBP5", "NKX3-1", "NKX3.1", "PLPP1", "PMEPA1", "PART1", "ALDH1A3", "STEAP4")
valid_custom_ar <- custom_ar_genes[custom_ar_genes %in% rownames(ncount.gc_symbol)]
if(length(valid_custom_ar) > 0) gene_sets[["Custom_AR_Gene_Set"]] <- valid_custom_ar

# 4.2 Hallmark Androgen Response
suppressWarnings({
  m_t2g_h <- msigdbr(species = "Homo sapiens", category = "H") %>% filter(gs_name == "HALLMARK_ANDROGEN_RESPONSE")
})
valid_hallmark <- unique(m_t2g_h$gene_symbol)[unique(m_t2g_h$gene_symbol) %in% rownames(ncount.gc_symbol)]
if(length(valid_hallmark) > 0) gene_sets[["HALLMARK_ANDROGEN_RESPONSE"]] <- valid_hallmark

# 4.3 GO MF/BP with ANDROGEN
suppressWarnings({
  m_t2g_go <- msigdbr(species = "Homo sapiens", category = "C5") %>%
    filter(grepl("^GO.*ANDROGEN", gs_name, ignore.case = TRUE))
})

for (pw in unique(m_t2g_go$gs_name)) {
  genes <- m_t2g_go$gene_symbol[m_t2g_go$gs_name == pw]
  valid_genes <- genes[genes %in% rownames(ncount.gc_symbol)]
  if (length(valid_genes) >= 3) {
    gene_sets[[pw]] <- valid_genes
  }
}

cat(sprintf("Analysis progress: %d\n", length(gene_sets)))

if (packageVersion("GSVA") >= "1.50.0") {
  ssgsea_res <- gsva(ssgseaParam(ncount.gc_symbol, gene_sets), verbose=FALSE)
} else {
  ssgsea_res <- gsva(ncount.gc_symbol, gene_sets, method="ssgsea", verbose=FALSE)
}

plot_df <- data.frame(
  Sample = colnames(ncount.tx),
  Iso1 = expr_iso1,
  Iso2 = expr_iso2,
  Delta = delta_expr
)
plot_df$AR_Expression <- if ("AR" %in% rownames(ncount.gc_symbol)) ncount.gc_symbol["AR", ] else NA

for (pw in rownames(ssgsea_res)) {
  plot_df[[pw]] <- ssgsea_res[pw, ]
}

# =====================================================================
# =====================================================================
cat("Assigning analysis groups\n")
top_cutoff <- quantile(plot_df$Delta, 0.75, na.rm = TRUE)
bot_cutoff <- quantile(plot_df$Delta, 0.25, na.rm = TRUE)

n_top <- sum(plot_df$Delta >= top_cutoff, na.rm=TRUE)
n_bot <- sum(plot_df$Delta <= bot_cutoff, na.rm=TRUE)

group_top_name <- paste0(iso1_id, " Dominant\n(n=", n_top, ")")
group_bot_name <- paste0(iso2_id, " Dominant\n(n=", n_bot, ")")

plot_df <- plot_df %>%
  mutate(Group = case_when(
    Delta >= top_cutoff ~ group_top_name,
    Delta <= bot_cutoff ~ group_bot_name,
    TRUE ~ "Intermediate"
  ))

box_df <- plot_df %>% filter(Group != "Intermediate")
box_df$Group <- factor(box_df$Group, levels = c(group_top_name, group_bot_name))

features_to_plot <- c("AR_Expression", rownames(ssgsea_res))

# =====================================================================
# =====================================================================
cat("Analysis progress\n")

my_comparisons <- list(c(group_top_name, group_bot_name))
colors <- c("#E78AC3", "#8DA0CB")

boxplots_list <- list()
for (feat in features_to_plot) {
  short_title <- gsub("^GO[A-Z]*_", "", feat)
  short_title <- gsub("_", " ", short_title)

  y_label <- ifelse(feat == "AR_Expression", "log2(CPM+1)", "ssGSEA Score")

  p <- ggboxplot(box_df, x = "Group", y = feat, fill = "Group", palette = colors,
                 add = "jitter", add.params = list(size = 0.2, alpha = 0.3), outlier.shape=NA) +
    stat_compare_means(comparisons = my_comparisons, method = "wilcox.test", size=3.5) +
    labs(title = stringr::str_wrap(short_title, width=30), x = NULL, y = y_label) +
    theme_bw(base_size = 12) +
    theme(legend.position = "none",
          plot.title = element_text(size=9, face="bold"),
          axis.text.x = element_text(angle = 15, hjust = 1))

  boxplots_list[[feat]] <- p
}

final_boxplots <- wrap_plots(boxplots_list, ncol = 3) +
  plot_annotation(title = "TRIM11 Isoform Dominance vs AR Pathways (ssGSEA)",
                  theme = theme(plot.title = element_text(size = 18, face = "bold")))
ggsave("./TRIM11_AR_Pathways_ssGSEA_Boxplots.pdf", final_boxplots, width = 15, height = 5 * ceiling(length(features_to_plot)/3))
ggsave("./TRIM11_AR_Pathways_ssGSEA_Boxplots.png", final_boxplots, width = 15, height = 5 * ceiling(length(features_to_plot)/3), dpi = 300)

# =====================================================================
# =====================================================================
cat("Analysis progress\n")

plot_scatter <- function(df, x_col, y_col, x_label, y_label, color, short_title) {
  ggscatter(df, x = x_col, y = y_col, color = color, alpha = 0.3, size = 1,
            add = "reg.line", add.params = list(color = "black", linetype = "dashed")) +
    stat_cor(method = "pearson", label.x.npc = "left", label.y.npc = "top", size=3) +
    labs(title = stringr::str_wrap(short_title, width=30), x = x_label, y = y_label) +
    theme_bw(base_size = 10) +
    theme(plot.title = element_text(size=8, face="bold"))
}

iso1_scatters <- list()
iso2_scatters <- list()

for (feat in features_to_plot) {
  short_title <- gsub("^GO[A-Z]*_", "", feat)
  short_title <- gsub("_", " ", short_title)
  y_label <- ifelse(feat == "AR_Expression", "Expression", "ssGSEA")

  iso1_scatters[[feat]] <- plot_scatter(plot_df, "Iso1", feat, paste0(iso1_id), y_label, "#66C2A5", short_title)
  iso2_scatters[[feat]] <- plot_scatter(plot_df, "Iso2", feat, paste0(iso2_id), y_label, "#FC8D62", short_title)
}

final_scatter_iso1 <- wrap_plots(iso1_scatters, ncol = 4) +
  plot_annotation(title = paste0("Correlation: ", iso1_id, " Expression vs AR Pathways (ssGSEA)"))
final_scatter_iso2 <- wrap_plots(iso2_scatters, ncol = 4) +
  plot_annotation(title = paste0("Correlation: ", iso2_id, " Expression vs AR Pathways (ssGSEA)"))

ggsave("./TRIM11_AR_Pathways_ssGSEA_Scatter_Iso1.pdf", final_scatter_iso1, width = 14, height = 3 * ceiling(length(features_to_plot)/4))
ggsave("./TRIM11_AR_Pathways_ssGSEA_Scatter_Iso2.pdf", final_scatter_iso2, width = 14, height = 3 * ceiling(length(features_to_plot)/4))

cat("Analysis completed and output saved\n")
```
