## Supplementary material for "An integrated, transcript-resolution atlas of prostate cancer progression unmasks associations with oncogenic pathways": Reproducible code: WNT2B_pathway_network.R

```
# Public submission copy.
# Analytical statements and parameter values are unchanged from the archived source.
# Legacy non-English comments and console messages were normalized for public release.
# AWSI–Hallmark pathway similarity network
#
# Purpose
# -------
# Build a pathway–pathway network from the same AWSI results used for
# Figure S9B. Nodes are Hallmark pathways and edges are based on Jaccard
# similarity of the exact gene sets stored in the GSVA workspace. The graph is
# drawn with igraph/ggraph, while following the similarity logic used by
# clusterProfiler/enrichplot::emapplot().
#
# This script does not rerun GSVA, AWSI correlations, or GSEA. It uses the
# existing *_Continuous_AWSI_with_CI.csv as node-level statistics.

suppressPackageStartupMessages({
  library(igraph)
  library(ggraph)
  library(ggplot2)
  library(dplyr)
  library(readr)
  library(scales)
})

# -------------------------------------------------------------------------
# 1. Paths and user-configurable parameters
# -------------------------------------------------------------------------

data_root <- "__PCATLAS_DOCUMENTS_ROOT__/Data_Harmanization"
output_root <- "__PCATLAS_PROJECT_ROOT__/AWSI_pathway_networks"
args <- commandArgs(trailingOnly = TRUE)
gene_id <- if (length(args) >= 1L) args[[1]] else "WNT2B"
standard_pdf <- identical(Sys.getenv("PDF_SUFFIX", ""), "_standardpdf")
aws_label <- if (standard_pdf) "AWSI-pathway\nSpearman rho" else "AWSI–pathway\nSpearman ρ"
minus_log_fdr_label <- if (standard_pdf) "-log10(FDR)" else "−log10(FDR)"
aws_results_file <- if (length(args) >= 2L) args[[2]] else {
  file.path(data_root, "WNT2B/WNT2B_Continuous_AWSI_with_CI.csv")
}
result_dir <- if (length(args) >= 3L) args[[3]] else {
  file.path(output_root, gene_id)
}
category_mode <- if (length(args) >= 4L) args[[4]] else "HALLMARK"
if (!category_mode %in% c("HALLMARK", "H_C2_C6")) {
  stop("category_mode must be 'HALLMARK' or 'H_C2_C6'.")
}
category_label <- if (category_mode == "HALLMARK") "Hallmark" else "H_C2_C6"
file_prefix <- paste0(gene_id, "_", category_label)
dir.create(result_dir, recursive = TRUE, showWarnings = FALSE)

gsva_workspace <- file.path(data_root, "Full_GSVA_Analysis_Workspace_2026-03-23.Rdata")

# S9B used Hallmark pathways with FDR <= 0.05.
fdr_cutoff <- 0.05
hallmark_prefix <- "HALLMARK_"

# Recommended primary threshold. A Jaccard edge is retained when two Hallmark
# gene sets share at least this fraction of their union.
jaccard_cutoff <- 0.05

# Additional minimum shared-gene safeguard. Set to 0 to use Jaccard alone.
min_shared_genes <- 0L

# TRUE: retain only FDR-significant Hallmark nodes, matching S9B.
# FALSE: retain all 50 Hallmark pathways and mark non-significant nodes.
significant_only <- TRUE

# To reproduce enrichplot-like edge semantics, use Jaccard. The alternative
# "overlap" option uses intersection/min(set sizes), but is not the default.
edge_metric <- "jaccard"

# Plotting options.
layout_method <- "fr"  # compact Fruchterman–Reingold layout
# Labels are shown only for strong, highly significant AWSI associations.
# Nodes and edges are retained independently of these display thresholds.
label_abs_cor_cutoff <- 0.4
label_p_cutoff <- 0.001
label_candidate_abs_cor_cutoff <- 0.2
label_candidate_p_cutoff <- 0.05
label_fraction_target <- 0.20
label_fraction_min <- 0.20
label_fraction_max <- 0.40
group_label_min_nodes <- 4L
group_label_max_terms <- 1L
node_size_min <- 2.5
node_size_max <- 5.5
edge_width_min <- 0.8
edge_width_max <- 2.0
figure_size_cm <- 10
figure_text_pt <- 5

# -------------------------------------------------------------------------
# 2. Load the original S9B results and the original gene-set workspace
# -------------------------------------------------------------------------

if (!file.exists(aws_results_file)) {
  stop("Missing AWSI result file: ", aws_results_file)
}
if (!file.exists(gsva_workspace)) {
  stop("Missing GSVA workspace: ", gsva_workspace)
}

aws <- readr::read_csv(aws_results_file, show_col_types = FALSE) %>%
  mutate(Pathway = as.character(Pathway))

workspace_env <- new.env(parent = emptyenv())
load(gsva_workspace, envir = workspace_env)

if (!exists("all_signatures_list", envir = workspace_env, inherits = FALSE)) {
  stop("The workspace does not contain all_signatures_list.")
}

all_signatures_list <- get("all_signatures_list", envir = workspace_env)
all_signatures_list <- lapply(all_signatures_list, function(x) unique(as.character(x)))

# Build the pathway catalogue used for node selection. For the expanded mode,
# C6 names do not carry a universal prefix, so use the saved PDX CS.names list
# to identify the exact 189 C6 signatures rather than guessing from names.
if (category_mode == "HALLMARK") {
  pathway_catalog <- tibble(
    pathway = grep("^HALLMARK_", names(all_signatures_list), value = TRUE),
    category = "Hallmark"
  )
} else {
  cs_file <- "__PCATLAS_DOCUMENTS_ROOT__/PDX/objects/CS.names.RDS"
  if (!file.exists(cs_file)) stop("Missing CS.names.RDS needed to identify C6 signatures.")
  cs_names <- readRDS(cs_file)
  h_names <- cs_names[grepl("^HALLMARK_", cs_names)]
  pid_names <- cs_names[grepl("^PID_", cs_names)]
  c6_names <- setdiff(cs_names, c("AR_GSET", "NE_GSET", h_names, pid_names))
  pathway_catalog <- bind_rows(
    tibble(pathway = h_names, category = "Hallmark"),
    tibble(pathway = pid_names, category = "C2_PID"),
    tibble(pathway = c6_names, category = "C6_oncogenic")
  )
}
pathway_catalog <- pathway_catalog %>%
  filter(pathway %in% names(all_signatures_list)) %>%
  distinct(pathway, .keep_all = TRUE)

# -------------------------------------------------------------------------
# 3. Select nodes exactly as in Figure S9B and attach node statistics
# -------------------------------------------------------------------------

hallmark_results <- aws %>%
  inner_join(pathway_catalog, by = c("Pathway" = "pathway"))

if (significant_only) {
  node_table <- hallmark_results %>% filter(!is.na(FDR), FDR <= fdr_cutoff)
} else {
  node_table <- hallmark_results
}

node_table <- node_table %>%
  filter(Pathway %in% names(all_signatures_list)) %>%
  mutate(
    pathway = Pathway,
    label = sub("^(HALLMARK_|PID_)", "", Pathway),
    rho = as.numeric(Correlation),
    fdr = as.numeric(FDR),
    neg_log10_fdr = -log10(pmax(fdr, .Machine$double.xmin)),
    gene_set_size = vapply(all_signatures_list[Pathway], length, integer(1))
  ) %>%
  distinct(pathway, .keep_all = TRUE)

if (nrow(node_table) < 2) {
  stop("Fewer than two usable pathways remain after filtering.")
}

gene_sets <- all_signatures_list[node_table$pathway]
names(gene_sets) <- node_table$pathway

# -------------------------------------------------------------------------
# 4. Construct pathway similarity edges
# -------------------------------------------------------------------------

pair_similarity <- function(a, b, metric = c("jaccard", "overlap")) {
  metric <- match.arg(metric)
  shared <- length(intersect(a, b))
  if (metric == "jaccard") {
    denom <- length(union(a, b))
  } else {
    denom <- min(length(a), length(b))
  }
  score <- if (denom == 0) 0 else shared / denom
  c(shared_genes = shared, similarity = score)
}

edge_rows <- list()
edge_index <- 1L
if (length(gene_sets) >= 2) {
  for (i in seq_len(length(gene_sets) - 1L)) {
    for (j in seq.int(i + 1L, length(gene_sets))) {
      sim <- pair_similarity(gene_sets[[i]], gene_sets[[j]], edge_metric)
      if (sim[["similarity"]] >= jaccard_cutoff &&
          sim[["shared_genes"]] >= min_shared_genes) {
        edge_rows[[edge_index]] <- data.frame(
          from = names(gene_sets)[i],
          to = names(gene_sets)[j],
          shared_genes = as.integer(sim[["shared_genes"]]),
          similarity = as.numeric(sim[["similarity"]]),
          stringsAsFactors = FALSE
        )
        edge_index <- edge_index + 1L
      }
    }
  }
}

edges <- if (length(edge_rows)) bind_rows(edge_rows) else {
  data.frame(from = character(), to = character(), shared_genes = integer(),
             similarity = numeric(), stringsAsFactors = FALSE)
}

# -------------------------------------------------------------------------
# 5. Build graph and calculate communities
# -------------------------------------------------------------------------

vertices <- node_table %>%
  select(pathway, label, category, rho, P_value, fdr, neg_log10_fdr, gene_set_size)

graph <- graph_from_data_frame(
  d = edges,
  directed = FALSE,
  vertices = vertices %>% rename(name = pathway)
)

if (ecount(graph) > 0) {
  graph <- set_edge_attr(graph, "weight", value = E(graph)$similarity)
  set.seed(20260811)
  community <- cluster_louvain(graph, weights = E(graph)$similarity)
  V(graph)$community <- as.character(membership(community)[V(graph)$name])
} else {
  V(graph)$community <- "1"
}

# Isolated nodes remain in the graph and are deliberately retained.
# They are biologically meaningful: no edge above the declared similarity
# threshold is not equivalent to missing data.

# -------------------------------------------------------------------------
# 6. Export underlying node and edge data for auditability
# -------------------------------------------------------------------------

node_export <- igraph::as_data_frame(graph, what = "vertices") %>%
  rename(pathway = name)
edge_export <- igraph::as_data_frame(graph, what = "edges")

# Give each numeric Louvain community a reproducible descriptive name based
# on its strongest AWSI-associated Hallmark terms.  Keep the numeric ID for
# auditability and add the human-readable name as a separate CSV column.
community_name_map <- node_export %>%
  group_by(community) %>%
  arrange(desc(abs(rho)), fdr, .by_group = TRUE) %>%
  summarise(
    community_name = paste(head(label, group_label_max_terms), collapse = " · "),
    .groups = "drop"
  )

# Author-confirmed community labels from the annotated review copy.  The
# numeric Louvain IDs remain unchanged; only the displayed descriptive names
# are replaced where a biological interpretation was explicitly supplied.
community_label_overrides <- list(
  ROCK2 = c(
    APOPTOSIS = "Inflammation; STAT signaling",
    ADIPOGENESIS = "Lipid metabolism",
    UV_RESPONSE_DN = "DNA repair; Epithelial-to-Mesenchymal Transition"
  ),
  RNF43 = c(
    OXIDATIVE_PHOSPHORYLATION = "Energy metabolism",
    UV_RESPONSE_UP = "DNA repair; Epithelial-to-Mesenchymal Transition",
    MITOTIC_SPINDLE = "Cell cycle; MYC"
  ),
  RELA = c(
    IL2_STAT5_SIGNALING = "Inflammation; STAT signaling",
    P53_PATHWAY = "DNA repair",
    MITOTIC_SPINDLE = "Cell cycle; MYC",
    PEROXISOME = "Lipid metabolism"
  ),
  TRIM11 = c(
    CHOLESTEROL_HOMEOSTASIS = "Cholesterol homeostasis; mTOR signalling",
    UV_RESPONSE_UP = "DNA repair; Inflammation",
    MITOTIC_SPINDLE = "Cell cycle; MYC",
    PEROXISOME = "Energy metabolism"
  ),
  WNT2B = c(
    ADIPOGENESIS = "Lipid metabolism",
    MYOGENESIS = "Epithelial-to-Mesenchymal Transition",
    E2F_TARGETS = "Cell cycle",
    APOPTOSIS = "Inflammation; STAT signaling"
  )
)
if (gene_id %in% names(community_label_overrides)) {
  override <- community_label_overrides[[gene_id]]
  community_name_map$community_name <- vapply(
    seq_len(nrow(community_name_map)),
    function(i) {
      old <- community_name_map$community_name[i]
      key <- strsplit(old, " · ", fixed = TRUE)[[1]][1]
      if (key %in% names(override)) unname(override[[key]]) else old
    },
    character(1)
  )
}
node_export <- node_export %>%
  left_join(community_name_map, by = "community")

write_csv(node_export, file.path(result_dir, paste0(file_prefix, "_network_nodes.csv")))
write_csv(edge_export, file.path(result_dir, paste0(file_prefix, "_network_edges.csv")))

writeLines(capture.output(sessionInfo()), file.path(result_dir, "sessionInfo.txt"))

network_summary <- data.frame(
  n_nodes = vcount(graph),
  n_edges = ecount(graph),
  category_mode = category_mode,
  category_label = category_label,
  n_candidate_signatures = nrow(pathway_catalog),
  edge_metric = edge_metric,
  similarity_cutoff = jaccard_cutoff,
  min_shared_genes = min_shared_genes,
  significant_only = significant_only,
  stringsAsFactors = FALSE
)
write_csv(network_summary, file.path(result_dir, paste0(file_prefix, "_network_summary.csv")))

# -------------------------------------------------------------------------
# 7. Custom ggraph plot, using enrichplot-like encodings
# -------------------------------------------------------------------------

set.seed(20260811)
layout_fun <- switch(
  layout_method,
  kk = "kk",
  fr = "fr",
  stop("layout_method must be 'kk' or 'fr'.")
)

# Normalize node-size and edge-width mappings explicitly so the plot is
# comparable across re-runs and across genes.
node_export <- node_export %>%
  mutate(
    node_size = rescale(neg_log10_fdr, to = c(node_size_min, node_size_max)),
    label_score = abs(rho) * pmax(-log10(pmax(as.numeric(P_value), .Machine$double.xmin)), 0),
    label_candidate = !is.na(rho) & !is.na(P_value) &
      abs(rho) >= label_candidate_abs_cor_cutoff &
      P_value <= label_candidate_p_cutoff
  )

# Select a comparable fraction of labels in every figure. Candidates still
# need a non-trivial correlation and nominal significance; ranking then favors
# the most extreme and most significant associations.
n_nodes <- nrow(node_export)
n_label_target <- min(
  ceiling(label_fraction_max * n_nodes),
  max(ceiling(label_fraction_min * n_nodes), ceiling(label_fraction_target * n_nodes))
)
label_candidates <- which(node_export$label_candidate)
if (length(label_candidates) > 0L) {
  label_candidates <- label_candidates[order(-node_export$label_score[label_candidates],
                                              -abs(node_export$rho[label_candidates]),
                                              node_export$P_value[label_candidates])]
}
label_keep <- head(label_candidates, n_label_target)
# Preserve the androgen-response label in the TRIM11 network even when the
# dynamic label quota would otherwise drop it.
if (identical(gene_id, "TRIM11")) {
  forced_pathway <- which(node_export$pathway == "HALLMARK_ANDROGEN_RESPONSE")
  if (length(forced_pathway) == 1L && !(forced_pathway %in% label_keep)) {
    label_keep <- c(head(setdiff(label_keep, forced_pathway), max(0L, n_label_target - 1L)), forced_pathway)
  }
}
node_export$node_label <- ""
node_export$node_label[label_keep] <- node_export$label[label_keep]

network_summary$n_labels <- length(label_keep)
network_summary$label_fraction <- length(label_keep) / n_nodes
network_summary$label_candidate_abs_cor_cutoff <- label_candidate_abs_cor_cutoff
network_summary$label_candidate_p_cutoff <- label_candidate_p_cutoff
write_csv(network_summary, file.path(result_dir, paste0(file_prefix, "_network_summary.csv")))

V(graph)$node_size <- node_export$node_size[match(V(graph)$name, node_export$pathway)]
V(graph)$node_label <- node_export$node_label[match(V(graph)$name, node_export$pathway)]
V(graph)$community_name <- node_export$community_name[match(V(graph)$name, node_export$pathway)]

if (ecount(graph) > 0) {
  # Keep the visual range narrow (max-min = 1.2) while retaining the ordering
  # of Jaccard similarities.
  E(graph)$edge_width <- rescale(E(graph)$similarity,
                                 to = c(edge_width_min, edge_width_max))
} else {
  E(graph)$edge_width <- numeric(0)
}

# Freeze the layout once so that ellipses, group labels and network layers use
# exactly the same coordinates.
# Explicitly reduce the force-directed layout area to remove large empty
# margins while retaining deterministic coordinates.
layout_matrix <- igraph::layout_with_fr(
  graph,
  weights = E(graph)$similarity,
  niter = 2000
)
layout_matrix <- igraph::norm_coords(
  layout_matrix, xmin = -1, xmax = 1, ymin = -1, ymax = 1
)
layout_data <- ggraph::create_layout(
  graph,
  layout = "manual",
  x = layout_matrix[, 1],
  y = layout_matrix[, 2]
)
# Keep TRIM11 androgen-response visually adjacent to the late estrogen-
# response node, as requested, while preserving the graph's other coordinates.
if (identical(gene_id, "TRIM11")) {
  ar_i <- which(layout_data$name == "HALLMARK_ANDROGEN_RESPONSE")
  er_i <- which(layout_data$name == "HALLMARK_ESTROGEN_RESPONSE_LATE")
  if (length(ar_i) == 1L && length(er_i) == 1L) {
    target <- c(layout_data$x[er_i] + 0.12, layout_data$y[er_i] + 0.10)
    layout_data$x[ar_i] <- max(-0.95, min(0.95, target[1]))
    layout_data$y[ar_i] <- max(-0.95, min(0.95, target[2]))
  }
}
layout_data <- layout_data %>%
  group_by(community) %>%
  mutate(group_n = n()) %>%
  ungroup()
group_centers <- layout_data %>%
  group_by(community) %>%
  summarise(
    x = mean(x),
    y = mean(y),
    n_nodes = n(),
    group_label = first(community_name),
    .groups = "drop"
  )
# Do not repeat a pathway label as a separate community label; the pathway
# label already identifies that region and the duplicate would add clutter.
group_centers <- group_centers %>%
  filter(!group_label %in% node_export$label[label_keep])

# Only communities with a rendered Group label receive a descriptive name in
# the node export. Numeric community IDs are retained for all nodes.
displayed_communities <- unique(group_centers$community[group_centers$n_nodes >= group_label_min_nodes])
node_export$community_name <- ifelse(
  node_export$community %in% displayed_communities,
  node_export$community_name,
  ""
)
write_csv(node_export, file.path(result_dir, paste0(file_prefix, "_network_nodes.csv")))

# Use one shared repel layer for pathway and community labels so the two label
# classes can avoid one another. Wrapping long pathway names improves the
# 10-cm presentation without changing their identity.
label_data <- bind_rows(
  layout_data %>%
    filter(node_label != "") %>%
    transmute(
      x, y,
      label = stringr::str_wrap(node_label, width = 22),
      label_type = "pathway"
    ),
    group_centers %>%
    filter(n_nodes >= group_label_min_nodes) %>%
    transmute(
      x, y, community,
      label = stringr::str_wrap(group_label, width = 22),
      label_type = "community"
    )
)

p_network <- ggraph(
  graph,
  layout = "manual",
  x = layout_data$x,
  y = layout_data$y
) +
  # Ellipses are drawn only for communities with at least three nodes; this
  # keeps isolated/singleton pathways visible without invalid hull geometry.
  ggforce::geom_mark_ellipse(
    data = layout_data %>% filter(group_n >= 3),
    aes(x = x, y = y, group = community, fill = community),
    inherit.aes = FALSE,
    alpha = 0.15,
    colour = NA,
    expand = unit(4, "mm"),
    show.legend = FALSE
  ) +
  geom_edge_link(
    aes(width = edge_width, alpha = similarity),
    colour = "grey55",
    show.legend = FALSE
  ) +
  geom_node_point(
    aes(size = neg_log10_fdr, colour = rho),
    alpha = 0.95
  ) +
  ggrepel::geom_text_repel(
    data = label_data %>% filter(label_type == "pathway"),
    aes(x = x, y = y, label = label),
    inherit.aes = FALSE,
    size = figure_text_pt / ggplot2::.pt,
    family = "sans",
    max.overlaps = Inf,
    box.padding = 1.0,
    point.padding = 0.8,
    force = 10,
    force_pull = 0.03,
    direction = "both",
    seed = 20260811,
    max.time = 5,
    max.iter = 50000,
    min.segment.length = 0,
    segment.colour = "black",
    segment.alpha = 1
  ) +
  ggrepel::geom_label_repel(
    data = label_data %>% filter(label_type == "community"),
    aes(x = x, y = y, label = label, fill = community),
    inherit.aes = FALSE,
    size = figure_text_pt / ggplot2::.pt,
    family = "sans",
    fontface = "bold",
    max.overlaps = Inf,
    box.padding = 1.2,
    point.padding = 0.8,
    force = 10,
    force_pull = 0.03,
    direction = "both",
    seed = 20260811,
    max.time = 5,
    max.iter = 50000,
    min.segment.length = 0,
    segment.colour = "black",
    segment.alpha = 1,
    label.padding = unit(0.08, "lines"),
    label.r = unit(0.05, "lines"),
    label.size = 0,
    show.legend = FALSE
  ) +
  scale_colour_gradient2(
    low = "#3953A4",
    mid = "white",
    high = "#B50404",
    midpoint = 0,
    name = aws_label
  ) +
  scale_size_continuous(
    range = c(node_size_min, node_size_max),
    name = minus_log_fdr_label,
  ) +
  theme_void(base_size = figure_text_pt) +
  theme(
    plot.background = element_rect(fill = "white", colour = NA),
    panel.background = element_rect(fill = "white", colour = NA),
    legend.position = "right",
    legend.title = element_text(face = "bold"),
    plot.margin = margin(8, 16, 8, 8),
    plot.title = element_text(hjust = 0.5, face = "bold", size = figure_text_pt),
    plot.subtitle = element_text(hjust = 0.5, size = figure_text_pt)
  ) +
  coord_cartesian(clip = "off") +
  labs(
    title = paste0(gene_id, if (standard_pdf) " AWSI-" else " AWSI–", category_label, " pathway network"),
  subtitle = paste0(
      if (standard_pdf) "FDR<=" else "FDR≤", fdr_cutoff,
      "; Jaccard", if (standard_pdf) ">=" else "≥", jaccard_cutoff,
      "; labels ", round(100 * length(label_keep) / n_nodes), "%"
    )
  )

ggsave(
  filename = file.path(
    result_dir,
    paste0(file_prefix, "_pathway_network", Sys.getenv("PDF_SUFFIX", ""), ".pdf")
  ),
  plot = p_network,
  width = figure_size_cm / 2.54,
  height = figure_size_cm / 2.54,
  device = function(filename, width, height, ...) {
    grDevices::pdf(
      file = filename,
      width = width,
      height = height,
      family = "Helvetica",
      useDingbats = FALSE,
      ...
    )
  }
)
ggsave(
  filename = file.path(result_dir, paste0(file_prefix, "_pathway_network.png")),
  plot = p_network,
  width = figure_size_cm / 2.54,
  height = figure_size_cm / 2.54,
  dpi = 300
)

# -------------------------------------------------------------------------
# 8. Native enrichplot::emapplot using a synthetic enrichResult object
# -------------------------------------------------------------------------
# This does not rerun over-representation analysis. It wraps the already
# selected S9B pathways, their original gene sets, AWSI correlations and FDRs
# in the class expected by enrichplot. Consequently pairwise_termsim(method=
# "JC") and emapplot() can provide the same k-means grouping/ellipse logic as
# the PTM_set1_network_BP.R workflow, while retaining the S9B node selection.

similarity_matrix <- matrix(
  0,
  nrow = length(gene_sets),
  ncol = length(gene_sets),
  dimnames = list(names(gene_sets), names(gene_sets))
)
if (length(gene_sets) >= 2) {
  for (i in seq_len(length(gene_sets) - 1L)) {
    for (j in seq.int(i + 1L, length(gene_sets))) {
      similarity_matrix[i, j] <- similarity_matrix[j, i] <-
        pair_similarity(gene_sets[[i]], gene_sets[[j]], edge_metric)[["similarity"]]
    }
  }
}
saveRDS(similarity_matrix,
        file.path(result_dir, paste0(file_prefix, "_pairwise_Jaccard_similarity.rds")))

if (requireNamespace("clusterProfiler", quietly = TRUE) &&
    requireNamespace("enrichplot", quietly = TRUE)) {
  suppressPackageStartupMessages({
    library(clusterProfiler)
    library(enrichplot)
  })

  # enrichplot::geneInCategory() reads result$geneID, so preserve every
  # pathway's exact gene set as a slash-delimited SYMBOL string.
  emap_result <- node_table %>%
    transmute(
      ID = pathway,
      Description = label,
      GeneRatio = paste0(gene_set_size, "/", max(gene_set_size)),
      BgRatio = paste0(gene_set_size, "/", max(gene_set_size)),
      RichFactor = gene_set_size / max(gene_set_size),
      FoldEnrichment = 1,
      zScore = rho,
      pvalue = fdr,
      p.adjust = fdr,
      qvalue = fdr,
      geneID = vapply(gene_sets[pathway], paste, collapse = "/", FUN.VALUE = character(1)),
      Count = gene_set_size,
      rho = rho,
      neg_log10_fdr = neg_log10_fdr
    )
  # enrichplot's JC implementation indexes gene sets by result$ID through
  # rownames(result); clusterProfiler objects conventionally use IDs here.
  # Setting these explicitly is essential when ID and the display Description
  # (the shortened Hallmark label) are different strings.
  emap_result <- as.data.frame(emap_result, stringsAsFactors = FALSE)
  rownames(emap_result) <- emap_result$ID

  emap_obj <- methods::new(
    "enrichResult",
    result = emap_result,
    pvalueCutoff = fdr_cutoff,
    pAdjustMethod = "BH",
    qvalueCutoff = fdr_cutoff,
    organism = "Homo sapiens",
    ontology = "HALLMARK",
    gene = unique(unlist(gene_sets, use.names = FALSE)),
    keytype = "SYMBOL",
    universe = unique(unlist(gene_sets, use.names = FALSE)),
    gene2Symbol = character(),
    geneSets = gene_sets,
    readable = TRUE,
    termsim = similarity_matrix,
    method = "JC",
    dr = list()
  )
  emap_obj <- pairwise_termsim(
    emap_obj,
    method = "JC",
    showCategory = nrow(emap_result)
  )

  emap_dir <- file.path(result_dir, "emapplot")
  dir.create(emap_dir, recursive = TRUE, showWarnings = FALSE)
  emap_ncluster <- max(1L, round(sqrt(nrow(emap_result)) / 2))
  set.seed(20260811)
  p_emap <- emapplot(
    emap_obj,
    showCategory = nrow(emap_result),
    node_label = "group",
    nCluster = emap_ncluster,
    layout = "fr",
    color = "rho",
    min_edge = jaccard_cutoff,
    color_edge = "grey70",
    size_edge = 0.5
  ) +
    # Match the custom ggraph plot: negative rho = blue, positive rho = red.
    # enrichplot's default colour scale is reversed relative to this convention.
    scale_colour_gradient2(
      low = "#3953A4",
      mid = "white",
      high = "#B50404",
      midpoint = 0,
      name = aws_label
    ) +
    labs(
      title = paste0(gene_id, if (standard_pdf) " AWSI-" else " AWSI–", category_label, " enrichment map"),
      subtitle = paste0(
        "Jaccard similarity ", if (standard_pdf) ">= " else "≥ ", jaccard_cutoff,
        "; k-means groups = ", emap_ncluster,
        "; node colour = ", if (standard_pdf) "AWSI-pathway Spearman rho" else "AWSI–pathway Spearman ρ"
      )
    ) +
    theme(
      plot.background = element_rect(fill = "white", colour = NA),
      panel.background = element_rect(fill = "white", colour = NA),
      plot.margin = margin(8, 16, 8, 8)
    )

  ggsave(
    file.path(emap_dir, paste0(file_prefix, "_emapplot_cluster.pdf")),
    p_emap, width = 12, height = 10, device = cairo_pdf
  )
  ggsave(
    file.path(emap_dir, paste0(file_prefix, "_emapplot_cluster.png")),
    p_emap, width = 12, height = 10, dpi = 300
  )
  write_csv(as.data.frame(emap_obj),
            file.path(emap_dir, paste0(file_prefix, "_emapplot_result.csv")))
  write_csv(p_emap$data,
            file.path(emap_dir, paste0(file_prefix, "_emapplot_node_data.csv")))
  saveRDS(emap_obj,
          file.path(emap_dir, paste0(file_prefix, "_enrichResult.rds")))
  writeLines(capture.output(sessionInfo()),
             file.path(emap_dir, "sessionInfo.txt"))
} else {
  warning("clusterProfiler/enrichplot not installed; native emapplot was skipped.")
}

message("Network complete. Results written to: ", normalizePath(result_dir))
```
