## Supplementary material for "An integrated, transcript-resolution atlas of prostate cancer progression unmasks associations with oncogenic pathways": Reproducible code: audit_primary_survival_variants.R

```
# Public submission copy.
# Analytical statements and parameter values are unchanged from the archived source.
# Legacy non-English comments and console messages were normalized for public release.
suppressPackageStartupMessages({library(fst); library(survival)})
atlas <- "__PCATLAS_DOCUMENTS_ROOT__/Data_Harmanization"
meta <- read.csv(file.path(atlas, "filtered_samples.csv"), check.names = FALSE)
clin <- read.csv("__PCATLAS_DOCUMENTS_ROOT__/WebToolStudy/combined_all_survival_updated_v5.csv", check.names = FALSE)
info <- read.csv(file.path(atlas, "transcripts_info.csv"))
tx <- read.fst(file.path(atlas, "txi.counts.fst"))
mat <- as.matrix(tx)
rownames(mat) <- sub("\\..*$", "", info$transcript_id)
sample_ids <- sub("^id\\.", "", colnames(mat))
lib_colsum <- colSums(mat)
lib_meta <- meta$totalcounts[match(sample_ids, meta$id)]
targets <- c("ENST00000584437", "ENST00000583753", "ENST00000357654", "ENST00000352993")

logrank <- function(d, group) {
  if (length(unique(group)) < 2) return(NA_real_)
  fit <- survdiff(Surv(PFS_Time, PFS_Event) ~ group, data = d)
  pchisq(fit$chisq, length(fit$n)-1, lower.tail=FALSE)
}

out <- list(); k <- 0
for (id in targets) {
  raw <- as.numeric(mat[id, ])
  for (norm in c("colsum", "metadata_totalcounts", "raw")) {
    expr <- switch(norm, colsum=log2(raw/(lib_colsum/1e6)+1), metadata_totalcounts=log2(raw/(lib_meta/1e6)+1), raw=log2(raw+1))
    dat <- merge(clin, data.frame(id=sample_ids, Expression=expr), by="id")
    primary <- dat[dat[["Molecular Type"]] == "PRIMARY" & !is.na(dat$PFS_Time) & dat$PFS_Time > 0 & !is.na(dat$PFS_Event),]
    for (scope in c("global","primary")) {
      ref <- if(scope=="global") dat$Expression else primary$Expression
      for (cut in c("median","q25q75","tertiles")) {
        d <- primary
        if(cut=="median") {
          z <- median(ref,na.rm=TRUE); g <- ifelse(d$Expression>=z,"High","Low")
        } else if(cut=="q25q75") {
          q <- quantile(ref,c(.25,.75),na.rm=TRUE); d <- d[d$Expression<=q[1] | d$Expression>=q[2],]; g <- ifelse(d$Expression>=q[2],"High","Low")
        } else {
          q <- quantile(ref,c(1/3,2/3),na.rm=TRUE); d <- d[d$Expression<=q[1] | d$Expression>=q[2],]; g <- ifelse(d$Expression>=q[2],"High","Low")
        }
        k<-k+1; out[[k]]<-data.frame(ENST=id,norm=norm,scope=scope,cut=cut,n=nrow(d),events=sum(d$PFS_Event==1),p=logrank(d,g))
      }
    }
  }
}
out <- do.call(rbind,out)
write.csv(out,"__PCATLAS_PROJECT_ROOT__/reproduced_supplementary_tables_v2/primary_survival_variant_audit.csv",row.names=FALSE)
print(out)
```
