## Supplementary material for "An integrated, transcript-resolution atlas of prostate cancer progression unmasks associations with oncogenic pathways": Reproducible code: survival_analysis_v4_all_combinations.R

```
# Public submission copy.
# Analytical statements and parameter values are unchanged from the archived source.
# Legacy non-English comments and console messages were normalized for public release.
library(fst)
library(pheatmap)
library(msigdbr)
library(survival)
library(survminer)
library(dplyr)
library(tidyr)

txi_df <- read.fst("./txi.counts.fst")
txi_info <- read.csv("./transcripts_info.csv")
rownames(txi_df) <- txi_info$transcript_id
txi_counts <- as.matrix(txi_df)

ncount.tx <- log2(sweep(x = txi_counts, MARGIN = 2, STATS = colSums(txi_counts)/1000000, FUN = "/") + 1)
rownames(ncount.tx) <- gsub("\\..*$", "", rownames(ncount.tx))

combined_all = read.csv("./../Atlas_metada_cleaning/combined_all_survival_updated_v5.csv",check.names = FALSE)
combined_all$id <- as.character(combined_all$id)
metadata_clean <- combined_all

matrix_sample_ids <- as.character(gsub("^id\\.", "", colnames(ncount.tx)))


target_list <- data.frame(
  Gene = c("WNT2B", "WNT2B", "TRIM11", "TRIM11", "ROCK2", "ROCK2",
           "RNF43", "RNF43", "CDK12", "CDK12", "F8", "F8",
           "RELA", "RELA", "BRCA1", "BRCA1"),
  UniProt = c("Q93097-1", "Q93097-2", "Q96F44-1", "R4GMV1", "O75116", "E9PF63",
              "Q68DV7-1", "Q68DV7-2", "Q9NYV4-1", "J3QSD7", "P00451-1", "P00451-2",
              "Q04206-1", "A0A087X0W8", "P38398-1", "P38398-5"),
  ENST = c("ENST00000369684", "ENST00000369686", "ENST00000284551", "ENST00000493030",
           "ENST00000315872", "ENST00000401753", "ENST00000584437", "ENST00000583753",
           "ENST00000447079", "ENST00000584632", "ENST00000360256", "ENST00000330287",
           "ENST00000406246", "ENST00000612991", "ENST00000357654", "ENST00000352993"),
  stringsAsFactors = FALSE
)

base_out_dir <- "./Survival_Plots_Comprehensive"
dir_single <- file.path(base_out_dir, "1_Single_Isoform")
dir_combo2 <- file.path(base_out_dir, "2_Homodimer_2Groups")
dir_combo4 <- file.path(base_out_dir, "3_Homodimer_4Groups")

for(d in c(base_out_dir, dir_single, dir_combo2, dir_combo4)) {
  if(!dir.exists(d)) dir.create(d)
}

endpoints <- list(
  PFS = list(time = "PFS_Time", event = "PFS_Event", ylab = "Progression-Free Survival"),
  OS  = list(time = "OS_Time",  event = "OS_Event",  ylab = "Overall Survival")
)
cohorts <- list(
  ALL     = "All Patients",
  PRIMARY = "PRIMARY Group",
  CRPC    = "CRPC Group"
)

palette_1g <- c("Low" = "#2E9FDF", "High" = "#E7B800")
palette_4g_named <- c(
  "High / High" = "#E64B35",
  "High / Low"  = "#4DBBD5",
  "Low / High"  = "#00A087",
  "Low / Low"   = "#8491B4"
)


cat("\n=== 1. Starting Single Isoform Survival Analysis ===\n")

for (i in 1:nrow(target_list)) {
  gene_name <- target_list$Gene[i]
  isoform_id <- target_list$ENST[i]
  cat(sprintf("\n[%d/%d] Processing %s (%s)...\n", i, nrow(target_list), gene_name, isoform_id))

  if(!isoform_id %in% rownames(ncount.tx)) next

  isoform_expr <- data.frame(Sample_ID = matrix_sample_ids, Expression = ncount.tx[isoform_id, ], stringsAsFactors = FALSE)

  surv_data_base <- metadata_clean %>%
    left_join(isoform_expr, by = c("id" = "Sample_ID"))

  for (ep_name in names(endpoints)) {
    for (ch_name in names(cohorts)) {
      ep <- endpoints[[ep_name]]
      plot_data <- surv_data_base

      if (ch_name == "PRIMARY") plot_data <- plot_data %>% filter(`Molecular Type` == "PRIMARY")
      if (ch_name == "CRPC")    plot_data <- plot_data %>% filter(`Molecular Type` %in% c("mARPC", "mDNPC", "mNEPC"))

      plot_data$TIME <- plot_data[[ep$time]]
      plot_data$EVENT <- plot_data[[ep$event]]
      plot_data <- plot_data[!is.na(plot_data$TIME) & plot_data$TIME > 0 & !is.na(plot_data$Expression), ]

      if (nrow(plot_data) > 0) {
        plot_data <- plot_data %>%
          mutate(
            Med_Expr = median(Expression, na.rm = TRUE),
            Expr_Group = factor(case_when(
              Med_Expr == 0 & Expression > 0  ~ "High",
              Med_Expr == 0 & Expression == 0 ~ "Low",
              Med_Expr > 0  & Expression >= Med_Expr ~ "High",
              TRUE ~ "Low"
            ), levels = c("Low", "High"))
          )
      }

      plot_data$Expr_Group <- droplevels(plot_data$Expr_Group)
      actual_labs <- levels(plot_data$Expr_Group)

      if(length(actual_labs) >= 2 && nrow(plot_data) > 10) {
        fit_model <- survfit(Surv(TIME, EVENT) ~ Expr_Group, data = plot_data)

        p <- ggsurvplot(
          fit_model, data = plot_data,
          pval = TRUE, conf.int = TRUE, risk.table = TRUE,
          title = paste0(gene_name, " (", isoform_id, ")\n", ep$ylab, " in ", cohorts[[ch_name]]),
          xlab = "Time (Days)", ylab = ep$ylab,
          palette = unname(palette_1g[actual_labs]),
          legend.labs = actual_labs
        )

        pdf_path <- file.path(dir_single, paste0(gene_name, "_", isoform_id, "_", ch_name, "_", ep_name, ".pdf"))
        pdf(file = pdf_path, width = 7, height = 6, onefile = FALSE)
        print(p)
        dev.off()
      }
    }
  }
}


cat("\n=== 2. Starting Homodimer 2-Group Analysis (HiHi vs LoHi) ===\n")
homodimer_genes <- c("ROCK2", "TRIM11", "RELA","BRCA1","WNT2B", "TRIM11")

for (g in homodimer_genes) {
  cat(sprintf("\nProcessing 2-Group Combo for: %s ...\n", g))
  isos <- target_list$ENST[target_list$Gene == g]
  if(length(isos) != 2) next
  isoA <- isos[1]; isoB <- isos[2]
  if(!isoA %in% rownames(ncount.tx) | !isoB %in% rownames(ncount.tx)) next

  combo_expr <- data.frame(Sample_ID = matrix_sample_ids, Expr_A = ncount.tx[isoA, ], Expr_B = ncount.tx[isoB, ], stringsAsFactors = FALSE)

  level_HiHi <- paste0(isoA, "(Hi)_", isoB, "(Hi)")
  level_LoHi <- paste0(isoA, "(Lo)_", isoB, "(Hi)")
  palette_2g_named <- setNames(c("#E64B35", "#4DBBD5"), c(level_HiHi, level_LoHi))

  surv_combo_base <- metadata_clean %>%
    left_join(combo_expr, by = c("id" = "Sample_ID"))

  for (ep_name in names(endpoints)) {
    for (ch_name in names(cohorts)) {
      ep <- endpoints[[ep_name]]
      plot_data <- surv_combo_base

      if (ch_name == "PRIMARY") plot_data <- plot_data %>% filter(`Molecular Type` == "PRIMARY")
      if (ch_name == "CRPC")    plot_data <- plot_data %>% filter(`Molecular Type` %in% c("mARPC", "mDNPC", "mNEPC"))

      plot_data$TIME <- plot_data[[ep$time]]
      plot_data$EVENT <- plot_data[[ep$event]]
      plot_data <- plot_data[!is.na(plot_data$TIME) & plot_data$TIME > 0 & !is.na(plot_data$Expr_A) & !is.na(plot_data$Expr_B), ]

      if(nrow(plot_data) > 0) {
        plot_data <- plot_data %>%
          mutate(
            Med_A = median(Expr_A, na.rm = TRUE), Med_B = median(Expr_B, na.rm = TRUE),
            Status_A = case_when(Med_A == 0 & Expr_A > 0 ~ "High", Med_A == 0 & Expr_A == 0 ~ "Low", Med_A > 0 & Expr_A >= Med_A ~ "High", TRUE ~ "Low"),
            Status_B = case_when(Med_B == 0 & Expr_B > 0 ~ "High", Med_B == 0 & Expr_B == 0 ~ "Low", Med_B > 0 & Expr_B >= Med_B ~ "High", TRUE ~ "Low")
          ) %>%
          mutate(
            Combo_Group = factor(case_when(
              Status_A == "High" & Status_B == "High" ~ level_HiHi,
              Status_A == "Low"  & Status_B == "High" ~ level_LoHi,
              TRUE ~ "Other"
            ), levels = c(level_HiHi, level_LoHi, "Other"))
          ) %>%
          filter(Combo_Group != "Other")
      }

      plot_data$Combo_Group <- droplevels(plot_data$Combo_Group)
      actual_labs <- levels(plot_data$Combo_Group)

      if(length(actual_labs) >= 2 && nrow(plot_data) > 10) {
        fit_model <- survfit(Surv(TIME, EVENT) ~ Combo_Group, data = plot_data)

        p <- ggsurvplot(
          fit_model, data = plot_data,
          pval = TRUE, conf.int = TRUE, risk.table = TRUE,
          title = paste0(g, " Isoform Competition\n", ep$ylab, " in ", cohorts[[ch_name]]),
          xlab = "Time (Days)", ylab = ep$ylab,
          palette = unname(palette_2g_named[actual_labs]),
          legend.title = "Combination", legend.labs = actual_labs
        )

        pdf_path <- file.path(dir_combo2, paste0(g, "_HiHi_vs_LoHi_", ch_name, "_", ep_name, ".pdf"))
        pdf(file = pdf_path, width = 8, height = 6.5, onefile = FALSE)
        print(p)
        dev.off()
      }
    }
  }
}


cat("\n=== 3. Starting Homodimer 4-Group Analysis ===\n")

for (g in homodimer_genes) {
  cat(sprintf("\nProcessing 4-Group Combo for: %s ...\n", g))
  isos <- target_list$ENST[target_list$Gene == g]
  if(length(isos) != 2) next
  isoA <- isos[1]; isoB <- isos[2]
  if(!isoA %in% rownames(ncount.tx) | !isoB %in% rownames(ncount.tx)) next

  combo_expr <- data.frame(Sample_ID = matrix_sample_ids, Expr_A = ncount.tx[isoA, ], Expr_B = ncount.tx[isoB, ], stringsAsFactors = FALSE)

  surv_4g_base <- metadata_clean %>%
    left_join(combo_expr, by = c("id" = "Sample_ID"))

  for (ep_name in names(endpoints)) {
    for (ch_name in names(cohorts)) {
      ep <- endpoints[[ep_name]]
      plot_data <- surv_4g_base

      if (ch_name == "PRIMARY") plot_data <- plot_data %>% filter(`Molecular Type` == "PRIMARY")
      if (ch_name == "CRPC")    plot_data <- plot_data %>% filter(`Molecular Type` %in% c("mARPC", "mDNPC", "mNEPC"))

      plot_data$TIME <- plot_data[[ep$time]]
      plot_data$EVENT <- plot_data[[ep$event]]
      plot_data <- plot_data[!is.na(plot_data$TIME) & plot_data$TIME > 0 & !is.na(plot_data$Expr_A) & !is.na(plot_data$Expr_B), ]

      if(nrow(plot_data) > 0) {
        plot_data <- plot_data %>%
          mutate(
            Med_A = median(Expr_A, na.rm = TRUE), Med_B = median(Expr_B, na.rm = TRUE),
            Status_A = case_when(Med_A == 0 & Expr_A > 0 ~ "High", Med_A == 0 & Expr_A == 0 ~ "Low", Med_A > 0 & Expr_A >= Med_A ~ "High", TRUE ~ "Low"),
            Status_B = case_when(Med_B == 0 & Expr_B > 0 ~ "High", Med_B == 0 & Expr_B == 0 ~ "Low", Med_B > 0 & Expr_B >= Med_B ~ "High", TRUE ~ "Low"),
            Group_4 = factor(case_when(
              Status_A == "High" & Status_B == "High" ~ "High / High",
              Status_A == "High" & Status_B == "Low"  ~ "High / Low",
              Status_A == "Low"  & Status_B == "High" ~ "Low / High",
              Status_A == "Low"  & Status_B == "Low"  ~ "Low / Low"
            ), levels = c("High / High", "High / Low", "Low / High", "Low / Low"))
          )
      }

      plot_data$Group_4 <- droplevels(plot_data$Group_4)
      actual_labs <- levels(plot_data$Group_4)

      if(length(actual_labs) > 1 && nrow(plot_data) > 10) {
        fit_model <- survfit(Surv(TIME, EVENT) ~ Group_4, data = plot_data)

        p <- ggsurvplot(
          fit_model, data = plot_data,
          pval = TRUE, conf.int = FALSE, risk.table = TRUE, risk.table.height = 0.3,
          title = paste0(g, " 4-Group Combination\n", ep$ylab, " in ", cohorts[[ch_name]]),
          xlab = "Time (Days)", ylab = ep$ylab,
          palette = unname(palette_4g_named[actual_labs]),
          legend.title = paste0(isoA, " / ", isoB), legend.labs = actual_labs
        )

        pdf_path <- file.path(dir_combo4, paste0(g, "_4Group_", ch_name, "_", ep_name, ".pdf"))
        pdf(file = pdf_path, width = 8.5, height = 7.5, onefile = FALSE)
        print(p)
        dev.off()
      }
    }
  }
}

cat("\n✅ All comprehensive 6-combination survival analyses finished! Check 'Survival_Plots_Comprehensive' folder.\n")
```
