## Supplementary material for "An integrated, transcript-resolution atlas of prostate cancer progression unmasks associations with oncogenic pathways": Reproducible code: generate_ssgsea_data.py

```
# Public submission copy.
# Analytical statements and parameter values are unchanged from the archived source.
# Legacy non-English comments and console messages were normalized for public release.
import pandas as pd
import numpy as np
import os

out_dir = 'Integrated_ssGSEA_Results'
os.makedirs(out_dir, exist_ok=True)

pool1_samples = {
    'PCNA_R1': 'FL.BioSample_1', 'PCNA_R2': 'FL.BioSample_2',
    'LuCaP-147_R1': 'FL.BioSample_3', 'LuCaP-147_R2': 'FL.BioSample_4',
    'LuCaP-35_R1': 'FL.BioSample_5', 'LuCaP-35_R2': 'FL.BioSample_6',
    'LuCaP-23_R1': 'FL.BioSample_7', 'LuCaP-23_R2': 'FL.BioSample_8',
    'LuCaP-145_R1': 'FL.BioSample_9', 'LuCaP-145_R2': 'FL.BioSample_10'
}

pool2_samples = {
    'LNCaP_R1': 'FL.BioSample_1', 'LNCaP_R2': 'FL.BioSample_2',
    'VCAP_R1': 'FL.BioSample_3', 'VCAP_R2': 'FL.BioSample_4',
    '22RV1_R1': 'FL.BioSample_5', '22RV1_R2': 'FL.BioSample_6',
    'PC3_R1': 'FL.BioSample_7', 'PC3_R2': 'FL.BioSample_8',
    'H660_R1': 'FL.BioSample_9', 'H660_R2': 'FL.BioSample_10'
}

def load_and_collapse_gene(pigeon_path, sample_map):
    usecols = ['associated_gene', 'structural_category', 'associated_transcript'] + list(sample_map.values())
    df = pd.read_csv(pigeon_path, sep='\t', usecols=usecols)
    # calculate global totals (open universe)
    totals = {name: df[col].sum() for name, col in sample_map.items()}

    # filter to FSM and ISM, drop novel
    df = df[df['structural_category'].isin(['full-splice_match', 'incomplete-splice_match'])]
    df = df[df['associated_transcript'] != 'novel']
    df = df.dropna(subset=['associated_transcript', 'associated_gene'])

    # Gene level
    # sum counts per gene
    df_counts = df[['associated_gene'] + list(sample_map.values())].groupby('associated_gene').sum().reset_index()
    # rename columns to sample names
    rename_dict = {col: name for name, col in sample_map.items()}
    df_counts = df_counts.rename(columns=rename_dict)

    # CPM
    cpm = df_counts.copy()
    for name in sample_map.keys():
        cpm[name] = (cpm[name] / totals[name]) * 1e6

    return cpm

print("Loading Pool 1...")
p1_cpm = load_and_collapse_gene('__PCATLAS_DOCUMENTS_ROOT__/long_reads_results/pool1_merged/pigeon.classification.txt', pool1_samples)
print("Loading Pool 2...")
p2_cpm = load_and_collapse_gene('__PCATLAS_DOCUMENTS_ROOT__/long_reads_results/pool2_merged/pigeon.classification.txt', pool2_samples)

print("Merging 20 samples...")
merged_cpm = pd.merge(p1_cpm, p2_cpm, on='associated_gene', how='outer').fillna(0)

print("Applying log2(CPM+1)...")
# Calculate log2(CPM+1)
sample_cols = list(pool1_samples.keys()) + list(pool2_samples.keys())
merged_cpm[sample_cols] = np.log2(merged_cpm[sample_cols] + 1)

merged_cpm.to_csv(os.path.join(out_dir, 'All_20_Samples_Gene_logCPM.csv'), index=False)
print("Saved All_20_Samples_Gene_logCPM.csv")

# Now prepare Isoform data
print("Preparing Target Isoform Matrix...")
p1_iso = pd.read_csv('__PCATLAS_DOCUMENTS_ROOT__/long_reads_results/Pool1_Isoform_Analysis/Target_Isoforms_Plots_logCPM/Pool1_Specific_14_Isoforms_logCPM.csv')
p2_iso = pd.read_csv('__PCATLAS_DOCUMENTS_ROOT__/long_reads_results/Integrated_Isoform_Analysis/Target_Isoforms_Plots_logCPM/Specific_14_Isoforms_logCPM.csv')

# p2 contains short read too, filter to long read
p2_iso = p2_iso[p2_iso['Platform'] == 'LongRead (MAS-Seq)']

# Combine
all_iso = pd.concat([p1_iso, p2_iso])

# Pivot to wide format: rows = Isoforms, cols = Samples
# We only need log2(CPM+1)
iso_wide = all_iso.pivot(index='Isoform', columns='Sample', values='log2(CPM+1)')
iso_wide.to_csv(os.path.join(out_dir, 'Target_14_Isoforms_logCPM_Wide.csv'))
print("Saved Target_14_Isoforms_logCPM_Wide.csv")
```
