## Supplementary material for "An integrated, transcript-resolution atlas of prostate cancer progression unmasks associations with oncogenic pathways": Reproducible code: plot_best_sample_scatter_publication.R

```
# Public submission copy.
# Analytical statements and parameter values are unchanged from the archived source.
# Legacy non-English comments and console messages were normalized for public release.
library(ggplot2)
library(ggpubr)
library(dplyr)

if (!requireNamespace("ggrastr", quietly = TRUE)) {
  install.packages("ggrastr", repos = "https://cloud.r-project.org", ask = FALSE, update = FALSE)
}
library(ggrastr)

out_dir <- "Publication_Quality_Plots/LNCaP_R1_Scatterplots"
dir.create(out_dir, recursive = TRUE, showWarnings = FALSE)

# 1. Load Data
df <- read.csv("Best_Sample_Scatter_Data_LNCaP_R1.csv")

# Ensure factor levels
conditions <- c(
  "All Genes", "protein_coding (Gene)", "lncRNA (Gene)",
  "All Transcripts", "protein_coding (Tx)", "lncRNA (Tx)"
)
df$Condition <- factor(df$Condition, levels=conditions)

for (cond in conditions) {
  df_cond <- df %>% filter(Condition == cond)

  if (nrow(df_cond) == 0) next

  # Set colors based on condition type (Gene vs Tx)
  # Match the color palette from the combined boxplot if possible, or just use a nice deep blue/purple.
  pt_color <- ifelse(grepl("Gene", cond), "#1f77b4", "#9467bd")

  # Remove points that are exactly zero in both?
  # Let's keep what's strictly in the data.

  p <- ggscatter(df_cond, x="SR_logCPM", y="LR_logCPM", color=pt_color, alpha=0.3, size=0.3,
                 add="reg.line", add.params=list(color="black", linetype="dashed", linewidth=0.3)) +
    stat_cor(method="pearson", label.x.npc="left", label.y.npc="top", size=2) +
    geom_abline(intercept=0, slope=1, linetype="dashed", color="red", linewidth=0.3) +
    labs(title=paste("LNCaP_R1:", cond),
         x="Short-Read log2(CPM + 1)",
         y="Long-Read log2(CPM + 1)") +
    coord_cartesian(xlim=c(0, max(df_cond$SR_logCPM, na.rm=TRUE)), ylim=c(0, max(df_cond$LR_logCPM, na.rm=TRUE))) +
    theme_bw(base_size=5) +
    theme(plot.title=element_text(face="bold", hjust=0.5, size=6),
          axis.text=element_text(size=5),
          axis.title=element_text(size=5),
          plot.margin=margin(t=2, r=2, b=2, l=2, unit="pt"))

  # Rasterize only the scatter points to keep file size small for Illustrator
  p <- rasterize(p, layers="Point", dpi=300)

  # Replace spaces and parentheses for filename
  safe_cond <- gsub(" \\(", "_", cond)
  safe_cond <- gsub("\\)", "", safe_cond)
  safe_cond <- gsub(" ", "_", safe_cond)

  ggsave(file.path(out_dir, paste0("LNCaP_R1_Scatterplot_", safe_cond, ".pdf")), p, width=7, height=5.25, units="cm")
  cat(paste("Saved plot for", cond, "\n"))
}

cat("ALL SCATTERPLOTS DONE!\n")
```
