## Supplementary material for "An integrated, transcript-resolution atlas of prostate cancer progression unmasks associations with oncogenic pathways": Reproducible code: plot_combined_correlation_boxplot_publication.R

```
# Public submission copy.
# Analytical statements and parameter values are unchanged from the archived source.
# Legacy non-English comments and console messages were normalized for public release.
library(ggplot2)
library(ggpubr)
library(dplyr)

out_dir <- "Publication_Quality_Plots"
dir.create(out_dir, showWarnings = FALSE)

# 1. Load Data
gene_all <- read.csv("Gene_Level_Correlation_Results/Gene_Correlation_Results.csv", check.names=FALSE)
gene_all$Group <- "All Genes"
gene_all$Level <- "Gene"

gene_type <- read.csv("Gene_Level_Correlation_ByType_Results/Gene_Correlation_ByType_Results.csv", check.names=FALSE)
gene_type$Group <- paste0(gene_type$GeneType, " (Gene)")
gene_type$Level <- "Gene"

tx_all <- read.csv("Integrated_Isoform_Analysis/Isoform_Correlation_Results.csv", check.names=FALSE)
tx_all$Group <- "All Transcripts"
tx_all$Level <- "Transcript"
colnames(tx_all)[which(names(tx_all) == "Pearson (logCPM)")] <- "Pearson (logCPM)"

tx_type <- read.csv("Transcript_Level_Correlation_ByType_Results/Transcript_Correlation_ByType_Results.csv", check.names=FALSE)
tx_type$Group <- paste0(tx_type$GeneType, " (Tx)")
tx_type$Level <- "Transcript"

# Combine data
df1 <- data.frame(Sample=gene_all$Sample, Pearson=gene_all[["Pearson (logCPM)"]], Group=gene_all$Group, Level=gene_all$Level)
df2 <- data.frame(Sample=gene_type$Sample, Pearson=gene_type[["Pearson (logCPM)"]], Group=gene_type$Group, Level=gene_type$Level)
df3 <- data.frame(Sample=tx_all$Sample, Pearson=tx_all[["Pearson (logCPM)"]], Group=tx_all$Group, Level=tx_all$Level)
df4 <- data.frame(Sample=tx_type$Sample, Pearson=tx_type[["Pearson (logCPM)"]], Group=tx_type$Group, Level=tx_type$Level)

df_combined <- rbind(df1, df2, df3, df4)

# Factorize Group for ordering
df_combined$Group <- factor(df_combined$Group, levels=c(
  "All Genes", "protein_coding (Gene)", "lncRNA (Gene)",
  "All Transcripts", "protein_coding (Tx)", "lncRNA (Tx)"
))

colors <- c(
  "#D32F2F", "#1976D2", "#388E3C",
  "#FBC02D", "#8E24AA", "#F57C00"
)

# 2. Generate Plot
p <- ggboxplot(df_combined, x="Group", y="Pearson", color="Group", palette=colors,
               add="jitter", add.params=list(size=0.3, alpha=0.6), outlier.shape=NA, size=0.3) +
  geom_hline(yintercept = 0, linetype="dashed", color="gray50", linewidth=0.3) +
  labs(title="Pearson Correlation (CPM vs CPM)",
       subtitle="Gene vs Tx | Protein Coding vs lncRNA",
       y="Pearson R (LR vs SR)", x=NULL) +
  theme_bw(base_size=5) +
  theme(legend.position="none",
        plot.title=element_text(face="bold", hjust=0.5, size=6),
        plot.subtitle=element_text(hjust=0.5, size=5),
        axis.text.x=element_text(angle=45, hjust=1, vjust=1, face="bold", size=5),
        axis.text.y=element_text(size=5),
        axis.title.y=element_text(size=5),
        plot.margin=margin(t=2, r=2, b=2, l=2, unit="pt"))

ggsave(file.path(out_dir, "Combined_6_Conditions_Correlation_Boxplot.pdf"), p, width=7.5, height=5.25, units="cm")
cat("Combined Boxplot saved to:", file.path(out_dir, "Combined_6_Conditions_Correlation_Boxplot.pdf"), "\n")
```
