## Supplementary material for "An integrated, transcript-resolution atlas of prostate cancer progression unmasks associations with oncogenic pathways": Reproducible code: plot_target_isoforms_boxplots_publication.R

```
# Public submission copy.
# Analytical statements and parameter values are unchanged from the archived source.
# Legacy non-English comments and console messages were normalized for public release.
library(ggplot2)
library(ggpubr)
library(dplyr)

out_dir <- "Publication_Quality_Plots/Target_Isoforms"
dir.create(out_dir, recursive = TRUE, showWarnings = FALSE)

# 1. Load Data
df <- read.csv("Integrated_Isoform_Analysis/Specific_12_Isoforms_CPM.csv")

# Apply log2(CPM+1) transformation for better visualization
df$logCPM <- log2(df$CPM + 1)

# Define explicit order: Canonical (Full-length) first, Truncated second
target_genes <- c("RELA", "RNF43", "ROCK2", "TRIM11", "WNT2B")
isoform_order <- list(
  RELA = c("ENST00000406246", "ENST00000612991"),
  RNF43 = c("ENST00000584437", "ENST00000583753"),
  ROCK2 = c("ENST00000401753", "ENST00000315872"),
  TRIM11 = c("ENST00000284551", "ENST00000493030"),
  WNT2B = c("ENST00000369686", "ENST00000369684")
)

for (gene in target_genes) {
  df_gene <- df %>% filter(Gene == gene)

  if (nrow(df_gene) == 0) next

  # Set factor levels explicitly: Canonical (left) then Truncated (right)
  ordered_isos <- isoform_order[[gene]]
  df_gene$Isoform <- factor(df_gene$Isoform, levels=ordered_isos)

  p <- ggboxplot(df_gene, x="Isoform", y="logCPM", color="Platform",
                 palette=c("#1f77b4", "#ff7f0e"), # Blue for LR, Orange for SR
                 add="jitter", add.params=list(size=0.3, alpha=0.6),
                 outlier.shape=NA, size=0.3) +
    labs(title=sprintf("%s Isoforms Expression", gene),
         subtitle="Long-Read vs Short-Read (CPM)",
         y="log2(CPM + 1)", x="Isoform ID") +
    theme_bw(base_size=5) +
    theme(legend.position="top",
          legend.title=element_blank(),
          legend.text=element_text(size=5),
          legend.key.size=unit(0.3, "cm"),
          legend.margin=margin(b=-5, unit="pt"),
          plot.title=element_text(face="bold", hjust=0.5, size=6),
          plot.subtitle=element_text(hjust=0.5, size=5),
          axis.text.x=element_text(face="bold", size=5),
          axis.text.y=element_text(size=5),
          axis.title=element_text(size=5),
          plot.margin=margin(t=2, r=2, b=2, l=2, unit="pt"))

  ggsave(file.path(out_dir, sprintf("%s_isoforms_boxplot_publication.pdf", gene)), p, width=7, height=5.25, units="cm")
  cat(sprintf("Generated plot for %s\n", gene))
}

cat("DONE ALL TARGET ISOFORMS!\n")
```
