## Supplementary material for "An integrated, transcript-resolution atlas of prostate cancer progression unmasks associations with oncogenic pathways": Reproducible code: prepare_sr_lr_intersect.py

```
# Public submission copy.
# Analytical statements and parameter values are unchanged from the archived source.
# Legacy non-English comments and console messages were normalized for public release.
import pandas as pd
import numpy as np
import os

out_dir = 'Integrated_ssGSEA_Results/Platform_Comparison_Inputs'
os.makedirs(out_dir, exist_ok=True)

pool2_samples = [
    'LNCaP_R1', 'LNCaP_R2', 'VCAP_R1', 'VCAP_R2',
    '22RV1_R1', '22RV1_R2', 'PC3_R1', 'PC3_R2',
    'H660_R1', 'H660_R2'
]

print("1. Loading Short-Read Gene Matrix...")
sr_df = pd.read_csv('PolyA_Bossi_10s_27July26/salmon.merged.gene_counts.tsv', sep='\t')
sr_df.columns = [col.replace('H8660', 'H660') for col in sr_df.columns]
sr_counts = sr_df[['gene_name'] + pool2_samples].groupby('gene_name').sum().reset_index()

sr_totals = {sample: sr_counts[sample].sum() for sample in pool2_samples}
sr_cpm = sr_counts.copy()
for sample in pool2_samples:
    sr_cpm[sample] = np.log2((sr_cpm[sample] / sr_totals[sample]) * 1e6 + 1)
sr_cpm.rename(columns={'gene_name': 'associated_gene'}, inplace=True)

print("2. Loading Long-Read Gene Matrix for Pool 2...")
lr_df = pd.read_csv('Integrated_ssGSEA_Results/All_20_Samples_Gene_logCPM.csv')
lr_pool2 = lr_df[['associated_gene'] + pool2_samples]

print("3. Intersecting Genes for Fair ssGSEA Background...")
common_genes = set(lr_pool2['associated_gene']).intersection(set(sr_cpm['associated_gene']))
print(f"Total Common Genes: {len(common_genes)}")

lr_intersect = lr_pool2[lr_pool2['associated_gene'].isin(common_genes)]
sr_intersect = sr_cpm[sr_cpm['associated_gene'].isin(common_genes)]

lr_intersect.to_csv(os.path.join(out_dir, 'Pool2_LR_Gene_logCPM_Intersect.csv'), index=False)
sr_intersect.to_csv(os.path.join(out_dir, 'Pool2_SR_Gene_logCPM_Intersect.csv'), index=False)
print("Saved Intersected Gene Matrices.")
```
