## Supplementary material for "An integrated, transcript-resolution atlas of prostate cancer progression unmasks associations with oncogenic pathways": Reproducible code: run_gene_correlation_by_type.py

```
# Public submission copy.
# Analytical statements and parameter values are unchanged from the archived source.
# Legacy non-English comments and console messages were normalized for public release.
import pandas as pd
import numpy as np
import plotly.express as px
from scipy.stats import pearsonr, spearmanr
import os

out_dir = 'Gene_Level_Correlation_ByType_Results'
os.makedirs(out_dir, exist_ok=True)

# 0. Load Gene Classifications
print("Loading Gene Classifications...")
genes_df = pd.read_csv('__USER_DOWNLOADS__/genes.csv')
genes_df = genes_df[['gene_name', 'gene_type']].drop_duplicates(subset=['gene_name'])

# 1. Load Long-Read Gene Matrix (already logCPM)
print("Loading Long-Read Gene Matrix...")
lr_df = pd.read_csv('Integrated_ssGSEA_Results/All_20_Samples_Gene_logCPM.csv')

pool2_samples = [
    'LNCaP_R1', 'LNCaP_R2', 'VCAP_R1', 'VCAP_R2',
    '22RV1_R1', '22RV1_R2', 'PC3_R1', 'PC3_R2',
    'H660_R1', 'H660_R2'
]

lr_df = lr_df[['associated_gene'] + pool2_samples].rename(columns={'associated_gene': 'gene_name'})

# 2. Load Short-Read Gene Matrix
print("Loading Short-Read Gene Matrix...")
sr_df = pd.read_csv('PolyA_Bossi_10s_27July26/salmon.merged.gene_counts.tsv', sep='\t')
sr_df.columns = [col.replace('H8660', 'H660') for col in sr_df.columns]

sr_counts = sr_df[['gene_name'] + pool2_samples]
sr_counts = sr_counts.groupby('gene_name').sum().reset_index()

sr_totals = {sample: sr_counts[sample].sum() for sample in pool2_samples}
sr_cpm = sr_counts.copy()
for sample in pool2_samples:
    sr_cpm[sample] = np.log2((sr_cpm[sample] / sr_totals[sample]) * 1e6 + 1)

# 3. Merge Long-Read, Short-Read and Gene Classifications
print("Merging on common genes...")
common_genes = set(lr_df['gene_name']).intersection(set(sr_cpm['gene_name']))
print(f"Found {len(common_genes)} common genes.")

lr_common = lr_df[lr_df['gene_name'].isin(common_genes)].sort_values('gene_name').reset_index(drop=True)
sr_common = sr_cpm[sr_cpm['gene_name'].isin(common_genes)].sort_values('gene_name').reset_index(drop=True)

assert all(lr_common['gene_name'] == sr_common['gene_name']), "Genes are not aligned!"

merged_df = pd.DataFrame({'gene_name': lr_common['gene_name']})
for sample in pool2_samples:
    merged_df[f'{sample}_LR'] = lr_common[sample]
    merged_df[f'{sample}_SR'] = sr_common[sample]

merged_df = merged_df.merge(genes_df, on='gene_name', how='inner')

target_types = ['protein_coding', 'lncRNA']
merged_df = merged_df[merged_df['gene_type'].isin(target_types)]

# 4. Calculate Correlations and Plot
results = []
for sample in pool2_samples:
    for gtype in target_types:
        subset = merged_df[merged_df['gene_type'] == gtype]
        x = subset[f'{sample}_SR'].values
        y = subset[f'{sample}_LR'].values

        if len(x) > 1 and x.std() > 0 and y.std() > 0:
            pearson_r, p_pearson = pearsonr(x, y)
            spearman_r, p_spearman = spearmanr(x, y)
        else:
            pearson_r, p_pearson = np.nan, np.nan
            spearman_r, p_spearman = np.nan, np.nan

        results.append({
            'Sample': sample,
            'GeneType': gtype,
            'Pearson (logCPM)': pearson_r, 'Pearson_P': p_pearson,
            'Spearman (logCPM)': spearman_r, 'Spearman_P': p_spearman
        })

        plot_df = pd.DataFrame({
            'Gene': subset['gene_name'],
            'ShortRead_logCPM': x,
            'LongRead_logCPM': y
        })

        fig = px.scatter(plot_df, x='ShortRead_logCPM', y='LongRead_logCPM',
                         hover_name='Gene',
                         title=f'Gene Correlation: LongRead vs ShortRead ({sample}) - {gtype}',
                         labels={'ShortRead_logCPM': 'Short-Read log2(CPM+1)', 'LongRead_logCPM': 'Long-Read log2(CPM+1)'},
                         opacity=0.3)
        fig.add_shape(type='line', line=dict(dash='dash', color='red'), x0=0, y0=0, x1=15, y1=15)

        if not np.isnan(pearson_r):
            if p_pearson < 1e-100:
                annot_text = f"Pearson: {pearson_r:.3f} (p < 1e-100)<br>Spearman: {spearman_r:.3f} (p < 1e-100)"
            else:
                annot_text = f"Pearson: {pearson_r:.3f} (p={p_pearson:.1e})<br>Spearman: {spearman_r:.3f} (p={p_spearman:.1e})"

            fig.add_annotation(
                x=0.05, y=0.95, xref="paper", yref="paper",
                text=annot_text, showarrow=False, align="left",
                bgcolor="rgba(255,255,255,0.8)", bordercolor="black", borderwidth=1
            )
        fig.write_html(os.path.join(out_dir, f'{sample}_{gtype}_Scatter.html'))

# 5. Save results and Summary Boxplot
res_df = pd.DataFrame(results)
res_df.to_csv(os.path.join(out_dir, 'Gene_Correlation_ByType_Results.csv'), index=False)
print(res_df)

melted_df = res_df.melt(id_vars=['Sample', 'GeneType'], value_vars=['Pearson (logCPM)', 'Spearman (logCPM)'],
                        var_name='Metric', value_name='Correlation')
fig = px.box(melted_df, x='Metric', y='Correlation', color='GeneType', points='all',
             hover_data=['Sample'],
             title='Gene-Level Correlation Distribution across 10 Samples (Pool2) by Biotype',
             labels={'Correlation': 'Correlation Coefficient', 'Metric': 'Metric Type'})
fig.update_layout(template='plotly_white')
fig.write_html(os.path.join(out_dir, 'Gene_Correlation_Summary_Boxplot_ByType.html'))

print("DONE!")
```
