## Supplementary material for "An integrated, transcript-resolution atlas of prostate cancer progression unmasks associations with oncogenic pathways": Reproducible code: run_integrated_analysis.py

```
# Public submission copy.
# Analytical statements and parameter values are unchanged from the archived source.
# Legacy non-English comments and console messages were normalized for public release.
import pandas as pd
import numpy as np
import plotly.express as px
from scipy.stats import pearsonr, spearmanr
import os
import time

out_dir = 'Integrated_Isoform_Analysis'
os.makedirs(out_dir, exist_ok=True)

print("1. Loading Pigeon Classification data (pool2_merged)...")
start_time = time.time()
usecols = ['isoform', 'structural_category', 'associated_transcript'] + [f'FL.BioSample_{i}' for i in range(1, 11)]
# Load pool2_merged
lr_df = pd.read_csv('__PCATLAS_DOCUMENTS_ROOT__/long_reads_results/pool2_merged/pigeon.classification.txt', sep='\t', usecols=usecols)
print(f"Loaded {len(lr_df)} rows in {time.time() - start_time:.2f} seconds.")

# Map sample names for pool2
sample_map = {
    'LNCaP_R1': 'FL.BioSample_1', 'LNCaP_R2': 'FL.BioSample_2',
    'VCAP_R1': 'FL.BioSample_3', 'VCAP_R2': 'FL.BioSample_4',
    '22RV1_R1': 'FL.BioSample_5', '22RV1_R2': 'FL.BioSample_6',
    'PC3_R1': 'FL.BioSample_7', 'PC3_R2': 'FL.BioSample_8',
    'H660_R1': 'FL.BioSample_9', 'H660_R2': 'FL.BioSample_10'
}

# Calculate Global Long-Read Denominators
lr_totals = {sample: lr_df[col].sum() for sample, col in sample_map.items()}

# Filter to known transcripts
lr_df = lr_df[lr_df['structural_category'].isin(['full-splice_match', 'incomplete-splice_match'])]
lr_df = lr_df[lr_df['associated_transcript'] != 'novel']
lr_df = lr_df.dropna(subset=['associated_transcript'])

for sample, col in sample_map.items():
    lr_df[sample] = lr_df[col]

lr_counts = lr_df[['associated_transcript'] + list(sample_map.keys())]

print("3. Collapsing ISM to FSM...")
lr_collapsed = lr_counts.groupby('associated_transcript').sum().reset_index()
lr_collapsed['tx_base'] = lr_collapsed['associated_transcript'].str.split('.').str[0]
lr_collapsed = lr_collapsed.drop('associated_transcript', axis=1).groupby('tx_base').sum().reset_index()

print("4. Calculating Open-Universe CPM for Long-Read...")
lr_cpm = lr_collapsed.copy()
for sample in sample_map.keys():
    lr_cpm[sample] = (lr_cpm[sample] / lr_totals[sample]) * 1e6
lr_cpm.to_csv(os.path.join(out_dir, 'LR_Isoform_CPM_OpenUniverse_Merged.csv'), index=False)

print("5. Processing Short-Read Data...")
sr_df = pd.read_csv('__PCATLAS_DOCUMENTS_ROOT__/long_reads_results/PolyA_Bossi_10s_27July26/salmon.merged.transcript_counts.tsv', sep='\t')
sr_df = sr_df.drop('gene_id', axis=1)
sr_df.columns = [col.replace('H8660', 'H660') for col in sr_df.columns]
sr_df['tx_base'] = sr_df['tx'].str.split('.').str[0]

sr_totals = {sample: sr_df[sample].sum() for sample in sample_map.keys()}
sr_collapsed = sr_df.drop('tx', axis=1).groupby('tx_base').sum().reset_index()
sr_cpm = sr_collapsed.copy()
for sample in sample_map.keys():
    sr_cpm[sample] = (sr_cpm[sample] / sr_totals[sample]) * 1e6

print("Finding Common Known Transcripts...")
common_tx = set(lr_cpm['tx_base']).intersection(set(sr_cpm['tx_base']))
print(f"Number of overlapping known transcripts: {len(common_tx)}")

lr_cpm_common = lr_cpm[lr_cpm['tx_base'].isin(common_tx)].sort_values('tx_base').reset_index(drop=True)
sr_cpm_common = sr_cpm[sr_cpm['tx_base'].isin(common_tx)].sort_values('tx_base').reset_index(drop=True)

print("Calculating Correlations and Generating Scatter Plots for ALL samples...")
results = []
for sample in sample_map.keys():
    x = np.log10(sr_cpm_common[sample] + 1)
    y = np.log10(lr_cpm_common[sample] + 1)
    pearson_r, p_pearson = pearsonr(x, y)
    spearman_r, p_spearman = spearmanr(x, y)
    results.append({
        'Sample': sample,
        'Pearson (logCPM)': pearson_r, 'Pearson_P': p_pearson,
        'Spearman (logCPM)': spearman_r, 'Spearman_P': p_spearman
    })
    plot_df = pd.DataFrame({
        'Transcript': lr_cpm_common['tx_base'],
        'ShortRead_logCPM': x,
        'LongRead_logCPM': y
    })
    fig = px.scatter(plot_df, x='ShortRead_logCPM', y='LongRead_logCPM',
                     hover_name='Transcript',
                     title=f'Isoform Correlation: LongRead vs ShortRead ({sample})',
                     labels={'ShortRead_logCPM': 'Short-Read log10(CPM+1)', 'LongRead_logCPM': 'Long-Read log10(CPM+1)'},
                     opacity=0.3)
    fig.add_shape(type='line', line=dict(dash='dash', color='red'), x0=0, y0=0, x1=5, y1=5)
    annot_text = f"Pearson: {pearson_r:.3f} (p < 1e-100)<br>Spearman: {spearman_r:.3f} (p < 1e-100)" if p_pearson < 1e-100 else f"Pearson: {pearson_r:.3f} (p={p_pearson:.1e})<br>Spearman: {spearman_r:.3f} (p={p_spearman:.1e})"
    fig.add_annotation(
        x=0.05, y=0.95, xref="paper", yref="paper",
        text=annot_text, showarrow=False, align="left",
        bgcolor="rgba(255,255,255,0.8)", bordercolor="black", borderwidth=1
    )
    fig.write_html(os.path.join(out_dir, f'{sample}_Isoform_Scatter.html'))

res_df = pd.DataFrame(results)
res_df.to_csv(os.path.join(out_dir, 'Isoform_Correlation_Results.csv'), index=False)

melted_df = res_df.melt(id_vars=['Sample'], value_vars=['Pearson (logCPM)', 'Spearman (logCPM)'],
                    var_name='Metric', value_name='Correlation')
fig = px.box(melted_df, x='Metric', y='Correlation', color='Metric', points='all',
             hover_data=['Sample'],
             title='Isoform-Level Correlation Distribution across 10 Samples',
             labels={'Correlation': 'Correlation Coefficient', 'Metric': 'Metric Type'})
fig.update_layout(showlegend=False, template='plotly_white')
fig.write_html(os.path.join(out_dir, 'Correlation_Summary_Boxplot.html'))

print("Part 2: Targeted Isoforms Plots")
genes = {
    'ERF': ['ENST00000440177', 'ENST00000222329'],
    'RELA': ['ENST00000406246', 'ENST00000612991'],
    'RNF43': ['ENST00000584437', 'ENST00000583753'],
    'ROCK2': ['ENST00000401753', 'ENST00000315872'],
    'TRIM11': ['ENST00000284551', 'ENST00000493030'],
    'WNT2B': ['ENST00000369686', 'ENST00000369684']
}
target_isoforms = [iso for isos in genes.values() for iso in isos]
isoform_to_gene = {iso: gene for gene, isos in genes.items() for iso in isos}

samples = list(sample_map.keys())

records = []
for iso in target_isoforms:
    gene = isoform_to_gene[iso]
    lr_row = lr_cpm[lr_cpm['tx_base'] == iso]
    sr_row = sr_cpm[sr_cpm['tx_base'] == iso]

    for sample in samples:
        lr_val = lr_row[sample].values[0] if not lr_row.empty else 0
        sr_val = sr_row[sample].values[0] if not sr_row.empty else 0
        cell_line = sample.split('_')[0]

        records.append({
            'Gene': gene, 'Isoform': iso, 'Sample': sample,
            'CellLine': cell_line, 'Platform': 'LongRead (MAS-Seq)', 'CPM': lr_val
        })
        records.append({
            'Gene': gene, 'Isoform': iso, 'Sample': sample,
            'CellLine': cell_line, 'Platform': 'ShortRead (Salmon)', 'CPM': sr_val
        })

df_plot = pd.DataFrame(records)
df_plot.to_csv(os.path.join(out_dir, 'Specific_12_Isoforms_CPM.csv'), index=False)

target_out_dir = os.path.join(out_dir, 'Target_Isoforms_Plots')
os.makedirs(target_out_dir, exist_ok=True)

for iso in target_isoforms:
    gene = isoform_to_gene[iso]
    df_iso = df_plot[df_plot['Isoform'] == iso]
    fig = px.box(df_iso, x='Platform', y='CPM', color='Platform', points='all',
                 hover_data=['Sample'],
                 title=f'{gene} - {iso} CPM (ShortRead vs LongRead)',
                 template='plotly_white')
    fig.update_traces(marker=dict(size=8))
    fig.update_layout(showlegend=False)
    fig.write_html(os.path.join(target_out_dir, f'{gene}_{iso}_boxplot.html'))

for gene in genes.keys():
    df_gene = df_plot[df_plot['Gene'] == gene]
    fig = px.box(df_gene, x='Isoform', y='CPM', color='Platform', points='all',
                 hover_data=['Sample'],
                 title=f'{gene} Isoforms CPM Comparison (ShortRead vs LongRead)',
                 template='plotly_white')
    fig.write_html(os.path.join(target_out_dir, f'{gene}_both_isoforms_boxplot.html'))

print("DONE ALL")
```
