## Supplementary material for "An integrated, transcript-resolution atlas of prostate cancer progression unmasks associations with oncogenic pathways": Reproducible code: BRCA1_C_experiment_atlas_concordance.R

```
# Public submission copy.
# Analytical statements and parameter values are unchanged from the archived source.
# Legacy non-English comments and console messages were normalized for public release.
library(dplyr)
library(readr)
library(stringr)
library(ggplot2)
library(ggrastr)

# =======================================================
# =======================================================
res_exp <- read.csv("./DEs_11q_vs_CTRL.csv")
res_atlas <- read.csv("../../Data_Harmanization/DEGs_ENST00000357654_Top25_vs_Low25.csv")

res_atlas <- res_atlas %>%
  mutate(ensembl_gene_id = str_replace(X, "\\.[0-9]+", ""))

merged_data <- inner_join(res_exp, res_atlas,
                          by = "ensembl_gene_id",
                          suffix = c("_exp", "_atlas"))

# =======================================================
# =======================================================
p <- ggplot(merged_data, aes(x = log2FoldChange_atlas, y = log2FoldChange_exp)) +
  geom_point_rast(alpha = 0.5, color = "#2c3e50", dpi = 300) +
  geom_hline(yintercept = 0, linetype = "dashed", color = "red", alpha = 0.6) +
  geom_vline(xintercept = 0, linetype = "dashed", color = "red", alpha = 0.6) +
  geom_smooth(method = "lm", color = "blue", se = TRUE, alpha = 0.2) +
  coord_fixed(ratio = 1) +
  labs(
    title = "Comparison of log2FoldChange",
    subtitle = "Exp (11q vs CTRL) vs Atlas (Top25 vs Low25)",
    x = "log2FoldChange (Atlas)",
    y = "log2FoldChange (Exp)"
  ) +
  theme_minimal() +
  theme(
    plot.title = element_text(hjust = 0.5, face = "bold"),
    plot.subtitle = element_text(hjust = 0.5)
  )

print(p)

cor_test <- cor.test(merged_data$log2FoldChange_atlas, merged_data$log2FoldChange_exp)
print(cor_test)

# =======================================================
# =======================================================
plot_data <- merged_data %>%
  mutate(
    is_sig_both = (padj_exp < 0.05 & padj_atlas < 0.05),
    is_same_dir = sign(log2FoldChange_exp) == sign(log2FoldChange_atlas),
    Gene_Category = case_when(
      is_sig_both & is_same_dir ~ "Significant & Consistent (Red)",
      is_sig_both & !is_same_dir ~ "Significant but Opposite (Blue)",
      TRUE ~ "Not Significant or Single (Grey)"
    )
  ) %>%
  mutate(Gene_Category = factor(Gene_Category, levels = c(
    "Not Significant or Single (Grey)",
    "Significant but Opposite (Blue)",
    "Significant & Consistent (Red)"
  ))) %>%
  arrange(Gene_Category)

cor_all <- cor.test(plot_data$log2FoldChange_atlas, plot_data$log2FoldChange_exp)
R_all <- round(cor_all$estimate, 3)
P_all <- format.pval(cor_all$p.value, digits = 3, eps = 0.001)

sig_both_data <- plot_data %>% filter(is_sig_both == TRUE)
cor_sig <- cor.test(sig_both_data$log2FoldChange_atlas, sig_both_data$log2FoldChange_exp)
R_sig <- round(cor_sig$estimate, 3)
P_sig <- format.pval(cor_sig$p.value, digits = 3, eps = 0.001)

stat_label_updated <- paste0(
  "All Genes: R = ", R_all, ", p = ", P_all, "\n",
  "Significant in Both: R = ", R_sig, ", p = ", P_sig
)

max_val <- max(abs(c(plot_data$log2FoldChange_atlas, plot_data$log2FoldChange_exp)), na.rm = TRUE)
axis_limit <- ceiling(max_val)

# =======================================================
# =======================================================
p_square <- ggplot(plot_data, aes(x = log2FoldChange_atlas, y = log2FoldChange_exp, color = Gene_Category)) +
  geom_point_rast(alpha = 0.7, size = 1.5, dpi = 300) +
  scale_color_manual(values = c(
    "Significant & Consistent (Red)" = "#e74c3c",
    "Significant but Opposite (Blue)" = "#3498db",
    "Not Significant or Single (Grey)" = "#ecf0f1"
  )) +
  geom_hline(yintercept = 0, linetype = "dashed", color = "black", alpha = 0.5) +
  geom_vline(xintercept = 0, linetype = "dashed", color = "black", alpha = 0.5) +

  coord_fixed(ratio = 1, xlim = c(-axis_limit, axis_limit), ylim = c(-axis_limit, axis_limit)) +

  labs(
    title = "Concordance of Gene Expression Changes",
    x = "log2FoldChange (Atlas)",
    y = "log2FoldChange (Exp)",
    color = "Gene Status"
  ) +
  theme_minimal(base_size = 12) +
  theme(
    plot.title = element_text(hjust = 0.5, face = "bold", size = 14),
    legend.position = "bottom"
  ) +
  annotate("text", x = -axis_limit, y = axis_limit, label = stat_label_updated,
           hjust = -0.05, vjust = 1.2, size = 4, fontface = "italic", color = "black")

print(p_square)

ggsave("p_final_scatterplot_BRCA_c.pdf", p_square, width = 8, height = 8)
ggsave("p_final_scatterplot_BRCA_c.png", p_square, width = 8, height = 8)
```
