## Supplementary material for "An integrated, transcript-resolution atlas of prostate cancer progression unmasks associations with oncogenic pathways": Reproducible code: RNA_seq.R

```
# Public submission copy.
# Analytical statements and parameter values are unchanged from the archived source.
# Legacy non-English comments and console messages were normalized for public release.
library(dplyr)
library(readr)
library(DESeq2)
library(ggplot2)
library(ggrepel)
library(msigdbr)
library(clusterProfiler)
library(stringr)
library(forcats)
library(limma)
### prepare dataset #######

salmon_merged_gene_counts <- read_delim("salmon.merged.gene_counts.tsv",
                                        delim = "\t", escape_double = FALSE,
                                        trim_ws = TRUE)


annot = data.frame( ID = colnames(salmon_merged_gene_counts)[c(3:8)],
                    REP = rep(1:2, each = 3),
                    TREAT =  c(rep("11q",3), rep("PCW",3)),
                    BATCH = as.factor(c(rep("batch1", 5), "batch2"))
)

rownames(annot) = annot$ID
counts = salmon_merged_gene_counts[,c(3:8)]              %>%
  mutate(across(is.numeric, round))


##### remove pcw1 #####

# counts = counts[,-4]
#
# annot = annot[-4,]
# 3 Pca plot PC1 vs PC2 ----
rownames(counts) = salmon_merged_gene_counts$gene_id

dds <- DESeqDataSetFromMatrix(countData = counts,
                              colData = annot,
                              design = ~ BATCH + TREAT )
dds <- dds[ rowSums(counts(dds)) > 0, ] # tolgo i geni non espressi in nessun sample
dds <- DESeq(dds, betaPrior = FALSE)
dko_vst <- vst(dds, blind = TRUE)
assay(dko_vst) <- limma::removeBatchEffect(assay(dko_vst), batch = annot$BATCH, design = model.matrix(~TREAT, data=annot))
dko_assay = assay(dko_vst)


gold_assay <- as.matrix(dko_assay)
rv = rowVars(gold_assay)
#selected = rownames(counts(dds))[order(rv, decreasing = TRUE)]
selected = order(rv, decreasing = TRUE)
# pcagenes = selected
# length(pcagenes) #10230
top2000 = selected[1:2000] #define top 2000 genes
#selected = top2000
pca = prcomp( t(assay(dko_vst[top2000,])) )
percentVar <- round(100* pca$sdev^2/sum(pca$sdev^2) )
pca = data.frame(PC1 = pca$x[,1], PC2 = pca$x[,2], PC3 = pca$x[,3], PC4 = pca$x[,4], GROUP = annot$ID,   name = rownames(pca$x))
p_gene1 = ggplot(pca, aes(PC1, PC2, color=annot$TREAT)) +
  #geom_point(size=4, aes(fill=annotations$ID)) +
  geom_point(size=8) +
  xlab(paste0("PC1: ",percentVar[1],"% variance")) +
  ylab(paste0("PC2: ",percentVar[2],"% variance")) +
  coord_fixed(ratio = 1) +
  theme( axis.text=element_text(size=12), axis.title=element_text(size=20,family = "Arial")) +
  geom_vline(xintercept = 0, linetype="dotted", size=0.3) + geom_hline(yintercept = 0,linetype="dotted", size=0.3) +
  theme_bw()  + theme(legend.position = "none") +
  geom_label_repel(aes(label = annot$ID),
                   box.padding   = 0.35,
                   point.padding = 0.5,
                   segment.color = 'grey50')
p_gene1
ggsave('./PCA_plot.png', p_gene1, width = 10, height = 10)
ggsave('./PCA_plot.pdf', p_gene1, width = 10, height = 10)


keep_genes = rownames(counts(dds))
SYMBOL = salmon_merged_gene_counts %>%
  filter(gene_id %in% keep_genes ) %>%
  dplyr::select(gene_name)


#5 DIFFERENTIAL EXPRESSION ----

comparisons <- c("11q")


gsea_results <- list()
m_t2g <- msigdbr(species = "Homo sapiens", category = "H") %>%
  dplyr::select(gs_name, ensembl_gene)
for (comp in comparisons) {

  res <- data.frame(results(dds, contrast=c("TREAT", comp, "PCW")))
  res$ensembl_gene_id =  sub("\\..*$", "", rownames(res))
  res$gene_name = SYMBOL$gene_name
  write.csv(res, paste0("./DEs_", comp, "_vs_CTRL.csv"), row.names = FALSE)
  geneList <- res %>%
    filter(!is.na(stat) & !is.na(ensembl_gene_id)) %>%
    distinct(ensembl_gene_id, .keep_all = TRUE) %>%
    arrange(desc(stat)) %>%
    dplyr::select(stat, ensembl_gene_id)
  ranked_genes <- geneList$stat
  names(ranked_genes) <- geneList$ensembl_gene_id
  gsea_result <- GSEA(geneList = ranked_genes,
                                TERM2GENE = m_t2g,
                                pvalueCutoff = 0.05,
                                eps=1e-50,
                                nPermSimple = 10000,
                                verbose = FALSE)
  gene_count <- gsea_result@result %>%
    group_by(ID) %>%
    summarise(count = sum(str_count(core_enrichment, "/")) + 1)
  dot_df <- left_join(gsea_result@result, gene_count, by = "ID") %>%
    mutate(GeneRatio = count/setSize)
  gsea_results[[paste0(comp, "_vs_PWC")]] <- dot_df
}

merged.res <- as.data.frame(merge_result(gsea_results))
merged.res$type = "upregulated"
merged.res$type[merged.res$NES < 0] = "downregulated"
merged.res$SIG = -log10(merged.res$p.adj)

p <- ggplot(merged.res, aes(x = NES, y = fct_reorder(Description, SIG))) +
  geom_point(aes(size = SIG, color = NES)) +
  theme_bw(base_size = 14) +
  scale_colour_gradient2(limits=c(-3.5, 3.5), low = "#3953A4", high = "#b50404") +
  ylab(NULL) + facet_grid(.~Cluster) +
  scale_size(range = c(3,12))
p

ggsave("./GSEA_NES.png",width=10,height=10)
ggsave("./GSEA_NES.pdf",width=10,height=10)
### save all ###
write.csv(merged.res,"./GSEA.csv",row.names=FALSE)
#write.csv(counts,"./counts_table.csv")


tx.counts_table = read_delim("salmon.merged.transcript_counts.tsv",
           delim = "\t", escape_double = FALSE,
           trim_ws = TRUE)


tx.counts = tx.counts_table[,c(3:8)]              %>%
  mutate(across(is.numeric, round))


tpm = log2(sweep(x=tx.counts, MARGIN=2, STATS=colSums(tx.counts)/1000000, FUN="/") + 1)
ERF = tpm[196098,]

ERF_long = data.frame(expression = as.numeric(ERF[1, ]),
                      sample = annot$ID,
                      group = annot$TREAT)
p <- ggplot(ERF_long, aes(x = group, y = expression, fill = group)) +
  geom_boxplot(width = 0.5, outlier.shape = NA, alpha = 0.7) +
  geom_jitter(width = 0.15, alpha = 0.6, size = 2) +

  labs(title = "Expression of BRCA1_truncated by Group",
       x = "Group",
       y = "Log2(TPM + 1)") +

  theme_classic() +

  theme(
    axis.text.x = element_text(angle = 45, hjust = 1),
    legend.position = "none",

    panel.grid.major = element_blank(),
    panel.grid.minor = element_blank(),

    panel.border = element_rect(colour = "black", fill = NA, linewidth = 1)
  )

print(p)

ggsave("./BRCA1_trun_expression.png",plot=p)
ggsave("./BRCA1_trun_expression.pdf",plot=p)
```
