## Supplementary material for "An integrated, transcript-resolution atlas of prostate cancer progression unmasks associations with oncogenic pathways": Reproducible code: reproduce_Fig4H.R

```
#!/usr/bin/env Rscript
# Public submission copy.
# Analytical statements and parameter values are unchanged from the archived source.
# Legacy non-English comments and console messages were normalized for public release.

# Reproduce Figure 4H from the original GraphPad Prism project and add five
# prespecified paired comparisons. All multiplicity-adjusted p values use the
# Sidak family-wise correction across these five comparisons.

suppressPackageStartupMessages({
  library(ggplot2)
})

base_pt <- 5
user_args <- commandArgs(trailingOnly = TRUE)
size_arg <- grep("^--size-cm=", user_args, value = TRUE)
figure_size_cm <- if (length(size_arg)) {
  as.numeric(sub("^--size-cm=", "", size_arg[[1]]))
} else {
  10
}
if (!is.finite(figure_size_cm) || figure_size_cm <= 0) {
  stop("--size-cm must be a positive number")
}
script_arg <- grep("^--file=", commandArgs(trailingOnly = FALSE), value = TRUE)
script_path <- if (length(script_arg)) sub("^--file=", "", script_arg[[1]]) else "reproduce_Fig4H.R"
# Some desktop launchers encode spaces in --file paths as "~+~".
script_path <- gsub("~\\+~", " ", script_path)
script_dir <- dirname(normalizePath(script_path, mustWork = FALSE))
prism_file <- file.path(script_dir, "RNF43_JZ.prism")
out_dir <- file.path(script_dir, "Fig4H_R_output")
dir.create(out_dir, recursive = TRUE, showWarnings = FALSE)

if (!file.exists(prism_file)) stop("Prism source file not found: ", prism_file)

# The Prism project stores the 5 x 12 source table as a CSV member. The column
# order is Control, RNF43-207, RNF43-208, each with four stimulation conditions.
member <- "data/tables/797813C5-5C27-4352-B6AB-CDCC25E42CEF/data.csv"
tmp_dir <- tempfile("rnf43_prism_")
dir.create(tmp_dir)
on.exit(unlink(tmp_dir, recursive = TRUE, force = TRUE), add = TRUE)
utils::unzip(prism_file, files = member, exdir = tmp_dir)
wide <- utils::read.csv(file.path(tmp_dir, member), header = FALSE, check.names = FALSE)
if (!identical(dim(wide), c(5L, 12L))) stop("Unexpected Prism table dimensions: ", paste(dim(wide), collapse = " x "))

source_construct_levels <- c("Control", "RNF43-207", "RNF43-208")
construct_levels <- c("Control", "RNF43-208", "RNF43-207")
construct_axis_labels <- c(
  "Control",
  "RNF43-208\n(ENST00000584437)",
  "RNF43-207\n(ENST00000583753)"
)
stim_levels <- c("EV", "EV + Rspo", "WNT3a", "WNT3a + Rspo")
column_map <- expand.grid(
  stimulation = stim_levels,
  construct = source_construct_levels,
  KEEP.OUT.ATTRS = FALSE,
  stringsAsFactors = FALSE
)
column_map <- column_map[order(match(column_map$construct, source_construct_levels),
                               match(column_map$stimulation, stim_levels)), ]

long <- do.call(rbind, lapply(seq_len(ncol(wide)), function(j) {
  data.frame(
    replicate = seq_len(nrow(wide)),
    construct = column_map$construct[j],
    stimulation = column_map$stimulation[j],
    value = wide[[j]],
    stringsAsFactors = FALSE
  )
}))
long$replicate <- factor(long$replicate)
long$construct <- factor(long$construct, levels = construct_levels)
long$stimulation <- factor(long$stimulation, levels = stim_levels)

# Match the three four-bar blocks in the current Figure 4H.
position_lookup <- data.frame(
  construct = rep(construct_levels, each = 4),
  stimulation = rep(stim_levels, times = 3),
  xpos = c(1:4, 6:9, 11:14),
  stringsAsFactors = FALSE
)
long <- merge(long, position_lookup, by = c("construct", "stimulation"), sort = FALSE)
long <- long[order(long$xpos, long$replicate), ]

summary_df <- aggregate(value ~ construct + stimulation + xpos, long, function(z) {
  c(mean = mean(z), sd = stats::sd(z), n = length(z))
})
summary_df <- data.frame(
  construct = summary_df$construct,
  stimulation = summary_df$stimulation,
  xpos = summary_df$xpos,
  mean = summary_df$value[, "mean"],
  sd = summary_df$value[, "sd"],
  n = summary_df$value[, "n"]
)
summary_df$lower <- pmax(1, summary_df$mean - summary_df$sd)
summary_df$upper <- summary_df$mean + summary_df$sd

get_values <- function(construct, stimulation) {
  z <- long[long$construct == construct & long$stimulation == stimulation, ]
  z$value[order(z$replicate)]
}

comparison_spec <- data.frame(
  comparison_id = paste0("C", 1:5),
  group_1 = c(
    "Control | WNT3a",
    "Control | WNT3a",
    "RNF43-208 | WNT3a",
    "RNF43-207 | WNT3a",
    "RNF43-208 | WNT3a"
  ),
  group_2 = c(
    "RNF43-207 | WNT3a",
    "RNF43-208 | WNT3a",
    "RNF43-207 | WNT3a",
    "RNF43-207 | WNT3a + Rspo",
    "RNF43-208 | WNT3a + Rspo"
  ),
  x1 = c(3, 3, 8, 13, 8),
  x2 = c(13, 8, 13, 14, 9),
  bracket_y = c(300, 150, 215, 112, 94),
  label_y = c(326, 163, 234, 122, 102),
  stringsAsFactors = FALSE
)

parse_group <- function(x) strsplit(x, " \\| ")[[1]]
test_rows <- lapply(seq_len(nrow(comparison_spec)), function(i) {
  a <- parse_group(comparison_spec$group_1[i])
  b <- parse_group(comparison_spec$group_2[i])
  x <- get_values(a[1], a[2])
  y <- get_values(b[1], b[2])
  tt <- stats::t.test(x, y, paired = TRUE, alternative = "two.sided")
  diff <- x - y
  data.frame(
    comparison_id = comparison_spec$comparison_id[i],
    group_1 = comparison_spec$group_1[i],
    group_2 = comparison_spec$group_2[i],
    n_pairs = length(diff),
    mean_group_1 = mean(x),
    mean_group_2 = mean(y),
    mean_difference = mean(diff),
    ci_low = unname(tt$conf.int[1]),
    ci_high = unname(tt$conf.int[2]),
    t = unname(tt$statistic),
    df = unname(tt$parameter),
    p_raw = tt$p.value,
    dz = mean(diff) / stats::sd(diff),
    stringsAsFactors = FALSE
  )
})
stats_df <- do.call(rbind, test_rows)
family_size <- nrow(stats_df)
stats_df$p_sidak <- pmin(1, 1 - (1 - stats_df$p_raw)^family_size)
stats_df$significance <- cut(
  stats_df$p_sidak,
  breaks = c(-Inf, 0.0001, 0.001, 0.01, 0.05, Inf),
  labels = c("****", "***", "**", "*", "ns"),
  right = FALSE
)

format_p <- function(p) {
  if (p < 0.0001) return("p_adj < 0.0001")
  if (p < 0.001) return(sprintf("p_adj = %.6f", p))
  if (p < 0.01) return(sprintf("p_adj = %.4f", p))
  return(sprintf("p_adj = %.3f", p))
}
comparison_spec$label <- paste0(stats_df$significance, "\n", vapply(stats_df$p_sidak, format_p, character(1)))

# Save transparent source and analysis tables.
utils::write.csv(long, file.path(out_dir, "Fig4H_source_data_long.csv"), row.names = FALSE)
utils::write.csv(summary_df, file.path(out_dir, "Fig4H_descriptive_statistics_mean_SD.csv"), row.names = FALSE)
utils::write.csv(stats_df, file.path(out_dir, "Fig4H_five_planned_comparisons_Sidak.csv"), row.names = FALSE)

# Repeated-measures two-factor ANOVA is retained as a global design check. The
# five displayed comparisons remain the prespecified paired contrasts above.
anova_fit <- stats::aov(
  value ~ construct * stimulation + Error(replicate / (construct * stimulation)),
  data = long
)
capture.output(
  summary(anova_fit),
  file = file.path(out_dir, "Fig4H_two_way_repeated_measures_ANOVA.txt")
)

palette <- c(
  "EV" = "#B7B7B7",
  "EV + Rspo" = "#F4A11A",
  "WNT3a" = "#A61B1B",
  "WNT3a + Rspo" = "#009C9A"
)

bracket_segments <- do.call(rbind, lapply(seq_len(nrow(comparison_spec)), function(i) {
  y <- comparison_spec$bracket_y[i]
  drop <- 2^(log2(y) - 0.09)
  data.frame(
    x = c(comparison_spec$x1[i], comparison_spec$x1[i], comparison_spec$x2[i]),
    xend = c(comparison_spec$x1[i], comparison_spec$x2[i], comparison_spec$x2[i]),
    y = c(drop, y, y),
    yend = c(y, y, drop)
  )
}))

label_x <- (comparison_spec$x1 + comparison_spec$x2) / 2
if (figure_size_cm <= 7) {
  label_x[c(1, 4)] <- c(5.5, 13.0)
}
label_df <- data.frame(
  x = label_x,
  y = comparison_spec$label_y,
  label = comparison_spec$label,
  stringsAsFactors = FALSE
)

set.seed(20260820)
p <- ggplot() +
  geom_col(
    data = summary_df,
    aes(x = xpos, y = mean, fill = stimulation),
    width = 0.72, color = "black", linewidth = 0.32
  ) +
  geom_errorbar(
    data = summary_df,
    aes(x = xpos, ymin = lower, ymax = upper),
    width = 0.25, linewidth = 0.32, color = "black"
  ) +
  geom_point(
    data = long[long$value >= 1, ],
    aes(x = xpos, y = value),
    position = position_jitter(width = 0.09, height = 0, seed = 20260820),
    shape = 21, size = 1.35, stroke = 0.25, fill = "black", color = "black"
  ) +
  geom_segment(
    data = bracket_segments,
    aes(x = x, xend = xend, y = y, yend = yend),
    linewidth = 0.28, color = "black", inherit.aes = FALSE
  ) +
  geom_text(
    data = label_df,
    aes(x = x, y = y, label = label),
    size = base_pt / 2.845276, family = "sans", lineheight = 0.9, vjust = 0,
    inherit.aes = FALSE
  ) +
  annotate(
    "text", x = c(2.5, 7.5, 12.5), y = 0.26,
    label = construct_axis_labels, family = "sans", size = base_pt / 2.845276,
    fontface = "bold", lineheight = 0.92, vjust = 1
  ) +
  scale_fill_manual(values = palette, guide = "none") +
  scale_x_continuous(
    breaks = position_lookup$xpos,
    labels = position_lookup$stimulation,
    limits = c(0.2, 14.8),
    expand = expansion(mult = c(0, 0))
  ) +
  scale_y_continuous(
    trans = "log2",
    breaks = c(1, 2, 4, 8, 16, 32, 64, 128),
    labels = c("1", "2", "4", "8", "16", "32", "64", "128"),
    expand = expansion(mult = c(0, 0.015))
  ) +
  labs(x = NULL, y = "TOPFlash reporter") +
  coord_cartesian(ylim = c(1, 400), clip = "off") +
  theme_classic(base_family = "sans", base_size = base_pt) +
  theme(
    text = element_text(family = "sans", size = base_pt, color = "black"),
    axis.title.y = element_text(size = base_pt, margin = margin(r = 3)),
    axis.text.y = element_text(size = base_pt, color = "black"),
    axis.text.x = element_text(
      size = base_pt, color = "black", angle = 48, hjust = 1, vjust = 1,
      margin = margin(t = 1)
    ),
    axis.ticks = element_line(linewidth = 0.3, color = "black"),
    axis.ticks.length = grid::unit(1.6, "pt"),
    axis.line = element_line(linewidth = 0.35, color = "black"),
    plot.margin = margin(t = 4, r = 4, b = 48, l = 6, unit = "pt")
  )

size_suffix <- if (identical(figure_size_cm, 10)) "" else paste0("_", format(figure_size_cm, trim = TRUE), "cm")
pdf_file <- file.path(out_dir, paste0("Fig4H_R_reproduced_with_5_comparisons", size_suffix, ".pdf"))
ggplot2::ggsave(
  filename = pdf_file,
  plot = p,
  width = figure_size_cm,
  height = figure_size_cm,
  units = "cm",
  device = grDevices::pdf,
  family = "Helvetica",
  useDingbats = FALSE,
  compress = FALSE,
  bg = "white"
)

message("Created: ", pdf_file)
message("Statistics: ", file.path(out_dir, "Fig4H_five_planned_comparisons_Sidak.csv"))
```
