## Supplementary material for "An integrated, transcript-resolution atlas of prostate cancer progression unmasks associations with oncogenic pathways": Reproducible code: BRAC1_customized.R

```
# Public submission copy.
# Analytical statements and parameter values are unchanged from the archived source.
# Legacy non-English comments and console messages were normalized for public release.

pfam_info <- getBM(attributes = c("ensembl_transcript_id", "pfam", "pfam_start", "pfam_end"),
                   filters    = "ensembl_transcript_id",
                   values     = up_rna,
                   mart = ensembl)
info <- getBM(attributes = c("ensembl_transcript_id", "cds_length" ,"exon_chrom_start","exon_chrom_end","rank","cds_start","cds_end","5_utr_start","5_utr_end","3_utr_start","3_utr_end"),
              filters    = "ensembl_transcript_id",
              values     = up_rna,
              mart = ensembl)

info$start = 1
info$end = info$cds_length
info$gene=name

pfam_info = na.omit(pfam_info)
info = info %>% filter(cds_length == max(cds_length))
pfam_aug <- pfam_info %>% left_join(Pfam_clans, by = "pfam") %>%
  mutate(label = ifelse(is.na(label), "Not Found", label))
pfam_aug$orientation =info$exon_chrom_start[info$rank==1]<
  info$exon_chrom_start[info$rank==2]
pfam_aug$start = 1
pfam_aug$end = max(info$cds_length)

pfam_info_2 <- getBM(attributes = c("ensembl_transcript_id", "pfam", "pfam_start", "pfam_end"),
                     filters    = "ensembl_transcript_id",
                     values     = down_rna,
                     mart = ensembl)
info_2 <- getBM(attributes = c("ensembl_transcript_id", "cds_length" ,"exon_chrom_start","exon_chrom_end","rank","cds_start","cds_end","5_utr_start","5_utr_end","3_utr_start","3_utr_end"),
                filters    = "ensembl_transcript_id",
                values     = down_rna,
                mart = ensembl)

info_2$start = 1
info_2$end = info_2$cds_length

info_2$gene=name

pfam_info_2 = na.omit(pfam_info_2)
info_2 = info_2 %>% filter(cds_length == max(cds_length))

pfam_aug_2 <- pfam_info_2 %>% left_join(Pfam_clans, by = "pfam") %>%
  mutate(label = ifelse(is.na(label), "Not Found", label))
pfam_aug_2$orientation =info_2$exon_chrom_start[info_2$rank==1]<
  info_2$exon_chrom_start[info_2$rank==2]
pfam_aug_2$start = 1
pfam_aug_2$end = max(info_2$cds_length)


###### customized pfam_aug and pfam_aug_2


pfam_aug[2,7] = "RING"
add_1 = pfam_aug[1,]

add_1$pfam_start = 1362
add_1$pfam_end = 1437
add_1$Domain= "CCD"
pfam_aug = rbind(pfam_aug,add_1)

add_1$pfam_start = 81
add_1$pfam_end = 99
add_1$Domain= "NES"
pfam_aug = rbind(pfam_aug,add_1)

add_1$pfam_start = 503
add_1$pfam_end = 508
add_1$Domain= "NLS"
pfam_aug = rbind(pfam_aug,add_1)


add_1$pfam_start = 607
add_1$pfam_end = 614
add_1$Domain= "NLS"
pfam_aug = rbind(pfam_aug,add_1)


pfam_aug_2[1,7] = "RING"

add_1 = pfam_aug_2[1,]


add_1$pfam_start = 1362-1658+515
add_1$pfam_end = 1437-1658+515
add_1$Domain= "CCD"
pfam_aug_2 = rbind(pfam_aug_2,add_1)

add_1$pfam_start = 81
add_1$pfam_end = 99
add_1$Domain= "NES"
pfam_aug_2 = rbind(pfam_aug_2,add_1)


#start drawing


height = 1/64
info_order = info %>% arrange(exon_chrom_start)
intron = data.frame(start = info_order$exon_chrom_end[1:dim(info_order)[1]-1],
                    end = info_order$exon_chrom_start[2:dim(info_order)[1]])
intron$mid = (intron$start + intron$end)/2


info_order_2 = info_2 %>% arrange(exon_chrom_start)
intron_2 = data.frame(start = info_order_2$exon_chrom_end[1:dim(info_order_2)[1]-1],
                      end = info_order_2$exon_chrom_start[2:dim(info_order_2)[1]])
intron_2$mid = (intron_2$start + intron_2$end)/2

padding = 1/400

t = ggplot() +
  geom_rect(data=info,aes(xmin = exon_chrom_start, xmax = exon_chrom_end, ymin = 1+padding, ymax = 1+height,fill = "white" ),color="black" )  +
  geom_rect(data=info_2,aes(xmin = exon_chrom_start, xmax = exon_chrom_end, ymin = 1-height, ymax = 1-padding,fill = "white") ,color="black" ) +
  geom_segment(data=intron,aes(x=start,xend=mid,y=1+height/2+padding,yend=1+4/5*height+padding))+
  geom_segment(data=intron,aes(x=mid,xend=end,y=1+4/5*height+padding,yend=1+height/2+padding))+
  geom_segment(data=intron_2,aes(x=start,xend=mid,y=1-height/2-padding,yend=1-4/5*height-padding))+
  geom_segment(data=intron_2,aes(x=mid,xend=end,y=1-4/5*height-padding,yend=1-height/2-padding))+
  theme_minimal()+
  theme(panel.grid = element_blank(),
        panel.grid.major = element_blank(),
        panel.grid.minor = element_blank(),
        panel.background = element_blank(),
        axis.text = element_blank(),
        axis.ticks = element_blank())

t
rectangle1_length <- info$cds_length
rectangle2_length <- info_2$cds_length

info_filter <- info %>% arrange(exon_chrom_start) %>% filter(!is.na(cds_start))
if (info_filter$rank[1]>info_filter$rank[2]){
  info_filter$exon_chrom_start[1] = info_filter$`3_utr_end`[1]
  info_filter$exon_chrom_end[dim(info_filter)[1]] = info_filter$`5_utr_start`[dim(info_filter)[1]]
}
info_filter_2 <- info_2 %>% arrange(exon_chrom_start) %>% filter(!is.na(cds_start))
if (info_filter_2$rank[1]>info_filter_2$rank[2]){
  info_filter_2$exon_chrom_start[1] = info_filter_2$`3_utr_end`[1]
  info_filter_2$exon_chrom_end[dim(info_filter_2)[1]] = info_filter_2$`5_utr_start`[dim(info_filter_2)[1]]
}

t2 = t +

  geom_rect(data=info_filter,aes(xmin = exon_chrom_start, xmax = exon_chrom_end, ymin = 1+padding, ymax = 1+height,fill = "lightgrey" ),color="black" )  +
  geom_rect(data=info_filter_2,aes(xmin = exon_chrom_start, xmax = exon_chrom_end, ymin = 1-height, ymax = 1-padding,fill = "lightgrey" ),color="black" )  +
  labs(x=NULL, y=NULL,fill="mRNA") +

  scale_fill_manual(values = c("darkgrey", "lightgrey"),
                    labels = c("CDS", "UTR"))

to = info_filter
V4 = sort(c(to$exon_chrom_start,to$exon_chrom_end))


start = V4[1]+500
end = tail(V4,1)-500
if(rectangle1_length[1]>=rectangle2_length[1]){
  end_2 = (end-start)/rectangle1_length*rectangle2_length+start
  end_2 = end_2[1]
  end_1 = end}
if(rectangle2_length[1]>rectangle1_length[1]){
  end_1 = (end-start)/rectangle2_length*rectangle1_length+start
  end_1 = end_1[1]
  end_2 = end
}

V5= rescale(sort(c(info$cds_start,info$cds_end),decreasing = T), to=c(1, (start-end_1)))
V5 = V5 + (end_1-tail(V5,1))
connection_points_upper <- data.frame(
  V4 = V4,
  V5 = V5
)

pfam_aug$start = end_1+pfam_aug$pfam_end*(start-end_1)/rectangle1_length[1]*3
pfam_aug$end = end_1+pfam_aug$pfam_start*(start-end_1)/rectangle1_length[1]*3

height2 = 1/32
top_start = 1.05
t3= t2 +new_scale_fill() + geom_segment(
  data = connection_points_upper,
  aes(x = V4, y = 1+height, xend = V5, yend = top_start),
  color = "gray", size = 0.5
) +
  geom_rect(data=pfam_aug,
            aes(xmin=start,xmax = end,ymin = top_start, ymax =top_start+height2,fill=Domain))+
  geom_rect(
    aes(xmin = start, xmax = end_1, ymin = top_start, ymax =top_start+height2),
    fill = "white", alpha = 0,color="black")
t3


start_2 = 1/2*min(info_2$exon_chrom_start)+1/2*(max(info_2$exon_chrom_end))

info_filter <- info_2 %>% arrange(exon_chrom_start) %>% filter(!is.na(cds_start))
if (info_filter$rank[1]>info_filter$rank[2]){
  info_filter$exon_chrom_start[1] = info_filter$`3_utr_end`[1]
  info_filter$exon_chrom_end[dim(info_filter)[1]] = info_filter$`5_utr_start`[dim(info_filter)[1]]
}
to = info_filter


V4 = sort(c(to$exon_chrom_start,to$exon_chrom_end))
V5= rescale(sort(c(info_2$cds_start,info_2$cds_end),decreasing = T), to=c(1, (start-end_2)))
V5 = V5 + (end_2-tail(V5,1))
connection_points_lower <- data.frame(
  V4 = V4,
  V5 = V5
)
pfam_aug_2$start = end_2+pfam_aug_2$pfam_end*(start-end_2)/rectangle2_length[1]*3
pfam_aug_2$end = end_2+pfam_aug_2$pfam_start*(start-end_2)/rectangle2_length[1]*3

height2 = 1/32
down_start = 0.95


t4 = t3 +geom_segment(
  data = connection_points_lower,
  aes(x = V5, y = down_start, xend = V4, yend = 1-height),
  color = "gray", size = 0.5
) +
  geom_rect(data=pfam_aug_2,
            aes(xmin=start,xmax = end,ymin = down_start-height2, ymax =down_start,fill=Domain))+geom_rect(
              aes(xmin = start, xmax = end_2, ymin = down_start-height2, ymax =down_start ),
              fill = "white", alpha = 0,color="black")+
  theme(
    legend.text = element_text(size = 24),
    legend.title = element_text(size = 30)
  )
t4
start_pro_up = start
end_pro_up = end_1
start_pro_down = start
end_pro_down = end_2
start_rna_up = min(info$exon_chrom_start)
end_rna_up = max(info$exon_chrom_end)
start_rna_down = min(info_2$exon_chrom_start)
end_rna_down = max(info_2$exon_chrom_end)
end_all = max(end_rna_up,end_pro_up,end_rna_down,end_pro_down)

max_length = (max(-start_rna_up+end_rna_up,-start_rna_down+end_rna_down))
ratio_label = max_length/40
ratio_text = max_length/2.5
t5 = t4+
  annotate("text", x=start_pro_down-ratio_label, y= down_start-height2/2, label= "C",size=24/.pt)+
  annotate("text", x=end_pro_down+ratio_label, y= down_start-height2/2, label= "N",size=24/.pt)+
  annotate("text", x=start_pro_up-ratio_label, y=top_start+height2/2, label= "C",size=24/.pt)+
  annotate("text", x=end_pro_up+ratio_label, y=top_start+height2/2, label= "N",size=24/.pt)+
  annotate("text", x=min(start_rna_up,start_rna_down)-ratio_label, y= 1, label= "3'",size=24/.pt)+
  annotate("text", x=max(end_rna_up,end_rna_down)+ratio_label, y= 1, label= "5'",size=24/.pt)+
  annotate("text", x=end_all+ratio_text, y= 1+padding*3+height/2, label= up_rna,size=24/.pt,hjust=0)+
  annotate("text", x=end_all+ratio_text, y= top_start+height2/2, label= up_pro,size=24/.pt,hjust=0)+
  annotate("text", x=end_all+ratio_text, y= 1-padding*3-height/2, label= down_rna,size=24/.pt,hjust=0)+
  annotate("text", x=end_all+ratio_text, y=down_start-height2/2, label= down_pro,size=24/.pt,hjust=0)+
  annotate("text",x=max(end_rna_up,end_pro_up,end_rna_down,end_pro_down)+ratio_text, y=1.2, label= paste0(name," isoforms"),size=30/.pt,hjust=0)


connection_points_lower$ystart =  down_start-height2
connection_points_lower$yend =  down_start
connection_points_upper$ystart = top_start
connection_points_upper$yend = top_start+height2

final=t5 + geom_segment(data = connection_points_lower, aes(x = V5, y = ystart, xend = V5, yend = yend), size = 0.5)+geom_segment(data = connection_points_upper, aes(x = V5, y = ystart, xend = V5, yend = yend), size = 0.5)+ ylim(0.5,1.5) +scale_x_reverse()+
  theme(plot.margin = margin(l = 100, unit = "pt"))+
  coord_cartesian(clip = "off")
final
ggsave(filename = paste0(name,".pdf"),plot = final,width=18,height=15)
```
