## Supplementary material for "An integrated, transcript-resolution atlas of prostate cancer progression unmasks associations with oncogenic pathways": Reproducible code: draw_init.R

```
# Public submission copy.
# Analytical statements and parameter values are unchanged from the archived source.
# Legacy non-English comments and console messages were normalized for public release.
library(magrittr)
library(dplyr)
library(ggplot2)
library(ggtranscript)
library(stringr)
library(readr)
library(scales)
library(ggnewscale)
library(biomaRt)
Pfam_clans <- read_delim("~/Downloads/Pfam-A.clans.tsv",
                         delim = "\t", escape_double = FALSE,
                         col_names = c("pfam","x2","x3","Domain","label"), trim_ws = TRUE)

ensembl = useMart("ensembl", dataset = "hsapiens_gene_ensembl")


### ERF #####
up_rna = "ENST00000440177"
down_rna = "ENST00000222329"
up_pro = "P50548-2"
down_pro = "P50548-1"
name = "ERF"

### TRIM11 #####
up_rna = "ENST00000284551"
down_rna = "ENST00000493030"
up_pro = "Q96F44-1"
down_pro = "R4GMV1"
name = "TRIM11"

### BRCA1 #####
up_rna = "ENST00000352993"
down_rna = "ENST00000357654"
up_pro = "P38398-5"
down_pro = "P38398-1"
name = "BRCA1"

### ROCK2 #####
up_rna = "ENST00000401753"
down_rna = "ENST00000315872"
up_pro = "E9PF63"
down_pro = "O75116"
name = "ROCK2"

### RELA #####
up_rna = "ENST00000406246"
down_rna = "ENST00000612991"
up_pro = "Q04206-1"
down_pro = "A0A087X0W8"
name = "RELA"

### F8 #####
up_rna = "ENST00000330287"
down_rna = "ENST00000360256"
up_pro = "P00451-2"
down_pro = "P00451-1"
name = "F8"

### RNF43 #####
up_rna = "ENST00000584437"
down_rna = "ENST00000583753"
up_pro = "Q68DV7-1"
down_pro = "Q68DV7-2"
name = "RNF43"

############ Positive ones

### CDK12 #####
up_rna = "ENST00000447079"
down_rna = "ENST00000584632"
up_pro = "Q9NYV4-1"
down_pro = "J3QSD7"
name = "CDK12"

### WNT2B #####
up_rna = "ENST00000369686"
down_rna = "ENST00000369684"
up_pro = "Q93097-2"
down_pro = "Q93097-1"
name = "WNT2B"

### AR #####
up_rna = "ENST00000504326"
down_rna = "ENST00000374690"
up_pro = "P10275-4"
down_pro = "P10275-1"
name = "AR"


#### PPP2R5C ######
up_rna = "ENST00000328724"
down_rna = "ENST00000445439"
up_pro = "Q13362-1"
down_pro = "Q13362-3"
name = "PPP2R5C"
```
