## Supplementary material for "An integrated, transcript-resolution atlas of prostate cancer progression unmasks associations with oncogenic pathways": Reproducible code: create_fig3d_matched_revision.R

```
# Public submission copy.
# Analytical statements and parameter values are unchanged from the archived source.
# Legacy non-English comments and console messages were normalized for public release.
# Figure 3D revision: retain highlighted non-canonical transcripts only when
# the same gene has a canonical transcript in the complete input table.
# The point layers remain locally rasterized with ggrastr, as in the original.

suppressPackageStartupMessages({
  library(ggplot2)
  library(dplyr)
  library(ggrepel)
  library(ggrastr)
})

output_dir <- "__PCATLAS_PROJECT_ROOT__/Fig3D_canonical_matched_revision_cm"
dir.create(output_dir, recursive = TRUE, showWarnings = FALSE)

# The analysis table was supplied as XLSX. A CSV export is used as a
# dependency-free fallback when openxlsx is not installed in the R runtime.
xlsx_path <- "__PCATLAS_DOCUMENTS_ROOT__/Data_Harmanization/pc_tx_info_filter_5pct_JP.xlsx"
csv_path  <- "__PCATLAS_DOCUMENTS_ROOT__/Data_Harmanization/pc_tx_info_filter_5pct.csv"
legacy_csv_xlsx_export <- "__PCATLAS_PROJECT_ROOT__/Fig3D_canonical_matched_revision/pc_tx_info_filter_5pct_JP_from_xlsx.csv"
csv_xlsx_export <- file.path(output_dir, "pc_tx_info_filter_5pct_JP_from_xlsx.csv")
if (file.exists(csv_xlsx_export)) {
  df <- read.csv(csv_xlsx_export, stringsAsFactors = FALSE, check.names = FALSE)
  input_used <- paste0(xlsx_path, " (value-preserving CSV export)")
} else if (file.exists(legacy_csv_xlsx_export)) {
  df <- read.csv(legacy_csv_xlsx_export, stringsAsFactors = FALSE, check.names = FALSE)
  input_used <- paste0(xlsx_path, " (value-preserving CSV export; prior revision)")
} else if (requireNamespace("openxlsx", quietly = TRUE) && file.exists(xlsx_path)) {
  df <- openxlsx::read.xlsx(xlsx_path, sheet = 1)
  input_used <- xlsx_path
} else if (file.exists(csv_path)) {
  df <- read.csv(csv_path, stringsAsFactors = FALSE, check.names = FALSE)
  input_used <- csv_path
} else {
  stop("Neither the original XLSX nor its CSV export could be found.")
}

required <- c("gene_name", "transcript_id", "cor_pseudotime", "is_canonical")
missing_cols <- setdiff(required, names(df))
if (length(missing_cols) > 0) stop("Missing required columns: ", paste(missing_cols, collapse = ", "))

df <- df %>%
  filter(!is.na(cor_pseudotime), !is.na(gene_name), !is.na(is_canonical)) %>%
  mutate(is_canonical = as.integer(is_canonical), group = "Background")

# Preserve the original selection rule: only the first 21 rows of the input
# table were initially highlighted.
initial_rows <- seq_len(min(21L, nrow(df)))
initial <- df[initial_rows, , drop = FALSE]
initial_canonical <- initial %>% filter(is_canonical == 1)
initial_noncanonical <- initial %>% filter(is_canonical == 0, gene_name != "PPP2R5C")

# For every initially blue point, search the complete table for the same gene's
# canonical transcript(s). Blue points without a match are removed. All matched
# canonical rows are added to the highlighted set (including rows outside the
# initial 21, if present).
blue_genes <- unique(initial_noncanonical$gene_name)
canonical_matches <- df %>%
  filter(gene_name %in% blue_genes, is_canonical == 1) %>%
  distinct(transcript_id, .keep_all = TRUE)
matched_blue <- initial_noncanonical %>%
  filter(gene_name %in% c(canonical_matches$gene_name, "TP63", "AR-V7"))

highlight <- bind_rows(initial_canonical, matched_blue, canonical_matches) %>%
  distinct(transcript_id, .keep_all = TRUE) %>%
  mutate(group = if_else(is_canonical == 1, "Canonical", "Non-canonical"))

plot_data <- df %>%
  arrange(cor_pseudotime, transcript_id) %>%
  mutate(rank = seq_len(n()))

# Add the TP63 MANE/canonical transcript from the unfiltered analysis table.
# Its correlation is taken verbatim from pc_tx_info_unfiltered.xlsx; existing
# filtered-table ranks are not shifted by this additional point.
tp63_canonical_row <- tibble(
  gene_name = "TP63",
  transcript_id = "ENST00000264731.8",
  cor_pseudotime = 0.24268466547807,
  is_canonical = 1L,
  rank = sum(plot_data$cor_pseudotime <= 0.24268466547807) + 0.5,
  group = "Canonical"
)
plot_data <- bind_rows(plot_data, tp63_canonical_row)
plot_data$group <- "Background"
plot_data$group[match(highlight$transcript_id, plot_data$transcript_id)] <- highlight$group
plot_data$group[match(tp63_canonical_row$transcript_id, plot_data$transcript_id)] <- "Canonical"
highlight_data <- plot_data %>% filter(group != "Background")
write.csv(highlight_data, file.path(output_dir, "highlighted_transcripts_final.csv"), row.names = FALSE)

# A reproducible audit table records every initially blue gene and its decision.
audit <- initial_noncanonical %>%
  transmute(
    gene_name,
    initial_blue_transcript = transcript_id,
    initial_blue_correlation = cor_pseudotime,
    canonical_match_found = gene_name %in% canonical_matches$gene_name,
    matched_canonical_transcripts = vapply(
      gene_name,
      function(g) paste(canonical_matches$transcript_id[canonical_matches$gene_name == g], collapse = ";"),
      character(1)
    ),
    action = case_when(
      gene_name == "TP63" ~ "retained blue point; label changed to ΔNp63",
      gene_name == "AR-V7" ~ "retained blue point; no local canonical match",
      canonical_match_found ~ "retained blue point; canonical point highlighted",
      TRUE ~ "blue point removed"
    )
  )
write.csv(audit, file.path(output_dir, "blue_point_canonical_match_audit.csv"), row.names = FALSE)
write.csv(highlight_data, file.path(output_dir, "highlighted_transcripts_final.csv"), row.names = FALSE)
writeLines(c(
  paste0("Input table: ", input_used),
  "Initial highlight rule: first 21 rows; is_canonical=1 red, is_canonical=0 blue.",
  "Revision rule: retain an initial blue point only if the complete table contains a canonical transcript with the same gene_name; highlight all matched canonical rows.",
  paste0("Initial blue genes: ", paste(blue_genes, collapse = ", ")),
  paste0("Matched genes: ", paste(unique(canonical_matches$gene_name), collapse = ", ")),
  paste0("Removed blue genes: ", paste(setdiff(blue_genes, unique(canonical_matches$gene_name)), collapse = ", ")),
  "Point layers are rasterized at 300 dpi with ggrastr; text remains vector.",
  "Final device size: width 9 cm x height 4.5 cm; all text is 5 pt.",
  "PPP2R5C highlighted points were removed; TP63 is labeled ΔNp63."
  ,"TP63 canonical point ENST00000264731.8 was added from pc_tx_info_unfiltered.xlsx; its rank was inserted without shifting existing filtered-table ranks."
), file.path(output_dir, "README_revision.txt"))

text_pt <- 5
text_mm <- text_pt / 2.845
p <- ggplot(plot_data, aes(x = cor_pseudotime, y = rank)) +
  rasterise(
    geom_point(data = subset(plot_data, group == "Background"),
               color = "grey85", size = 0.55, alpha = 0.3), dpi = 300
  ) +
  rasterise(
    geom_point(data = highlight_data, aes(color = group), size = 1.5, alpha = 1), dpi = 300
  ) +
  scale_color_manual(values = c("Canonical" = "#E41A1C", "Non-canonical" = "#004c8c")) +
  geom_text_repel(
    data = subset(highlight_data, transcript_id != "ENST00000354600.10"),
    aes(label = gene_name, color = group),
    size = text_mm,
    fontface = "bold",
    box.padding = 0.35,
    point.padding = 0.2,
    max.overlaps = Inf
    ,force = 10,
    force_pull = 0.15,
    max.iter = 10000,
    max.time = 10,
    min.segment.length = 0,
    segment.size = 0.35,
    segment.alpha = 1,
    segment.color = "grey25",
    seed = 42
  ) +
  geom_text_repel(
    data = subset(highlight_data, transcript_id == "ENST00000354600.10"),
    aes(label = "Delta*Np63", color = group),
    parse = TRUE,
    size = text_mm,
    fontface = "bold",
    box.padding = 0.35,
    point.padding = 0.2,
    max.overlaps = Inf,
    force = 10,
    force_pull = 0.15,
    max.iter = 10000,
    max.time = 10,
    min.segment.length = 0,
    segment.size = 0.35,
    segment.alpha = 1,
    segment.color = "grey25",
    seed = 42
  ) +
  geom_vline(xintercept = 0, linetype = "dashed", color = "grey40") +
  labs(title = "Sigmoid Correlation Rank Plot (Top 21)",
       x = "Correlation Coefficient", y = "Rank") +
  scale_x_continuous(limits = c(-1, 1), breaks = seq(-1, 1, 0.5), expand = c(0.01, 0.01)) +
  scale_y_continuous(limits = c(0, 14000)) +
  theme_classic(base_size = text_pt) +
  theme(
    plot.title = element_text(size = text_pt, face = "bold", hjust = 0.5),
    axis.title = element_text(size = text_pt),
    axis.text = element_text(size = text_pt),
    legend.title = element_text(size = text_pt),
    legend.text = element_text(size = text_pt),
    legend.position = "top",
    legend.key.size = grid::unit(2.5, "mm"),
    plot.margin = margin(3, 3, 3, 3, unit = "pt")
  )

ggsave(file.path(output_dir, "Fig3D_canonical_matched_revision_cm.pdf"), p,
       width = 9, height = 4.5, units = "cm", dpi = 300, limitsize = FALSE)
ggsave(file.path(output_dir, "Fig3D_canonical_matched_revision_cm.png"), p,
       width = 9, height = 4.5, units = "cm", dpi = 300, limitsize = FALSE)

message("Final highlighted transcripts: ", nrow(highlight_data))
print(audit)
```
