## Supplementary material for "An integrated, transcript-resolution atlas of prostate cancer progression unmasks associations with oncogenic pathways": Reproducible code: plot_TRIM11_isoforms_vs_pseudotime.R

```
#!/usr/bin/env Rscript
# Public submission copy.
# Analytical statements and parameter values are unchanged from the archived source.
# Legacy non-English comments and console messages were normalized for public release.

# Reproduction-style plot based on Figure 5B:
# gene-level TRIM11 and the two indicated TRIM11 transcripts versus atlas pseudotime.
# Output dimensions: 9 x 6 cm; base font size: 5 pt.

suppressPackageStartupMessages({
  library(fst)
  library(ggplot2)
  library(dplyr)
  library(tidyr)
})

atlas_dir <- "__PCATLAS_DOCUMENTS_ROOT__/WebToolStudy"
out_pdf <- "__PCATLAS_PROJECT_ROOT__/TRIM11_isoforms_vs_pseudotime_5B_style.pdf"
out_png <- "__PCATLAS_PROJECT_ROOT__/TRIM11_isoforms_vs_pseudotime_5B_style.png"

# The atlas matrices are stored with columns in the same sample order as filtered_samples.csv.
mdata <- read.csv(file.path(atlas_dir, "filtered_samples.csv"), stringsAsFactors = FALSE)
tx_counts <- as.matrix(read.fst(file.path(atlas_dir, "txi.counts.fst")))
gene_counts <- as.matrix(read.fst(file.path(atlas_dir, "gc_txi.counts.fst")))
tx_info <- read.csv(file.path(atlas_dir, "transcripts_info.csv"), stringsAsFactors = FALSE)
gene_info <- read.csv(file.path(atlas_dir, "genes_info.csv"), stringsAsFactors = FALSE)

# fst matrices do not carry feature row names; restore them from their annotation tables.
stopifnot(nrow(tx_counts) == nrow(tx_info), nrow(gene_counts) == nrow(gene_info))
rownames(tx_counts) <- tx_info$transcript_id
rownames(gene_counts) <- gene_info$gene_id

stopifnot(ncol(tx_counts) == nrow(mdata), ncol(gene_counts) == nrow(mdata))

isoforms <- c("ENST00000284551.11", "ENST00000493030.6")
gene_id <- gene_info$gene_id[match("TRIM11", gene_info$gene_name)]
if (length(gene_id) != 1L || is.na(gene_id)) stop("TRIM11 gene ID was not uniquely identified.")
if (!all(isoforms %in% tx_info$transcript_id)) stop("One or more TRIM11 transcript IDs are absent from transcripts_info.csv.")
if (!all(isoforms %in% rownames(tx_counts))) stop("One or more TRIM11 transcript IDs are absent from txi.counts.fst.")
if (!gene_id %in% rownames(gene_counts)) stop("TRIM11 gene ID is absent from gc_txi.counts.fst.")

# Match the atlas analysis convention: log2(CPM + 1).
tx_expr <- log2(sweep(tx_counts, 2, colSums(tx_counts) / 1e6, "/") + 1)
gene_expr <- log2(sweep(gene_counts, 2, colSums(gene_counts) / 1e6, "/") + 1)

sample_info <- mdata %>%
  transmute(
    sample = name,
    pseudotime = as.numeric(pseudotime),
    subtype = Molecular.Type
  ) %>%
  filter(is.finite(pseudotime), !is.na(subtype))

idx <- match(sample_info$sample, mdata$name)
if (anyNA(idx)) stop("Sample identifiers could not be aligned.")

plot_df <- bind_rows(
  data.frame(id = "TRIM11", count = gene_expr[gene_id, idx]),
  data.frame(id = isoforms[1], count = tx_expr[isoforms[1], idx]),
  data.frame(id = isoforms[2], count = tx_expr[isoforms[2], idx])
) %>%
  mutate(
    pseudotime = rep(sample_info$pseudotime, times = 3),
    subtype = rep(sample_info$subtype, times = 3),
    id = factor(id, levels = c("TRIM11", isoforms[1], isoforms[2])),
    subtype = factor(subtype, levels = c("NORMAL", "PRIMARY", "ARPC", "DNPC", "NEPC"))
  )

cor_gene <- cor(sample_info$pseudotime, gene_expr[gene_id, idx], use = "complete.obs", method = "pearson")
cor_iso <- vapply(isoforms, function(x) {
  cor(sample_info$pseudotime, tx_expr[x, idx], use = "complete.obs", method = "pearson")
}, numeric(1))
delta_corr <- max(cor_iso) - min(cor_iso)

palette_subtypes <- c(
  NORMAL = "#25681E",
  PRIMARY = "#F63A21",
  ARPC = "#4889C7",
  DNPC = "#1010C0",
  NEPC = "#93355A"
)

p <- ggplot(plot_df, aes(x = pseudotime, y = count)) +
  geom_smooth(
    method = "gam",
    formula = y ~ s(x, bs = "cs"),
    color = "grey35",
    fill = "grey80",
    linewidth = 0.25,
    alpha = 0.55,
    na.rm = TRUE
  ) +
  geom_point(
    aes(fill = subtype),
    shape = 21,
    size = 0.85,
    stroke = 0.12,
    alpha = 0.9,
    na.rm = TRUE
  ) +
  facet_wrap(~ id, nrow = 1, scales = "free_y") +
  scale_fill_manual(values = palette_subtypes, drop = FALSE, name = "subtype") +
  labs(
    title = paste0(
      "TRIM11\n",
      "Delta corr= ", formatC(delta_corr, format = "f", digits = 2), "\n",
      "Gene corr= ", formatC(cor_gene, format = "f", digits = 2)
    ),
    x = "pseudotime",
    y = "count"
  ) +
  theme_classic(base_size = 5) +
  theme(
    plot.title = element_text(size = 5, face = "plain", lineheight = 0.9, hjust = 0),
    axis.title = element_text(size = 5),
    axis.text = element_text(size = 5),
    strip.background = element_rect(fill = "white", colour = "black", linewidth = 0.25),
    strip.text = element_text(size = 5, face = "plain"),
    panel.border = element_rect(colour = "black", fill = NA, linewidth = 0.25),
    panel.spacing = grid::unit(0.8, "mm"),
    legend.title = element_text(size = 5),
    legend.text = element_text(size = 5),
    legend.key.size = grid::unit(2.5, "mm"),
    legend.spacing.y = grid::unit(0, "mm"),
    plot.margin = margin(1.5, 1.5, 1.5, 1.5, unit = "mm")
  )

# Use the base R PDF device with a standard PostScript font.  This avoids
# cairo's per-glyph text encoding, which can produce abnormal inter-letter
# spacing in some PDF viewers while leaving the figure geometry unchanged.
pdf(out_pdf, width = 9 / 2.54, height = 6 / 2.54, family = "Helvetica", useDingbats = FALSE)
print(p)
dev.off()
ggsave(out_png, p, width = 9, height = 6, units = "cm", dpi = 600, limitsize = FALSE)

write.csv(
  data.frame(
    gene = "TRIM11",
    transcript_A = isoforms[1],
    transcript_B = isoforms[2],
    gene_correlation = cor_gene,
    transcript_A_correlation = cor_iso[1],
    transcript_B_correlation = cor_iso[2],
    delta_correlation = delta_corr,
    n_samples = nrow(sample_info)
  ),
  "__PCATLAS_PROJECT_ROOT__/TRIM11_isoforms_vs_pseudotime_5B_style_statistics.csv",
  row.names = FALSE
)

message("Wrote: ", out_pdf)
message("Wrote: ", out_png)
```
