## Supplementary material for "An integrated, transcript-resolution atlas of prostate cancer progression unmasks associations with oncogenic pathways": Reproducible code: reproduce_firre_fendrr_lasso.R

```
# Public submission copy.
# Analytical statements and parameter values are unchanged from the archived source.
# Legacy non-English comments and console messages were normalized for public release.
suppressPackageStartupMessages({
  library(Biostrings)
  library(glmnet)
  library(dplyr)
})

source_dir <- "__PCATLAS_DOCUMENTS_ROOT__/Data_Harmanization"
output_dir <- "__PCATLAS_PROJECT_ROOT__/reproduced_supplementary_tables_v2"
dir.create(output_dir, recursive = TRUE, showWarnings = FALSE)

pheno_all <- read.csv(file.path(source_dir, "lncRNA_with_pseudotime_corr.csv"), stringsAsFactors = FALSE) %>%
  filter(!is.na(cor_pseudotime))

run_exact_model <- function(target_gene, kmer_length, min_prevalence = 0.2) {
  gene_lnc <- pheno_all %>% filter(gene_name == target_gene)
  seqs <- readDNAStringSet(file.path(source_dir, "lncRNA_FASTA_output", paste0(target_gene, "_transcripts.fasta")))
  names(seqs) <- sapply(strsplit(names(seqs), "\\|"), `[`, 1)

  kmer_matrix <- oligonucleotideFrequency(seqs, width = kmer_length, as.prob = TRUE)
  prevalence <- colMeans(kmer_matrix > 0)
  kmer_matrix_filtered <- kmer_matrix[, prevalence >= min_prevalence, drop = FALSE]
  kmer_df <- as.data.frame(kmer_matrix_filtered)
  kmer_df$transcript_id <- names(seqs)
  model_data <- inner_join(gene_lnc[, c("transcript_id", "cor_pseudotime")], kmer_df, by = "transcript_id")

  X <- as.matrix(model_data %>% select(-transcript_id, -cor_pseudotime))
  Y <- model_data$cor_pseudotime
  set.seed(42)
  n_folds <- min(10, nrow(X))
  cv_fit <- cv.glmnet(X, Y, alpha = 1, family = "gaussian", nfolds = n_folds)
  coef_matrix <- coef(cv_fit, s = cv_fit$lambda.min)
  coef_df <- data.frame(
    gene = target_gene,
    k = kmer_length,
    K_mer = rownames(coef_matrix),
    Coefficient = as.vector(coef_matrix),
    stringsAsFactors = FALSE
  ) %>%
    filter(Coefficient != 0, K_mer != "(Intercept)") %>%
    arrange(desc(abs(Coefficient))) %>%
    mutate(abs_rank = row_number(), direction = ifelse(Coefficient > 0, "positive", "negative"))

  model_info <- data.frame(
    gene = target_gene,
    k = kmer_length,
    min_prevalence = min_prevalence,
    n_transcripts = nrow(X),
    n_kmers_before_filter = ncol(kmer_matrix),
    n_kmers_after_filter = ncol(X),
    nfolds = n_folds,
    seed = 42,
    alpha = 1,
    family = "gaussian",
    lambda_min = cv_fit$lambda.min,
    lambda_1se = cv_fit$lambda.1se,
    glmnet_version = as.character(packageVersion("glmnet")),
    Biostrings_version = as.character(packageVersion("Biostrings")),
    R_version = R.version.string,
    stringsAsFactors = FALSE
  )

  write.csv(coef_df, file.path(output_dir, paste0(target_gene, "_LASSO_coefficients_exact_reproduction.csv")), row.names = FALSE)
  write.csv(model_info, file.path(output_dir, paste0(target_gene, "_LASSO_model_metadata_exact_reproduction.csv")), row.names = FALSE)
  list(coefficients = coef_df, metadata = model_info)
}

firre <- run_exact_model("FIRRE", 10)
fendrr <- run_exact_model("FENDRR", 5)
combined <- bind_rows(firre$coefficients, fendrr$coefficients)
write.csv(combined, file.path(output_dir, "FIRRE_FENDRR_LASSO_coefficients_exact_reproduction.csv"), row.names = FALSE)

print(firre$metadata)
print(firre$coefficients)
print(fendrr$metadata)
print(fendrr$coefficients)
```
