## Supplementary material for "An integrated, transcript-resolution atlas of prostate cancer progression unmasks associations with oncogenic pathways": Reproducible code: run_reviewer_statistics.R

```
#!/usr/bin/env Rscript
# Public submission copy.
# Analytical statements and parameter values are unchanged from the archived source.
# Legacy non-English comments and console messages were normalized for public release.
suppressPackageStartupMessages({
  library(fst)
  library(survival)
})

atlas_dir <- "__PCATLAS_DOCUMENTS_ROOT__/Data_Harmanization"
pdx_dir <- "__PCATLAS_DOCUMENTS_ROOT__/PDX"
out_dir <- "__PCATLAS_PROJECT_ROOT__/reviewer_response_analysis"
dir.create(out_dir, recursive = TRUE, showWarnings = FALSE)

message("Loading metadata and annotations")
meta <- read.csv(file.path(atlas_dir, "filtered_samples.csv"))
tx_info <- read.csv(file.path(atlas_dir, "transcripts_info.csv"))
g_info <- read.csv(file.path(atlas_dir, "genes_info.csv"))
tx_prod <- read.table(file.path(pdx_dir, "metadata/transcript_protein.tsv"), sep = "\t", quote = "", fill = TRUE,
                      col.names = c("transcript_id", "protein_id", "transcript_name", "CDS"))
tx_info <- merge(tx_info, tx_prod, by = "transcript_id", all.x = TRUE, sort = FALSE)
tx_info <- tx_info[match(read.csv(file.path(atlas_dir, "transcripts_info.csv"))$transcript_id, tx_info$transcript_id), ]

paired_idx <- which(meta$Library.layout == "PAIRED")
sample_cols <- paste0("id.", meta$id[paired_idx])
subtype <- meta$Molecular.Type[paired_idx]
n <- length(paired_idx)
message("Paired samples: ", n, "; NEPC: ", sum(subtype == "NEPC"))

message("Loading transcript counts")
tx_df <- read_fst(file.path(atlas_dir, "txi.counts.fst"), columns = sample_cols)
tx_mat <- as.matrix(tx_df)
rm(tx_df); gc()
rownames(tx_mat) <- tx_info$transcript_id

count_keep <- tx_info$transcript_type == "protein_coding" & rowSums(tx_mat > 1) > n * 0.05
tx_mat <- tx_mat[count_keep, , drop = FALSE]
tx_ann <- tx_info[count_keep, , drop = FALSE]
message("Protein-coding transcripts passing count filter: ", nrow(tx_mat))

message("Loading gene counts and calculating transcript contributions")
gene_df <- read_fst(file.path(atlas_dir, "gc_txi.counts.fst"), columns = sample_cols)
gene_mat <- as.matrix(gene_df)
rm(gene_df); gc()
rownames(gene_mat) <- g_info$gene_id
gene_index <- match(tx_ann$gene_id, rownames(gene_mat))
stopifnot(!anyNA(gene_index))
ratio <- tx_mat / gene_mat[gene_index, , drop = FALSE]
ratio[!is.finite(ratio)] <- 0
contrib_keep <- rowSums(ratio > 0.4) > n * 0.05
ratio <- ratio[contrib_keep, , drop = FALSE]
tx_mat <- tx_mat[contrib_keep, , drop = FALSE]
tx_ann <- tx_ann[contrib_keep, , drop = FALSE]
variable_keep <- apply(ratio, 1, sd) > 0
ratio <- ratio[variable_keep, , drop = FALSE]
tx_mat <- tx_mat[variable_keep, , drop = FALSE]
tx_ann <- tx_ann[variable_keep, , drop = FALSE]
rm(gene_mat); gc()
message("Eligible transcripts after contribution filter: ", nrow(ratio))

message("Loading pathway scores")
env <- new.env(parent = emptyenv())
load(file.path(atlas_dir, "Full_GSVA_Analysis_Workspace_2026-03-23.Rdata"), envir = env)
gsva <- env$full_gsva_res
rm(env); gc()
reported <- read.csv(file.path(pdx_dir, "results_06_IsoformSwitch/IsoformSwitchAll/all_results.csv"))
reported_pairs <- unique(reported[c("gene_id", "target")])
pathways <- intersect(unique(reported$target), rownames(gsva))
Y <- gsva[pathways, sample_cols, drop = FALSE]
Y <- rbind(Y, pseudotime = meta$pseudotime[paired_idx])
Y <- Y[apply(Y, 1, sd, na.rm = TRUE) > 0, , drop = FALSE]
pathways <- rownames(Y)
reported_pairs_analyzed <- reported_pairs[reported_pairs$target %in% pathways, , drop = FALSE]
rm(gsva); gc()

scale_rows <- function(x) {
  mu <- rowMeans(x)
  ss <- sqrt(rowSums((x - mu)^2))
  (x - mu) / ss
}

message("Computing transcript-target correlations: ", nrow(ratio), " x ", nrow(Y))
Xz <- scale_rows(ratio)
Yz <- scale_rows(Y)
C <- Xz %*% t(Yz)
rm(Xz, Yz); gc()
colnames(C) <- pathways
rownames(C) <- tx_ann$transcript_id

# Two-sided Pearson p values and global BH across the full transcript-target family.
Tstat <- C * sqrt((n - 2) / pmax(1e-15, 1 - C^2))
P <- 2 * pt(-abs(Tstat), df = n - 2)
Q <- matrix(p.adjust(as.vector(P), method = "BH"), nrow = nrow(P), ncol = ncol(P),
            dimnames = dimnames(P))
rm(Tstat, P); gc()

gene_groups <- split(seq_len(nrow(tx_ann)), tx_ann$gene_id)

product_count <- function(idx) {
  cds <- tx_ann$CDS[idx]
  prot <- tx_ann$protein_id[idx]
  # Deliberately mirror the discovery script, including its treatment of NA as a unique value.
  if (any(is.na(cds) | cds == "")) length(unique(prot)) else length(unique(cds))
}

call_switches <- function(cormat, threshold, qmat = NULL, require_fdr = FALSE) {
  ans <- vector("list", length(gene_groups))
  ai <- 0L
  for (gene in names(gene_groups)) {
    idx <- gene_groups[[gene]]
    if (length(idx) < 2L) next
    cg <- cormat[idx, , drop = FALSE]
    mx <- apply(cg, 2, max, na.rm = TRUE)
    mn <- apply(cg, 2, min, na.rm = TRUE)
    candidates <- which(mx > threshold & mn < -threshold)
    if (!length(candidates)) next
    for (j in candidates) {
      qual <- idx[abs(cormat[idx, j]) > threshold]
      if (length(qual) < 2L || product_count(qual) <= 1L) next
      pos_local <- idx[which.max(cormat[idx, j])]
      neg_local <- idx[which.min(cormat[idx, j])]
      pair_q <- if (is.null(qmat)) NA_real_ else max(qmat[c(pos_local, neg_local), j], na.rm = TRUE)
      if (require_fdr && !(is.finite(pair_q) && pair_q < 0.05)) next
      ai <- ai + 1L
      ans[[ai]] <- data.frame(gene_id = gene, target = colnames(cormat)[j],
                              max_corr = mx[j], min_corr = mn[j], delta_corr = mx[j] - mn[j],
                              iso_pos = rownames(cormat)[pos_local], iso_neg = rownames(cormat)[neg_local],
                              pair_q = pair_q)
    }
  }
  if (!ai) return(data.frame())
  do.call(rbind, ans[seq_len(ai)])
}

thresholds <- c(0.40, 0.45, 0.50, 0.55, 0.60)
switch_lists <- list()
sens <- data.frame()
for (thr in thresholds) {
  message("Calling switches at |r| > ", thr)
  sw <- call_switches(C, thr)
  switch_lists[[as.character(thr)]] <- sw
  key <- paste(sw$gene_id, sw$target, sep = "|")
  base_reported <- paste(reported_pairs_analyzed$gene_id, reported_pairs_analyzed$target, sep = "|")
  sens <- rbind(sens, data.frame(threshold = thr, n_switches = nrow(sw), n_genes = length(unique(sw$gene_id)),
                                 overlap_reported = sum(key %in% base_reported),
                                 reported_covered = mean(base_reported %in% key)))
  write.csv(sw, file.path(out_dir, sprintf("isoform_switches_r%.2f.csv", thr)), row.names = FALSE)
}

message("Applying global transcript-target BH-FDR at baseline threshold")
sw_fdr <- call_switches(C, 0.50, qmat = Q, require_fdr = TRUE)
write.csv(sw_fdr, file.path(out_dir, "isoform_switches_r0.50_global_BH.csv"), row.names = FALSE)
sens$global_BH_pair_q_lt_0.05 <- NA_integer_
sens$global_BH_pair_q_lt_0.05[sens$threshold == 0.50] <- nrow(sw_fdr)
write.csv(sens, file.path(out_dir, "isoform_switch_threshold_sensitivity.csv"), row.names = FALSE)

message("NEPC exclusion analysis")
no_nepc <- subtype != "NEPC"
n2 <- sum(no_nepc)
C_no_nepc <- scale_rows(ratio[, no_nepc, drop = FALSE]) %*% t(scale_rows(Y[, no_nepc, drop = FALSE]))
colnames(C_no_nepc) <- pathways
rownames(C_no_nepc) <- tx_ann$transcript_id
sw_no_nepc <- call_switches(C_no_nepc, 0.50)
write.csv(sw_no_nepc, file.path(out_dir, "isoform_switches_r0.50_excluding_NEPC.csv"), row.names = FALSE)

baseline <- switch_lists[["0.5"]]
base_key <- paste(baseline$gene_id, baseline$target, sep = "|")
nonepc_key <- paste(sw_no_nepc$gene_id, sw_no_nepc$target, sep = "|")
key_genes <- c("WNT2B", "RNF43", "RELA", "BRCA1", "ROCK2", "TRIM11")
gene_name_map <- setNames(g_info$gene_name, g_info$gene_id)
baseline$gene_name <- gene_name_map[baseline$gene_id]
key_base <- baseline[baseline$gene_name %in% key_genes, ]
key_base$retained_excluding_NEPC <- paste(key_base$gene_id, key_base$target, sep = "|") %in% nonepc_key
write.csv(key_base, file.path(out_dir, "key_switches_NEPC_exclusion.csv"), row.names = FALSE)
nepc_summary <- data.frame(
  analysis = c("paired_atlas", "baseline_switches", "baseline_retained_excluding_NEPC", "key_gene_switches", "key_gene_retained_excluding_NEPC"),
  value = c(n, length(base_key), sum(base_key %in% nonepc_key), nrow(key_base), sum(key_base$retained_excluding_NEPC)),
  denominator = c(NA, NA, length(base_key), NA, nrow(key_base))
)
write.csv(nepc_summary, file.path(out_dir, "NEPC_exclusion_summary.csv"), row.names = FALSE)

message("Molecular-state-balanced sensitivity analysis")
weighted_scale_rows <- function(x, w) {
  w <- w / sum(w)
  mu <- as.numeric(x %*% w)
  centered <- x - mu
  denom <- sqrt(rowSums(centered^2 * rep(w, each = nrow(x))))
  centered * rep(sqrt(w), each = nrow(x)) / denom
}
state_weights <- 1 / as.numeric(table(subtype)[subtype])
C_balanced <- weighted_scale_rows(ratio, state_weights) %*%
  t(weighted_scale_rows(Y, state_weights))
colnames(C_balanced) <- pathways
rownames(C_balanced) <- tx_ann$transcript_id
sw_balanced <- call_switches(C_balanced, 0.50)
write.csv(sw_balanced, file.path(out_dir, "isoform_switches_r0.50_molecular_state_balanced.csv"), row.names = FALSE)
balanced_key <- paste(sw_balanced$gene_id, sw_balanced$target, sep = "|")
key_balanced <- key_base
key_balanced$retained_state_balanced <- paste(key_balanced$gene_id, key_balanced$target, sep = "|") %in% balanced_key
write.csv(key_balanced, file.path(out_dir, "key_switches_state_balanced.csv"), row.names = FALSE)
balanced_summary <- data.frame(
  analysis = c("molecular_states", "balanced_switches", "baseline_retained_state_balanced",
               "key_gene_switches", "key_gene_retained_state_balanced"),
  value = c(length(unique(subtype)), length(balanced_key), sum(base_key %in% balanced_key),
            nrow(key_balanced), sum(key_balanced$retained_state_balanced)),
  denominator = c(NA, NA, length(base_key), NA, nrow(key_balanced))
)
write.csv(balanced_summary, file.path(out_dir, "molecular_state_balanced_summary.csv"), row.names = FALSE)

# Isolate the effect of NEPC representation by increasing its total analytic weight
# while retaining the observed relative weights among all non-NEPC samples.
nepc_weight_results <- list()
for (target_fraction in c(0.10, 0.20)) {
  w_nepc <- ifelse(subtype == "NEPC",
                   target_fraction / sum(subtype == "NEPC"),
                   (1 - target_fraction) / sum(subtype != "NEPC"))
  C_nepc_weighted <- weighted_scale_rows(ratio, w_nepc) %*%
    t(weighted_scale_rows(Y, w_nepc))
  colnames(C_nepc_weighted) <- pathways
  rownames(C_nepc_weighted) <- tx_ann$transcript_id
  sw_nepc_weighted <- call_switches(C_nepc_weighted, 0.50)
  weighted_key <- paste(sw_nepc_weighted$gene_id, sw_nepc_weighted$target, sep = "|")
  tag <- sprintf("%02d", round(100 * target_fraction))
  write.csv(sw_nepc_weighted,
            file.path(out_dir, paste0("isoform_switches_r0.50_NEPC_weight_", tag, "pct.csv")),
            row.names = FALSE)
  key_tmp <- key_base
  key_tmp$retained_NEPC_weighted <- paste(key_tmp$gene_id, key_tmp$target, sep = "|") %in% weighted_key
  write.csv(key_tmp, file.path(out_dir, paste0("key_switches_NEPC_weight_", tag, "pct.csv")), row.names = FALSE)
  nepc_weight_results[[tag]] <- data.frame(
    target_NEPC_weight = target_fraction,
    n_switches = length(weighted_key),
    baseline_retained = sum(base_key %in% weighted_key),
    baseline_denominator = length(base_key),
    key_retained = sum(key_tmp$retained_NEPC_weighted),
    key_denominator = nrow(key_tmp)
  )
}
nepc_weight_summary <- do.call(rbind, nepc_weight_results)
write.csv(nepc_weight_summary, file.path(out_dir, "NEPC_upweighting_summary.csv"), row.names = FALSE)

message("Kendall multiplicity")
kendall <- read.csv("__PCATLAS_DOCUMENTS_ROOT__/WebToolStudy/kendall_tau_results_pc_isoform.csv")
kendall$BH_FDR <- p.adjust(kendall$p_value, method = "BH")
write.csv(kendall, file.path(out_dir, "kendall_tau_results_with_BH_FDR.csv"), row.names = FALSE)
kendall_summary <- data.frame(
  family = "all protein-coding transcripts",
  n_tests = nrow(kendall),
  nominal_p_lt_0.05 = sum(kendall$p_value < 0.05, na.rm = TRUE),
  BH_FDR_lt_0.05 = sum(kendall$BH_FDR < 0.05, na.rm = TRUE),
  BH_FDR_lt_0.01 = sum(kendall$BH_FDR < 0.01, na.rm = TRUE)
)
write.csv(kendall_summary, file.path(out_dir, "kendall_multiplicity_summary.csv"), row.names = FALSE)

message("Survival multiplicity")
surv_meta <- read.csv("__PCATLAS_DOCUMENTS_ROOT__/WebToolStudy/combined_all_survival_updated_v5.csv", check.names = FALSE)
surv_meta$id <- as.character(surv_meta$id)
targets <- data.frame(
  Gene = rep(c("WNT2B", "TRIM11", "ROCK2", "RNF43", "CDK12", "F8", "RELA", "BRCA1"), each = 2),
  ENST = c("ENST00000369684", "ENST00000369686", "ENST00000284551", "ENST00000493030",
           "ENST00000315872", "ENST00000401753", "ENST00000584437", "ENST00000583753",
           "ENST00000447079", "ENST00000584632", "ENST00000360256", "ENST00000330287",
           "ENST00000406246", "ENST00000612991", "ENST00000357654", "ENST00000352993")
)
all_sample_cols <- paste0("id.", meta$id)
lib <- meta$totalcounts
tx_base <- sub("\\..*$", "", tx_info$transcript_id)
surv_res <- list()
si <- 0L
for (i in seq_len(nrow(targets))) {
  idx <- match(targets$ENST[i], tx_base)
  stopifnot(!is.na(idx))
  expr <- as.numeric(read_fst(file.path(atlas_dir, "txi.counts.fst"), from = idx, to = idx)[1, ])
  # Match the original script's log2(CPM+1) and all-atlas median grouping.
  val <- log2(expr / (lib / 1e6) + 1)
  ex <- data.frame(id = as.character(meta$id), Expression = val)
  dat <- merge(surv_meta, ex, by = "id")
  med <- median(dat$Expression, na.rm = TRUE)
  dat$Expr_Group <- if (med == 0) ifelse(dat$Expression > 0, "High", "Low") else ifelse(dat$Expression >= med, "High", "Low")
  for (endpoint in c("PRIMARY_PFS", "CRPC_OS")) {
    if (endpoint == "PRIMARY_PFS") {
      dd <- dat[dat[["Molecular Type"]] == "PRIMARY" & !is.na(dat$PFS_Time) & dat$PFS_Time > 0, ]
      time <- dd$PFS_Time; event <- dd$PFS_Event
    } else {
      dd <- dat[dat[["Molecular Type"]] %in% c("mARPC", "mDNPC", "mNEPC") & !is.na(dat$OS_Time) & dat$OS_Time > 0, ]
      time <- dd$OS_Time; event <- dd$OS_Event
    }
    if (length(unique(dd$Expr_Group)) < 2L) next
    dd$TIME <- time
    dd$EVENT <- event
    fit <- survdiff(Surv(TIME, EVENT) ~ Expr_Group, data = dd)
    p <- pchisq(fit$chisq, df = length(fit$n) - 1, lower.tail = FALSE)
    si <- si + 1L
    surv_res[[si]] <- data.frame(Gene = targets$Gene[i], ENST = targets$ENST[i], endpoint = endpoint,
                                 n = nrow(dd), events = sum(event == 1, na.rm = TRUE), logrank_p = p)
  }
}
surv_res <- do.call(rbind, surv_res)
surv_res$BH_within_endpoint <- ave(surv_res$logrank_p, surv_res$endpoint, FUN = function(x) p.adjust(x, "BH"))
surv_res$BH_across_all_32 <- p.adjust(surv_res$logrank_p, "BH")
write.csv(surv_res, file.path(out_dir, "survival_logrank_multiplicity.csv"), row.names = FALSE)

summary_lines <- c(
  paste("Paired atlas samples:", n, "including NEPC:", sum(subtype == "NEPC")),
  paste("Eligible transcripts:", nrow(ratio), "targets analyzed:", nrow(Y)),
  paste("Baseline r=0.50 switches:", nrow(baseline)),
  paste("Baseline switches passing global transcript-target BH pair-q<0.05:", nrow(sw_fdr)),
  paste("Baseline switches retained after excluding NEPC:", sum(base_key %in% nonepc_key), "/", length(base_key)),
  paste("Baseline switches retained with equal molecular-state weighting:", sum(base_key %in% balanced_key), "/", length(base_key)),
  paste("Baseline switches retained when NEPC has 20% total weight:",
        nepc_weight_summary$baseline_retained[nepc_weight_summary$target_NEPC_weight == 0.20], "/", length(base_key)),
  paste("Kendall BH q<0.05:", kendall_summary$BH_FDR_lt_0.05, "/", kendall_summary$n_tests),
  paste("Survival BH q<0.05 within endpoint:", sum(surv_res$BH_within_endpoint < 0.05), "/", nrow(surv_res))
)
writeLines(summary_lines, file.path(out_dir, "analysis_summary.txt"))
cat(paste(summary_lines, collapse = "\n"), "\n")
```
