## Supplementary material for "An integrated, transcript-resolution atlas of prostate cancer progression unmasks associations with oncogenic pathways": Reproducible code: filter_four_way_T_supported_GO_BP.R

```
# Public submission copy.
# Analytical statements and parameter values are unchanged from the archived source.
# Legacy non-English comments and console messages were normalized for public release.
suppressPackageStartupMessages(library(dplyr))

script_arg <- grep("^--file=", commandArgs(), value = TRUE)
script_dir <- if (length(script_arg)) {
  dirname(normalizePath(sub("^--file=", "", script_arg)))
} else {
  normalizePath(getwd())
}

atlas_experiment <- read.delim(
  file.path(script_dir, "BRCA1_C_T_significant_opposite_GO_BP_with_experimental_RNAseq.tsv"),
  check.names = FALSE,
  stringsAsFactors = FALSE
)

bioid <- read.delim(
  file.path(script_dir, "BioID_BRCA1_T_GO_BP_GSEA_all.tsv"),
  check.names = FALSE,
  stringsAsFactors = FALSE
) %>%
  transmute(
    pathway,
    BioID_T_NES = NES,
    BioID_T_FDR = padj,
    BioID_T_effective_size = size,
    BioID_T_leading_edge = leadingEdge,
    BioID_T_direction = direction
  )

localization_file <- file.path(
  dirname(script_dir),
  "bioID_BRCA", "BRCA1_T_cytoplasmic_reanalysis",
  "BRCA1_T_GO_screened_UniProt_localization_verification.tsv"
)
verified_localization <- read.delim(
  localization_file,
  check.names = FALSE,
  stringsAsFactors = FALSE
) %>%
  filter(final_non_nuclear_signature) %>%
  distinct(gene, uniprot_location_summary)

verified_non_nuclear <- verified_localization$gene
verified_location_lookup <- setNames(
  verified_localization$uniprot_location_summary,
  verified_localization$gene
)

extract_verified_non_nuclear <- function(leading_edge) {
  if (is.na(leading_edge) || !nzchar(leading_edge)) return(character())
  intersect(strsplit(leading_edge, ";", fixed = TRUE)[[1]], verified_non_nuclear)
}

format_verified_details <- function(genes) {
  if (!length(genes)) return("")
  paste0(genes, " [", verified_location_lookup[genes], "]", collapse = "; ")
}

annotated <- atlas_experiment %>%
  left_join(bioid, by = "pathway") %>%
  mutate(
    RNAseq_same_as_Atlas_T = sign(Experiment_T_OE_NES) == sign(Atlas_T_NES),
    BioID_same_as_Atlas_T = sign(BioID_T_NES) == sign(Atlas_T_NES),
    RNAseq_significant = !is.na(Experiment_T_OE_FDR) & Experiment_T_OE_FDR < 0.05,
    BioID_significant = !is.na(BioID_T_FDR) & BioID_T_FDR < 0.05,
    All_four_FDR_lt_0.05 = Atlas_T_FDR < 0.05 & Atlas_C_FDR < 0.05 &
      RNAseq_significant & BioID_significant,
    worst_FDR_four_way = pmax(
      Atlas_T_FDR, Atlas_C_FDR, Experiment_T_OE_FDR, BioID_T_FDR,
      na.rm = TRUE
    ),
    mean_abs_NES = rowMeans(
      abs(cbind(Atlas_T_NES, Atlas_C_NES, Experiment_T_OE_NES, BioID_T_NES)),
      na.rm = TRUE
    ),
    BioID_verified_non_nuclear_proteins = vapply(
      BioID_T_leading_edge,
      function(x) paste(extract_verified_non_nuclear(x), collapse = ";"),
      FUN.VALUE = character(1)
    ),
    BioID_verified_non_nuclear_count = vapply(
      BioID_T_leading_edge,
      function(x) length(extract_verified_non_nuclear(x)),
      FUN.VALUE = integer(1)
    ),
    BioID_verified_non_nuclear_details = vapply(
      BioID_T_leading_edge,
      function(x) format_verified_details(extract_verified_non_nuclear(x)),
      FUN.VALUE = character(1)
    ),
    Experimental_assay_alignment = case_when(
      RNAseq_same_as_Atlas_T & BioID_same_as_Atlas_T ~ "Both align with Atlas-T",
      sign(Experiment_T_OE_NES) == sign(Atlas_C_NES) &
        sign(BioID_T_NES) == sign(Atlas_C_NES) ~ "Both align with Atlas-C",
      TRUE ~ "RNA-seq and BioID discordant"
    )
  )

direction_matched <- annotated %>%
  filter(RNAseq_same_as_Atlas_T, BioID_same_as_Atlas_T) %>%
  arrange(pmax(Experiment_T_OE_FDR, BioID_T_FDR), worst_FDR_four_way)

four_way <- direction_matched %>%
  filter(
    RNAseq_significant,
    BioID_significant
  ) %>%
  arrange(desc(All_four_FDR_lt_0.05), worst_FDR_four_way, desc(mean_abs_NES), pathway) %>%
  select(
    pathway, pathway_label,
    Atlas_T_NES, Atlas_T_FDR,
    Atlas_C_NES, Atlas_C_FDR,
    Experiment_T_OE_NES, Experiment_T_OE_FDR,
    BioID_T_NES, BioID_T_FDR,
    BioID_verified_non_nuclear_count,
    BioID_verified_non_nuclear_proteins,
    BioID_verified_non_nuclear_details,
    BioID_T_leading_edge,
    All_four_FDR_lt_0.05, worst_FDR_four_way, mean_abs_NES,
    Atlas_T_effective_size, Atlas_C_effective_size,
    Experiment_T_OE_effective_size, BioID_T_effective_size
  )

strict_four <- four_way %>% filter(All_four_FDR_lt_0.05)

both_assays_significant_audit <- annotated %>%
  filter(RNAseq_significant, BioID_significant) %>%
  arrange(Experimental_assay_alignment, worst_FDR_four_way, pathway)

relaxed_rnaseq_cytoplasmic_shortlist <- annotated %>%
  filter(
    RNAseq_same_as_Atlas_T,
    RNAseq_significant,
    BioID_verified_non_nuclear_count >= 1
  ) %>%
  arrange(
    Experiment_T_OE_FDR,
    max_Atlas_FDR,
    desc(BioID_verified_non_nuclear_count),
    pathway
  ) %>%
  select(
    pathway, pathway_label,
    Atlas_T_NES, Atlas_T_FDR,
    Atlas_C_NES, Atlas_C_FDR,
    Experiment_T_OE_NES, Experiment_T_OE_FDR,
    BioID_T_NES, BioID_T_FDR, BioID_T_direction,
    BioID_verified_non_nuclear_count,
    BioID_verified_non_nuclear_proteins,
    BioID_verified_non_nuclear_details,
    BioID_T_leading_edge
  )

exploratory_three_fdr_0.10_shortlist <- annotated %>%
  filter(
    Atlas_T_FDR < 0.10,
    Atlas_C_FDR < 0.10,
    Experiment_T_OE_FDR < 0.10,
    RNAseq_same_as_Atlas_T,
    BioID_verified_non_nuclear_count >= 1
  ) %>%
  arrange(
    Experiment_T_OE_FDR,
    max_Atlas_FDR,
    desc(BioID_verified_non_nuclear_count),
    pathway
  ) %>%
  select(
    pathway, pathway_label,
    Atlas_T_NES, Atlas_T_FDR,
    Atlas_C_NES, Atlas_C_FDR,
    Experiment_T_OE_NES, Experiment_T_OE_FDR,
    BioID_T_NES, BioID_T_FDR, BioID_T_direction,
    BioID_verified_non_nuclear_count,
    BioID_verified_non_nuclear_proteins,
    BioID_verified_non_nuclear_details,
    BioID_T_leading_edge
  )

write.table(
  four_way,
  file.path(script_dir, "BRCA1_GO_BP_T_direction_supported_by_RNAseq_and_BioID.tsv"),
  sep = "\t", quote = FALSE, row.names = FALSE
)

write.table(
  strict_four,
  file.path(script_dir, "BRCA1_GO_BP_all_four_FDR_lt_0.05_T_direction_supported.tsv"),
  sep = "\t", quote = FALSE, row.names = FALSE
)

write.table(
  direction_matched,
  file.path(script_dir, "BRCA1_GO_BP_RNAseq_BioID_same_as_Atlas_T_any_significance.tsv"),
  sep = "\t", quote = FALSE, row.names = FALSE
)

write.table(
  both_assays_significant_audit,
  file.path(script_dir, "BRCA1_GO_BP_RNAseq_and_BioID_both_FDR_lt_0.05_direction_audit.tsv"),
  sep = "\t", quote = FALSE, row.names = FALSE
)

write.table(
  relaxed_rnaseq_cytoplasmic_shortlist,
  file.path(script_dir, "BRCA1_GO_BP_relaxed_RNAseq_supported_with_BioID_non_nuclear_protein.tsv"),
  sep = "\t", quote = FALSE, row.names = FALSE
)

write.table(
  exploratory_three_fdr_0.10_shortlist,
  file.path(script_dir, "BRCA1_GO_BP_three_contrasts_FDR_lt_0.10_with_BioID_non_nuclear_protein.tsv"),
  sep = "\t", quote = FALSE, row.names = FALSE
)

cat("T-direction supported by significant RNA-seq and BioID:", nrow(four_way), "\n")
cat("All four comparisons FDR < 0.05:", nrow(strict_four), "\n")
cat("Same T direction regardless of experimental significance:", nrow(direction_matched), "\n")
cat("RNA-seq and BioID both FDR < 0.05 regardless of direction:", nrow(both_assays_significant_audit), "\n")
cat("Relaxed RNA-seq-supported shortlist with >=1 verified non-nuclear BioID protein:", nrow(relaxed_rnaseq_cytoplasmic_shortlist), "\n")
cat("Three contrasts FDR < 0.10 with >=1 verified non-nuclear BioID protein:", nrow(exploratory_three_fdr_0.10_shortlist), "\n")
print(four_way %>% select(
  pathway_label,
  Atlas_T_NES, Atlas_T_FDR,
  Atlas_C_NES, Atlas_C_FDR,
  Experiment_T_OE_NES, Experiment_T_OE_FDR,
  BioID_T_NES, BioID_T_FDR,
  BioID_verified_non_nuclear_count,
  BioID_verified_non_nuclear_proteins,
  All_four_FDR_lt_0.05
) %>% head(30), row.names = FALSE)
```
