## Supplementary material for "An integrated, transcript-resolution atlas of prostate cancer progression unmasks associations with oncogenic pathways": Reproducible code: plot_BRCA1_GO_BP_bipartite_network_v5_compact.R

```
#!/usr/bin/env Rscript
# Public submission copy.
# Analytical statements and parameter values are unchanged from the archived source.
# Legacy non-English comments and console messages were normalized for public release.

# Compact 8 x 10 cm version of the final mirrored network. Text remains 5 pt.

script_arg <- grep("^--file=", commandArgs(), value = TRUE)
script_dir <- if (length(script_arg)) {
  dirname(normalizePath(sub("^--file=", "", script_arg)))
} else {
  normalizePath(getwd())
}

source(
  file.path(script_dir, "plot_BRCA1_GO_BP_bipartite_network_v4_final.R"),
  local = TRUE
)

evidence_x <- c("Atlas-T" = -0.13, "Atlas-C" = 0.00, "RNA-seq" = 0.13)
pathway_node_x <- 0.94
pathway_label_x <- 0.86
protein_node_x <- 1.45
protein_label_x <- 1.51

# Recalculate y coordinates with substantially smaller between-module gaps.
pathway_compact <- pathway_final %>%
  arrange(desc(y)) %>%
  mutate(
    row_index = row_number(),
    theme_index = as.integer(theme),
    y_raw_compact = row_index + (theme_index - 1) * 0.28,
    y = max(y_raw_compact) + 1 - y_raw_compact,
    x = pathway_node_x,
    display_label = recode(
      pathway_label,
      "Organic Acid Metabolic Process" = "Organic acid metabolism",
      "Lipid Metabolic Process" = "Lipid metabolism",
      "Carbohydrate Metabolic Process" = "Carbohydrate metabolism",
      "Homeostatic Process" = "Homeostasis",
      "Transmembrane Transport" = "Transmembrane transport",
      "Biological Process Involved in Interspecies Interaction Between Organisms" = "Interspecies interaction",
      "Defense Response to Other Organism" = "Defense response",
      "Negative Regulation of Immune System Process" = "Negative immune regulation",
      "Inflammatory Response" = "Inflammatory response",
      "Regulation of Inflammatory Response" = "Inflammatory regulation",
      "Regulation of Programmed Cell Death" = "Programmed-death regulation",
      "Apoptotic Signaling Pathway" = "Apoptotic signaling",
      "Positive Regulation of Programmed Cell Death" = "Positive cell-death regulation",
      "Mitochondrion Organization" = "Mitochondrion organization",
      "Response to Oxidative Stress" = "Oxidative-stress response",
      "Cellular Response to Oxidative Stress" = "Cellular oxidative response",
      "Regulation of Protein Catabolic Process" = "Protein-catabolism regulation",
      "Positive Regulation of Protein Metabolic Process" = "Positive protein metabolism",
      "Regulation of Cell Population Proliferation" = "Cell-proliferation regulation"
    ),
    label = stringr::str_wrap(display_label, width = 31)
  )

protein_compact <- protein_final %>%
  arrange(desc(y)) %>%
  mutate(
    y = seq(max(pathway_compact$y), min(pathway_compact$y), length.out = n()),
    x = protein_node_x
  )

edges_compact <- edges %>%
  left_join(
    pathway_compact %>% select(pathway, pathway_y = y),
    by = "pathway"
  ) %>%
  left_join(
    protein_compact %>% select(protein = node_id, protein_y = y),
    by = "protein"
  ) %>%
  transmute(
    pathway, protein, BioID_leading_edge,
    x = pathway_node_x, y = pathway_y,
    xend = protein_node_x, yend = protein_y
  )

evidence_compact <- bind_rows(
  pathway_compact %>% transmute(
    pathway, pathway_label, y,
    evidence = "Atlas-T", NES = Atlas_T_NES, FDR = Atlas_T_FDR
  ),
  pathway_compact %>% transmute(
    pathway, pathway_label, y,
    evidence = "Atlas-C", NES = Atlas_C_NES, FDR = Atlas_C_FDR
  ),
  pathway_compact %>% transmute(
    pathway, pathway_label, y,
    evidence = "RNA-seq", NES = RNAseq_NES, FDR = RNAseq_FDR
  )
) %>%
  mutate(
    evidence = factor(evidence, levels = c("Atlas-T", "Atlas-C", "RNA-seq")),
    x = unname(evidence_x[as.character(evidence)]),
    neglog10_FDR = -log10(pmax(FDR, .Machine$double.xmin))
  )

rectangles_compact <- pathway_compact %>%
  group_by(theme) %>%
  summarise(
    ymin = min(y) - 0.31,
    ymax = max(y) + 0.31,
    label_y = (min(y) + max(y)) / 2,
    .groups = "drop"
  ) %>%
  mutate(
    xmin = -0.20,
    xmax = 1.01,
    label_x = -0.255,
    short_label = recode(
      as.character(theme),
      "Metabolism and transport" = "Met",
      "Immune and inflammatory response" = "Imm",
      "Programmed cell death" = "Death",
      "Oxidative stress and mitochondria" = "Ox",
      "Protein homeostasis" = "Prot",
      "Cell proliferation" = "Prolif"
    )
  )

write.table(
  evidence_compact,
  file.path(script_dir, "BRCA1_GO_BP_v5_compact_three_evidence_matrix.tsv"),
  sep = "\t", quote = FALSE, row.names = FALSE
)
write.table(
  edges_compact,
  file.path(script_dir, "BRCA1_GO_BP_v5_compact_edges.tsv"),
  sep = "\t", quote = FALSE, row.names = FALSE
)
write.table(
  pathway_compact,
  file.path(script_dir, "BRCA1_GO_BP_v5_compact_pathway_nodes_with_full_names.tsv"),
  sep = "\t", quote = FALSE, row.names = FALSE
)

base_pt <- 5
label_mm <- base_pt / 2.845276
plot_y_min <- min(pathway_compact$y) - 0.48
plot_y_max <- max(pathway_compact$y) + 1.15
header_y <- max(pathway_compact$y) + 1.00
column_y <- max(pathway_compact$y) + 0.62
evidence_limit <- max(abs(evidence_compact$NES), na.rm = TRUE)

p_compact <- ggplot() +
  geom_rect(
    data = rectangles_compact,
    aes(xmin = xmin, xmax = xmax, ymin = ymin, ymax = ymax, fill = theme),
    alpha = 0.075,
    color = NA,
    show.legend = FALSE
  ) +
  geom_rect(
    data = rectangles_compact,
    aes(xmin = xmin, xmax = xmax, ymin = ymin, ymax = ymax, color = theme),
    fill = NA,
    linewidth = 0.30,
    alpha = 0.62,
    show.legend = FALSE
  ) +
  geom_text(
    data = rectangles_compact,
    aes(x = label_x, y = label_y, label = short_label, color = theme),
    hjust = 1,
    angle = 0,
    family = "Helvetica",
    fontface = "bold",
    size = label_mm,
    show.legend = FALSE
  ) +
  scale_fill_manual(values = module_colors, guide = "none") +
  scale_color_manual(values = module_colors, guide = "none") +
  ggnewscale::new_scale_fill() +
  geom_segment(
    data = edges_compact,
    aes(x = x, y = y, xend = xend, yend = yend),
    linewidth = 0.20,
    color = "#6F7782",
    alpha = 0.40,
    lineend = "round"
  ) +
  geom_point(
    data = pathway_compact,
    aes(x = x, y = y),
    shape = 22,
    size = 2.7,
    fill = "#F2F2F2",
    color = "#333333",
    stroke = 0.30
  ) +
  geom_text(
    data = pathway_compact,
    aes(y = y, label = label),
    x = pathway_label_x,
    hjust = 1,
    lineheight = 0.76,
    size = label_mm,
    family = "Helvetica",
    color = "black"
  ) +
  geom_point(
    data = protein_compact,
    aes(x = x, y = y, size = neglog10_q, fill = log2FC),
    shape = 21,
    color = "#1F2933",
    stroke = 0.23
  ) +
  scale_size_continuous(
    range = c(1.8, 4.8),
    breaks = c(1.5, 2.5),
    name = "P q",
    guide = guide_legend(order = 1, title.position = "top", nrow = 1)
  ) +
  scale_fill_gradient(
    low = "#DCEAF6",
    high = "#2166AC",
    name = "P FC",
    guide = guide_colorbar(
      order = 2,
      direction = "horizontal",
      title.position = "top",
      title.hjust = 0.5,
      barwidth = grid::unit(8, "mm"),
      barheight = grid::unit(2.0, "mm")
    )
  ) +
  ggnewscale::new_scale("size") +
  ggnewscale::new_scale_fill() +
  geom_point(
    data = evidence_compact,
    aes(x = x, y = y, size = neglog10_FDR, fill = NES),
    shape = 21,
    color = "#333333",
    stroke = 0.18
  ) +
  scale_size_continuous(
    range = c(1.2, 4.0),
    breaks = c(2.5, 7.5),
    name = "GO FDR",
    guide = guide_legend(order = 3, title.position = "top", nrow = 1)
  ) +
  scale_fill_gradient2(
    low = "#2166AC",
    mid = "#F7F7F7",
    high = "#B2182B",
    midpoint = 0,
    limits = c(-evidence_limit, evidence_limit),
    name = "GO NES",
    guide = guide_colorbar(
      order = 4,
      direction = "horizontal",
      title.position = "top",
      title.hjust = 0.5,
      barwidth = grid::unit(9, "mm"),
      barheight = grid::unit(2.0, "mm")
    )
  ) +
  geom_text(
    data = protein_compact,
    aes(y = y, label = label),
    x = protein_label_x,
    hjust = 0,
    size = label_mm,
    family = "Helvetica",
    color = "black"
  ) +
  annotate(
    "text", x = mean(evidence_x), y = header_y,
    label = "Evidence",
    family = "Helvetica", fontface = "bold", size = label_mm
  ) +
  annotate(
    "text",
    x = unname(evidence_x), y = column_y,
    label = c("T", "C", "RNA"),
    family = "Helvetica", size = label_mm,
    hjust = 0.5, lineheight = 0.80
  ) +
  annotate(
    "text", x = 0.66, y = header_y,
    label = "GO:BP pathways",
    family = "Helvetica", fontface = "bold", size = label_mm
  ) +
  annotate(
    "text", x = protein_node_x, y = header_y,
    label = "BioID proteins",
    family = "Helvetica", fontface = "bold", size = label_mm
  ) +
  coord_cartesian(
    xlim = c(-0.42, 1.76),
    ylim = c(plot_y_min, plot_y_max),
    clip = "off"
  ) +
  theme_void(base_size = base_pt, base_family = "Helvetica") +
  theme(
    text = element_text(size = base_pt, family = "Helvetica", color = "black"),
    plot.margin = margin(1.3, 1.5, 1.2, 1.5, unit = "mm"),
    legend.position = "bottom",
    legend.box = "horizontal",
    legend.box.just = "center",
    legend.direction = "horizontal",
    legend.title = element_text(size = base_pt),
    legend.text = element_text(size = base_pt),
    legend.key.height = grid::unit(2.8, "mm"),
    legend.key.width = grid::unit(2.6, "mm"),
    legend.spacing.x = grid::unit(0.2, "mm"),
    legend.spacing.y = grid::unit(0.1, "mm"),
    legend.margin = margin(0, 0, 0, 0)
  )

output_pdf <- file.path(
  script_dir,
  "BRCA1_GO_BP_bipartite_network_v5_compact_8x10cm.pdf"
)

ggsave(
  filename = output_pdf,
  plot = p_compact,
  width = 8,
  height = 10,
  units = "cm",
  device = grDevices::pdf,
  family = "Helvetica",
  useDingbats = FALSE,
  compress = FALSE,
  bg = "white"
)

writeLines(c(
  "BRCA1 GO:BP bipartite network v5 compact",
  "Compact version of v4 final mirrored layout",
  "Page: 8 cm x 10 cm; all visible text remains 5 pt",
  "Reduced module gaps, horizontal spacing, margins, node ranges and legend widths",
  "Compact display labels are abbreviated for fit; the pathway-node TSV preserves all full GO names",
  "Module codes: Met, Imm, Death, Ox, Prot and Prolif",
  "Pathway inclusion and all statistical values are unchanged",
  "PDF only; Helvetica; standard R PDF device; Dingbats and compression disabled"
), file.path(script_dir, "BRCA1_GO_BP_bipartite_network_v5_compact_summary.txt"))

cat("Created:", output_pdf, "\n")
```
