## Supplementary material for "An integrated, transcript-resolution atlas of prostate cancer progression unmasks associations with oncogenic pathways": Reproducible code: redraw_BRCA1_T_vs_GFP_BioID_volcano_9x6cm.R

```
#!/usr/bin/env Rscript
# Public submission copy.
# Analytical statements and parameter values are unchanged from the archived source.
# Legacy non-English comments and console messages were normalized for public release.

# Redraw the existing BRCA1-T versus GFP BioID volcano plot at 9 x 6 cm.
# Statistical values, thresholds, status classes, and labelled proteins are
# read from the archived analysis table and are not recalculated here.

suppressPackageStartupMessages({
  library(ggplot2)
  library(ggrepel)
})

script_arg <- grep("^--file=", commandArgs(), value = TRUE)
script_dir <- if (length(script_arg)) {
  dirname(normalizePath(sub("^--file=", "", script_arg)))
} else {
  normalizePath(getwd())
}

input_file <- file.path(script_dir, "BRCA1_T_vs_GFP_BioID_analysis_table.tsv")
output_file <- file.path(script_dir, "BRCA1_T_vs_GFP_BioID_volcano_9x6cm.pdf")

base_pt <- 5
text_mm <- base_pt / 2.845276
fc_cutoff <- 0.5
seed <- 20260826

dat <- read.delim(
  input_file,
  check.names = FALSE,
  quote = "",
  comment.char = "",
  stringsAsFactors = FALSE
)

dat$status <- factor(
  dat$status,
  levels = c("Not significant", "GFP enriched", "BRCA1-T enriched")
)
dat$volcano_label[dat$volcano_label == ""] <- NA_character_

plot_cols <- c(
  `Not significant` = "#B8BDC5",
  `GFP enriched` = "#2C73B9",
  `BRCA1-T enriched` = "#D9684F"
)

p <- ggplot(
  dat,
  aes(x = log2_fc_BRCA1_T_minus_GFP, y = neglog10_p, color = status)
) +
  geom_vline(
    xintercept = c(-fc_cutoff, fc_cutoff),
    color = "#888888",
    linetype = "dashed",
    linewidth = 0.22
  ) +
  geom_point(size = 0.62, alpha = 0.82, na.rm = TRUE) +
  geom_label_repel(
    data = subset(dat, !is.na(volcano_label)),
    aes(label = volcano_label),
    color = "#222222",
    fill = scales::alpha("white", 0.94),
    size = text_mm,
    label.size = 0.06,
    label.padding = grid::unit(0.08, "lines"),
    box.padding = 0.16,
    point.padding = 0.06,
    min.segment.length = 0,
    max.overlaps = Inf,
    seed = seed,
    force = 1.2,
    max.time = 5,
    segment.color = "#888888",
    segment.size = 0.12,
    show.legend = FALSE
  ) +
  scale_color_manual(values = plot_cols, drop = FALSE) +
  scale_x_continuous(expand = expansion(mult = c(0.04, 0.06))) +
  scale_y_continuous(expand = expansion(mult = c(0.02, 0.08))) +
  labs(
    title = "BRCA1-T vs GFP BioID",
    subtitle = "q < 0.05 and |log2 FC| >= 0.5",
    x = "BRCA1-T - GFP log2 LFQ difference",
    y = "-log10 Student t-test p value",
    color = NULL
  ) +
  theme_classic(base_size = base_pt, base_family = "Helvetica") +
  theme(
    text = element_text(size = base_pt, family = "Helvetica", color = "black"),
    plot.title = element_text(size = base_pt, face = "bold", hjust = 0.5,
                              margin = margin(b = 0.6, unit = "pt")),
    plot.subtitle = element_text(size = base_pt, hjust = 0.5,
                                 margin = margin(b = 1.5, unit = "pt")),
    axis.title = element_text(size = base_pt),
    axis.text = element_text(size = base_pt),
    axis.ticks = element_line(linewidth = 0.25),
    axis.line = element_line(linewidth = 0.3),
    legend.position = "bottom",
    legend.direction = "horizontal",
    legend.text = element_text(size = base_pt),
    legend.key.height = grid::unit(0.18, "cm"),
    legend.key.width = grid::unit(0.28, "cm"),
    legend.spacing.x = grid::unit(0.10, "cm"),
    legend.margin = margin(t = -1.5, unit = "pt"),
    plot.margin = margin(2.2, 2.5, 1.5, 2.5, unit = "pt")
  ) +
  guides(color = guide_legend(
    nrow = 1,
    byrow = TRUE,
    override.aes = list(size = 0.9, alpha = 1)
  ))

ggsave(
  filename = output_file,
  plot = p,
  width = 9,
  height = 6,
  units = "cm",
  device = grDevices::pdf,
  family = "Helvetica",
  useDingbats = FALSE,
  compress = FALSE,
  bg = "white",
  limitsize = FALSE
)

message("Wrote: ", output_file)
```
