## Supplementary material for "An integrated, transcript-resolution atlas of prostate cancer progression unmasks associations with oncogenic pathways": Reproducible code: run_BRCA1_GO_BP_joint_continuous.R

```
#!/usr/bin/env Rscript
# Public submission copy.
# Analytical statements and parameter values are unchanged from the archived source.
# Legacy non-English comments and console messages were normalized for public release.

# Integrated GO Biological Process GSEA for:
# 1) Atlas BRCA1-T conditional continuous association, adjusted only for BRCA1-C
# 2) Atlas BRCA1-C conditional continuous association, adjusted only for BRCA1-T
# 3) Experimental BRCA1-T overexpression versus pCW107 (Figure S10I)
# 4) BRCA1-T BioID versus GFP (cross-modality comparison)
#
# The atlas model intentionally contains no technical or biological covariates:
#   gene count ~ z(log2CPM(BRCA1-C) + 1) + z(log2CPM(BRCA1-T) + 1)

suppressPackageStartupMessages({
  library(fst)
  library(DESeq2)
  library(ggplot2)
  library(dplyr)
  library(tidyr)
  library(stringr)
  library(msigdbr)
  library(fgsea)
  library(cowplot)
})

script_arg <- grep("^--file=", commandArgs(), value = TRUE)
script_dir <- if (length(script_arg)) {
  dirname(normalizePath(sub("^--file=", "", script_arg)))
} else {
  normalizePath(getwd())
}
root_dir <- dirname(script_dir)
output_dir <- script_dir

source_dir <- file.path(
  root_dir, "reproducible_code_submission_v1.0", "source_data", "brca1"
)
input_files <- c(
  Experiment_BRCA1_T = file.path(source_dir, "DEs_11q_vs_CTRL.csv"),
  BioID_BRCA1_T = file.path(
    root_dir, "bioID_BRCA", "BRCA1_T_vs_GFP_BioID_analysis",
    "BRCA1_T_vs_GFP_GSEA_ranking_unique_genes.tsv"
  )
)
stopifnot(all(file.exists(input_files)))

atlas_dir <- "__PCATLAS_DOCUMENTS_ROOT__/Data_Harmanization"
atlas_files <- c(
  transcript_counts = file.path(atlas_dir, "txi.counts.fst"),
  transcript_info = file.path(atlas_dir, "transcripts_info.csv"),
  gene_counts = file.path(atlas_dir, "gc_txi.counts.fst"),
  gene_info = file.path(atlas_dir, "genes_info.csv")
)
stopifnot(all(file.exists(atlas_files)))

page_cm <- 10
base_pt <- 5
seed <- 20260827
fdr_cutoff <- 0.05
min_size <- 15L
max_size <- Inf
jaccard_cutoff <- 0.50
representative_n <- 5L

# Reuse the pinned local MSigDB 2026.1 cache already used for BioID GSEA.
msigdb_cache <- file.path(
  root_dir, "bioID-ERF", "ERF202_vs_ERF201_GO_GSEA", ".r-cache"
)
Sys.setenv(R_USER_CACHE_DIR = msigdb_cache)

comparison_labels <- c(
  Atlas_BRCA1_T = "Atlas BRCA1-T conditional association",
  Atlas_BRCA1_C = "Atlas BRCA1-C conditional association",
  Experiment_BRCA1_T = "BRCA1-T OE vs pCW107",
  BioID_BRCA1_T = "BRCA1-T BioID vs GFP"
)

positive_labels <- c(
  Atlas_BRCA1_T = "Positive BRCA1-T conditional association",
  Atlas_BRCA1_C = "Positive BRCA1-C conditional association",
  Experiment_BRCA1_T = "BRCA1-T OE enriched",
  BioID_BRCA1_T = "BRCA1-T BioID enriched"
)
negative_labels <- c(
  Atlas_BRCA1_T = "Negative BRCA1-T conditional association",
  Atlas_BRCA1_C = "Negative BRCA1-C conditional association",
  Experiment_BRCA1_T = "pCW107 enriched",
  BioID_BRCA1_T = "GFP enriched"
)

save_pdf <- function(filename, plot, width_cm = page_cm, height_cm = page_cm) {
  ggsave(
    filename = file.path(output_dir, filename), plot = plot,
    width = width_cm, height = height_cm, units = "cm",
    device = grDevices::pdf, family = "Helvetica",
    useDingbats = FALSE, compress = FALSE, bg = "white"
  )
}

theme_5pt <- function() {
  theme_classic(base_size = base_pt, base_family = "Helvetica") +
    theme(
      text = element_text(size = base_pt, family = "Helvetica", color = "black"),
      plot.title = element_text(size = base_pt, face = "bold", hjust = 0.5),
      plot.subtitle = element_text(size = base_pt, hjust = 0.5),
      axis.title = element_text(size = base_pt),
      axis.text = element_text(size = base_pt),
      strip.text = element_text(size = base_pt, face = "bold"),
      legend.title = element_text(size = base_pt),
      legend.text = element_text(size = base_pt)
    )
}

clean_rank <- function(gene, score, source_row = seq_along(gene)) {
  z <- data.frame(
    gene = toupper(trimws(as.character(gene))),
    score = suppressWarnings(as.numeric(score)),
    source_row = source_row,
    stringsAsFactors = FALSE
  )
  z <- z[!is.na(z$gene) & z$gene != "" & !is.na(z$score) & is.finite(z$score), ]
  z <- z[order(abs(z$score), decreasing = TRUE), ]
  z$duplicate_gene <- duplicated(z$gene)
  unique_z <- z[!z$duplicate_gene, ]
  ranks <- sort(setNames(unique_z$score, unique_z$gene), decreasing = TRUE)
  list(all = z, unique = unique_z, ranks = ranks)
}

# Build the two continuous atlas predictors exactly from the transcript-count
# matrix used by the preceding quartile analysis. Counts are converted to
# log2(CPM + 1), then standardized across all 1,365 atlas samples.
tx_counts <- as.matrix(read.fst(atlas_files[["transcript_counts"]]))
storage.mode(tx_counts) <- "double"
tx_info <- read.csv(atlas_files[["transcript_info"]], check.names = FALSE)
stopifnot(nrow(tx_counts) == nrow(tx_info))
rownames(tx_counts) <- tx_info$transcript_id

brca1_ids <- c(
  BRCA1_T = "ENST00000352993.7",
  BRCA1_C = "ENST00000357654.9"
)
stopifnot(all(brca1_ids %in% rownames(tx_counts)))
library_sizes <- colSums(tx_counts)
stopifnot(all(is.finite(library_sizes)), all(library_sizes > 0))
brca1_cpm <- sweep(
  tx_counts[unname(brca1_ids), , drop = FALSE],
  2, library_sizes / 1e6, "/"
)
rownames(brca1_cpm) <- names(brca1_ids)
brca1_logcpm <- log2(brca1_cpm + 1)

sample_data <- data.frame(
  BRCA1_C_z = as.numeric(scale(brca1_logcpm["BRCA1_C", ])),
  BRCA1_T_z = as.numeric(scale(brca1_logcpm["BRCA1_T", ])),
  row.names = colnames(tx_counts),
  check.names = FALSE
)

# Gene-level raw counts are modeled jointly with both continuous predictors.
# No dataset, state, purity, library, or other covariate is included by design.
gene_counts <- as.matrix(read.fst(atlas_files[["gene_counts"]]))
storage.mode(gene_counts) <- "double"
gene_info <- read.csv(atlas_files[["gene_info"]], check.names = FALSE)
stopifnot(
  nrow(gene_counts) == nrow(gene_info),
  ncol(gene_counts) == nrow(sample_data),
  identical(colnames(gene_counts), rownames(sample_data))
)
rownames(gene_counts) <- gene_info$gene_id
gene_counts <- round(gene_counts)
keep_gene <- rowSums(gene_counts) > 0
gene_counts <- gene_counts[keep_gene, , drop = FALSE]
gene_info_model <- gene_info[keep_gene, , drop = FALSE]

dds <- DESeqDataSetFromMatrix(
  countData = gene_counts,
  colData = sample_data,
  design = ~ BRCA1_C_z + BRCA1_T_z
)
dds <- DESeq(dds, quiet = TRUE)
stopifnot(all(c("BRCA1_C_z", "BRCA1_T_z") %in% resultsNames(dds)))

format_atlas_result <- function(coefficient) {
  result <- as.data.frame(results(dds, name = coefficient))
  result$gene_id <- rownames(result)
  result$SYMBOL <- gene_info_model$gene_name[
    match(result$gene_id, gene_info_model$gene_id)
  ]
  result
}

atlas_t <- format_atlas_result("BRCA1_T_z")
atlas_c <- format_atlas_result("BRCA1_C_z")
write.table(
  atlas_t,
  file.path(output_dir, "Atlas_BRCA1_T_joint_continuous_DESeq2_results.tsv"),
  sep = "\t", quote = FALSE, row.names = FALSE
)
write.table(
  atlas_c,
  file.path(output_dir, "Atlas_BRCA1_C_joint_continuous_DESeq2_results.tsv"),
  sep = "\t", quote = FALSE, row.names = FALSE
)

predictor_r <- cor(sample_data$BRCA1_T_z, sample_data$BRCA1_C_z)
predictor_vif <- 1 / (1 - predictor_r^2)
writeLines(c(
  "BRCA1 atlas joint continuous model",
  "Design: gene count ~ BRCA1_C_z + BRCA1_T_z",
  "No additional covariates were included.",
  paste0("Samples: ", nrow(sample_data)),
  paste0("Tested nonzero genes: ", nrow(gene_counts)),
  paste0("BRCA1-T transcript: ", brca1_ids[["BRCA1_T"]]),
  paste0("BRCA1-C transcript: ", brca1_ids[["BRCA1_C"]]),
  "Predictor transform: z score of log2(CPM + 1)",
  paste0("Predictor Pearson r: ", signif(predictor_r, 6)),
  paste0("Two-predictor VIF: ", signif(predictor_vif, 6)),
  "Interpretation: each coefficient is a conditional association adjusted only for the other BRCA1 isoform."
), file.path(output_dir, "Atlas_BRCA1_joint_continuous_model_summary.txt"))

experiment_t <- read.csv(input_files[["Experiment_BRCA1_T"]], check.names = FALSE)
bioid_t <- read.delim(input_files[["BioID_BRCA1_T"]], check.names = FALSE)

rank_objects <- list(
  Atlas_BRCA1_T = clean_rank(atlas_t$SYMBOL, atlas_t$stat),
  Atlas_BRCA1_C = clean_rank(atlas_c$SYMBOL, atlas_c$stat),
  Experiment_BRCA1_T = clean_rank(experiment_t$gene_name, experiment_t$stat),
  BioID_BRCA1_T = clean_rank(bioid_t$gene, bioid_t$t_statistic)
)

for (nm in names(rank_objects)) {
  write.table(
    rank_objects[[nm]]$unique,
    file.path(output_dir, paste0(nm, "_GO_BP_ranking_unique_genes.tsv")),
    sep = "\t", quote = FALSE, row.names = FALSE
  )
}

msigdb_c5 <- msigdbr(species = "Homo sapiens", collection = "C5")
go_bp <- msigdb_c5 %>%
  filter(gs_subcollection == "GO:BP") %>%
  select(gs_name, gene_symbol) %>%
  distinct()
pathways <- split(go_bp$gene_symbol, go_bp$gs_name)
msigdb_version <- unique(msigdb_c5$db_version)

run_gsea <- function(ranks, comparison, seed_offset) {
  set.seed(seed + seed_offset)
  fg <- fgseaMultilevel(
    pathways = pathways,
    stats = ranks,
    minSize = min_size,
    maxSize = max_size,
    eps = 0,
    nPermSimple = 10000
  )
  out <- as.data.frame(fg)
  out$leadingEdge <- vapply(out$leadingEdge, paste, collapse = ";", FUN.VALUE = character(1))
  out$comparison <- comparison
  out$comparison_label <- unname(comparison_labels[[comparison]])
  out$direction <- ifelse(
    out$NES >= 0,
    unname(positive_labels[[comparison]]),
    unname(negative_labels[[comparison]])
  )
  out <- out[order(out$padj, -abs(out$NES)), ]
  out
}

gsea_results <- Map(
  function(obj, nm, i) run_gsea(obj$ranks, nm, i),
  rank_objects, names(rank_objects), seq_along(rank_objects)
)
names(gsea_results) <- names(rank_objects)

for (nm in names(gsea_results)) {
  all_file <- file.path(output_dir, paste0(nm, "_GO_BP_GSEA_all.tsv"))
  sig_file <- file.path(output_dir, paste0(nm, "_GO_BP_GSEA_FDR_lt_0.05.tsv"))
  write.table(gsea_results[[nm]], all_file, sep = "\t", quote = FALSE, row.names = FALSE)
  write.table(
    subset(gsea_results[[nm]], !is.na(padj) & padj < fdr_cutoff),
    sig_file, sep = "\t", quote = FALSE, row.names = FALSE
  )
}

all_gsea <- bind_rows(gsea_results)
sig_gsea <- all_gsea %>% filter(!is.na(padj), padj < fdr_cutoff)
write.table(
  all_gsea, file.path(output_dir, "BRCA1_four_comparisons_GO_BP_GSEA_all.tsv"),
  sep = "\t", quote = FALSE, row.names = FALSE
)
write.table(
  sig_gsea, file.path(output_dir, "BRCA1_four_comparisons_GO_BP_GSEA_FDR_lt_0.05.tsv"),
  sep = "\t", quote = FALSE, row.names = FALSE
)

counts <- sig_gsea %>%
  mutate(sign = ifelse(NES >= 0, "Positive NES", "Negative NES")) %>%
  count(comparison, comparison_label, sign, name = "significant_terms") %>%
  complete(
    comparison = names(comparison_labels), sign = c("Positive NES", "Negative NES"),
    fill = list(significant_terms = 0)
  ) %>%
  mutate(comparison_label = unname(comparison_labels[comparison]))
write.table(
  counts, file.path(output_dir, "BRCA1_GO_BP_significant_term_counts.tsv"),
  sep = "\t", quote = FALSE, row.names = FALSE
)

# Exact GO-term membership and direction matrices.
all_sig_paths <- sort(unique(sig_gsea$pathway))
membership <- expand.grid(
  pathway = all_sig_paths,
  comparison = names(comparison_labels),
  stringsAsFactors = FALSE
) %>%
  left_join(
    sig_gsea %>% select(pathway, comparison, NES, padj, direction),
    by = c("pathway", "comparison")
  ) %>%
  mutate(significant = as.integer(!is.na(padj)))

membership_wide <- membership %>%
  select(pathway, comparison, significant) %>%
  pivot_wider(names_from = comparison, values_from = significant, values_fill = 0)
direction_wide <- membership %>%
  mutate(direction_code = ifelse(is.na(NES), 0, sign(NES))) %>%
  select(pathway, comparison, direction_code) %>%
  pivot_wider(names_from = comparison, values_from = direction_code, values_fill = 0)
nes_wide <- membership %>%
  select(pathway, comparison, NES) %>%
  pivot_wider(names_from = comparison, values_from = NES)
padj_wide <- membership %>%
  select(pathway, comparison, padj) %>%
  pivot_wider(names_from = comparison, values_from = padj)

write.table(
  membership_wide, file.path(output_dir, "BRCA1_GO_BP_significant_membership_matrix.tsv"),
  sep = "\t", quote = FALSE, row.names = FALSE
)
write.table(
  direction_wide, file.path(output_dir, "BRCA1_GO_BP_significant_direction_matrix.tsv"),
  sep = "\t", quote = FALSE, row.names = FALSE
)
write.table(
  nes_wide, file.path(output_dir, "BRCA1_GO_BP_significant_NES_matrix.tsv"),
  sep = "\t", quote = FALSE, row.names = FALSE
)
write.table(
  padj_wide, file.path(output_dir, "BRCA1_GO_BP_significant_FDR_matrix.tsv"),
  sep = "\t", quote = FALSE, row.names = FALSE
)

set_list <- lapply(names(comparison_labels), function(nm) {
  sig_gsea$pathway[sig_gsea$comparison == nm]
})
names(set_list) <- names(comparison_labels)

pairwise_rows <- list()
pair_i <- 1L
for (i in seq_along(set_list)) {
  for (j in seq_along(set_list)) {
    a <- names(set_list)[i]
    b <- names(set_list)[j]
    shared <- intersect(set_list[[a]], set_list[[b]])
    union_terms <- union(set_list[[a]], set_list[[b]])
    if (length(shared)) {
      da <- direction_wide[match(shared, direction_wide$pathway), a]
      db <- direction_wide[match(shared, direction_wide$pathway), b]
      same_n <- sum(da[[1]] == db[[1]])
    } else {
      same_n <- 0L
    }
    pairwise_rows[[pair_i]] <- data.frame(
      comparison_1 = a,
      comparison_2 = b,
      n_1 = length(set_list[[a]]),
      n_2 = length(set_list[[b]]),
      shared = length(shared),
      same_direction = same_n,
      opposite_direction = length(shared) - same_n,
      jaccard = if (length(union_terms)) length(shared) / length(union_terms) else NA_real_,
      stringsAsFactors = FALSE
    )
    pair_i <- pair_i + 1L
  }
}
pairwise <- bind_rows(pairwise_rows)
write.table(
  pairwise, file.path(output_dir, "BRCA1_GO_BP_pairwise_overlap.tsv"),
  sep = "\t", quote = FALSE, row.names = FALSE
)

make_intersections <- function(groups) {
  m <- membership_wide[, c("pathway", groups), drop = FALSE]
  binary <- as.matrix(m[, groups, drop = FALSE])
  patterns <- apply(binary, 1, function(z) paste(groups[z == 1], collapse = ";"))
  tab <- as.data.frame(table(patterns), stringsAsFactors = FALSE)
  names(tab) <- c("pattern", "n_pathways")
  tab <- tab[tab$pattern != "" & tab$n_pathways > 0, ]
  tab <- tab[order(tab$n_pathways, decreasing = TRUE), ]
  tab$intersection_id <- paste0("I", seq_len(nrow(tab)))
  members <- do.call(rbind, lapply(seq_len(nrow(tab)), function(i) {
    data.frame(
      intersection_id = tab$intersection_id[i],
      comparison = groups,
      member = as.integer(groups %in% strsplit(tab$pattern[i], ";", fixed = TRUE)[[1]]),
      stringsAsFactors = FALSE
    )
  }))
  list(summary = tab, members = members)
}

plot_upset <- function(groups, prefix, title) {
  ints <- make_intersections(groups)
  write.table(
    ints$summary,
    file.path(output_dir, paste0(prefix, "_intersection_counts.tsv")),
    sep = "\t", quote = FALSE, row.names = FALSE
  )
  ord <- ints$summary$intersection_id
  ints$summary$intersection_id <- factor(ints$summary$intersection_id, levels = ord)
  ints$members$intersection_id <- factor(ints$members$intersection_id, levels = ord)
  ints$members$comparison <- factor(ints$members$comparison, levels = rev(groups))
  ints$members$comparison_num <- as.numeric(ints$members$comparison)

  p_bar <- ggplot(ints$summary, aes(intersection_id, n_pathways)) +
    geom_col(width = 0.7, fill = "#4B6F8A") +
    geom_text(aes(label = n_pathways), vjust = -0.25, size = base_pt / 2.845276) +
    scale_y_continuous(expand = expansion(mult = c(0, 0.15))) +
    labs(title = title, x = NULL, y = "Significant GO:BP terms") +
    theme_5pt() +
    theme(
      axis.text.x = element_blank(), axis.ticks.x = element_blank(),
      plot.margin = margin(2, 2, 0, 8, unit = "pt")
    )

  active <- ints$members %>% filter(member == 1)
  spans <- active %>%
    group_by(intersection_id) %>%
    summarise(
      ymin = min(comparison_num),
      ymax = max(comparison_num),
      .groups = "drop"
    )
  p_mat <- ggplot(ints$members, aes(intersection_id, comparison_num)) +
    geom_segment(
      data = spans,
      aes(x = intersection_id, xend = intersection_id, y = ymin, yend = ymax),
      inherit.aes = FALSE, color = "#333333", linewidth = 0.3
    ) +
    geom_point(aes(fill = factor(member)), shape = 21, size = 1.4, stroke = 0.25) +
    scale_fill_manual(values = c(`0` = "white", `1` = "#222222"), guide = "none") +
    scale_y_continuous(
      breaks = seq_along(groups),
      labels = rev(unname(comparison_labels[groups])),
      expand = expansion(add = 0.45)
    ) +
    labs(x = "Exact significant-term intersection", y = NULL) +
    theme_5pt() +
    theme(
      axis.line.y = element_blank(), axis.ticks.y = element_blank(),
      plot.margin = margin(0, 2, 2, 8, unit = "pt")
    )

  combined <- plot_grid(
    p_bar, p_mat, ncol = 1, align = "v", axis = "lr",
    rel_heights = c(1.7, 1)
  )
  save_pdf(paste0(prefix, ".pdf"), combined, 10, 10)
}

transcript_groups <- c("Atlas_BRCA1_T", "Atlas_BRCA1_C", "Experiment_BRCA1_T")
all_groups <- names(comparison_labels)
plot_upset(
  transcript_groups,
  "BRCA1_GO_BP_overlap_3_transcriptomic_groups",
  "GO:BP overlap across atlas and perturbation RNA-seq"
)
plot_upset(
  all_groups,
  "BRCA1_GO_BP_overlap_4_groups_including_BioID",
  "GO:BP overlap including BRCA1-T BioID"
)

# Pairwise Jaccard heatmap.
pairwise_plot <- pairwise %>%
  mutate(
    comparison_1 = factor(comparison_1, levels = all_groups),
    comparison_2 = factor(comparison_2, levels = rev(all_groups)),
    label = sprintf("%.2f\n(n=%d)", jaccard, shared)
  )
p_jaccard <- ggplot(pairwise_plot, aes(comparison_1, comparison_2, fill = jaccard)) +
  geom_tile(color = "white", linewidth = 0.35) +
  geom_text(aes(label = label), size = base_pt / 2.845276) +
  scale_fill_gradient(low = "white", high = "#375A7F", limits = c(0, 1)) +
  scale_x_discrete(labels = comparison_labels) +
  scale_y_discrete(labels = comparison_labels) +
  coord_fixed() +
  labs(
    title = "Pairwise overlap of significant GO:BP terms",
    subtitle = "Cell value: Jaccard index and exact shared-term count",
    x = NULL, y = NULL, fill = "Jaccard"
  ) +
  theme_5pt() +
  theme(
    axis.text.x = element_text(angle = 45, hjust = 1),
    axis.ticks = element_blank()
  )
save_pdf("BRCA1_GO_BP_pairwise_overlap_Jaccard.pdf", p_jaccard, 10, 10)

# Direction-aware NES heatmap for the strongest exact shared terms.
shared_terms <- membership_wide %>%
  mutate(n_groups = rowSums(across(all_of(all_groups)))) %>%
  filter(n_groups >= 2) %>%
  select(pathway, n_groups)

if (nrow(shared_terms)) {
  shared_long <- membership %>%
    filter(pathway %in% shared_terms$pathway) %>%
    left_join(shared_terms, by = "pathway") %>%
    group_by(pathway) %>%
    mutate(best_fdr = min(padj, na.rm = TRUE)) %>%
    ungroup()
  selected_shared <- shared_long %>%
    distinct(pathway, n_groups, best_fdr) %>%
    arrange(desc(n_groups), best_fdr) %>%
    slice_head(n = 30) %>%
    pull(pathway)
  shared_plot <- shared_long %>%
    filter(pathway %in% selected_shared) %>%
    mutate(
      pathway_label = str_to_sentence(str_replace_all(str_remove(pathway, "^GOBP_"), "_", " ")),
      pathway_label = factor(pathway_label, levels = rev(unique(pathway_label[order(match(pathway, selected_shared))]))),
      comparison = factor(comparison, levels = all_groups),
      label = ifelse(is.na(NES), "", sprintf("%.1f", NES))
    )
  heat_height <- max(10, min(18, 4 + 0.28 * length(selected_shared)))
  p_shared <- ggplot(shared_plot, aes(comparison, pathway_label, fill = NES)) +
    geom_tile(color = "white", linewidth = 0.25) +
    geom_text(aes(label = label), size = base_pt / 2.845276) +
    scale_fill_gradient2(
      low = "#2C73B9", mid = "white", high = "#D9684F", midpoint = 0,
      na.value = "#F2F2F2"
    ) +
    scale_x_discrete(labels = comparison_labels) +
    labs(
      title = "Strongest GO:BP terms significant in at least two comparisons",
      subtitle = "Numbers are NES; blank cells are not significant at FDR < 0.05",
      x = NULL, y = NULL, fill = "NES"
    ) +
    theme_5pt() +
    theme(axis.text.x = element_text(angle = 45, hjust = 1), axis.ticks = element_blank())
  save_pdf("BRCA1_GO_BP_shared_terms_NES_heatmap.pdf", p_shared, 10, heat_height)
}

# Representative nonredundant significant terms for a compact GSEA overview.
jaccard <- function(a, b) {
  u <- union(a, b)
  if (!length(u)) return(0)
  length(intersect(a, b)) / length(u)
}

select_representatives <- function(x, n = representative_n) {
  if (!nrow(x)) return(x)
  x <- x[order(x$padj, -abs(x$NES)), ]
  keep <- integer(0)
  leading <- strsplit(x$leadingEdge, ";", fixed = TRUE)
  for (i in seq_len(nrow(x))) {
    if (!length(keep) || all(vapply(keep, function(k) {
      jaccard(leading[[i]], leading[[k]]) < jaccard_cutoff
    }, logical(1)))) {
      keep <- c(keep, i)
    }
    if (length(keep) >= n) break
  }
  x[keep, , drop = FALSE]
}

representatives <- sig_gsea %>%
  mutate(nes_sign = ifelse(NES >= 0, "Positive NES", "Negative NES")) %>%
  group_split(comparison, nes_sign) %>%
  lapply(select_representatives) %>%
  bind_rows()

write.table(
  representatives,
  file.path(output_dir, "BRCA1_GO_BP_representative_terms_for_bubble.tsv"),
  sep = "\t", quote = FALSE, row.names = FALSE
)

if (nrow(representatives)) {
  rep_plot <- representatives %>%
    mutate(
      pathway_label = str_to_sentence(str_replace_all(str_remove(pathway, "^GOBP_"), "_", " ")),
      pathway_label = factor(pathway_label, levels = rev(unique(pathway_label[order(NES)]))),
      comparison_label = factor(comparison_label, levels = comparison_labels),
      sig_size = pmin(-log10(pmax(padj, 1e-300)), 12)
    )
  p_bubble <- ggplot(rep_plot, aes(NES, pathway_label)) +
    geom_vline(xintercept = 0, color = "#777777", linewidth = 0.25) +
    geom_point(aes(size = sig_size, fill = NES), shape = 21, color = "#333333", stroke = 0.25) +
    scale_fill_gradient2(low = "#2C73B9", mid = "white", high = "#D9684F", midpoint = 0) +
    scale_size(range = c(1.2, 3.2)) +
    facet_grid(comparison_label ~ ., scales = "free_y", space = "free_y") +
    labs(
      title = "Representative GO:BP GSEA",
      subtitle = "Positive NES follows the positive association or named perturbation direction",
      x = "Normalized enrichment score (NES)", y = NULL,
      fill = "NES", size = "-log10 FDR"
    ) +
    theme_5pt() +
    theme(panel.grid.major.x = element_line(color = "#E5E5E5", linewidth = 0.2))
  bubble_height <- max(12, min(20, 5 + 0.24 * nrow(representatives)))
  save_pdf("BRCA1_four_comparisons_GO_BP_representative_bubble.pdf", p_bubble, 10, bubble_height)
}

transcript_intersection <- Reduce(intersect, set_list[transcript_groups])
four_way_intersection <- Reduce(intersect, set_list[all_groups])
four_way_direction_rows <- direction_wide[
  match(four_way_intersection, direction_wide$pathway),
  all_groups,
  drop = FALSE
]
four_way_all_same_direction <- if (nrow(four_way_direction_rows)) {
  sum(apply(four_way_direction_rows, 1, function(v) length(unique(v)) == 1))
} else {
  0L
}
writeLines(
  c("pathway", transcript_intersection),
  file.path(output_dir, "BRCA1_GO_BP_3_transcriptomic_groups_exact_intersection.tsv")
)
writeLines(
  c("pathway", four_way_intersection),
  file.path(output_dir, "BRCA1_GO_BP_4_groups_exact_intersection.tsv")
)

summary_lines <- c(
  "Integrated BRCA1 GO Biological Process GSEA and overlap analysis",
  paste0("Run date: ", Sys.Date()),
  paste0("MSigDB version: ", paste(msigdb_version, collapse = ", ")),
  paste0("Gene-set size filter: minimum ", min_size, "; no upper limit"),
  paste0("GSEA method: fgseaMultilevel; eps = 0; nPermSimple = 10000"),
  paste0("Significance threshold: FDR < ", fdr_cutoff),
  "",
  paste0(
    comparison_labels[names(rank_objects)], ": ",
    vapply(rank_objects, function(x) length(x$ranks), integer(1)),
    " unique ranked genes; ",
    vapply(set_list, length, integer(1)), " significant GO:BP terms"
  ),
  paste0(
    "Tied-statistic fraction (", comparison_labels[names(rank_objects)], "): ",
    sprintf(
      "%.2f%%",
      100 * vapply(rank_objects, function(x) {
        1 - length(unique(x$ranks)) / length(x$ranks)
      }, numeric(1))
    )
  ),
  "",
  paste0("Exact three-transcriptomic-group intersection: ", length(transcript_intersection), " terms"),
  paste0("Exact four-group intersection including BioID: ", length(four_way_intersection), " terms"),
  paste0(
    "Four-way shared terms with the same direction in all comparisons: ",
    four_way_all_same_direction, "; mixed direction: ",
    length(four_way_intersection) - four_way_all_same_direction
  ),
  paste0(
    "Atlas BRCA1-T versus experiment: ",
    pairwise$shared[pairwise$comparison_1 == "Atlas_BRCA1_T" & pairwise$comparison_2 == "Experiment_BRCA1_T"],
    " shared terms; ",
    pairwise$same_direction[pairwise$comparison_1 == "Atlas_BRCA1_T" & pairwise$comparison_2 == "Experiment_BRCA1_T"],
    " same-direction and ",
    pairwise$opposite_direction[pairwise$comparison_1 == "Atlas_BRCA1_T" & pairwise$comparison_2 == "Experiment_BRCA1_T"],
    " opposite-direction"
  ),
  paste0(
    "Atlas BRCA1-C versus experiment: ",
    pairwise$shared[pairwise$comparison_1 == "Atlas_BRCA1_C" & pairwise$comparison_2 == "Experiment_BRCA1_T"],
    " shared terms; ",
    pairwise$same_direction[pairwise$comparison_1 == "Atlas_BRCA1_C" & pairwise$comparison_2 == "Experiment_BRCA1_T"],
    " same-direction and ",
    pairwise$opposite_direction[pairwise$comparison_1 == "Atlas_BRCA1_C" & pairwise$comparison_2 == "Experiment_BRCA1_T"],
    " opposite-direction"
  ),
  paste0(
    "Atlas BRCA1-T versus BioID: ",
    pairwise$shared[pairwise$comparison_1 == "Atlas_BRCA1_T" & pairwise$comparison_2 == "BioID_BRCA1_T"],
    " shared terms; ",
    pairwise$same_direction[pairwise$comparison_1 == "Atlas_BRCA1_T" & pairwise$comparison_2 == "BioID_BRCA1_T"],
    " same-direction and ",
    pairwise$opposite_direction[pairwise$comparison_1 == "Atlas_BRCA1_T" & pairwise$comparison_2 == "BioID_BRCA1_T"],
    " opposite-direction"
  ),
  paste0(
    "Experiment versus BioID: ",
    pairwise$shared[pairwise$comparison_1 == "Experiment_BRCA1_T" & pairwise$comparison_2 == "BioID_BRCA1_T"],
    " shared terms; ",
    pairwise$same_direction[pairwise$comparison_1 == "Experiment_BRCA1_T" & pairwise$comparison_2 == "BioID_BRCA1_T"],
    " same-direction and ",
    pairwise$opposite_direction[pairwise$comparison_1 == "Experiment_BRCA1_T" & pairwise$comparison_2 == "BioID_BRCA1_T"],
    " opposite-direction"
  ),
  paste0(
    "Four-way exact shared term(s): ",
    if (length(four_way_intersection)) paste(four_way_intersection, collapse = "; ") else "none"
  ),
  "",
  "Interpretation note: the two atlas ranks are conditional continuous associations from one joint model with no additional covariates. They are not causal effects and remain potentially associated with disease state, dataset, or sample composition. The BioID rank reflects protein proximity, whereas the other three ranks reflect gene-expression changes; exact cross-modality overlap is therefore conservative."
)
writeLines(summary_lines, file.path(output_dir, "BRCA1_GO_BP_overlap_analysis_summary.txt"))
capture.output(sessionInfo(), file = file.path(output_dir, "R_sessionInfo.txt"))

cat("Completed integrated BRCA1 GO:BP comparison in:", output_dir, "\n")
```
