## Supplementary material for "An integrated, transcript-resolution atlas of prostate cancer progression unmasks associations with oncogenic pathways": Reproducible code: run_BRCA1_T_vs_GFP_BioID_analysis.R

```
#!/usr/bin/env Rscript
# Public submission copy.
# Analytical statements and parameter values are unchanged from the archived source.
# Legacy non-English comments and console messages were normalized for public release.

# Single-bait BioID analysis: truncated BRCA1 construct (BRCA1-T) versus GFP.
# Positive statistics indicate enrichment toward BRCA1-T.

suppressPackageStartupMessages({
  library(ggplot2)
  library(ggrepel)
  library(pheatmap)
  library(msigdbr)
  library(fgsea)
  library(stringr)
})

script_arg <- grep("^--file=", commandArgs(), value = TRUE)
script_dir <- if (length(script_arg)) {
  dirname(normalizePath(sub("^--file=", "", script_arg)))
} else {
  normalizePath(getwd())
}
output_dir <- script_dir
input_file <- normalizePath(file.path(
  script_dir, "..", "t_test_BRCA-BioID_vs_GFP-BioID_BH5FDR.txt"
))

page_cm <- 10
page_in <- page_cm / 2.54
base_pt <- 5
geom_text_mm <- base_pt / 2.845276
seed <- 20260826
fdr_cutoff <- 0.05
fc_cutoff <- 0.5
min_size <- 15L
max_size <- 500L
max_terms_per_direction <- 6L
jaccard_cutoff <- 0.50

# Reuse the previously downloaded pinned MSigDB cache.
msigdb_cache <- file.path(
  dirname(dirname(script_dir)), "bioID-ERF",
  "ERF202_vs_ERF201_GO_GSEA", ".r-cache"
)
Sys.setenv(R_USER_CACHE_DIR = msigdb_cache)

save_pdf <- function(filename, plot, width_cm = page_cm, height_cm = page_cm) {
  ggsave(
    filename = filename,
    plot = plot,
    width = width_cm,
    height = height_cm,
    units = "cm",
    device = grDevices::pdf,
    family = "Helvetica",
    useDingbats = FALSE,
    compress = FALSE,
    bg = "white"
  )
}

theme_5pt <- function() {
  theme_classic(base_size = base_pt, base_family = "Helvetica") +
    theme(
      text = element_text(size = base_pt, family = "Helvetica", color = "black"),
      plot.title = element_text(size = base_pt, face = "bold", hjust = 0.5),
      plot.subtitle = element_text(size = base_pt, hjust = 0.5),
      axis.title = element_text(size = base_pt),
      axis.text = element_text(size = base_pt),
      legend.title = element_text(size = base_pt),
      legend.text = element_text(size = base_pt),
      legend.key.height = grid::unit(0.25, "cm"),
      legend.key.width = grid::unit(0.35, "cm")
    )
}

read_perseus <- function(path) {
  x <- read.delim(
    path, check.names = FALSE, quote = "", comment.char = "",
    na.strings = c("", "NaN"), stringsAsFactors = FALSE
  )
  x[-c(1, 2), , drop = FALSE]
}

first_symbol <- function(x, fallback) {
  z <- toupper(trimws(sub(";.*$", "", x)))
  bad <- is.na(z) | z == "" | z == "NA"
  z[bad] <- fallback[bad]
  z
}

raw <- read_perseus(input_file)
lfq_cols <- grep("^LFQ intensity ", names(raw), value = TRUE)
sig_col <- grep("^Student's T-test Significant ", names(raw), value = TRUE)
p_col <- grep("^-Log10 Student's T-test p-value ", names(raw), value = TRUE)
q_col <- grep("^Student's T-test q-value ", names(raw), value = TRUE)
fc_col <- grep("^Student's T-test Log2 FC ", names(raw), value = TRUE)
t_col <- grep("^Student's T-test Test statistic ", names(raw), value = TRUE)
stopifnot(
  length(lfq_cols) == 8,
  length(sig_col) == 1,
  length(p_col) == 1,
  length(q_col) == 1,
  length(fc_col) == 1,
  length(t_col) == 1
)

dat <- data.frame(
  protein_id = raw[["Protein IDs"]],
  gene_names_original = raw[["Gene names"]],
  gene = first_symbol(raw[["Gene names"]], raw[["Protein names"]]),
  protein = raw[["Protein names"]],
  significant_export = !is.na(raw[[sig_col]]) & raw[[sig_col]] == "+",
  neglog10_p = as.numeric(raw[[p_col]]),
  q_value = as.numeric(raw[[q_col]]),
  log2_fc_BRCA1_T_minus_GFP = as.numeric(raw[[fc_col]]),
  t_statistic = as.numeric(raw[[t_col]]),
  stringsAsFactors = FALSE
)

sample_mat_all <- as.matrix(data.frame(
  lapply(raw[, lfq_cols, drop = FALSE], as.numeric), check.names = FALSE
))
colnames(sample_mat_all) <- c(
  paste0("BRCA1_T_", 1:4), paste0("GFP_", 1:4)
)
rownames(sample_mat_all) <- make.unique(dat$gene)

dat$status <- "Not significant"
dat$status[
  !is.na(dat$q_value) & dat$q_value < fdr_cutoff &
    dat$log2_fc_BRCA1_T_minus_GFP >= fc_cutoff
] <- "BRCA1-T enriched"
dat$status[
  !is.na(dat$q_value) & dat$q_value < fdr_cutoff &
    dat$log2_fc_BRCA1_T_minus_GFP <= -fc_cutoff
] <- "GFP enriched"
dat$status <- factor(
  dat$status,
  levels = c("Not significant", "GFP enriched", "BRCA1-T enriched")
)

plot_cols <- c(
  `Not significant` = "#B8BDC5",
  `GFP enriched` = "#2C73B9",
  `BRCA1-T enriched` = "#D9684F"
)

choose_labels <- function(d, n_each = 8L) {
  labels <- rep(NA_character_, nrow(d))
  for (direction in c("GFP enriched", "BRCA1-T enriched")) {
    ix <- which(d$status == direction)
    if (!length(ix)) next
    score <- d$neglog10_p[ix] + 0.15 * abs(d$log2_fc_BRCA1_T_minus_GFP[ix])
    keep <- head(ix[order(score, decreasing = TRUE)], n_each)
    labels[keep] <- d$gene[keep]
  }
  brca_ix <- which(d$gene == "BRCA1")
  if (length(brca_ix)) {
    best <- brca_ix[which.min(d$q_value[brca_ix])]
    labels[best] <- "BRCA1"
  }
  labels
}

dat$volcano_label <- choose_labels(dat, 8L)

volcano <- ggplot(
  dat,
  aes(log2_fc_BRCA1_T_minus_GFP, neglog10_p, color = status)
) +
  geom_vline(
    xintercept = c(-fc_cutoff, fc_cutoff),
    color = "#888888", linetype = "dashed", linewidth = 0.25
  ) +
  geom_point(size = 0.75, alpha = 0.82, na.rm = TRUE) +
  geom_label_repel(
    data = subset(dat, !is.na(volcano_label)),
    aes(label = volcano_label),
    color = "#222222", fill = scales::alpha("white", 0.92),
    size = geom_text_mm, label.size = 0.08,
    box.padding = 0.25, point.padding = 0.1,
    min.segment.length = 0, max.overlaps = Inf, seed = seed,
    segment.color = "#888888", segment.size = 0.15,
    show.legend = FALSE
  ) +
  scale_color_manual(values = plot_cols, drop = FALSE) +
  labs(
    title = "BRCA1-T vs GFP BioID",
    subtitle = "Color: q < 0.05 and absolute log2 FC >= 0.5",
    x = "BRCA1-T - GFP log2 LFQ difference",
    y = "-log10 Student t-test p value",
    color = NULL
  ) +
  theme_5pt() +
  theme(legend.position = "bottom")

save_pdf(file.path(output_dir, "BRCA1_T_vs_GFP_BioID_volcano.pdf"), volcano)

# Mean-intensity scatter for the single bait versus control design.
dat$mean_BRCA1_T <- rowMeans(sample_mat_all[, 1:4, drop = FALSE], na.rm = TRUE)
dat$mean_GFP <- rowMeans(sample_mat_all[, 5:8, drop = FALSE], na.rm = TRUE)
dat$scatter_label <- choose_labels(dat, 6L)
pearson_r <- cor(dat$mean_GFP, dat$mean_BRCA1_T, use = "complete.obs")

scatter <- ggplot(dat, aes(mean_GFP, mean_BRCA1_T)) +
  geom_abline(slope = 1, intercept = 0, color = "#BBBBBB", linewidth = 0.25) +
  geom_point(
    data = subset(dat, status == "Not significant"),
    color = plot_cols[["Not significant"]], size = 0.6, alpha = 0.55
  ) +
  geom_point(
    data = subset(dat, status != "Not significant"),
    aes(color = status), size = 0.9, alpha = 0.92
  ) +
  geom_label_repel(
    data = subset(dat, !is.na(scatter_label)),
    aes(label = scatter_label, color = status),
    fill = scales::alpha("white", 0.92),
    size = geom_text_mm, label.size = 0.08,
    box.padding = 0.25, point.padding = 0.1,
    min.segment.length = 0, max.overlaps = Inf, seed = seed,
    segment.color = "#888888", segment.size = 0.15,
    show.legend = FALSE
  ) +
  annotate(
    "text", x = -Inf, y = Inf, hjust = -0.08, vjust = 1.25,
    label = sprintf("Pearson R = %.2f, n = %d", pearson_r, nrow(dat)),
    size = geom_text_mm
  ) +
  scale_color_manual(values = plot_cols, drop = FALSE) +
  labs(
    title = "BRCA1-T and GFP BioID mean profiles",
    x = "Mean GFP log2 LFQ intensity",
    y = "Mean BRCA1-T log2 LFQ intensity",
    color = NULL
  ) +
  theme_5pt() +
  theme(legend.position = "bottom")

save_pdf(file.path(output_dir, "BRCA1_T_vs_GFP_BioID_mean_scatter.pdf"), scatter)

# Heatmap rows: up to 10 significant proteins per direction. There are fewer
# GFP-enriched proteins at the predeclared q and effect-size thresholds, so no
# non-significant rows are inserted to fill that group.
brca_ix <- which(dat$status == "BRCA1-T enriched")
gfp_ix <- which(dat$status == "GFP enriched")
brca_ix <- head(brca_ix[order(
  dat$q_value[brca_ix], -dat$log2_fc_BRCA1_T_minus_GFP[brca_ix]
)], 10)
gfp_ix <- head(gfp_ix[order(
  dat$q_value[gfp_ix], dat$log2_fc_BRCA1_T_minus_GFP[gfp_ix]
)], 10)
heat_ix <- c(brca_ix, gfp_ix)

heat_dat <- dat[heat_ix, , drop = FALSE]
heat_mat_raw <- sample_mat_all[heat_ix, , drop = FALSE]
rownames(heat_mat_raw) <- make.unique(heat_dat$gene)
heat_mat_z <- t(scale(t(heat_mat_raw)))

heat_mean_raw <- cbind(
  GFP = rowMeans(heat_mat_raw[, 5:8, drop = FALSE], na.rm = TRUE),
  `BRCA1-T` = rowMeans(heat_mat_raw[, 1:4, drop = FALSE], na.rm = TRUE)
)
heat_mean_z <- cbind(
  GFP = rowMeans(heat_mat_z[, 5:8, drop = FALSE], na.rm = TRUE),
  `BRCA1-T` = rowMeans(heat_mat_z[, 1:4, drop = FALSE], na.rm = TRUE)
)

ann_row <- data.frame(Direction = heat_dat$status, row.names = rownames(heat_mat_raw))
ann_col_8 <- data.frame(
  Bait = factor(
    c(rep("BRCA1-T", 4), rep("GFP", 4)),
    levels = c("GFP", "BRCA1-T")
  ),
  row.names = colnames(heat_mat_raw)
)
ann_col_2 <- data.frame(
  Bait = factor(c("GFP", "BRCA1-T"), levels = c("GFP", "BRCA1-T")),
  row.names = colnames(heat_mean_raw)
)
heat_ann_colors <- list(
  Bait = c(GFP = "#777777", `BRCA1-T` = "#D9684F"),
  Direction = c(
    `GFP enriched` = "#2C73B9",
    `BRCA1-T enriched` = "#D9684F"
  )
)
heat_cols <- colorRampPalette(c("#2C73B9", "#F7F7F7", "#D9684F"))(101)

centered_breaks <- function(x) {
  midpoint <- median(x, na.rm = TRUE)
  lim <- max(abs(x - midpoint), na.rm = TRUE)
  seq(midpoint - lim, midpoint + lim, length.out = 102)
}

sym_breaks <- function(x) {
  lim <- max(abs(x), na.rm = TRUE)
  seq(-lim, lim, length.out = 102)
}

raw_breaks <- centered_breaks(heat_mat_raw)
z_breaks <- sym_breaks(heat_mat_z)

draw_heatmap <- function(filename, matrix, title, breaks, annotation_col) {
  grDevices::pdf(
    filename, width = page_in, height = page_in,
    family = "Helvetica", useDingbats = FALSE, compress = FALSE,
    bg = "white"
  )
  pheatmap(
    matrix,
    color = heat_cols,
    breaks = breaks,
    cluster_rows = FALSE,
    cluster_cols = FALSE,
    border_color = "white",
    annotation_row = ann_row,
    annotation_col = annotation_col,
    annotation_colors = heat_ann_colors,
    fontsize = base_pt,
    fontsize_row = base_pt,
    fontsize_col = base_pt,
    cellwidth = 10,
    cellheight = 10,
    angle_col = 45,
    main = title,
    legend = TRUE,
    annotation_legend = TRUE
  )
  dev.off()
}

draw_heatmap(
  file.path(output_dir, "BRCA1_T_vs_GFP_heatmap_2col_unscaled.pdf"),
  heat_mean_raw,
  "Condition means (unscaled log2 LFQ)",
  raw_breaks,
  ann_col_2
)
draw_heatmap(
  file.path(output_dir, "BRCA1_T_vs_GFP_heatmap_2col_row_normalized.pdf"),
  heat_mean_z,
  "Condition means (mean row z score)",
  z_breaks,
  ann_col_2
)
draw_heatmap(
  file.path(output_dir, "BRCA1_T_vs_GFP_heatmap_8col_unscaled.pdf"),
  heat_mat_raw,
  "Eight replicates (unscaled log2 LFQ)",
  raw_breaks,
  ann_col_8
)
draw_heatmap(
  file.path(output_dir, "BRCA1_T_vs_GFP_heatmap_8col_row_normalized.pdf"),
  heat_mat_z,
  "Eight replicates (row z score)",
  z_breaks,
  ann_col_8
)

# Preranked GSEA using the full t-statistic-ranked list.
rank_table <- dat[
  !is.na(dat$gene) & dat$gene != "" & is.finite(dat$t_statistic),
  c(
    "protein_id", "gene_names_original", "gene", "protein",
    "t_statistic", "log2_fc_BRCA1_T_minus_GFP", "q_value"
  ),
  drop = FALSE
]
rank_table <- rank_table[
  order(rank_table$gene, -abs(rank_table$t_statistic)),
  , drop = FALSE
]
rank_table$duplicate_gene <- duplicated(rank_table$gene)
rank_unique <- rank_table[!rank_table$duplicate_gene, , drop = FALSE]
rank_unique <- rank_unique[order(rank_unique$t_statistic, decreasing = TRUE), , drop = FALSE]
ranks <- rank_unique$t_statistic
names(ranks) <- rank_unique$gene

write.table(
  rank_table,
  file.path(output_dir, "BRCA1_T_vs_GFP_GSEA_ranking_all_mapped_rows.tsv"),
  sep = "\t", quote = FALSE, row.names = FALSE, na = ""
)
write.table(
  rank_unique,
  file.path(output_dir, "BRCA1_T_vs_GFP_GSEA_ranking_unique_genes.tsv"),
  sep = "\t", quote = FALSE, row.names = FALSE, na = ""
)

collections <- list(
  BP = list(collection = "C5", subcollection = "GO:BP", label = "Biological process"),
  CC = list(collection = "C5", subcollection = "GO:CC", label = "Cellular component"),
  MF = list(collection = "C5", subcollection = "GO:MF", label = "Molecular function"),
  HALLMARK = list(collection = "H", subcollection = NULL, label = "Hallmark")
)

all_results <- list()
msigdb_versions <- character()

for (i in seq_along(collections)) {
  domain <- names(collections)[i]
  info <- collections[[domain]]
  genesets <- if (is.null(info$subcollection)) {
    msigdbr(species = "Homo sapiens", collection = info$collection)
  } else {
    msigdbr(
      species = "Homo sapiens",
      collection = info$collection,
      subcollection = info$subcollection
    )
  }
  msigdb_versions <- unique(c(msigdb_versions, genesets$db_version))
  descriptions <- unique(genesets[, c("gs_name", "gs_description")])
  pathways <- lapply(split(genesets$gene_symbol, genesets$gs_name), unique)
  set.seed(seed + i)
  fg <- fgseaMultilevel(
    pathways = pathways,
    stats = ranks,
    minSize = min_size,
    maxSize = max_size,
    eps = 0,
    scoreType = "std",
    nPermSimple = 10000
  )
  fg <- as.data.frame(fg)
  fg$leadingEdge <- vapply(
    fg$leadingEdge, paste, collapse = ";", FUN.VALUE = character(1)
  )
  fg$GO_domain <- domain
  fg$GO_domain_label <- info$label
  fg <- merge(
    fg, descriptions,
    by.x = "pathway", by.y = "gs_name",
    all.x = TRUE, sort = FALSE
  )
  fg$Direction <- ifelse(fg$NES >= 0, "BRCA1-T enriched", "GFP enriched")
  fg <- fg[order(fg$padj, -abs(fg$NES)), , drop = FALSE]
  all_results[[domain]] <- fg
  write.table(
    fg,
    file.path(output_dir, paste0("BRCA1_T_vs_GFP_GSEA_", domain, "_all.tsv")),
    sep = "\t", quote = FALSE, row.names = FALSE, na = ""
  )
  write.table(
    fg[!is.na(fg$padj) & fg$padj < fdr_cutoff, , drop = FALSE],
    file.path(output_dir, paste0("BRCA1_T_vs_GFP_GSEA_", domain, "_FDR_lt_0.05.tsv")),
    sep = "\t", quote = FALSE, row.names = FALSE, na = ""
  )
}

gsea_res <- do.call(rbind, all_results)
rownames(gsea_res) <- NULL
gsea_res$minus_log10_FDR <- -log10(pmax(gsea_res$padj, .Machine$double.xmin))
write.table(
  gsea_res,
  file.path(output_dir, "BRCA1_T_vs_GFP_GSEA_all_collections.tsv"),
  sep = "\t", quote = FALSE, row.names = FALSE, na = ""
)

leading_set <- function(x) {
  if (is.na(x) || x == "") character() else unique(strsplit(x, ";", fixed = TRUE)[[1]])
}

jaccard <- function(a, b) {
  if (!length(a) || !length(b)) return(0)
  length(intersect(a, b)) / length(union(a, b))
}

select_representatives <- function(x, n = max_terms_per_direction) {
  x <- x[!is.na(x$padj) & x$padj < fdr_cutoff, , drop = FALSE]
  if (!nrow(x)) return(x)
  x <- x[order(x$padj, -abs(x$NES)), , drop = FALSE]
  selected <- integer()
  selected_sets <- list()
  for (i in seq_len(nrow(x))) {
    current <- leading_set(x$leadingEdge[i])
    overlap <- if (length(selected_sets)) {
      max(vapply(selected_sets, function(z) jaccard(current, z), numeric(1)))
    } else 0
    if (overlap < jaccard_cutoff) {
      selected <- c(selected, i)
      selected_sets[[length(selected_sets) + 1L]] <- current
    }
    if (length(selected) >= n) break
  }
  x[selected, , drop = FALSE]
}

clean_term <- function(pathway, domain) {
  prefix <- if (domain == "HALLMARK") "HALLMARK_" else paste0("GO", domain, "_")
  z <- sub(paste0("^", prefix), "", pathway)
  z <- str_replace_all(z, "_", " ")
  z <- str_to_sentence(str_to_lower(z))
  z <- str_replace_all(z, regex("\\bdna\\b", ignore_case = TRUE), "DNA")
  z <- str_replace_all(z, regex("\\brna\\b", ignore_case = TRUE), "RNA")
  z <- str_replace_all(z, regex("\\batp\\b", ignore_case = TRUE), "ATP")
  z <- str_replace_all(z, regex("\\bnadh\\b", ignore_case = TRUE), "NADH")
  str_replace_all(z, regex("\\bswi[ /]snf\\b", ignore_case = TRUE), "SWI/SNF")
}

make_bubble <- function(domains, labels, title, filename, height_cm = page_cm) {
  x <- gsea_res[gsea_res$GO_domain %in% domains, , drop = FALSE]
  plot_terms <- do.call(
    rbind,
    lapply(split(x, interaction(x$GO_domain, x$Direction, drop = TRUE)), function(z) {
      select_representatives(z)
    })
  )
  if (is.null(plot_terms) || !nrow(plot_terms)) {
    blank <- ggplot() +
      annotate(
        "text", x = 0.5, y = 0.5,
        label = "No gene sets with FDR < 0.05",
        size = geom_text_mm
      ) +
      xlim(0, 1) + ylim(0, 1) +
      labs(title = title) +
      theme_void(base_size = base_pt, base_family = "Helvetica") +
      theme(plot.title = element_text(size = base_pt, face = "bold", hjust = 0.5))
    save_pdf(filename, blank, page_cm, height_cm)
    return(invisible(NULL))
  }
  rownames(plot_terms) <- NULL
  plot_terms$term_label <- mapply(
    clean_term, plot_terms$pathway, plot_terms$GO_domain
  )
  plot_terms$term_label <- str_wrap(plot_terms$term_label, width = 50)
  plot_terms$GO_domain_label <- factor(
    plot_terms$GO_domain_label, levels = labels
  )
  plot_terms$term_key <- paste(
    plot_terms$GO_domain, plot_terms$term_label, sep = "___"
  )
  ordered_keys <- unlist(lapply(domains, function(domain) {
    z <- plot_terms[plot_terms$GO_domain == domain, , drop = FALSE]
    z$term_key[order(z$NES)]
  }), use.names = FALSE)
  plot_terms$term_factor <- factor(
    plot_terms$term_key, levels = unique(ordered_keys)
  )

  bubble <- ggplot(plot_terms, aes(NES, term_factor)) +
    geom_vline(xintercept = 0, color = "#777777", linewidth = 0.2) +
    geom_point(
      aes(size = minus_log10_FDR, fill = NES),
      shape = 21, color = "#333333", stroke = 0.2, alpha = 0.95
    ) +
    facet_grid(
      GO_domain_label ~ ., scales = "free_y", space = "free_y", switch = "y"
    ) +
    scale_y_discrete(labels = function(z) sub("^.*___", "", z)) +
    scale_fill_gradient2(
      low = "#2C73B9", mid = "#F7F7F7", high = "#D85B4B",
      midpoint = 0, name = "NES"
    ) +
    scale_size_continuous(range = c(1.2, 3.5), name = "-log10 FDR") +
    guides(
      fill = guide_colorbar(display = "rectangles"),
      size = guide_legend()
    ) +
    labs(
      title = title,
      subtitle = "Positive NES: BRCA1-T enriched; negative NES: GFP enriched",
      x = "Normalized enrichment score (NES)", y = NULL
    ) +
    theme_bw(base_size = base_pt, base_family = "Helvetica") +
    theme(
      text = element_text(size = base_pt, family = "Helvetica", color = "black"),
      plot.title = element_text(size = base_pt, face = "bold", hjust = 0.5),
      plot.subtitle = element_text(size = base_pt, hjust = 0.5),
      axis.title = element_text(size = base_pt),
      axis.text = element_text(size = base_pt),
      axis.text.y = element_text(size = base_pt, lineheight = 0.85),
      strip.text = element_text(size = base_pt, face = "bold"),
      strip.background = element_rect(fill = "#F1F1F1", color = "#BBBBBB"),
      strip.placement = "outside",
      panel.grid.major.y = element_blank(),
      panel.grid.minor = element_blank(),
      legend.title = element_text(size = base_pt),
      legend.text = element_text(size = base_pt),
      legend.key.height = grid::unit(0.25, "cm")
    )
  save_pdf(filename, bubble, page_cm, height_cm)
  write.table(
    plot_terms,
    sub("\\.pdf$", "_terms.tsv", filename),
    sep = "\t", quote = FALSE, row.names = FALSE, na = ""
  )
}

make_bubble(
  c("BP", "CC"),
  c("Biological process", "Cellular component"),
  "GO BP and CC GSEA of BRCA1-T vs GFP BioID",
  file.path(output_dir, "BRCA1_T_vs_GFP_GO_BP_CC_GSEA_bubble.pdf"),
  height_cm = 10
)
make_bubble(
  c("CC", "HALLMARK"),
  c("Cellular component", "Hallmark"),
  "GO CC and Hallmark GSEA of BRCA1-T vs GFP BioID",
  file.path(output_dir, "BRCA1_T_vs_GFP_GO_CC_Hallmark_GSEA_bubble.pdf"),
  height_cm = 10
)

write.table(
  dat,
  file.path(output_dir, "BRCA1_T_vs_GFP_BioID_analysis_table.tsv"),
  sep = "\t", quote = FALSE, row.names = FALSE, na = ""
)
write.table(
  cbind(
    protein_id = heat_dat$protein_id,
    gene = rownames(heat_mat_raw),
    direction = as.character(heat_dat$status),
    heat_mat_raw
  ),
  file.path(output_dir, "BRCA1_T_vs_GFP_heatmap_selected_matrix.tsv"),
  sep = "\t", quote = FALSE, row.names = FALSE, na = ""
)

sig_counts <- aggregate(
  pathway ~ GO_domain_label + Direction,
  data = gsea_res[!is.na(gsea_res$padj) & gsea_res$padj < fdr_cutoff, , drop = FALSE],
  FUN = length
)
names(sig_counts)[3] <- "FDR_lt_0.05_terms"
write.table(
  sig_counts,
  file.path(output_dir, "BRCA1_T_vs_GFP_GSEA_significant_term_counts.tsv"),
  sep = "\t", quote = FALSE, row.names = FALSE
)

summary_lines <- c(
  "BRCA1-T versus GFP BioID analysis",
  paste0("Run date: ", Sys.Date()),
  paste0("Input: ", input_file),
  "Design: 4 BRCA1-T BioID replicates versus 4 GFP BioID replicates",
  paste0("Protein rows: ", nrow(dat)),
  paste0("Complete eight-sample LFQ rows: ", sum(complete.cases(sample_mat_all))),
  paste0("Perseus q < 0.05 rows: ", sum(dat$q_value < fdr_cutoff, na.rm = TRUE)),
  paste0(
    "BRCA1-T enriched (q < 0.05, log2 FC >= 0.5): ",
    sum(dat$status == "BRCA1-T enriched")
  ),
  paste0(
    "GFP enriched (q < 0.05, log2 FC <= -0.5): ",
    sum(dat$status == "GFP enriched")
  ),
  paste0("Heatmap proteins: ", length(heat_ix)),
  paste0("Unique ranked gene symbols: ", nrow(rank_unique)),
  paste0("MSigDB version: ", paste(msigdb_versions, collapse = ", ")),
  "GSEA collections: C5 GO:BP, GO:CC, GO:MF and H Hallmark",
  paste0("GSEA gene-set size filter: ", min_size, "-", max_size),
  paste0("GSEA significance threshold: FDR < ", fdr_cutoff),
  paste0("Representative terms per collection and direction: up to ", max_terms_per_direction),
  paste0("Leading-edge Jaccard redundancy cutoff: ", jaccard_cutoff),
  "",
  "Significant GSEA term counts:",
  capture.output(print(sig_counts, row.names = FALSE))
)
writeLines(
  summary_lines,
  file.path(output_dir, "BRCA1_T_vs_GFP_BioID_analysis_summary.txt")
)
capture.output(sessionInfo(), file = file.path(output_dir, "R_sessionInfo.txt"))

cat("BRCA1-T versus GFP BioID analysis completed in:", output_dir, "\n")
```
