## Supplementary material for "An integrated, transcript-resolution atlas of prostate cancer progression unmasks associations with oncogenic pathways": Reproducible code: configure_paths.py

```
#!/usr/bin/env python3
import argparse
from pathlib import Path
parser = argparse.ArgumentParser(description='Replace repository path placeholders in a working copy of the code.')
parser.add_argument('documents_root', help='Local equivalent of the original Documents analysis root')
parser.add_argument('--project-root', default=str(Path(__file__).resolve().parent), help='Root for project-local inputs and outputs; defaults to this package')
parser.add_argument('--desktop-root', default='', help='Optional replacement for legacy Desktop-only inputs')
parser.add_argument('--downloads-root', default='', help='Optional replacement for legacy Downloads-only inputs')
args = parser.parse_args()
root = Path(__file__).resolve().parent
for path in (root/'code').rglob('*'):
    if not path.is_file():
        continue
    text = path.read_text(encoding='utf-8', errors='replace')
    updated = text.replace('__PCATLAS_DOCUMENTS_ROOT__', args.documents_root)
    updated = updated.replace('__PCATLAS_PROJECT_ROOT__', args.project_root)
    if args.desktop_root:
        updated = updated.replace('__USER_DESKTOP__', args.desktop_root)
    if args.downloads_root:
        updated = updated.replace('__USER_DOWNLOADS__', args.downloads_root)
    if updated != text:
        path.write_text(updated, encoding='utf-8')
print('Configured code paths under', root/'code')
```
