## Supplementary material for "An integrated, transcript-resolution atlas of prostate cancer progression unmasks associations with oncogenic pathways": Reproducible code: README.txt

Package versions should be taken from archived sessionInfo files when available. The historical scripts span multiple environments; a single current-machine lockfile would be misleading until clean-environment reruns are completed.
