## Supplementary material for "An integrated, transcript-resolution atlas of prostate cancer progression unmasks associations with oncogenic pathways": Reproducible code: alphafold_info.txt

In silico structural modelling. Full-length human TRIM11 (UniProt Q96F44, 468 aa) and full-length human androgen receptor (UniProt P10275) were submitted as a single two-chain job at 1:1 stoichiometry to the AlphaFold Server (https://alphafoldserver.com), which runs AlphaFold 3 (Abramson et al., Nature 630:493–500, 2024). Prediction used model version AlphaFold-beta-20231127 with server default settings: automatically generated multiple-sequence alignments, 10 recycling iterations, five returned models per job, and no seed specification available to the user. No ligands, metal ions, nucleic acids or post-translational modifications were included. The model shown is the top-ranked of the five selected by the server ranking score

I calculated the parameters only for the part of the model displayed into the picture.
