## Supplementary material for "An integrated, transcript-resolution atlas of prostate cancer progression unmasks associations with oncogenic pathways": Reproducible code: Fig5_FigS10_panel_revision_record_2026-09-04.md

# Figure 5 and Figure S10 panel revision record

Date first recorded: 2026-09-04  
Latest panel update: 2026-09-07

## Purpose and authority

This file records the author-directed reassignment of Figure 5 and Figure S10 panels and the provenance of two new Figure 5 panels. It is the panel-mapping authority for the next update of the manuscript, Figure Legends, Methods, supplementary tables, and reproducible-code package.

The **2026-09-07 Figure S10 mapping supersedes the 2026-09-04 Figure S10 mapping** recorded earlier in this file. The Figure 5 mapping is unchanged by the 2026-09-07 update.

The current Word files were not modified during this recording step. Their pre-revision panel descriptions remain in:

- `${PROJECT_ROOT}/Figure_legends_Cell_Press_code_completed_v6.0.docx`
- `${PROJECT_ROOT}/Methods_Cell_Press_code_completed_v6.0.docx`

## Author-directed panel changes

| Previous location | New location | Content |
|---|---|---|
| Figure 5H | Figure S10H | Atlas-only Hallmark GSEA comparing BRCA1-T- and BRCA1-C-associated expression signatures; moved through temporary S10J in the 2026-09-04 layout |
| Figure 5I | Figure 5H | BRCA1 and HA-tagged BRCA1-T/Delta11q immunoblot in LNCaP cells |
| Figure 5J | Figure 5I | Representative anti-HA immunofluorescence of BRCA1-T/Delta11q-expressing LNCaP cells |
| Figure 5K | Figure 5J | Concordance between atlas BRCA1-T-associated gene-expression changes and experimental BRCA1-T/Delta11q overexpression |
| Figure S10J | Figure 5K | Concordance between experimental BRCA1-T overexpression and atlas BRCA1-C-associated gene-expression changes |
| New panel | Figure 5L | BRCA1 GO Biological Process bipartite network integrating atlas, BRCA1-T RNA-seq, and BioID evidence |
| New panel | Figure 5M | BRCA1-T versus GFP BioID differential-proximity volcano plot |
| Figure 5L | Figure 5N | CRPC overall-survival analysis stratified by BRCA1-C expression |
| Figure 5M | Figure 5O | CRPC overall-survival analysis stratified by BRCA1-T expression |
| Original Figure S10H | Removed | Atlas-only GO cellular-component GSEA; retain underlying data/code but remove the panel citation |
| Figure S10I | Figure S10I | Experimental BRCA1-T/Delta11q-overexpression Hallmark GSEA remains in place |
| Figure S10K | Figure S10J | Primary-prostate-cancer survival stratified by BRCA1-C expression |
| Figure S10L | Figure S10K | Primary-prostate-cancer survival stratified by BRCA1-T expression |

## Revised Figure 5 panel order

- **A-G:** unchanged from Figure Legends v6.0.
- **H:** previous Figure 5I, BRCA1/HA-BRCA1-T immunoblot.
- **I:** previous Figure 5J, BRCA1-T immunofluorescence.
- **J:** previous Figure 5K, atlas BRCA1-T versus BRCA1-T-overexpression concordance.
- **K:** previous Figure S10J, experimental BRCA1-T overexpression versus atlas BRCA1-C concordance.
- **L:** new BRCA1 GO:BP bipartite network.
- **M:** new BRCA1-T versus GFP BioID volcano plot.
- **N:** previous Figure 5L, BRCA1-C CRPC overall survival.
- **O:** previous Figure 5M, BRCA1-T CRPC overall survival.

Accordingly, future grouped references to the CRPC survival panels must change from **Figure 5L-M** to **Figure 5N-O**.

## Revised Figure S10 panel order

- **A-G:** unchanged from Figure Legends v6.0.
- **H:** previous revised Figure S10J, originally Figure 5H: atlas-only Hallmark GSEA comparing BRCA1-T- and BRCA1-C-associated expression signatures.
- **I:** unchanged original Figure S10I: Hallmark GSEA comparing experimental BRCA1-T/Delta11q overexpression with empty-vector control.
- **J:** previous Figure S10K: primary-prostate-cancer survival stratified by BRCA1-C expression.
- **K:** previous Figure S10L: primary-prostate-cancer survival stratified by BRCA1-T expression.

The original Figure S10H GO cellular-component GSEA panel has been removed from Figure S10. Its underlying analysis files may remain in the supplementary-data and reproducibility archives but should no longer be cited as a displayed figure panel.

The original Figure S10J concordance panel is no longer present in Figure S10 because it became Figure 5K in the 2026-09-04 revision. The temporary replacement Figure S10J from that revision has now moved to Figure S10H.

### Figure S10 revision history

| Revision date | Change |
|---|---|
| 2026-09-04 | Original Figure 5H moved to Figure S10J; original Figure S10J moved to Figure 5K. |
| 2026-09-07 | Figure S10H removed; the temporary Figure S10J moved to H; Figure S10K and L moved forward to J and K. Original Figure S10I remained I. |

## New Figure 5L provenance

### Final artwork

`${PROJECT_ROOT}/BRCA1_GO_BP_bipartite_network_v1/BRCA1_GO_BP_bipartite_network_v5_compact_8x10cm.pdf`

### Final plotting script and script chain

1. `${PROJECT_ROOT}/BRCA1_GO_BP_bipartite_network_v1/plot_BRCA1_GO_BP_bipartite_network_v5_compact.R`
2. The v5 script sources `plot_BRCA1_GO_BP_bipartite_network_v4_final.R`.
3. The v4 script sources `plot_BRCA1_GO_BP_bipartite_network.R`, which performs the audited data preparation.

### Direct inputs

- Pathway evidence table: `${PROJECT_ROOT}/BRCA1_GO_BP_joint_continuous_no_covariates/BRCA1_GO_BP_expanded_RNAseq_FDR0.10_with_significant_nonbait_BioID_proteins.tsv`
- BioID protein annotation table: `${PROJECT_ROOT}/bioID_BRCA/BRCA1_T_cytoplasmic_reanalysis/BRCA1_T_significant_protein_localization.tsv`
- MSigDB membership source: pinned MSigDB 2026.1.Hs C5 GO:BP cache used by the BRCA1 GO:BP analysis.

### Upstream generation scripts

- `BRCA1_GO_BP_joint_continuous_no_covariates/run_BRCA1_GO_BP_joint_continuous.R`
- `BRCA1_GO_BP_joint_continuous_no_covariates/filter_four_way_T_supported_GO_BP.R`
- `BRCA1_GO_BP_joint_continuous_no_covariates/make_expanded_significant_BioID_protein_table.R`
- `bioID_BRCA/BRCA1_T_vs_GFP_BioID_analysis/run_BRCA1_T_vs_GFP_BioID_analysis.R`

### Selection rules encoded in the source

- Atlas gene-level models use all 1,365 samples and fit `gene count ~ BRCA1_C_z + BRCA1_T_z`, without additional covariates.
- `BRCA1_C_z` and `BRCA1_T_z` are z scores of transcript-level `log2(CPM + 1)` expression.
- GO:BP GSEA uses `fgseaMultilevel`, MSigDB 2026.1.Hs C5 GO:BP, `minSize = 15`, no upper gene-set-size limit, and `eps = 0`.
- Network pathway inclusion requires Atlas-T FDR < 0.10, Atlas-C FDR < 0.10, experimental BRCA1-T-overexpression RNA-seq FDR < 0.10, an RNA-seq NES direction matching Atlas-T, and at least one significant non-bait BRCA1-T-enriched BioID protein mapped to the pathway.
- A significant BRCA1-T-enriched BioID protein is defined as Perseus q < 0.05 and log2 fold change >= 0.5 for BRCA1-T versus GFP.
- The final compact network contains 19 pathways, 14 non-bait proteins, and 38 GO-membership edges. BRCA1 bait itself is excluded from the protein-node set.

### Visual encodings

- Three evidence columns show Atlas-T, Atlas-C, and experimental BRCA1-T RNA-seq.
- Evidence-node fill denotes NES; evidence-node size denotes `-log10(FDR)`.
- Protein-node fill denotes BRCA1-T-versus-GFP BioID log2 fold change; protein-node size denotes `-log10(q)`.
- Edges denote GO:BP membership of the displayed BioID proteins.
- Pathways are arranged in six modules: metabolism and transport, immune and inflammatory response, programmed cell death, oxidative stress and mitochondria, protein homeostasis, and cell proliferation.
- The final PDF is 8 x 10 cm, uses 5-point Helvetica, and was written with the standard R PDF device using `useDingbats = FALSE` and `compress = FALSE`.

### Legend seed for a future revision

**(L)** Bipartite network linking GO Biological Process terms supported by BRCA1 isoform-associated atlas expression, BRCA1-T-overexpression RNA-seq, and BRCA1-T-versus-GFP BioID to significant non-bait BRCA1-T-enriched proximity-labeled proteins. The three evidence columns represent Atlas-T, Atlas-C, and experimental BRCA1-T RNA-seq; fill denotes NES and size denotes -log10(FDR). Protein-node fill denotes BioID log2 fold change and protein-node size denotes -log10(q value); edges indicate GO:BP membership. Pathways are grouped into the indicated biological-process modules.

Interpret this network as integrated association and proximity-labeling evidence, not as proof of direct protein interaction, pathway activation, or causality.

## New Figure 5M provenance

### Final artwork

`${PROJECT_ROOT}/bioID_BRCA/BRCA1_T_vs_GFP_BioID_analysis/BRCA1_T_vs_GFP_BioID_volcano_9x6cm.pdf`

The duplicated filename text supplied in the panel note is interpreted as the existing final file above.

### Scripts

- Full analysis: `${PROJECT_ROOT}/bioID_BRCA/BRCA1_T_vs_GFP_BioID_analysis/run_BRCA1_T_vs_GFP_BioID_analysis.R`
- Final 9 x 6 cm redraw: `${PROJECT_ROOT}/bioID_BRCA/BRCA1_T_vs_GFP_BioID_analysis/redraw_BRCA1_T_vs_GFP_BioID_volcano_9x6cm.R`

### Inputs and archived outputs

- Perseus-exported input: `${PROJECT_ROOT}/bioID_BRCA/t_test_BRCA-BioID_vs_GFP-BioID_BH5FDR.txt`
- Frozen analysis table used by the redraw: `${PROJECT_ROOT}/bioID_BRCA/BRCA1_T_vs_GFP_BioID_analysis/BRCA1_T_vs_GFP_BioID_analysis_table.tsv`
- Analysis summary: `${PROJECT_ROOT}/bioID_BRCA/BRCA1_T_vs_GFP_BioID_analysis/BRCA1_T_vs_GFP_BioID_analysis_summary.txt`
- R environment: `${PROJECT_ROOT}/bioID_BRCA/BRCA1_T_vs_GFP_BioID_analysis/R_sessionInfo.txt`

### Design and statistics encoded in the source

- Four BRCA1-T BioID replicates were compared with four GFP BioID replicates.
- The analysis contains 1,174 protein rows with complete eight-sample LFQ values.
- The plot uses continuous two-group Student t-test p values, Perseus-exported Benjamini-Hochberg q values, and the BRCA1-T-minus-GFP log2 LFQ difference.
- Proteins are classified as significant at q < 0.05 and absolute log2 fold change >= 0.5.
- Positive values denote enrichment toward BRCA1-T; negative values denote enrichment toward GFP.
- The archived analysis reports 36 BRCA1-T-enriched and 5 GFP-enriched proteins at these joint thresholds.
- The final redraw preserves the archived statistics and classifications; it does not recalculate them.
- The final PDF is 9 x 6 cm, uses 5-point Helvetica, and was written with the standard R PDF device using `useDingbats = FALSE` and `compress = FALSE`.

### Legend seed for a future revision

**(M)** Volcano plot of differential proximity labeling between BRCA1-T BioID and GFP BioID controls across four replicate samples per group. The x axis shows the BRCA1-T-minus-GFP log2 LFQ difference and the y axis shows -log10(Student t-test p value). Proteins meeting Benjamini-Hochberg q < 0.05 and absolute log2 fold change >= 0.5 are colored by enrichment direction; selected proteins are labeled.

BioID measurements are processed LFQ values exported from Perseus. Enrichment indicates differential proximity labeling and should not be described as a direct physical interaction without orthogonal validation.

## Required downstream updates

When the next versions are generated, update all of the following together:

1. Figure 5 and Figure S10 legends, including all panel letters and grouped panel references.
2. Methods sections covering BRCA1 perturbation RNA-seq, BioID differential-proximity analysis, GO:BP GSEA, the continuous two-isoform atlas model, and the bipartite-network construction.
3. Main-text citations to Figure 5H-O and Figure S10H-K.
4. Supplementary-table/source-data indexes for the network node, edge, evidence, BioID protein-level, and volcano-analysis tables.
5. `reproducible_code_submission_v1.0/manifest/panel_code_map.csv`, code and input manifests, README coverage statements, and packaged scripts/data.
6. Any figure-provenance summary that still assigns old Figure 5H-M, Figure S10H, or Figure S10J-L content.
7. Final exported Figure 5 and Figure S10 PDFs after the revised `.ai` artwork is frozen.

Do not describe BioID as missing in future manuscript or provenance versions: the BRCA1-T-versus-GFP BioID analysis, source table, figure code, and downstream GO:BP integration are now present. Any remaining BioID gap should be stated narrowly and only if specific raw-instrument files or experimental metadata are still unavailable.
