## Supplementary material for "An integrated, transcript-resolution atlas of prostate cancer progression unmasks associations with oncogenic pathways": Reproducible code: README_revision.txt

Input table: ${DOCUMENTS_ROOT}/Data_Harmanization/pc_tx_info_filter_5pct_JP.xlsx (value-preserving CSV export)
Initial highlight rule: first 21 rows; is_canonical=1 red, is_canonical=0 blue.
Revision rule: retain an initial blue point only if the complete table contains a canonical transcript with the same gene_name; highlight all matched canonical rows.
Initial blue genes: AR-V7, TP53, TP63
Matched genes: TP53
Removed blue genes: AR-V7, TP63
Point layers are rasterized at 300 dpi with ggrastr; text remains vector.
Final device size: width 9 cm x height 4.5 cm; all text is 5 pt.
PPP2R5C highlighted points were removed; TP63 is labeled ΔNp63.
TP63 canonical point ENST00000264731.8 was added from pc_tx_info_unfiltered.xlsx; its rank was inserted without shifting existing filtered-table ranks.
