## Supplementary material for "An integrated, transcript-resolution atlas of prostate cancer progression unmasks associations with oncogenic pathways": Reproducible code: rnaseq_info.txt

 
RNA sequencing for all experiments involving the samples was performed at the Institute of Oncology Research using NEBNext Ultraexpress RNA Library Prep Kit for Illumina (E3330S - New England BioLabs Inc.) was employed with the NEBNext Multiplex Oligos for Illumina (New England BioLabs Inc.) and NEBNext® Poly(A) mRNA Magnetic Isolation Module for cDNA synthesis (E7490 -  New England BioLabs Inc.) with the addition of barcode sequences and sequenced on the SURFSeq 5000 platform (Genemind) with single-end, 120 base pair long reads.
 
Before the library prep the samples were tested for QC using Bioanalyzer 2100 (Agilent)
