## Supplementary material for "An integrated, transcript-resolution atlas of prostate cancer progression unmasks associations with oncogenic pathways": Reproducible code: STRUCTURE_MODELING_SCOPE.md

# Structure modeling scope

Internal exploratory docking notes are excluded from the public submission package because they were not used as evidence for the final manuscript claims. The reported protein structures are qualitative AlphaFold-derived visualizations. The authoritative software, model, interface-display and interpretation details are provided in Methods v7.2.
