## Supplementary material for "An integrated, transcript-resolution atlas of prostate cancer progression unmasks associations with oncogenic pathways": Reproducible code: README.md

# Reproducible code package — submission version 1.3

Generated: 2026-09-14

This repository-ready package organizes the recovered analysis code and compact machine-readable source data for the manuscript **An integrated, transcript-resolution atlas of prostate cancer progression unmasks associations with oncogenic pathways**. The authoritative textual descriptions are Methods v7.3, Figure Legends v7.2 and Manuscript v1.5 in the final submission package.

## What this package is

- An evidence-preserving archive of 59 located analysis scripts, grouped by analysis stage.
- A panel-level code map suitable for a public repository README, with explicit coverage labels.
- Compact source-data copies for analyses that can be audited without redistributing the large atlas matrices or controlled clinical data.
- Exact or deterministic recovery outputs for FIRRE/FENDRR LASSO, RNF43 Figure 4H, survival endpoint checks, BRCA1 concordance, AWSI networks, the Figure 5L GO:BP network, Figure 5M BioID analysis and reviewer robustness analyses.

## What this package is not

- It is not yet a single-command workflow. Many recovered historical scripts depend on serialized R objects and large matrices that are not redistributed here. Scientific logic is preserved, while machine-specific paths are tokenized in packaged copies; `manifest/code_manifest.csv` records absolute-path occurrences in the originals.
- It does not include raw FASTQ/BAM files or controlled clinical data.
- “Source-linked” means a script and its role were matched confidently, not that the complete panel has been rerun in a clean environment.

## Recommended order

Before running historical scripts in a working copy, execute `python3 configure_paths.py /path/to/Documents --project-root /path/to/project` to replace the repository placeholders with local roots. The manifest retains SHA-256 hashes of the untouched originals for provenance.

1. `01_atlas_core`: atlas correction, PCA, heatmaps, clustering metrics and trajectory.
2. `02_lncrna_motif`: lncRNA analyses and motif/LASSO models.
3. `03_isoform_switch`: Kendall analysis and pathway-associated switch detection.
4. `04_gene_specific_atlas`: gene-specific correlations, AWSI and networks.
5. `05_survival`: survival generation and endpoint audit.
6. `06_long_read`: long-/short-read concordance.
7. `07_brca1_experiment`: BRCA1 perturbation DE/GSEA and atlas concordance.
8. `08_rnf43_reporter`: exact Figure 4H reproduction.
9. `09_structures`: transcript/domain plotting; AlphaFold/ChimeraX provenance is documented separately.
10. `10_audits`: deterministic recovery and reviewer-facing robustness analyses.
11. `11_brca1_bioid_go`: BRCA1-T BioID statistics and integrated GO biological-process network.

## Current reproducibility boundary

- **Exact/deterministic:** FIRRE/FENDRR LASSO coefficients; RNF43 TOPFlash source-data plot and five Šidák-adjusted comparisons; core isoform-switch summaries; BRCA1 experiment-atlas joins; selected long-read correlation summaries; AWSI network exports; Figure 5L integrated GO:BP network; Figure 5M BioID volcano plot from the supplied protein-level table.
- **Source-linked but not clean-room rerun:** several final PCA, heatmap, gene-specific atlas and structure-display branches.
- **Upstream experimental boundary:** BioID protein-level statistics and the downstream analysis code are included. The digestion, LC–MS/MS, MaxQuant and Perseus parameters are documented in Methods v7.3, but raw spectra and search/quantification output are intentionally deferred for this initial round. Wet-lab image panels are outside the computational-code package.

## Critical scientific caveats

- The 41-gene presentation set is recoverable as the union of 34 prespecified candidates and the ten genes with the largest representative delta correlations (three genes overlap).
- Survival endpoints are fixed as progression-free survival for primary tumors and overall survival for CRPC.
- AlphaFold structures were used for qualitative visualization in UCSF ChimeraX v1.12; these panels are not quantitative validation of structural interfaces.
- Newly generated short-read and long-read RNA-seq data have been submitted under private BioProject PRJNA1519081; reviewer access will be generated through NCBI during peer review.
- This versioned package is suitable for editorial and peer review. A permanent public repository URL/DOI will be assigned no later than publication. Processed BRCA1 BioID protein-level results are supplied; raw mass-spectrometry files are intentionally deferred for this initial round and will be supplied upon request and deposited in ProteomeXchange/PRIDE before publication.
- Software versions in archived `sessionInfo.txt` files are preferred. Versions inferred from the current machine must not be presented as historical analysis versions.

See `manifest/code_manifest.csv`, `manifest/input_manifest.csv`, and `manifest/panel_code_map.csv` for the audited details.

## Public-code language normalization

Version 1.3 uses English-only script filenames and executable text. Legacy non-English comments and console messages were removed or normalized without changing analytical expressions, thresholds, statistical tests, random seeds, object names or data values. The code manifest retains the archived source checksum and separately records the checksum of each public copy.
