## Supplementary material for "An integrated, transcript-resolution atlas of prostate cancer progression unmasks associations with oncogenic pathways": Reproducible code: VALIDATION_REPORT.md

# Validation report — version 1.3

Validated on 2026-09-14.

- Manifest coverage: 59/59 listed analysis scripts located; 28/28 listed source-data inputs located.
- Syntax checks: 54 R scripts parsed successfully with `Rscript`; 7 Python files parsed successfully with Python `ast`.
- Portability scan: no unresolved machine-specific user-home paths were found outside the preserved Word documentation. Packaged script paths use repository placeholders.
- Integrity: source files recorded in the manifests include size, modification time and SHA-256 checksum where applicable.
- Figure map: updated for Figure 5A–O, Figure S10A–K, the Figure S2/S3 exchange, and the new Figure 5L/M analyses.
- Reproducibility boundary: figure-generating computational analyses are code-linked; panels labelled `exact-rerun` or `deterministic-core` include both code and compact source data. Historical atlas display branches labelled `source-linked` or `partial-final-branch` may require the original large matrices or serialized objects.

Short-read and long-read RNA-sequencing data have been submitted under private BioProject PRJNA1519081; reviewer access will be generated through NCBI during peer review. The remaining publication-stage release dependencies are a permanent public code/processed-data repository URL or DOI and deposition of the raw BRCA1 BioID mass-spectrometry files plus search/quantification output in ProteomeXchange/PRIDE. BioID digestion, LC–MS/MS, MaxQuant and Perseus parameters are documented in Methods v7.3.

- Public-language audit: script filenames and executable text contain no Chinese characters; legacy source and public-copy checksums are both recorded.
